# Hyperactive neuronal networks enhance tau spread in an Alzheimer’s disease mouse model

**DOI:** 10.1101/2024.12.01.625514

**Authors:** Aaron J. Barbour, Keegan Hoag, Eli J. Cornblath, Abigail Chavez, Alfredo Lucas, Xiaofan Li, Sydney Zebrowitz, Chloe Hassman, Omar Vazquez, Sharon X. Xie, Edward B. Lee, Kathryn A. Davis, Virginia M.Y. Lee, Delia M. Talos, Frances E. Jensen

**Affiliations:** Department of Neurology, Perelman School of Medicine, University of Pennsylvania; Philadelphia, PA 19104, USA; Department of Biostatistics, Epidemiology, and Informatics, Perelman School of Medicine, University of Pennsylvania; Philadelphia, PA 19104, USA; Translational Neuropathology Research Laboratory, Perelman School of Medicine, University of Pennsylvania; Philadelphia, PA 19104, USA; Department of Pathology and Laboratory Medicine, Perelman School of Medicine, University of Pennsylvania; Philadelphia, PA 19104, USA; Institute on Aging, Perelman School of Medicine, University of Pennsylvania; Philadelphia, PA 19104, USA; Center for Neurodegenerative Disease Research, Perelman School of Medicine, University of Pennsylvania; Philadelphia, PA 19104, USA

**Author notes:** Corresponding authors: Frances E. Jensen, MD, FACP Arthur Knight Asbury MD Professor Chair, Department of Neurology, Hospital of the University of Pennsylvania, 3400 Spruce Street, Dulles 3 Philadelphia, PA 19104, PA USA, Delia Talos, MD, 415 Curie Boulevard, 263 Clinical Research Building Philadelphia, PA, 19104.

## Abstract

Pathological tau spreads via neuronal connections in Alzheimer’s disease (AD). Given the high incidence and deleterious consequences of epileptiform activity in AD, we hypothesized that neuronal hyperactivity and seizures exacerbate tau spread. To examine the impacts of brain-wide network and population hyperactivity on tau spread, we created a novel mouse model involving the cross of targeted recombination in active populations (TRAP) and the 5 times familial AD mice (5X-TRAP) that allows for the permanent labelling of seizure-activated neurons. To explore the effects of seizures on tau spread, we injected these mice with human AD brain-derived tau to induce pathological tau spread, and induced seizures with pentylenetetrazol (PTZ) kindling. Brain mapping revealed that seizures increased tau spread in 5X-TRAP mice, which correlated extensively with memory deficits in PTZ kindled 5X-TRAP mice. Using computational models, we found data supportive of increased anterograde tau spread in 5X-TRAP mice and that regional neuronal activity levels were predictive of tau pathology. On a cellular level, we found that hyperactive neurons drive elevated tau propagation in 5X-TRAP mice. We also found corroborating evidence of increased tau spread in AD patients with a seizure history compared to those without. Our study identifies neuronal hyperactivity and seizures as key, targetable factors underlying AD progression.

## Introduction

The spread of the pathological tau protein is central to the progression of Alzheimer’s disease (AD), and proceeds along functional and neuroanatomical connections (1–6), with pathological tau burden closely correlating with cognitive decline (7–10). Tau pathology in AD begins in the locus coeruleus, and spreads to the entorhinal cortex, followed by the limbic system and neocortex (10–12). In mice, intracerebral injection of human AD brain-derived tau lysate (AD-tau seeding) results in misfolding and propagation of endogenous tau throughout the brain along neuroanatomical connections (13), as seen in human AD. Tau spread is worsened by β– amyloid (Aβ) pathology in humans (1, 14) and the five times familial AD (5XFAD) and other amyloidogenic rodent models (15, 16). Studies have shown that neuronal activity can induce synaptic tau release *in vitro* and can worsen local tau pathology *in vivo* (17), but direct evidence that neuronal hyperactivity causes pathological tau spread throughout the brain *in vivo* is still missing. A number of studies in mice and humans have demonstrated that the progressive spread of tau pathology throughout the brain can be recapitulated in silico by computational models of tau spread along structural connections between brain regions (18). Indeed, these models provide quantitative evidence that tau spreads in both retrograde and anterograde directions in seeded mice (19) and humans (1, 6, 20, 21). Understanding the factors that drive both network and cellular level tau spread throughout the brain is poised to yield novel therapeutic targets and strategies to slow disease progression.

Neuronal network hyperexcitability is prevalent in AD, particularly at early stages (22), with unprovoked seizures occurring in up to 22% of AD patients (23). We and others have found worsened AD pathology and cognitive outcomes in AD patients with a seizure history (23–28), and AD-like pathology in temporal lobe epilepsy (29), suggestive of a bidirectional relationship between network hyperactivity and AD. Similarly, neuronal hyperactivity is found in 5XFAD mice, beginning at early, prodromal stages (24, 28) and Aβ-producing mouse models also show neuronal hyperactivity at the cellular level (28, 30), with a positive feedback loop occurring between neuronal hyperactivity and Aβ pathology (31, 32). Furthermore, seizure induction with pentylenetetrazol (PTZ) kindling early in the disease progression worsens the progression of amyloid pathology, excitatory: inhibitory imbalance, and cognitive deficits (24, 28, 33). While network to cellular level neuronal hyperexcitability are well-characterized phenomena in AD, the role of neuronal hyperactivity in the progression of tau spread is not well understood.

Recently developed genetic tools enable the permanent fluorescent (tdTomato, tdT) labeling and tracking of transiently activated neuronal populations (Targeted Recombination in Active Populations; TRAP), using the immediate early gene cFos as a proxy marker for neuronal activity (34, 35). In murine epilepsy models, we and others have previously demonstrated that neurons that underwent activity dependent tdT labelling during seizures have long-lasting hyperexcitability, synaptic alterations, and demonstrate unique pathological effects (36–38) that are promising therapeutic targets. However, the TRAP tool has not been used to date to investigate hyperexcitability in neurodegeneration. Here, we created a novel model by crossing TRAP mice with 5XFAD (5X-TRAP/WT-TRAP) to examine the interactions between tau spread and neuronal hyperactivity on both a network and cellular level and track the contributions of hyperactive populations to subsequent tau spread.

We hypothesized that neuronal hyperactivity in 5X-TRAP mice would be associated with increased anterograde tau spread along neuronal networks and would be exacerbated by seizure induction. To establish a causal role of seizures and neuronal hyperactivity in tau propagation, we seeded WT-TRAP and 5X-TRAP mice with AD-tau in the right hippocampus and overlying cortex to induce tau spread, induced seizures with PTZ kindling, and labelled seizure-activated neurons with tdT. Using comprehensive brain mapping and computational models of tau spread throughout neuroanatomical networks, we found that PTZ kindling in early disease stages resulted in exacerbation of tau spread in 5X-TRAP mice through seizure-aactivated circuits, and that network hyperactivity was associated with increased anterograde tau propagation. Memory deficits in PTZ kindled 5X-TRAP mice were highly correlated with tau spread and neuronal hyperactivity. Further, we found that hyperactive tdT+ neurons specifically drove increased tau spread in 5X-TRAP mice. To increase translational relevance, we examined the extent of tau pathology in human AD brain tissue and found evidence suggestive of increased tau spread in AD patients with a clinical seizure history compared to those without. Overall, these studies causally demonstrate that neuronal hyperactivity, via seizure induction, drives tau spread and suggest that hyperactive networks and neurons promote anterograde tau spread in AD. Thus, seizures and neuronal hyperexcitability are mitigatable and translatable targets to slow tau spread and AD progression.

## Results

### Novel 5X-TRAP mouse to study the interactions between AD pathology and neuronal activity

As our prior results revealed that seizure induction in 5XFAD mice early in the disease progression resulted in exacerbation of later memory deficits and brain pathology, we aimed to further study the interactions between neurodegeneration and neuronal activity at a population and network level. We thus developed a novel model by crossing hemizygous 5XFAD mice with mice homozygous for Fos^2A-iCreER^ (TRAP2) and B6.Cg-*Gt(ROSA)26Sor*^tm14(CAG-tdTomato)Hze^/J (Ai14) (Fig 1A). The resulting 5X-TRAP and WT-TRAP mice allow for robust, permanent tdT expression in neurons that express the activity-dependent cFos protein during a 4–6-hour window after 4-hydroxytamoxifen (4-OHT) induced CreER expression (Fig 1A). This novel 5X-TRAP model allows for the examination of the interactions between individual and brain-wide neuronal activity patterns and AD pathology.

**Figure 1.**
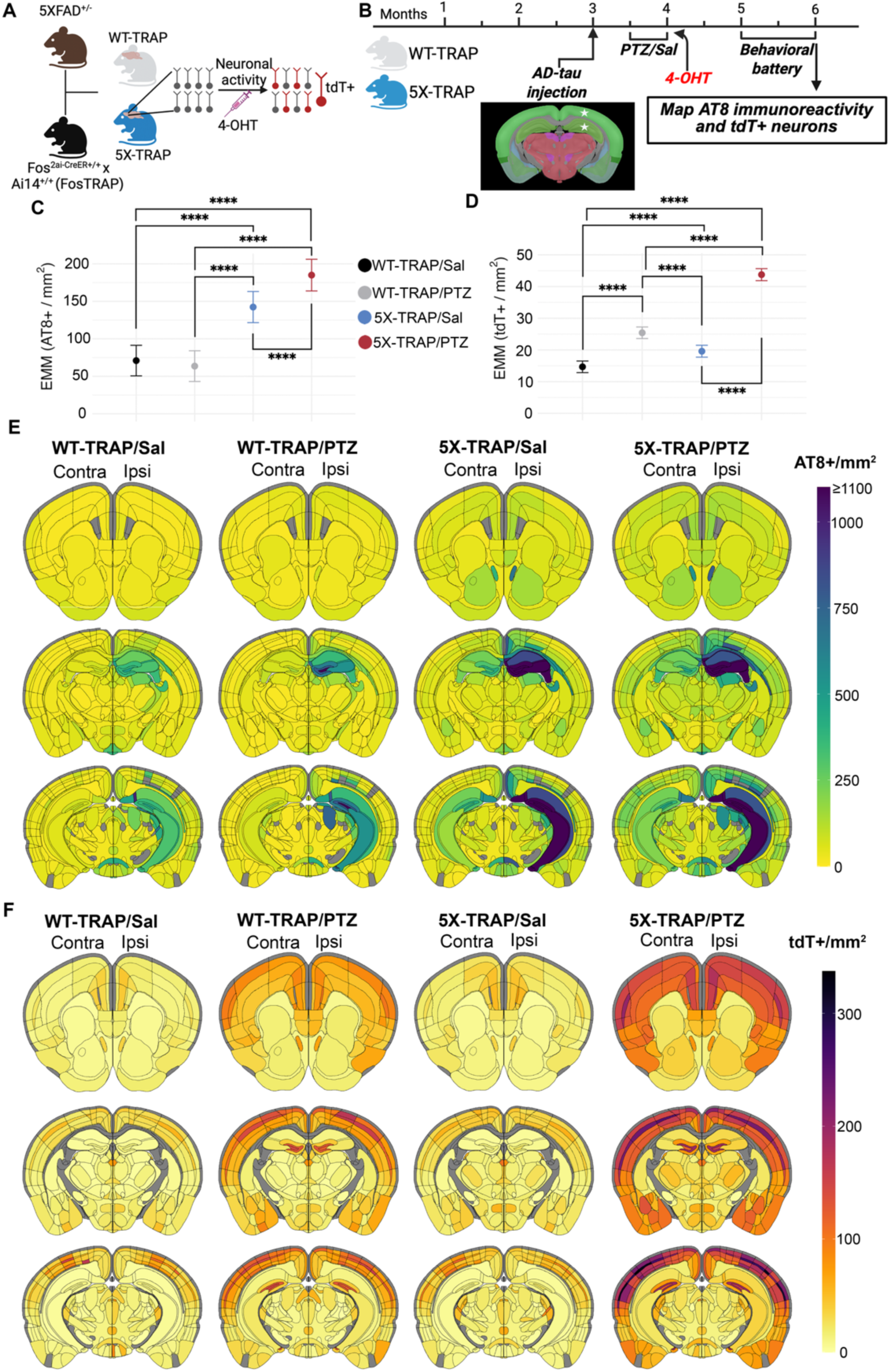
Brain mapping reveals increased tau spread in 5X-TRAP mice that is exacerbated by seizure induction and increased tdT+ labeling in PTZ and saline treated 5X-TRAP mice. (**A**) Schematic for the generation of, and activity-dependent labelling in WT-TRAP and 5X-TRAP mice. (**B**) Experimental timeline: WT-TRAP and 5X-TRAP mice underwent AD-tau seeding at three months of age in the right dorsal hippocampus and overlying cortex (posterior parietal association area) (star=injection sites), PTZ seizure kindling (or control, Sal injection) at 3.5-4 months of age, and behavioral battery at 5-6 months of age prior to brain collection for mapping studies. We performed linear mixed models with (**C**) AT8+/mm^2^ and (**D**) tdT+ cells/mm^2^ as dependent variables, genotype and treatment as fixed effects, and brain region/animal as random effects. We found significant interactions between 5XFAD genotype and PTZ treatment to increase brain-wide AT8+ levels (β=49.97, SE=9.07, t (15938)=5.51, p<0.0001) and tdT+ counts (β=13.42, SE=1.05, t (15938)=12.81, p<0.0001). (**C**) Pairwise comparisons were conducted using estimated marginal means (EMM) which revealed a significant increase in brain-wide AT8+ levels in 5X-TRAP groups compared to WT-TRAP and in PTZ kindled 5X-TRAP compared to Sal treated 5X-TRAP. Points represent EMM and bars represent 95% confidence intervals. Tukey’s post hoc: ****=p<0.0001. (**D**) Pairwise comparisons of EMM revealed a significant increase in tdT+ counts in Sal and PTZ treated 5X-TRAP compared to Sal and PTZ Treated WT-TRAP, respectively, and in PTZ treated 5X-TRAP and WT-TRAP compared to Sal treated 5X-TRAP and WT-TRAP. Three coronal sections from the Allen Brain Atlas for each experimental group displaying heatmaps for mean detected (**E**) AT8+ immunoreactive aggregates/mm^2^ and (**F**) tdT+ counts/mm^2^. Representative images of AT8 and tdT fluorescence can be found in Fig S3 and the mean, SEM, sample size, and statistical results for region/subregion AT8+ levels and tdT+ counts can be found in Tables S1, S2.

5X-TRAP and WT-TRAP mice underwent unilateral AD-tau injection into the right hippocampus and overlying cortex (posterior parietal association area) at 3 months of age, to initiate the spread of pathological tau throughout the brain (13, 16) in mice that otherwise lack overt AT8 (phospho-tau Ser202/205) pathology (39). 2-3 weeks following AD-tau seeding, PTZ kindling or control (saline, Sal) protocols were performed. On the final day of kindling (4 months of age), mice were injected with 4-OHT to permanently label all basal or seizure activated neurons with tdT. At 5-6 months of age, mice underwent behavioral testing and were euthanized at 6 months of age (3 months post AD-tau injection) (Fig 1B). This model enabled the study of post-seeding tau accumulation in basal and seizure activated networks, and in activated (tdT+) neurons to determine whether these neurons were differentially affected from tdT-neurons.

We have previously demonstrated that PTZ induced seizures worsen Aβ pathology and memory in the 5XFAD mouse (24). Here, we sought to establish whether seizures in WT-TRAP and 5X-TRAP mice are responsible for worsening of tau spread throughout the brain, as tau is central to cognitive decline and AD progression (7–10). We also examined brain-wide activity dependent labeling (tdT) to determine whether 5X-TRAP mice display neuronal hyperactivity, and whether early hyperactivity (tdT labeling at 4 months of age) is associated with subsequent tau spread.

Serially sectioned coronal samples spanning the majority of the brain from AD-tau seeded WT-TRAP and 5X-TRAP mice were registered to the Allen Brain Atlas (ABA) and tau pathology, assessed by AT8+ immunoreactive aggregates (AT8+ levels), along with detected tdT+ cells, were mapped and quantified throughout the brain (Fig 1, S1, S2, S3, S4 and Tables S1, S2).

To assess the effects of PTZ treatment and 5XFAD genotype on brain-wide AT8+ levels and tdT+ counts, linear mixed-effects statistical models were fit across all brain regions. We found a significant increase in the estimated marginal means (EMM; a model adjusted means to control for variations due to brain region and animal) in brain-wide AT8+ levels in Sal and PTZ treated 5X-TRAP compared to Sal and PTZ treated WT-TRAP mice and in PTZ treated 5X-TRAP compared to Sal treated 5X-TRAP mice (Fig 1C), consistent with our hypothesis that seizures drive tau spread in 5X-TRAP mice. With respect to activity dependent labeling, we found that PTZ induced seizures increased EMM of brain-wide tdT+ counts in both genotypes compared to Sal treated mice (Fig 1D), demonstrating increased activity-dependent neuronal labelling during seizures, as we have previously observed (36). We also found increased brain-wide tdT+ counts in Sal treated 5X-TRAP mice compared to WT-TRAP, and in PTZ treated 5X-TRAP compared to PTZ treated WT-TRAP (Fig 1D), demonstrating elevated basal and PTZ-induced neuronal activity in 5X-TRAP mice.

To examine the effects of 5XFAD genotype and PTZ on tau spread and neuronal activity levels in major brain structures, we used Bayesian statistics, as they provide full posterior distributions of parameter estimates and are robust to relatively low sample sizes and hierarchical data structures (40). Bayesian statistics identify credible effects, rather than p value-based significance used in frequentist approaches (e.g. ANOVA). Models that accounted for region-subregion relationships revealed credible interactions between 5XFAD genotype and PTZ kindling to increase AT8+ levels in all 10 major brain structures examined (Fig S1). Credible increases in tdT+ counts were also observed in all major brain structures examined due to PTZ treatment, while interactions between 5XFAD genotype and PTZ treatment to increase tdT+ counts were limited to isocortex, olfactory areas, hippocampus, cortical subplate, and striatum (Fig S2). Together, these models demonstrate substantial impacts of 5XFAD genotype and PTZ kindling to increase tau spread and activity dependent labeling in all major brain structures examined.

Given our results at the brain-wide and major brain structure levels, we next determined the effects of 5XFAD genotype and PTZ kindling on AT8+ pathology at the subregion level both ipsilateral and contralateral to tau injection site (Fig 1E, S3, Table S1). Using Bayesian models, we compared EMMs (posterior means) to determine group level differences in subregions where credible genotype, treatment, or interaction effects were found (Table S1). In the isocortex, we observed increased AT8+ levels in both PTZ and Sal treated 5X-TRAP mice compared to WT-TRAP mice across several ipsilateral and contralateral subregions. We also found that PTZ treated 5X-TRAP mice had higher AT8+ levels compared to Sal treated 5X-TRAP, including in the ipsilateral somatosensory cortex, the contralateral somatosensory, auditory, anterior cingulate, as well as dorsal retrosplenial and posterior parietal areas (Fig 1E, S3, Table S1). PTZ treatment also increased AT8+ levels in WT-TRAP mice in the ipsilateral secondary motor and contralateral anterior cingulate areas. In the hippocampus, 5XFAD genotype drove increased AT8+ levels in the ipsilateral dentate gyrus (DG) and in several subregions contralaterally (Fig 1E, S3, Table S1), with little change due to PTZ treatment. In the thalamus, we found credible increases in AT8+ levels in 5X-TRAP mice compared to WT-TRAP across several subregions. PTZ kindling resulted in increased AT8+ levels in 5X-TRAP mice in several dorsal, ventral, and medial thalamic nuclei, particularly in the contralateral hemisphere, and in bilateral anterior thalamic nuclei in WT-TRAP mice (Fig 1E, S3, Table S1). In the fiber tracts, 5X-TRAP mice showed increased AT8+ levels compared to WT-TRAP in both hemispheres, and PTZ treatment increased AT8+ levels contralaterally in 5X-TRAP and bilaterally in WT-TRAP mice (Fig 1E, Table S1), suggesting a trans-neuronal mechanism in increased AT8+ levels found due 5XFAD genotype and PTZ. Effects in the cortical subplate, striatum, hypothalamus, pallidum, and midbrain were largely driven by 5XFAD genotype and exacerbated by PTZ in select subregions (Fig 1E, Table S1). Overall, our AT8 mapping data demonstrate increased tau spread due to 5XFAD genotype brain wide. In WT-TRAP mice, PTZ worsened tau spread in select cortical and thalamic subregions and fiber tracts. In 5X-TRAP mice, PTZ kindling worsened tau spread in subregions in the majority of major brain structures, typically in the contralateral hemisphere, demonstrating that seizures drove tau spread to distal (from AD-tau injection) brain regions, which otherwise have little pathology at this stage (16). Taken together, these data support a causal role for seizures in enhancing the spread of tau.

Given we observed neuronal hyperactivity brain-wide and in major brain structures in 5X-TRAP mice, we next examined the effects of 5XFAD genotype and PTZ kindling on counts of tdT+ neurons at the subregion level (Fig 1F, S3, Table S2). In the isocortex, we found credible elevations in tdT+ counts bilaterally in PTZ treated WT-TRAP and 5X-TRAP compared to Sal treated mice in the majority of subregions, including the motor, somatosensory, auditory, and agranular insular areas (Fig 1F, Table S2). We also found increased tdT+ counts in PTZ treated 5X-TRAP mice compared to PTZ treated WT-TRAP, in these cortical subregions. PTZ kindling increased tdT+ counts in the dentate gyrus of the hippocampus, with little change due to genotype (Fig 1F, S3, Table S2). In the thalamus, we observed increased tdT+ counts in several dorsal, ventral, and medial nuclei, where PTZ treated WT-TRAP mice showed increased tdT+ counts in select regions, and PTZ kindled 5X-TRAP mice showed increased tdT+ counts compared to all other groups in most of these subregions (Fig 1F, S3, Table S2). In addition, Sal treated 5X-TRAP mice showed credibly increased tdT+ counts compared to Sal treated WT-TRAP across several thalamic nuclei (Fig 1F, S3, Table S2). Effects in the olfactory areas, cortical subplate, pallidum, hypothalamus, and midbrain were largely driven by interactions between 5XFAD genotype and PTZ kindling (Fig 1F, Table S2), similar to the isocortex and thalamus. In addition, genotype effects were found in several striatal subregions with increased tdT+ levels found in PTZ and Sal treated 5X-TRAP mice compared to WT-TRAP (Fig 1F, Table S2). Overall, our tdT+ mapping data demonstrate increased tdT+ labeling due to PTZ induced seizures in circuits relevant to AD, including limbic and thalamocortical networks. These data also demonstrate basal neuronal hyperactivity and increased PTZ induced neuronal hyperactivity in 5X-TRAP mice.

When taken together, our mapping data demonstrate that regions with increased relative AT8+ levels tended to be hyperactive (increased tdT+ counts) in both Sal and PTZ treated 5X-TRAP mice (Fig 1, S3, Table S2, S3, S4). Of note, PTZ kindled 5X-TRAP mice showed increased AT8+ levels in the contralateral intralaminar nuclei of the dorsal thalamus (rhomboid, central medial, paracentral, and parafascicular nuclei) of the thalamus and prefrontal areas (anterior cingulate, agranular insula), which are areas where we also found credibly increased levels of tdT+ counts, has previously been shown to be activated by PTZ (41, 42), and are highly interconnected, demonstrating that seizures drive tau spread through hyperactive networks. Sal treated 5X-TRAP mice also showed increased tau spread in areas with basal hyperactivity compared to Sal treated WT-TRAP, including in striatal and thalamic subregions. These data establish the first causal evidence that neuronal hyperactivity promotes tau spread preferentially through hyperactive networks.

To determine whether sex may influence tau spread and activity dependent labeling, we included sex as an independent variable in multiple linear regressions for AT8+ and tdT+ levels in all major brain structures examined (Table S3). We found that female sex was associated with AT8+ levels in the ipsilateral and contralateral hippocampi and ipsilateral fiber tracts. In contrast, sex had a less pronounced effect on tdT+ counts, but was found to be a predictor in the ipsilateral olfactory areas with elevations found in males. These results suggest that female sex is associated with increased tau spread, but these effects occurred independently of neuronal hyperactivity.

### Worsened seizure severity is associated with increased tau spread and activity-dependent labeling

As we have previously shown that 5XFAD mice have increased seizure susceptibility (24), we determined whether these effects were present in AD-tau seeded 5X-TRAP mice and whether seizure severity during the earlier kindling was correlated with AT8+ and tdT+ counts brain-wide (Fig 2, S5, S6, S7, S8). We performed PTZ kindling and activity-dependent labeling 2-3 weeks following AD-tau injection (3.5 – 4 months of age) (Fig 1B) and found that 5X-TRAP mice had significantly worsened seizure severity compared to WT-TRAP, measured by daily maximal Racine score and area under the curve (AUC) on the final day of PTZ administration (Fig 2A, B, C), confirming our prior report (24).

**Figure 2.**
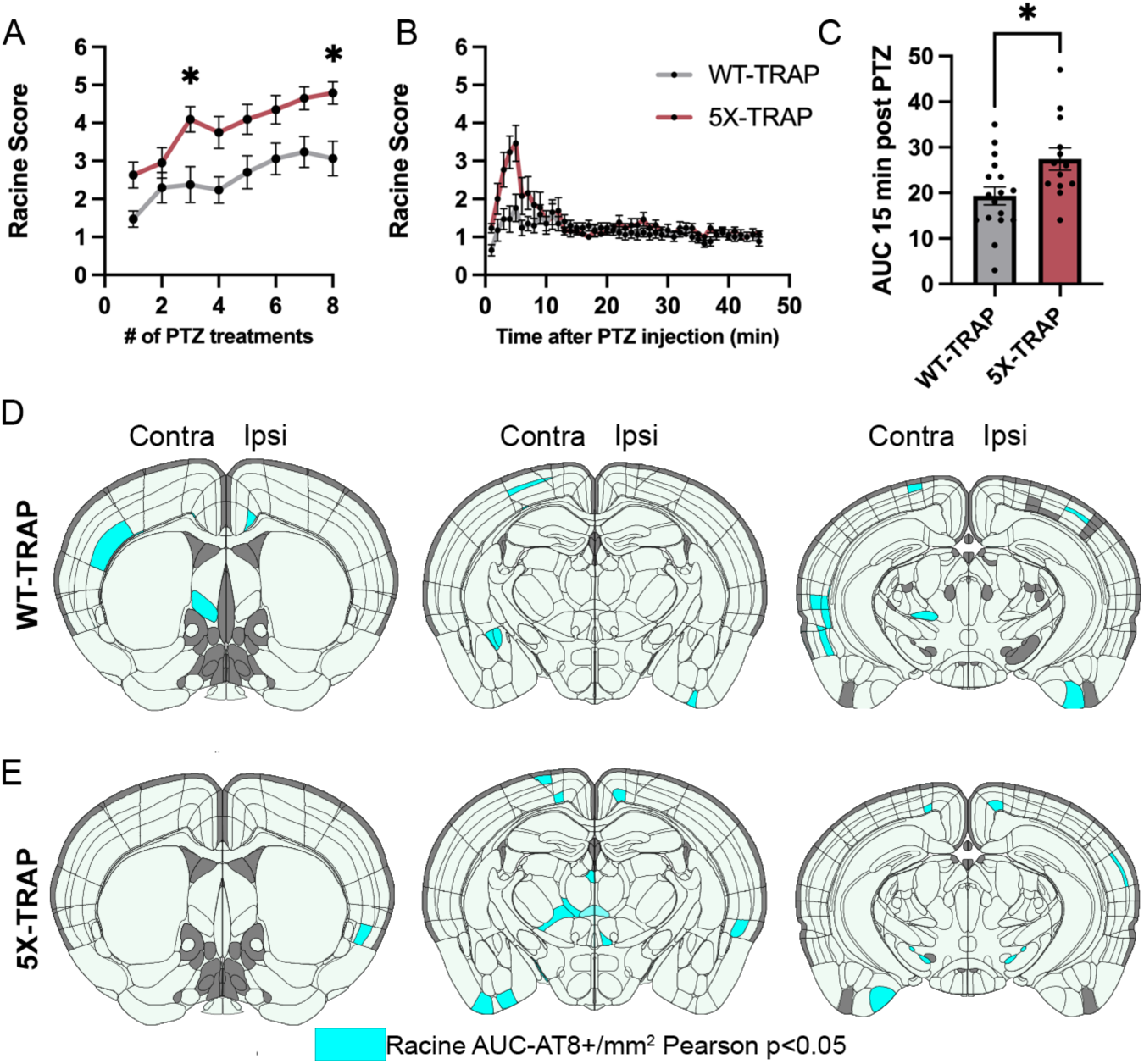
Seizure severity is increased in 5X-TRAP mice and is correlated with tau spread and activity dependent labeling. (**A**) Repeated measure (RM) two-way ANOVA with number of treatments (RM) and genotype as independent variables was performed for maximal daily seizure severity (Racine score). WT-TRAP and 5X-TRAP mice showed increasing seizure severity over the course of kindling (treatment #: F_4.95, 171.6_=15.27, p<0.0001), with 5X-TRAP mice reaching higher maximal Racine score in the 45 minutes following PTZ administration compared to WT-TRAP mice (genotype: F_1, 35_=9.71, p<0.01). Sidak’s post hoc: *=p<0.05, n= 17 (WT-TRAP)-21 (5X-TRAP). (**B**) Maximal Racine score per minute was recorded during the final PTZ administration. (**C**) 5X-TRAP mice have significantly increased area under the curve (AUC) during the first 15 minutes of the final PTZ administration. Two-tailed t-test: *=p<0.05. Brain atlas heatmaps highlight brain regions with significant positive Pearson correlations between AT8+/mm^2^ and seizure severity (Racine AUC) in PTZ kindled (**D**) 5X-TRAP mice and (**E**) WT-TRAP mice. No significant negative correlations were found. Scatter plots for these correlations can be found in Figure S5, S6. Histograms display group mean ± SEM.

Given our mapping data providing causal evidence that seizures enhance tau spread, we performed Pearson correlations between Racine AUC and regional AT8+ levels in PTZ kindled WT-TRAP and 5X-TRAP mice to determine whether seizure severity was associated with tau spread (Fig 2D, 2E, S5, S6). We found several significant positive correlations between seizure severity (Racine AUC) and AT8+ levels in both WT-TRAP and 5X-TRAP mice, largely in thalamocortical networks (Fig 2D, 2E, S5, S6). These data further indicate a critical role of seizures in enhanced tau spread, particularly in bilateral thalamocortical brain regions.

Next, since our prior work demonstrated associations between hippocampal activity dependent labeling and seizure severity (36), we performed Pearson correlations between seizure severity and tdT+ counts in 5X-TRAP and WT-TRAP mice (Fig S7, S8). We found significant positive correlations largely in cortical subregions (Fig S7, S8), demonstrating that seizure severity is associated with elevated activity dependent labeling.

### Tau spread and neuronal hyperactivity are correlated with worsened memory in PTZ kindled 5X-TRAP mice

All WT-TRAP and 5X-TRAP underwent the novel object recognition (NOR), contextual fear conditioning (CFC), and open field behavioral assays at 5 – 6 months of age (Fig 1B). In the NOR paradigm, the % time spent with the novel object as compared to the familiar object was significantly decreased in PTZ kindled 5X-TRAP mice compared to Sal treated 5X-TRAP and WT-TRAP mice (Fig 3A), demonstrating memory deficits in this group. CFC testing at 1 day (recent memory) and 14 days (remote memory) following conditioning revealed no differences between groups in recent or remote recall (Fig S9). In the open field assay, we observed decreased rearing behavior in 5X-TRAP mice, overall, compared to WT-TRAP, suggestive of decreased exploratory behavior and motor activity (Fig 3B), but the total distance traveled and the time spent in the center of the arena, a measure of anxiety-like behavior, were similar across all groups (Fig 3B). Female sex was associated with decreased rearing behavior in the open field assay, while no effects of sex or cohort were found on NOR or CFC (Table S3, S4). These results demonstrate that seizures significantly worsen memory in AD-tau seeded 5X-TRAP mice.

**Figure 3.**
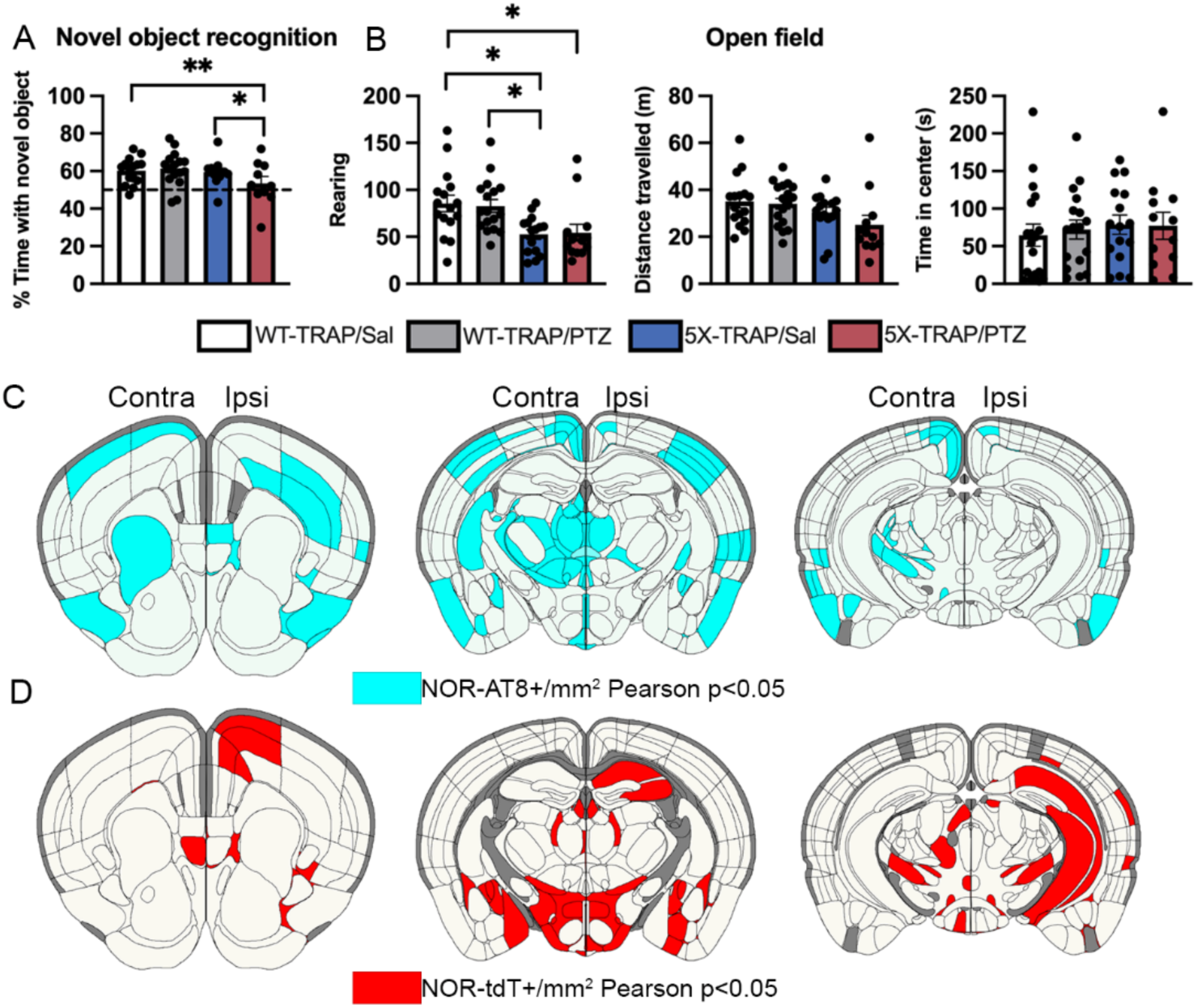
Memory deficits in PTZ kindled 5X-TRAP mice are correlated with tau spread and network hyperactivity. Two-way ANOVA with genotype (WT-TRAP vs 5X-TRAP) and kindling (Sal vs PTZ) were performed for behavioral tests (**A**, **B**). (**A**) PTZ-kindled 5X-TRAP mice had significantly decreased % time spent with novel object compared to saline treated 5X-TRAP and WT-TRAP mice (interaction: F_1,49_=4.45, p<0.05). (**B**) 5X-TRAP mice showed reduced rearing behavior (left; genotype effect: F_1,58_=17.21, p<0.0001) and total distance travelled (center; genotype effect: F_1,56_=6.01, p<0.05) without change in time spent in center (right). *, **=Sidak’s post hoc p<0.05, 0.01; n=10-16/group. Brain atlas heatmaps highlight brain regions with significant negative Pearson correlations between % time spent with novel object in the NOR task and (**C**) AT8+/mm^2^ and (**D**) tdT+ aggregates/mm^2^ in PTZ kindled 5X-TRAP mice. No significant positive correlations were found. Scatter plots for these correlations can be found in Figures S10, S11. Histograms display group mean ± SEM.

To determine the associations between tau spread and neuronal hyperactivity on cognition, we performed Pearson correlations between NOR performance and regional AT8+ levels and tdT+ counts in PTZ kindled 5X-TRAP mice (Fig 3C, D, S10, S11). We found strong inverse correlations between % time spent with novel object and AT8+ levels extensively throughout the brain, including in the ipsilateral intralaminar nuclei of the dorsal thalamus and agranular insular area, and in the contralateral intralaminar nuclei of the dorsal thalamus, as well as bilateral amygdalar, striatal, and hypothalamic nuclei and fiber tracts (Fig 3C, S10). Notably, many of these regions are areas in which we found elevated tau spread in PTZ kindled 5X-TRAP mice (Fig 1E, Table S1). In addition, we found that decreased NOR performance was significantly correlated with tdT+ counts in the ipsilateral hippocampal subfields, bilaterally in the intralaminar nuclei of the thalamus, prefrontal areas (agranular insula and motor), and in striatal and hypothalamic nuclei of PTZ kindled 5X-TRAP mice (Fig 3D, S11). These results demonstrate that seizure-driven exacerbation of tau spread and neuronal hyperactivity, particularly in thalamocortical circuits, are strongly associated with memory deficits in 5X-TRAP mice.

### Network dynamics of tau spread are altered by 5XFAD genotype and seizure induction

Given our data demonstrating increased tau propagation in PTZ kindled and 5X-TRAP mice, we next employed well-validated computational models (19, 43) to study whether 5XFAD genotype or PTZ kindling impacted the pattern of tau spread between brain regions. Prior studies using computational modeling of tau pathological progression have demonstrated that tau spreads throughout the brain via the neuroanatomical connectome, predominantly in a retrograde direction (19). We implemented linear diffusion models of tau spread from AD-tau injection sites (hippocampal and posterior parietal association area (PTLp) seeds) along a network defined by either anterograde connectivity, retrograde connectivity, or Euclidean (geometric) distance between brain areas derived from Oh and colleagues (44). We then compared the performance of each of these spread models in predicting regional AT8+ levels from each experimental group (Fig 4, S12).

**Figure 4.**
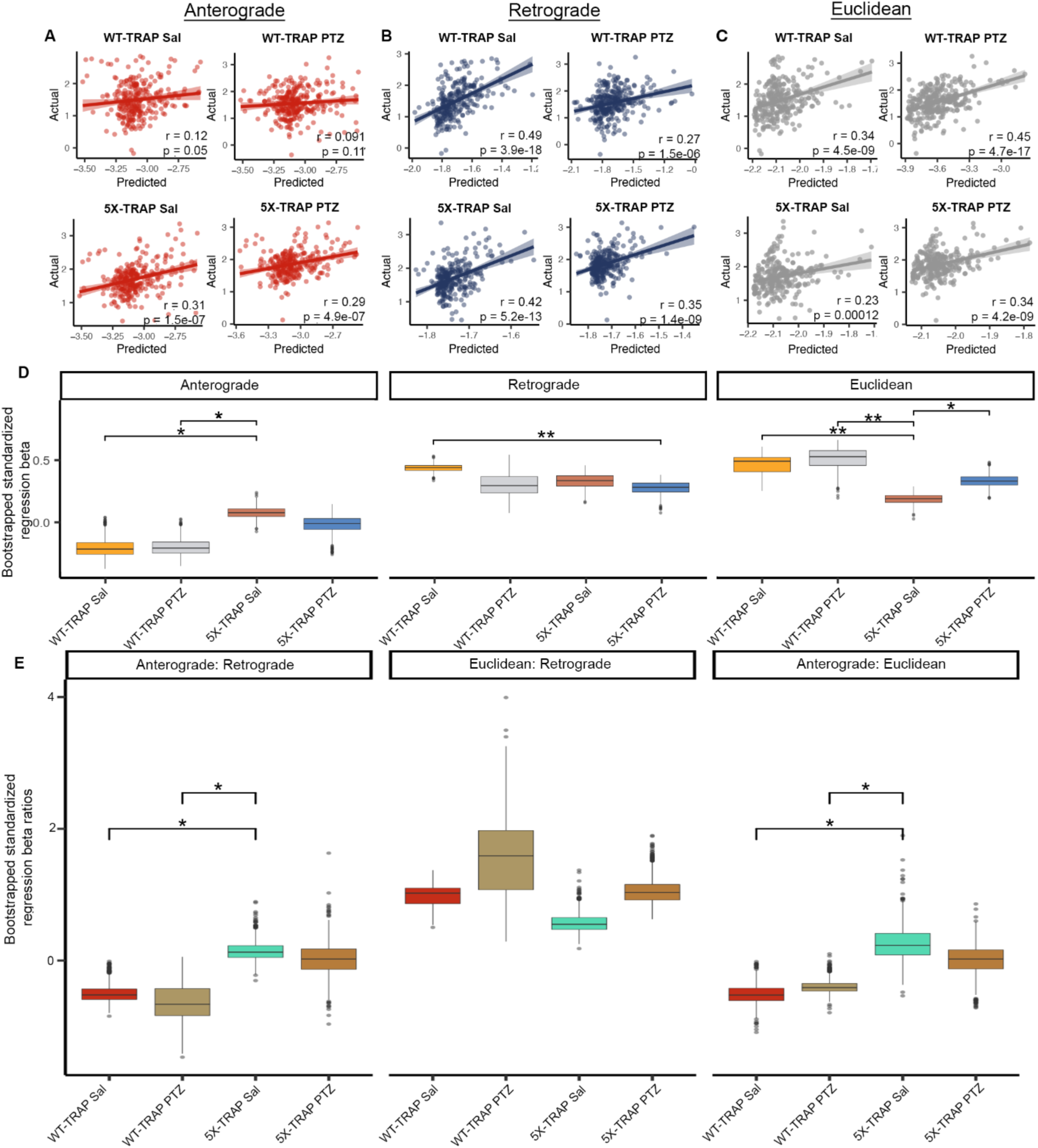
Computational modeling reveals an increased impact of anterograde tau spread in 5X-TRAP mice. Predictions of tau spread based on (**A**) anterograde, (**B**) retrograde, and (**C**) Euclidean models of spread plotted against our measured levels for each experimental group. Solid lines represent best fit and shaded regions are the 95% confidence interval. (**D**) For the Bidirectional Euclidean + tdT spread model architecture, we performed a bootstrap comparison of mean standardized regression beta weights between groups and adjusted for multiple comparisons. (**E**) For the Bidirectional Euclidean spread model architecture, we performed a bootstrap comparison of mean ratio of standardized anterograde: retrograde, Euclidean: retrograde, and anterograde: Euclidean regression beta weights between groups and adjusted for multiple comparisons. p-values were adjusted for multiple comparisons in each panel using FDR correction: *p<0.05, **p<0.01. Box plots boxes display the median, 25^th^ and 75^th^ percentiles, and whiskers extend to the data point that is at most 1.5 x interquartile range, and data beyond this point are plotted individually.

Consistent with prior data (19), we found that retrograde models tended to outperform anterograde and Euclidean models (Fig 4A-C, Fig S12). However, anterograde models produced a statistically significant fit with 5X-TRAP mice but not with WT-TRAP mice (Fig 4A). A model combining anterograde, retrograde, and Euclidean distance called “Bidirectional Euclidean” fit better than any individual model in all groups (Fig S12). To further examine the relative importance of anterograde, retrograde, and Euclidean tau spread, we compared standardized regression beta weights from the “Bidirectional Euclidean” model (Fig S12) between groups, which provide a quantitative measure of the relative importance of each mode of spread to the overall model fit (Fig 4D). We found that anterograde beta weights (Fig 4D) and anterograde: retrograde and anterograde: Euclidean beta weight ratios (Fig 4E) were increased in Sal treated 5X-TRAP mice compared to WT-TRAP, suggesting that anterograde connectivity specifically has an increased contribution to tau spread in 5X-TRAP mice relative to WT-TRAP mice. Beta weights were similar between PTZ and Sal treated 5X-TRAP mice (Fig 4D).

Retrograde betas were lower in PTZ kindled 5X-TRAP mice compared to Sal treated WT-TRAP mice, while Euclidean spread betas were lower in Sal treated 5X-TRAP compared to all other groups (Fig 4D). To confirm the regional specificity of the model, we repeated the retrograde spread modeling in WT-TRAP mice using 500 random seed sites with similar spatial clustering to the seed sites (i.e. the dorsal hippocampus and posterior parietal association area (PTLp)). We found that the model performance with the experimental seed site was better than 99% of the randomly chosen seed sites (Fig S13). In addition, we implemented linear diffusion models using hippocampus only and PTLp only seed sites (Fig S14). In general, the hippocampal model recapitulated the dual seed site model, while the PTLp model performed poorly, suggesting that the hippocampus is the primary seed site for tau spread across our experimental groups.

Overall, these computational analyses show that all groups fit with a pattern of connectome-based spread and that primarily anterograde spread contributes to the elevations in AT8+ levels quantified in our brain mapping data in Sal and PTZ treated 5X-TRAP mice (Fig 1).

### Regional neuronal activity levels at 4 months of age are predictive of tau spread at 6 months of age

Given that we have established a causal role of seizures in elevated tau spread in 5X-TRAP mice and that elevations in tau pathology tended to occur in hyperactive brain regions, we next determined whether tdT+ counts were predictive of tau spread. To do so, we correlated regional counts of tdT+ neurons with detected AT8+ immunoreactive aggregates while accounting for predicted Bidirectional Euclidean spread (Fig 5). We found significant positive relationships between tdT+ counts and relative AT8+ levels in Sal treated WT-TRAP mice, demonstrating that basal neuronal activity is predictive of tau spread. tdT+ counts were not predictive of relative tau spread in PTZ treated WT-TRAP mice, since PTZ increased tdT+ counts with relatively little change in AT8+ levels in these mice (Fig 1, S3, Table S1, S2). Additionally, tdT+ counts were predictive of tau spread in Sal treated 5X-TRAP mice, suggesting that basal hyperactivity found in Sal treated 5X-TRAP mice contributes to increased tau spread. A positive relationship between tdT+ counts and AT8+ levels was also found in PTZ treated 5X-TRAP mice, further strengthening the causal role of neuronal hyperactivity in tau spread established in this group. Overall, given that activity dependent labeling took place soon after tau seeding (prior to extensive tau spread (16, 19) (Fig 1B)), these data demonstrate a preferential vulnerability of hyperactive circuits to tau propagation, and provide additional evidence of hyperactivity driven tau spread in 5X-TRAP mice.

**Figure 5.**
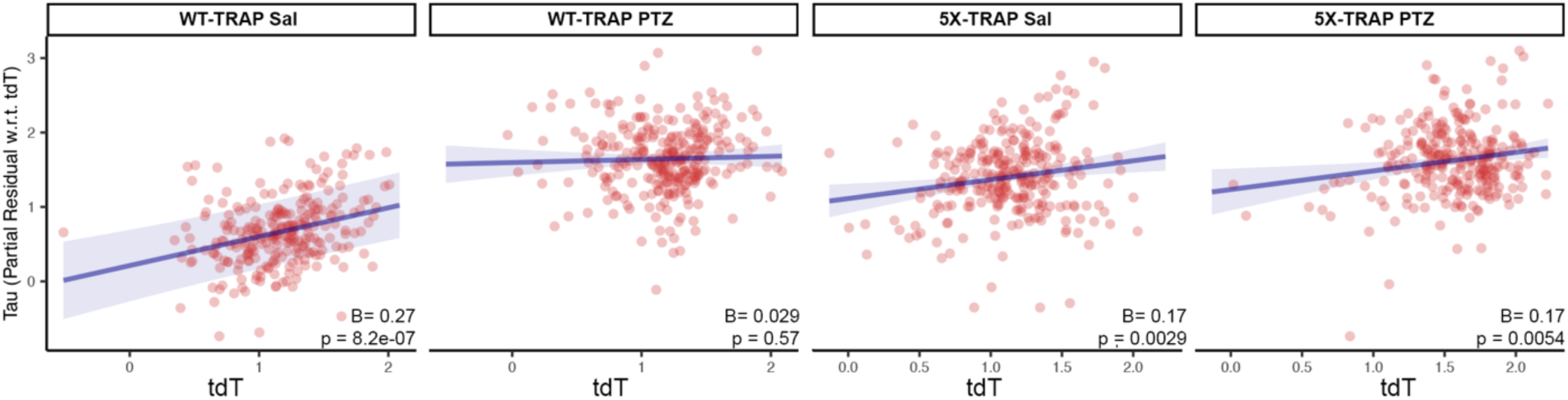
Regional tdT+ levels are predictive of tau spread. We fit a linear regression model separately for WT-TRAP and 5X-TRAP mice to estimate regional tau pathology using anterograde spread, retrograde spread, Euclidean distance-based spread, tdT+ cell density, and interactions between PTZ treatment and tdT positive cell density as predictor variables. In the plots above, we show partial residual plots of tau pathology (y-axis) versus tdT cell density (x-axis) for each level of the interaction with PTZ treatment (i.e. PTZ and Saline) while controlling for estimated anterograde, retrograde, and Euclidean distance-based spread. Individual points correspond to anatomical regions with available tdT+ cell density (n = 6-15 mice/group/brain region; 261-288 brain regions/group). tdT+ levels were predictive of tau spread in all groups except PTZ treated WT-TRAP mice. p-values test the significance of simple slopes of tdT as a predictor of AT8+ levels for each group.

### Hyperactive neuronal populations labelled at 4 months of age are more likely to exhibit tau pathology than surrounding neurons at 6 months of age

Given our results establishing that tau spreads via hyperactive networks in 5X-TRAP mice, we next determined whether hyperactive neurons accumulate more pathological tau. We examined whether tdT+ neurons were more likely to develop somatic AT8+ immunoreactivity compared to surrounding tdT-(NeuN+) neurons. Detected and mapped tdT+ and tdT-(NeuN+) neurons were colocalized with detected AT8+ immunoreactive aggregates across all sampled brain regions to calculate the percentage of tdT+ and tdT-neurons with somatic tau pathology in 5X-TRAP (Fig 6) and WT-TRAP (Fig S15) mice. Ipsilaterally, we found increased AT8+ colocalization with tdT+ neurons compared to tdT-in Sal treated 5X-TRAP mice in the isocortex, cortical subplate, striatum, and thalamus and increased AT8+ colocalization in tdT+ neurons compared to tdT-in PTZ kindled 5X-TRAP mice in the striatum and thalamus. In the contralateral hemisphere, we found that tdT+ neurons were more likely to colocalize with AT8 immunoreactivity than surrounding tdT-neurons in the thalamus of PTZ kindled 5X-TRAP mice (Fig 6). Given our results showing increased AT8+ levels and high tdT+ counts in the cortex of PTZ treated 5X-TRAP mice, we expected to see increased tdT+ colocalization with AT8 in the cortex, but we found no differences in these mice. These results are likely due to increased neuronal death (Fig S16) in seizure activated cortical neurons carrying a high tau load. In contrast, WT-TRAP mice did not exhibit any significant differences in AT8 colocalization due to kindling or tdT labelling (Fig S15). In summary, both Sal and PTZ treated 5X-TRAP mice exhibited increased tau pathology in tdT+ neurons compared to tdT-neurons in regions with increased overall AT8+ levels and tdT+ counts, demonstrating that seizure and basally hyperactive populations disproportionately contribute to tau spread in these mice.

**Figure 6.**
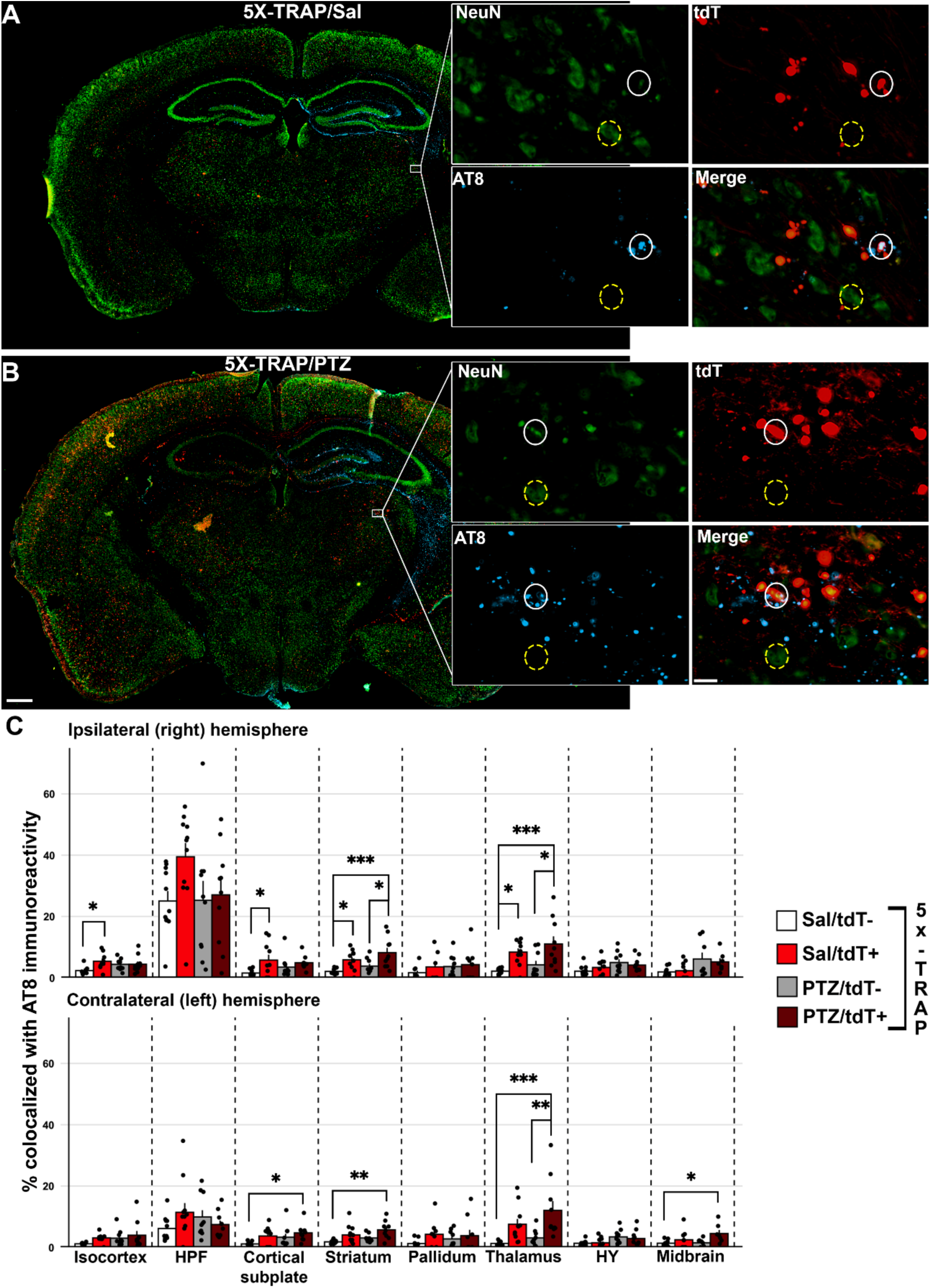
tdT+ neurons are more likely to colocalize with tau pathology in both saline and PTZ treated 5X-TRAP mice. Representative slide scanned coronal sections from (**A**) saline and (**B**) PTZ treated 5X-TRAP mice. Inset displays thalamic tdT+ (white circles) and tdT-(NeuN+) (yellow dashed circles) neurons colocalization with AT8+ immunoreactive aggregates. Scale bars = 500 µm, 20 µm (inset). (**C**) tdT+, tdT-, and colocalized AT8+ cells were mapped to the Allen Brain Atlas and % of detected tdT+/tdT-cells were quantified in 9 major brain structures, bilaterally. A two-way ANOVA was performed with treatment (Sal vs PTZ) and cell type (tdT-vs tdT+) as independent variables for each brain region. There was a significant increase in percentage of tdT+ neurons colocalized with AT8 than surrounding tdT-in Sal treated 5X-TRAP mice in the ipsilateral thalamus (cell type: F_1,38_=19.17, p<0.0001), striatum (treatment: F_1,38_=4.39, p<0.05; cell type: F_1,38_=17.62, p<0.001), and isocortex (F_1,34_=4.12, p<0.05). There was a significant increase in % tdT+ neurons colocalized with AT8+ than surrounding tdT-cells in PTZ treated mice in the ipsilateral thalamus and striatum and contralateral thalamus (cell type: F_1,38_=18.60, p<0.001). Tukey’s post hoc: *p<0.05, **p<0.01, ***p<0.001; n=7-11/group/brain region. Histograms display group mean ± SEM.

### Human postmortem AD brains exhibit increased tau pathology in association with a history of seizures

To determine whether seizures worsen indications of tau spread in AD, we analyzed postmortem pathological ratings across 18 brain regions from AD patients and stratified subjects based on the presence or absence of clinical seizure history. We found that AD patients with clinical seizure history (AD+Sz) exhibited more severe tau pathological ratings (phosphorylated tau (Ser396/404); PHF1 antibody) compared to AD patients without a clinical seizure history (AD-Sz). We found these alterations in regions affected at late Braak stages, the angular gyrus and a strong trend in the middle frontal gyrus (Fig 7). We did not find significant differences in regions involved in early Braak stages including the locus coeruleus and entorhinal cortex (Fig 7). We additionally performed ordinal linear regressions and found that male sex was associated with increased tau pathology in the substantia nigra and locus coeruleus (Table S5). These patterns indicate that a history of seizures is associated with increased tau pathology in brain regions impacted at advanced disease stages in AD, supporting our causal evidence that seizures increase tau spread in 5X-TRAP mice.

**Figure 7.**
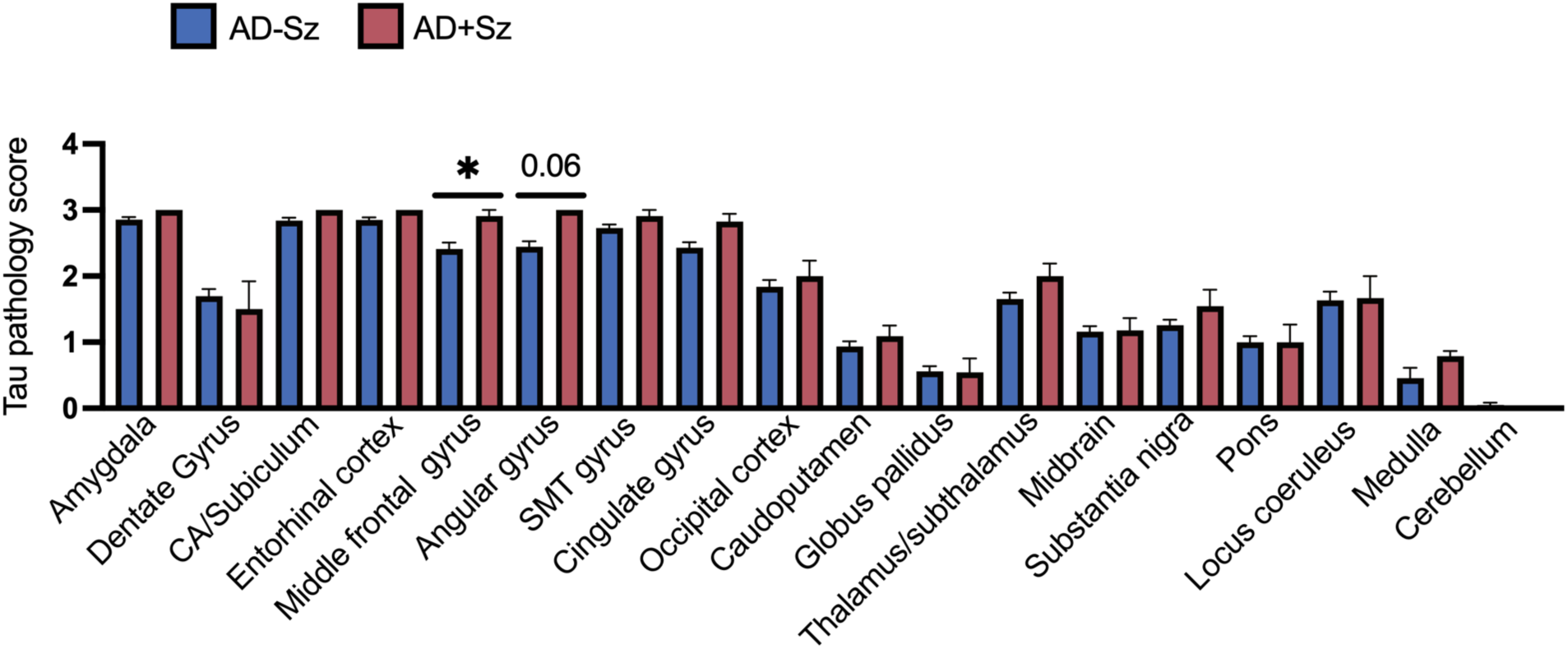
Worsened tau pathology in AD patients with a history of seizures. Tau pathology severity ratings from postmortem analysis of AD-Sz and AD+Sz were compared by two-tailed Mann-Whitney test. AD+Sz had significantly increased tau rating compared to AD-Sz in the middle frontal gyrus and a strong trend in the angular gyrus. *p<0.05, AD-Sz n= 74-95/brain region, AD+Sz n=8-11/brain region. SMT gyrus= superior/middle temporal gyrus. Histograms display group mean ± SEM.

## Discussion

Our study demonstrates for the first time that neuronal hyperactivity increases pathological tau spread through interconnected brain regions in an AD model. Our data also highlight that tau preferentially spreads through hyperactivated brain networks and neuronal populations in the anterograde direction in 5X-TRAP mice. Further, we found that tau spread and neuronal hyperactivity are highly and extensively correlated with worsened memory caused by seizures. We found corroborating data in postmortem human AD brain tissue, suggestive of increased tau spread in patients with a history of seizures. Together, our data demonstrate hyperactive networks and neurons as therapeutic targets to slow AD progression.

TRAP models are widely used in basic research and our recent work demonstrated that tdT labelling during seizure induction allows for the long-term tracking of hyperactive neuronal populations (36). Here we report the development of a novel model involving the cross of these mice with a neurodegenerative mouse. With tamoxifen-inducible, neuronal activity (Fos) driven labeling, the 5X-TRAP model can be used to examine the interactions between neuronal activity and AD pathology. For these studies, we used the 5X-TRAP mouse in conjunction with AD-tau seeding and PTZ seizure kindling as a robust model to test our hypothesis that neuronal hyperactivity drives subsequent tau spread and to determine the contribution of activated networks and neuronal populations to tau spread.

Prior studies have demonstrated that induced neuronal activity can worsen endogenous pathology in a tauopathy model (17). Here, using a tau seeding paradigm, we provide the first *in vivo* evidence of a causal relationship between neuronal hyperactivity, through seizure induction, and increased tau spread in mice that otherwise lack overt phospho-tau pathology (16, 39). In addition, we show that injection of pathological tau in the mouse brain recapitulates aspects of the connectome-directed propagation observed in human AD (13, 19). We performed AD-tau seeding unilaterally in the right dorsal hippocampus and overlying cortex (posterior parietal association area), as prior studies have shown robust tau spread following seeding at these sites (13, 16, 19). In contrast, tau seeding in the locus coeruleus, the earliest region impacted in AD, results in limited spread, with the hippocampus and entorhinal cortex largely spared of tau pathology (45, 46). Following AD-tau seeding, mice underwent PTZ seizure induction or control (saline) administration and all seizure or basally activated neurons labeled with tdT using the TRAP methodology. We found that 4 month old 5X-TRAP mice show basal increases in activity-dependent tdT labeling, corroborating prior reports of hyperexcitability in amyloidosis models (28, 47), which may underlie increased seizure susceptibility in 5XFAD (24) and 5X-TRAP mice.

For an unbiased approach to measure the spread of tau, we performed comprehensive brain mapping on sectioned tissue from AD-tau seeded WT-TRAP and 5X-TRAP mice. First, we found that Sal treated 5X-TRAP mice had significantly increased tau spread in cortical and subcortical regions and fiber tracts, relative to WT-TRAP mice, expanding upon prior reports of increased spread in amyloidosis models, including 5XFAD mice (15, 16). Next, we observed that PTZ kindling caused further increases in tau spread in 5X-TRAP mice in several cortical regions, the thalamus, striatum, and fiber tracts. The majority of PTZ driven effects in 5X-TRAP mice were found contralateral to tau injection, demonstrating that seizures drove tau spread to regions that would otherwise have minimal pathology at this stage (3 months post seeding) (16). In both PTZ and saline-treated 5X-TRAP mice, regions with high tdT counts tended to be associated with relative increases in tau pathology. PTZ induces thalamocortical seizure activity (41, 42), and our data show increased spread particularly in hyperactive interconnected regions of the thalamus and cortex, such as intralaminar nuclei of the dorsal thalamus and prefrontal regions, and in fiber tracts. These data suggest that seizure induced hyperactivity (increased tdT+ counts) drives axonal transport and tau spread through activated networks.

Thalamocortical Ttau spread in 5X-TRAP mice was associated with worsened cognitive performance in the NOR task, consistent with human studies linking cognitive decline and tau progression in AD (7–10). Of note, we did not find significant differences in CFC, which may indicate a selective deterioration of circuit function. CFC is largely mediated by the amygdala and hippcompus, while NOR is cortically driven. These behavioral data further establish a role of seizure-induced tau spread in overall disease progression.

PTZ kindled WT-TRAP mice also showed increased tau spread through thalamocortical networks and fiber tracts, consistent with seizure driven spread through activated circuits. However, compared to 5X-TRAP mice, PTZ kindling had relatively little effect on tau pathology in WT-TRAP mice. One plausible explanation for this may be that interactions between Aβ and neuronal hyperactivity promote enhanced seeding effects. Notably, we have found that PTZ kindling increases Aβ pathology in 5XFAD mice (24), which in turn can further increase and promote sustained hyperactivity, creating a positive feedback loop (32), which may result in exacerbated tau spread. Indeed, amyloid-related hyperconnectivity was recently associated with tau spread in AD (14). A second explanation is that WT-TRAP mice had significantly reduced seizure severity compared to 5X-TRAP mice. Future studies should continue kindling in WT-TRAP mice until similar levels of tonic-clonic seizures are achieved to determine whether increased seizure severity could increase tau spread in WT mice. Together, our mapping data suggest that Aβ and neuronal hyperactivity interactions promote the spread of tau in 5X-TRAP mice.

Notably, we found a significant effect of female sex on increased tau spread in mice, although similar changes were not found in our human cohort. This may indicate that effects of sex on tau spread may be dependent on disease severity, as these cases were late-stage. Indeed, our results in mice are consistent with tau positron emission tomography (PET) studies in patients with mild to moderate cognitive impairment due to AD, in which females had accelerated tau propagation (48, 49). Our data do not suggest that sex effects on tau spread in mice are due to changes in neuronal hyperactivity as we found little difference between males and females in seizure severity or tdT labeling. However, since we have previously demonstrated that female 5XFAD mice had increased Aβ plaques and mTORC1 activation compared to males (24), it is plausible that increased tau spread in females found here may be due to increased neuritic plaque tau or a reduction in autophagy, caused by elevated mTOR activity (50), resulting in increased intracellular tau accumulation.

To determine whether genotype or seizure induction altered the patterns of tau spread, we utilized connectomic data made available from the Allen Institute (44). Consistent with prior results (19), models incorporating anterograde, retrograde, and Euclidean distance-based spread had better fit with measured AT8+ levels than any model individually. These data are indicative, overall, of a connectome driven spread of tau, which is further supported by our findings of high AT8+ levels in fiber tracts, particularly in PTZ treated 5X-TRAP mice. These models also indicate that, while we used two seed sites, the hippocampus was the primary initial site of tau seeding effects. From the dual seed site analyses, measured AT8+ levels in PTZ treated 5X-TRAP and WT-TRAP mice tended to have worse fit with anterograde and retrograde models, compared to their Sal treated counterparts, which may be indicative of altered anatomical and functional long-range connectivity (reducing fit with a standardized connectomic dataset) and enhanced local connectivity caused by seizures (51–54). We also found that measured AT8+ levels in 5X-TRAP mice, but not WT-TRAP, fit with anterograde models of spread. Indeed, compared to anterograde, we found lower relative contribution of Euclidean and retrograde models to spread in Sal and PTZ treated 5X-TRAP mice, respectively. When taken with our mapping data, these results suggest that increased AT8+ levels in 5X-TRAP mice are primarily due to increased anterograde spread.

We next used these connectome-based models to further link neuronal activity to tau spread. We found that regional neuronal activity levels (tdT+ counts) were predictive of relative tau spread in all groups except PTZ treated WT-TRAP mice. The lack of an effect in PTZ treated WT-TRAP mice was expected since there was increased tdT+ counts due to seizure induction with relatively little change in tau spread. The relationship between neuronal activity level and predictivity of tau spread in Sal treated WT-TRAP mice suggest that tau preferentially travels between basally active networks. Similarly, tau has been shown to spread through the default mode network in human patients, which are more metabolically active at rest (1, 4, 6). Our results also indicate that regional neuronal activity levels are predictive of relative tau levels in 5X-TRAP mice, further suggesting that neuronal hyperactivity drives increased tau spread in these mice, demonstrating a vulnerability of hyperactive networks to tau propagation.

Another novel aspect of our study was the ability to specifically label and track hyperactivated neuronal populations to determine their contribution to tau spread months after their original activation. We found that tdT+ neurons were more likely to colocalize with AT8 than the surrounding tdT-neurons in the thalamus and striatum of both PTZ and Sal treated 5X-TRAP mice, indicating that these populations drove enhanced overall tau spread. Increased pathology found in these populations may also be associated with increases in tau kinases, as seen in temporal lobe epilepsy (29). TRAP labeling utilizes cFos as a proxy marker of neuronal activity limiting the temporal precision of neuronal dynamics. While elevated tdT+ levels indicate early population hyperactivity and we have previously demonstrated sustained hyperexcitability in seizure-activated neurons (36), the mechanisms underlying cellular hyperexcitability in tdT+ 5X-TRAP neurons remain to be explored. Nevertheless, the ability to label and track these populations provides a robust framework through which mechanistic studies may be performed to identify the pathophysiological alterations that underlie tau propagation in these vulnerable populations to identify therapeutic targets to slow tau spread.

To expand our investigations of the role of neuronal hyperactivity in tau pathology distribution in human AD, we performed a retrospective analysis of postmortem tau pathological ratings brain-wide. We found changes consistent with increased tau spread in AD patients with seizure history compared to those without, consistent with a clinical study demonstrating that the development of seizures in AD is associated with increased CSF tau levels (55). While clinical examination of the relationship between seizures and tau localization in the brain is limited, a recent study from Lam and colleagues (56) found associations between seizure foci and spatial development of both tau and amyloid pathology, additionally corroborating our results that neuronal activity levels were predictive of tau spread in that region. These data suggest that there may be clinical value in screening AD patients for epileptiform activity or other electrographic biomarkers, along with tracking tau progression by PET.

Overall, our data support neuronal hyperexcitability as a target to slow AD progression, especially early in the disease. Indeed, the antiseizure medication (ASM), levetiracetam, slowed cognitive decline in mild cognitive impairment (57) and AD patients with epileptiform activity (58), but both studies used short term treatments (≤ 4 weeks) and did not examine pathological progression. Based on our results, studies with long term ASM therapy and serial tau PET scans would be warranted to link seizure control with modulation of tau spread. Other drugs targeting hyperexcitability may also hold promise. Indeed, we have found that modulating network dysfunction via inhibition of mTORC1 activity with rapamycin reduced Aβ and cognitive deficits in PTZ kindled 5XFAD mice (24).

Together, these studies indicate that in the presence of Aβ, neuronal hyperactivity promotes anterograde tau spread through hyperactive networks and neurons, with rates of tau spread associated with cognitive dysfunction. Thus, we contribute to a model of AD progression whereby amyloid-neuronal hyperactivity interactions enhance the spread of tau, advancing disease progression and cognitive decline, a process which is exacerbated by seizures (Fig 8). Targeting neuronal hyperactivity, particularly at early, Aβ-predominant stages in AD, should be prioritized to slow overall disease progression.

**Figure 8.**
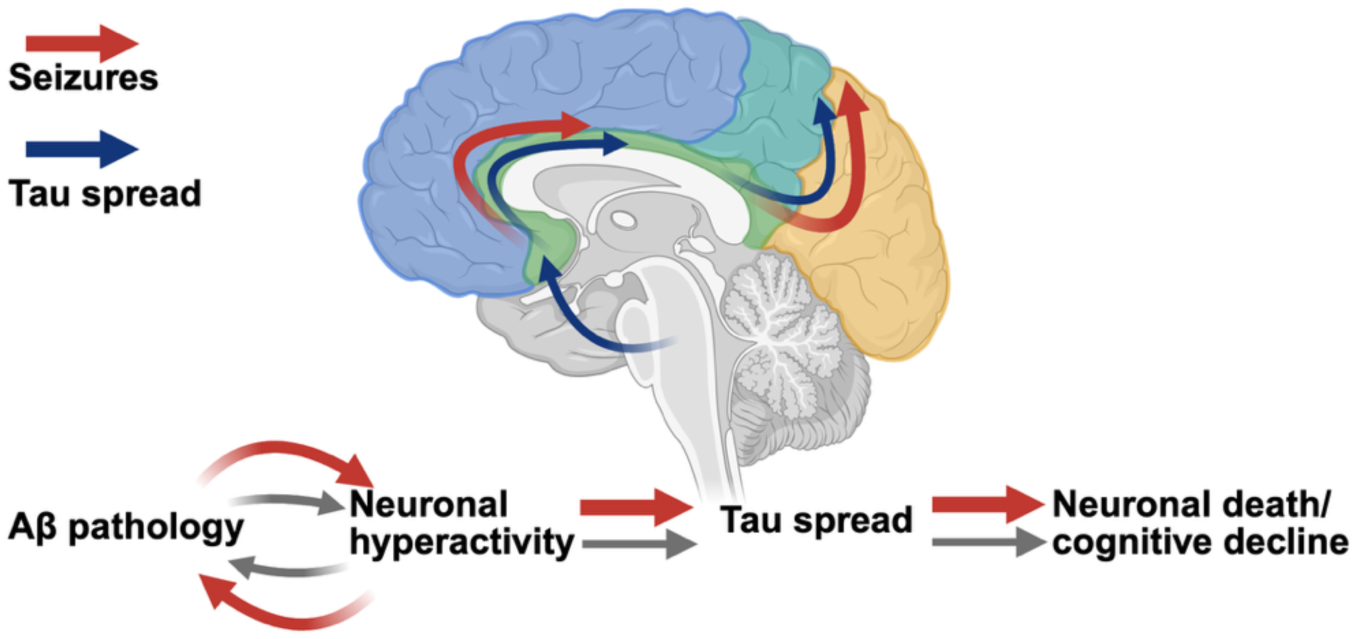
Schematic of proposed model for AD progression.

## Methods

### Mice

All mouse procedures and protocols were approved by the University of Pennsylvania Institutional Animal Care and Use Committee (IACUC) Office of Animal Welfare. Fos^2A-iCreER^ (TRAP2) and B6.Cg-*Gt(ROSA)26Sor*^tm14(CAG-tdTomato)Hze^/J (Ai14) were obtained from Jackson Laboratory (stock 030323, 007914). TRAP2 and Ai14 mice were crossed for two generations, producing mice homozygous for Fos^2A-iCreER^ and B6.Cg-*Gt(ROSA)26Sor*^tm14(CAG-tdTomato)Hze^/J. 5XFAD on a mixed B6SJL background first generation (F1) generated from mice purchased from Jackson Laboratory (MMRRC strain #034840, Bar Harbor ME) were bred with mice homozygous for TRAP2/Ai14 to produce 5XFAD x TRAP (5X-TRAP) and littermate WT x TRAP (WT-TRAP) experimental mice that were used in all biochemical and behavioral experiments.

### AD-tau isolation

AD-tau isolation was performed as previously established (13) from cortical tissue from patients confirmed to have AD by neuropathological diagnosis. Total protein concentrations were determined by BCA and tau, Aβ, and alpha synuclein concentrations were determined by ELISA: purified tau= 0.4-1.1 μg/μL, Aβ_42_ = 32-132 ng/mL, Aβ_40_ =3-46 ng/mL, and α synuclein= nondetectable – 0.54 μg/mL.

### Stereotaxic surgery

AD-tau was injected into 3-month-old mice using previously established protocols (16). Briefly, mice were anesthetized with ketamine-xylazine-acepromazine (90 mg/kg, 10 mg/kg, 2 mg/kg) and mounted in a stereotaxic apparatus. A hole was bore from 2.5 mm posterior from Bregma and 2.0 mm lateral (right) from midline. A Hamilton syringe was lowered 2.4 mm and 2.5 μL of 0.4 μg/μL (1 μg) AD-tau was injected into the dorsal hippocampus. The syringe was then retracted 1 mm (1.4 mm depth) and an additional 1 μg of AD-tau was injected into the posterior parietal association area. An example of tau injection sites can be found in Figure S4.

### PTZ kindling

Two to three weeks following AD-tau injection, mice underwent PTZ kindling using previously established protocols (24, 28). Briefly, mice were injected (I.P.) every 48 hours for 15 days (8 injections) with 35 mg/kg PTZ (Sigma Aldrich, St Louis MO) or vehicle (0.9% saline, Sigma-Aldrich). Mice were monitored and video-recorded for 45 minutes after PTZ administration and Racine scored for seizure severity. We used a modified Racine score (ref): 0=normal behavior, 1=freezing, 2=hunching with facial automatism, 3=rearing and forelimb clonus, 4=rearing with forearm clonus and falling, 5=tonic-clonic seizure, 6=death. To minimize death, mice received 31.5 mg/kg PTZ following a stage 5 seizure and if mice reached three consecutive days of stage 5 (tonic-clonic) seizures, they underwent one additional PTZ treatment and were then removed from kindling. To generate area under the curve (AUC) for seizure activity, we used the maximal score reached per minute over 15 minutes following the final PTZ administration.

### 4-OHT administration

5X-TRAP and WT-TRAP mice were injected with 4-hydroxytamoxifen (50 mg/kg, I.P.) to induce permanent tdTomato expression in neurons that were activated (Fos-expressing) in the 4-6 hour window after administration. AD-tau seeded 5X-TRAP and WT-TRAP mice were injected with 4-OHT 30 min prior to the final PTZ or saline treatment to induce tdTomato expression in seizure or basally activated (Fos expressing) neurons (36).

### Mouse behavior

Behavioral experiments were carried out in the Neurobehavior Testing Core in the Translational Research Center at the University of Pennsylvania. Experimenters were blind to genotype and condition. Mice underwent a behavioral battery consisting of the open field assay, NOR, and CFC. Procedures took place from 5 – 6 months of age in all WT-TRAP and 5X-TRAP mice. Mice that were single housed were excluded from NOR and CFC.

#### Open field assay

Spontaneous locomotion and rearing activity were assessed in an open field arena (14” x 14” x 18”). The Photobeam Activity System (San Diego Instruments) was used to acquire data. The arena is fitted with a scaffold of IR emitters and detectors to collect peripheral, center and vertical (rearing) beam breaks. After a thirty-minute habituation to the testing room, a ten-minute trial began with a mouse placed in the center of the arena. All trials were recorded by high-definition camcorders. Digitally recorded trials were processed for automated analysis by ANYmaze software (Stoelting Co, Il) to obtain additional measures.

#### Novel object recognition

The novel object recognition procedure consisted of habituation/pre-exposure, acquisition and recall trials. Before habituation, mice were handled daily for two minutes for three consecutive days. The habituation/pre-exposure phase consists of daily five-minute trials where the mouse is allowed to freely explore the empty NOR arena (approx. 1 square foot). During the acquisition trial, mice are returned to an arena containing a pair of the same objects. The object pairs used are either glass bottles or metal bar (2”x 2” x five”) placed about three inches from the arena walls. Mice freely explore the object pairs for fifteen minutes. The arena and objects are cleaned with 70% EtOH between all trials. 24 hours after acquisition, mice returned to the arena, with one of the now-familiar objects replaced with a novel object (for example a bottle may be swapped with a metal bar or vice versa). During the recall trial, mice explore the two objects for 15 minutes. All trials are digitally recorded for automated off-line grading by ANYmaze software (Stoelting Co, Il). Time spent exploring (approaches and sniffing) and was determined as the primary dependent variable. Mice that did not explore each object for a minimum of 10 seconds during learning and recall were excluded from analysis.

#### Contextual fear conditioning

Mice were habituated to experimenter handling for three days prior to acquisition. For acquisition, mice were placed in a soundproof conditioning chamber (Med Associates, Fairfax, VT) for 4.5 minutes, during which time they received to unsignaled foot shocks (1 mA, 2s) at 1-minute intervals. For recall trials, mice were returned to the conditioning chamber for 5 minutes at 1 day (recent memory) and 14 days (remote memory) after acquisition, without any stimulation. Behavior was recorded during all trials and freezing behavior (lack of movement except for breathing) was assessed with FreezeScan software (Clever Systems, Reston, VA). Results from recall trials were analyzed relative to baseline freezing levels while the mice were in the recording chamber prior to foot shock during the acquisition trial to control for baseline differences in freezing behavior.

### Euthanasia

Mice were anesthetized by I.P. injection of pentobarbital (50 mg/kg) (Sagent Pharmaceuticals, Schaumburg, IL) and underwent cardiac perfusion with ice cold 4% paraformaldehyde.

### Immunohistochemistry

Brains were incubated at 4°C for 24 h in 4% PFA, followed by sequential incubations in 10%, 20%, and 30% sucrose at 4°C. Samples were then cryoprotected in OCT Tissue Tek and sectioned at 20 µm. Immunohistochemistry (IHC) was performed using standard, previously published protocols (24). Briefly, slices were permeabilized (0.02% triton-X100 in PBS) for 30 minutes, blocked for 1 hour (5% normal goat serum, 0.01% triton X-100, 1% bovine serum albumin), and incubated with primary antibody (see Table S6) in blocking solution overnight at 4°C and for one hour with secondary antibodies Alexa Fluor (Thermo Fisher Scientific, Waltham MA) at a concentration of 1:1000.

### Fluorojade-B

Cryosectioned (20 µm) tissue was stained for fluorojade-B following manufacturer’s instructions (EMD Millipore, Burlington, MA). Briefly, slides were sequentially incubated in 80% ethanol for 5 min, 70% ethanol for 2 min, distilled water for 2 min, 0.06% KMnO4 for 15 min, distilled water for 2 min, 0.001% fluorojade-B solution for 20 min, and distilled water for 1 min (3 times). Images were acquired using a Nikon Eclipse 80i microscope and a digital Nikon DS-Fi2 camera (Micro Video Instruments, Avon MA) with a 20X objective. Fluorojade B+ cells were counted using Image J.

### Slide scanning and quantifications

Fluorescent slide scanning for sectioned tissue was performed with a 3DHISTECH Lamina Scanner (Perkin Elmer, Waltham, MA) with a 20X objective. We used NeuroInfo software (MBF, Williston, VT) to register brain slices to the Allen Brain Atlas (ABA) and detect and map tau AT8+ immunoreactive aggregates and tdT+ and tdT-/NeuN+ neuronal populations. All AT8+ immunoreactive aggregates and tdT+ neurons were detected. Given that NeuN staining was inconsistent within samples, only tdT-/NeuN+ cells with sharp, defined edges were detected providing a representative sampling of tdT-/NeuN+ cells for each region. Data containing each detected object were exported from NeuroInfo and analyzed in R. Each object was summed through the ABA hierarchy (e. g. Somatosensory area and auditory area counts were added to the isocortex counts). Brain regions that contained imperfections (e.g. folds, hole in tissue) were excluded from analysis for that sample. Only brain regions with at least six samples per group were included for analysis. Additionally, for cell type (tdT+/tdT-) colocalization with tau pathology, regions were excluded in samples that did not contain at least 8 tdT+ and 8 tdT-cells. ABA heatmaps displayed in Figures 1, 2, and 3 were generated using Mouse Brain Heatmap (59).

### Computational analysis

As in prior work (43), we used linear diffusion models to predict the spatial distribution of tau pathology, *x*(*t*), across *N* regions defined by the Allen Brain Atlas, using an *N* x *N* adjacency matrix *A*, whose edge weights are the directed structural connectivity strength from region *i* to region *j.* Connectivity strength was defined by fluorescence intensity from retrograde viral tract tracing (44). We computed the estimated regional tau pathology *x^^^*(*t*) using an equation of the form *x^^^*(*t*) = *e*^−cLt^*x*_0_, where *t* is 3 months post injection, *x*_0_ is a vector with 1 unit of pathology in the hippocampal and cortical injection sites, *L* is the out-degree Laplacian of *A*, and *c* is a free parameter obtained through model fitting. The injection sites were the right dentate gyrus, CA1, CA3, and posterior parietal association area. We generated distributions of model fit, i.e. the spatial correlation between *x*(*t*) and *x^^^*(*t*), using bootstrapped samples of mice to compare models where *A* is defined by retrograde connectivity, anterograde connectivity, or interregional Euclidean distance. Finally, we used multivariate regression to construct a “Bidirectional Euclidean tdT” model which combined predicted anterograde spread, retrograde spread, and Euclidean distance-based spread from optimized models with tdT positive cell counts using the following form: *x*(*t*) = β_a_ *x^^^*(*t*)_a_ + β_r_*x^^^*(*t*)_r_ + β_e_(*x^^^*(*t*)_e_ + β_T_T + ε, where *x^^^*(*t*)_&_ is predicted anterograde spread, *x^^^*(*t*)_’_ is predicted retrograde spread, *x^^^*(*t*)_’_ is spread based on Euclidean distance, T is a vector of tdT positive cell counts, and ε is an error term. Distributions of bootstrapped model fits or regression beta weights were compared using a two-tailed non-parametric test (60).

### Human subjects

All human procedures and protocols were approved by the ethical standards of the Institutional Review Board (IRB) at the University of Pennsylvania. Subjects were diagnosed with AD based on clinical history, and neurological and neuropsychological assessment. Clinical diagnoses were confirmed post-mortem based on staging of Aβ_42_ and tau pathology. For assessment of tau pathology distribution in AD patients and control subjects, numeral ratings across 18 brain regions were retrieved from the Center for Neurodegenerative Disease Research (CNDR) database at the University of Pennsylvania. The frequency of tau pathology was scored by expert pathologists using a standard semi-quantitative scale: 0 (absent), 1 (sparse), 2 (moderate), and 3 (frequent). Clinical data from these patients are reported in Table S7.

### Statistics

AT8+ and tdT+ mapping data were first analyzed by linear mixed models with genotype and treatment as fixed effects and brain region (subregion level) and subject as random effects and group contrasts were compared by EMM. We then performed hierarchical Bayesian modeling separately for each of the major parent brain regions (isocortex, hippocampus, cortical subplate, striatum, thalamus, hypothalamus, midbrain, and fiber tracts) incorporating subregions nested within parent regions as random effects. In major brain structures where credible genotype, treatment or genotype x treatment effects were found, we performed additional Bayesian statistical modeling with genotype, treatment, and brain subregion as fixed effects to determine genotype, treatment, and interaction effects at individual brain subregions. Where credible genotype, treatment, or genotype x treatment effects were found, we computed EMMs to determine group level differences for each identified brain region. For Bayesian statistics, credible effects and contrasts were identified when 95% credible intervals did not include 0.

Results from Bayesian modeling and group contrasts can be found in Tables S1 and S2. For all Bayesian models, weakly informative priors based on pilot studies were used to indicate interactions between 5XFAD genotype and PTZ kindling to increase AT8+ and tdT+ levels. Details of Bayesian models, including convergence and effective sample sizes can be found in Table S8. Daily maximal Racine scores were compared between WT-TRAP and 5X-TRAP mice by repeated measure 2-way ANOVA with PTZ administration (repeated measure) and genotype (WT-TRAP vs 5X-TRAP) as independent variables with Sidak’s post hoc to identify group differences. Area under the curve of Racine scores were compared between WT-TRAP and 5X-TRAP mice by two-tailed t-test. Behavior comparisons were made by 2-way ANOVA with genotype (WT-TRAP vs 5X-TRAP) and kindling (saline vs PTZ) as independent variables with Sidak’s post hoc to identify group differences when main effects or interactions were found.

Correlations between regional AT8+/tdT+ counts and behavior were identified by Pearson correlation. For AT8+ colocalization experiments (Fig 6, S15), WT-TRAP and 5X-TRAP were analyzed separately by 2-way ANOVA with kindling (saline vs PTZ) and cell type (tdT+ vs tdT-) as independent variables for each sampled brain region with Tukey’s post hoc where main effects or interactions were found. Human pathological ratings were compared by two-tailed Mann-Whitney test. Males and females were included in all experiments. Sex was added as an independent variable for each dataset and analyzed by multiple or ordinal linear regressions and are included in Table S3 and S5. Homogeneity of variance was confirmed for all 2-way ANOVA analyses. Statistical analyses for brain mapping, AT8 colocalization, and ordinal regressions were performed in R (4.3.2). All other statistical analyses were performed in Prism 10 (GraphPad, Boston, MA). Multiple comparisons were adjusted where appropriate as described above. All statistical tests were two-sided.

## Data availability

The data that support the findings of this study are available upon reasonable request from the corresponding authors.

## Code availability

The code used for computational analyses is available at https://github.com/ejcorn/tau-seizures.

## Acknowledgements

These studies were funded by the National Institutes of Health (NIH) National Institute on Aging (NIA) T32AG000255 (AJB) and R01AG077092 (FEJ and DMT), the Alzheimer’s Association AARF-22-972333 (AJB), the NIH National Institute of Neurological Disorders and Stroke (NINDS): R01NS101156 (DMT) and R37NS115439 (FEJ). Postmortem tissue and clinical data from control and Alzheimer’s disease was collected with funding from the NIH NIA: P30AG072979, P01AG084497, and R01AG054519.

We would like to thank The Neurobehavior Testing Core at UPenn/ITMAT and IDDRC at CHOP/Penn U54 HD086984 for assistance with the behavior procedures. We also thank the Center for Neurodegenerative Disease Research (CNDR) at the University of Pennsylvania for providing human Alzheimer’s disease and control samples and relevant clinical and neuropathology information. We also would like to thank Lakshmi Changolkar for her work to isolate the AD-tau used in these studies. Schematics in Figures 1 and 8 were created in BioRender.

## Author Contributions

AJB, VMYL, DMT, and FEJ conceptualized the studies in the manuscript. AJB, KH, AC, XL, SZ, and CH performed experiments, analyzed data, and prepared figures. EJC, AL, and KAD performed computational analyses and prepared figures. EBL performed pathological ratings for human postmortem analysis and assisted with clinical data reports. AJB, DMT, and FEJ wrote the manuscript. All authors read and approved the manuscript.

## Notes

### Competing Interest Statement

The authors have declared no competing interest.

### Summary of Updates

Updated figures, new data analyses, and expanded discussion.

