## Supplementary Material for "Hyperactive neuronal networks enhance tau spread in an Alzheimer’s disease mouse model"

### Supplemental material

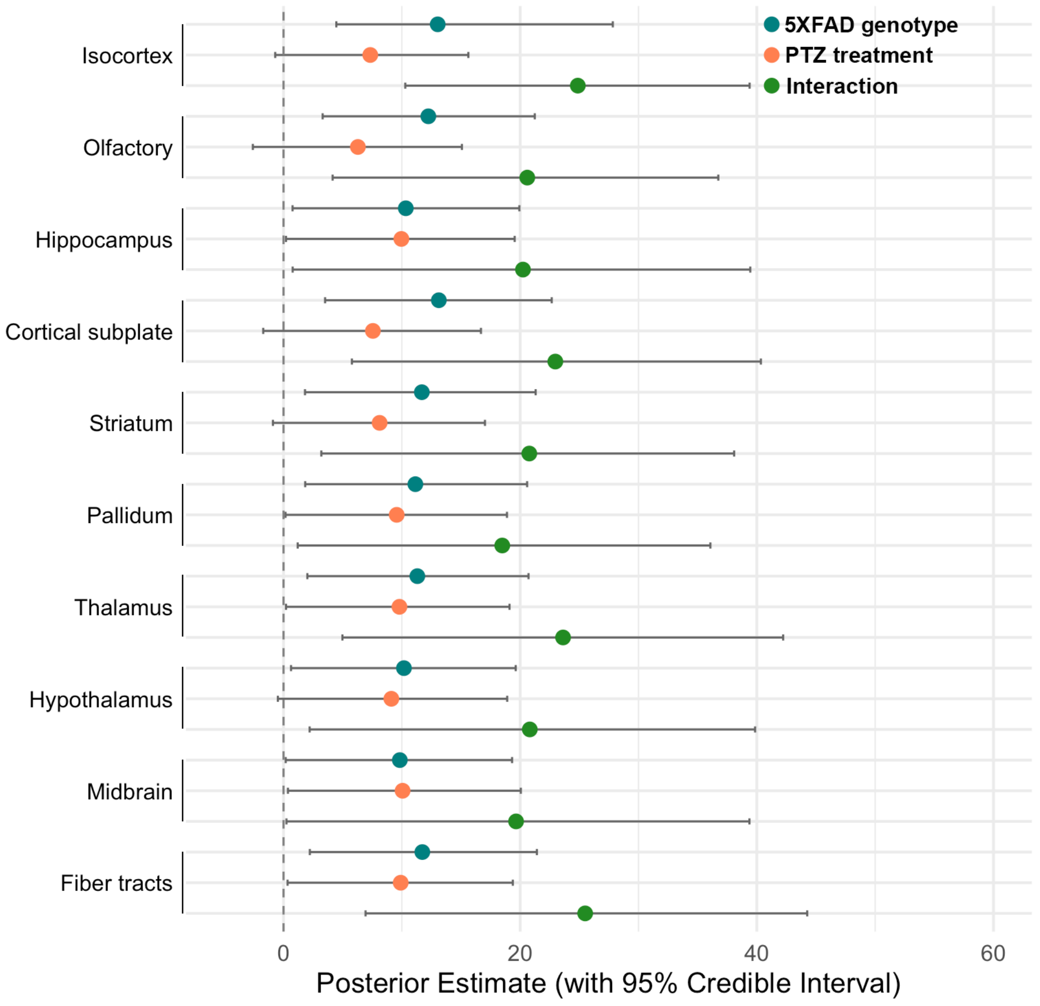

**Figure S1. Hierarchical Bayesian models demonstrate interactions between PTZ treatment and 5XFAD genotype increase tau pathology in 10 major brain structures.** Posterior means (points) and 95% credible intervals (bars) were estimated using Bayesian models for AT8+ levels including genotype, treatment, and their interactions as fixed effects, and animal and the region (nested within parent structure) as random effects. Each point represents the estimated effect on tau pathology (AT8+/$mm^2$) within the given brain region. Effects with 95% credible intervals that do not cross zero are considered to have credible differences. Interactions between 5XFAD genotype and PTZ kindling credibly elevated AT8+ levels in all 10 major brain structures examined.

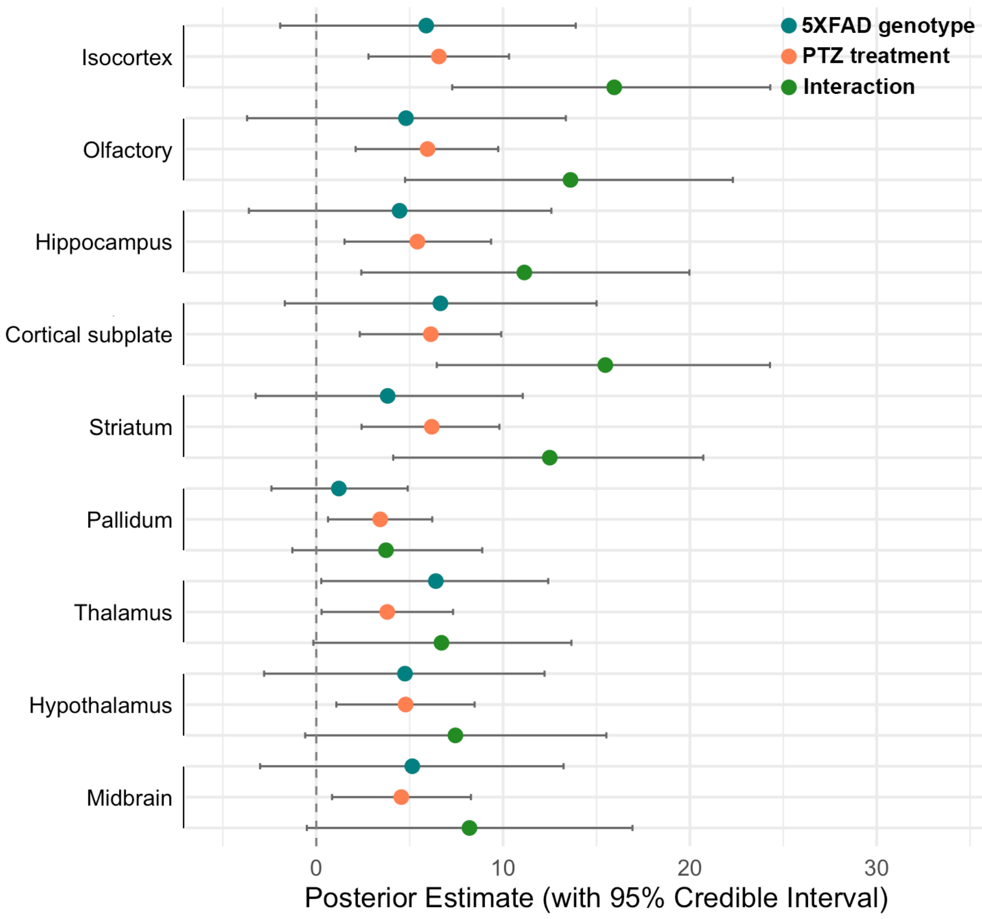

**Figure S2. Hierarchical Bayesian models demonstrate that PTZ and its interactions with 5XFAD genotype increase neuronal activity in 9 major brain structures.** Posterior means (points) and 95% credible intervals (bars) were estimated using Bayesian models for tdT+ counts including genotype, treatment, and their interactions as fixed effects, and animal and the region (nested within parent structure) as random effects. Each point represents the estimated effect on tdT+ counts (tdT+/mm$^2$) within the given brain region. Effects with 95% credible intervals that do not cross zero are considered to have credible differences. PTZ kindling credibly elevated tdT+ cells across all 10 parent brain regions and interacts with 5XFAD genotype to increase tdT+ cells in the isocortex, olfactory areas, hippocampus, cortical subplate, and striatum.

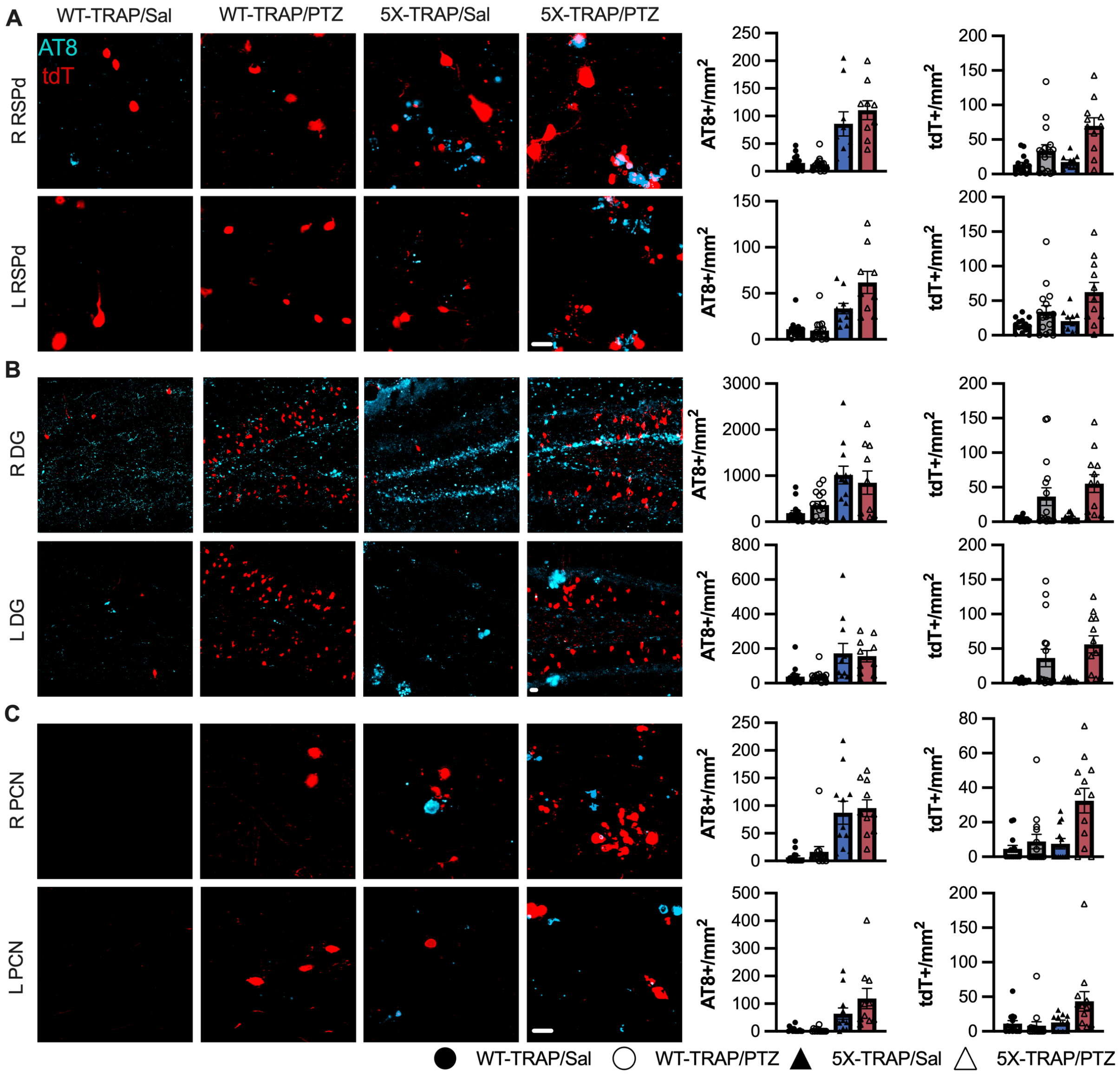

**Figure S3. Representative staining and histograms from AT8 and tdT mapping.**

Representative images and histograms from AT8 and tdTomato (tdT) brain mapping (Figure 1, Table S1, S2). AT8 IHC (cyan) and tdT (red) from all experimental groups in the ipsilateral (R) and contralateral (L) (to tau injection site) (**A**) dorsal retrosplenial cortex (RSPd), (**B**) dentate gyrus (DG), and (**C**) paracentral nucleus of the thalamus. N=9-16/group/brain region. Scale bars = 20 μm.

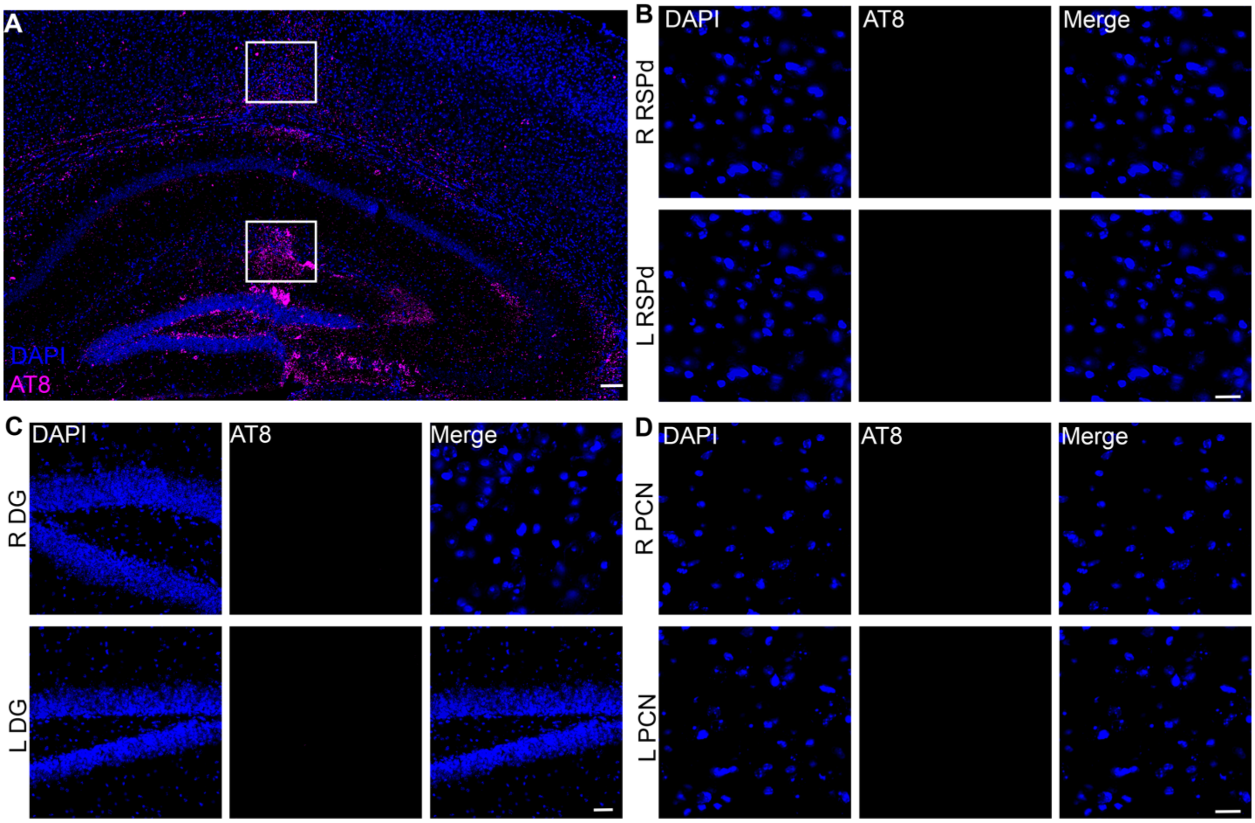

**Figure S4. Validation of AD-tau injection site and AT8 specificity.** (**A**) Representative AT8 (magenta) and DAPI (blue) image showing AD-tau injection sites (boxes) into the dorsal hippocampus and overlying cortex. Representative images of control sections omitting the primary antibody in the ipsilateral (R) and contralateral (L) (to AD-tau injection) (**B**) dorsal retrosplenial cortex (RSPd), (**C**) dentate gyrus (DG), and (**D**) paracentral nuclei of the thalamus from AD-tau seeded and slide scanned sections. Scale bars = 100 μm (**A**), 20 μm (**B, D**), 50 μm (**C**).

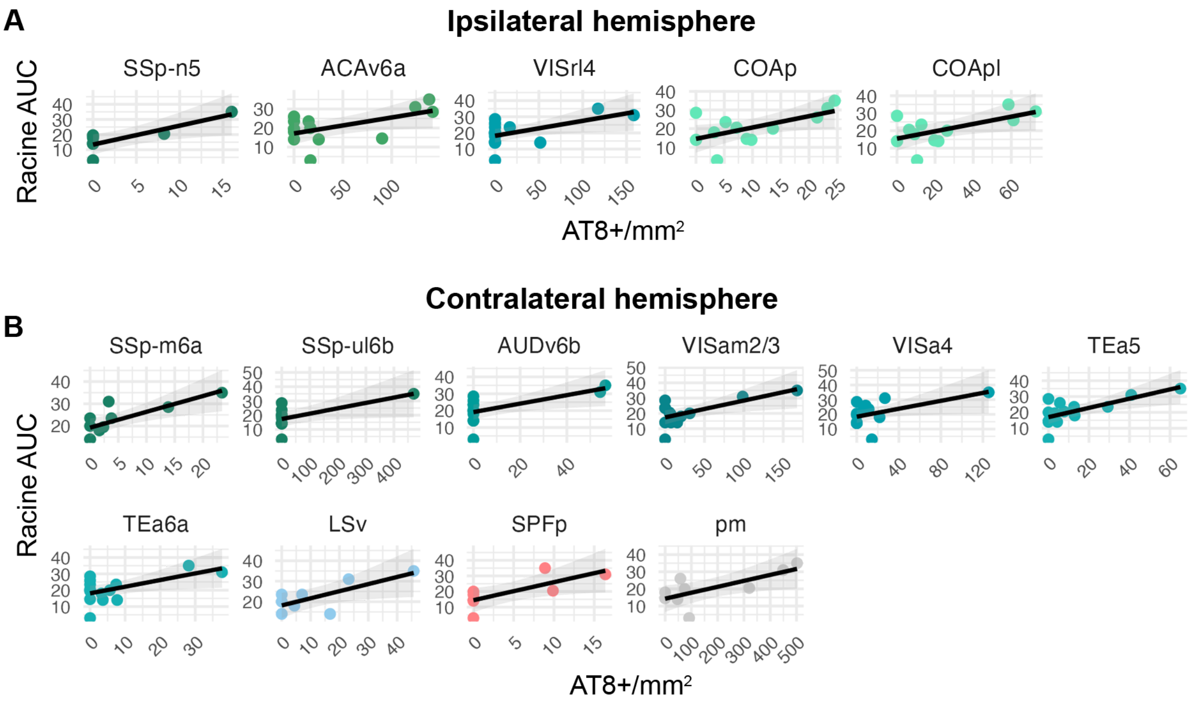

**Figure S5. Seizure severity is correlated with tau spread in WT-TRAP mice.** Pearson correlations were performed between area under the curve (AUC) for Racine score in the 15 min following the final PTZ administration and regional AT8+ immunoreactive aggregate levels in WT-TRAP mice. We found significant positive correlations in the (**A**) ipsilateral (to AD-tau injection) (right) isocortex: primary somatosensory area, nose, layer 5 (SSp-n5, r=0.82, p=0.019, n=6), anterior cingulate area, ventral part, layer 6a (ACAv6a, r=0.57, p=0.044, n=12), rostrolateral area, layer 4 (VISrl4, r=0.57, p=0.028, n=13), and cortical suplate: cortical amygdalar area posterior part (COAp, r=0.61, p=0.037, n=11). (**B**) Significant positive correlations were found in contralateral (left) isocortex: SSP, mouth, layer 6a (SSP-m6a, r=0.78, p=0.0076, n=9), SSp, upper limb, layer 6b (SSP-ul6b, r=0.64, p=0.048, n=9), ventral auditory area, layer 6b (AUDv6b, r=0.63, p=0.021, n=10), anteromedial visual area, layer 2/3 (VISam2/3, r=0.66, p=0.015, n=12), anterior area, layer 4 (VISa4, r=0.54, p=0.040, n=12), temporal association area, layer 5 (TEa5, r=0.69, p=0.0064, n=14) and 6a (TEa6a, r=0.58, p=0.030, n=14), striatum: lateral septal nucleus, ventral part (LSv, r=0.74, p=0.037, n=8), thalamus: subparafascicular nucleus, parvicellular part (SPFp, r=0.74, p=0.035, n=8), and fiber tracts:

primary mammillary tract (pm, r=0.71, p=0.032, n=9). Shaded areas display 95% confidence interval.

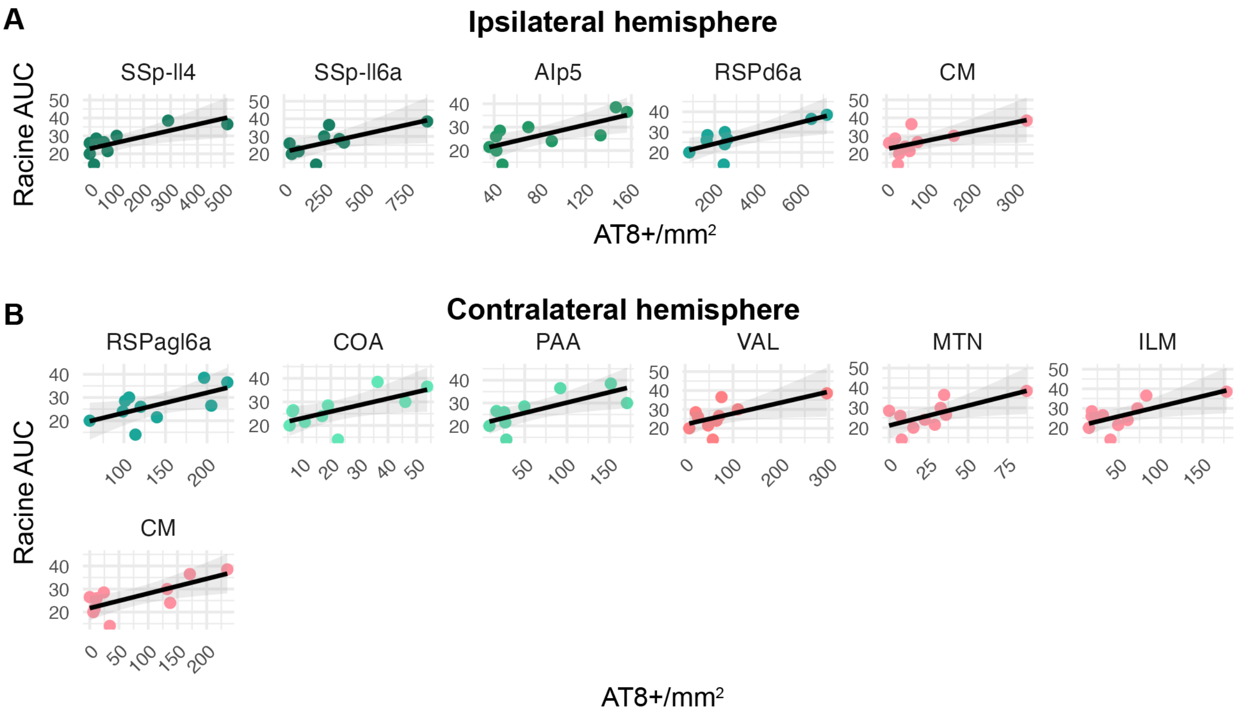

**Figure S6. Seizure severity is correlated with tau spread in 5X-TRAP mice.** Pearson correlations were performed between area under the curve (AUC) for Racine score in the 15 min following the final PTZ administration and regional AT8+ immunoreactive aggregate levels in 5X-TRAP mice. We found significant positive correlations in the (**A**) ipsilateral (to AD-tau injection) (right) isocortex: primary somatosensory area, lower limb layer 4 (SSp-ll4, r=0.76, p=0.041, n=9) and layer 6a (SSp-ll6a, r=0.68, p=0.046, n=9) agranular insular area, posterior part layer 5 (AIp5, r=0.73, p=0.015, n=11), and dorsal retrosplenial area layer 6a (RSPd6a, r=0.77, p=0.015, n=9), and the thalamus: central medial nucleus (CM, r=0.65, p=0.043, n=9). (**B**) Significant positive correlations were found in the contralateral (left) isocortex: retrosplenial area, lateral agranular part, layer 6a (RSPagl6a, r=0.64, p=0.047, n=10), olfactory areas: cortical amygdalar area (COA, r=0.65, p=0.043, n=9), piriform amygdalar area (PAA, r=0.72, p=0.019, n=9), and thalamus: ventral anterior-lateral complex (VAL, r=0.64, p=0.045, n=10), midline group of the dorsal thalamus (MTN, r=0.67, p=0.035, n=10), intralaminar nuclei of the dorsal thalamus (r=0.69, p=0.029, n=10), and CM (r=0.73, p=0.014, n=9). Shaded areas display 95% confidence interval.

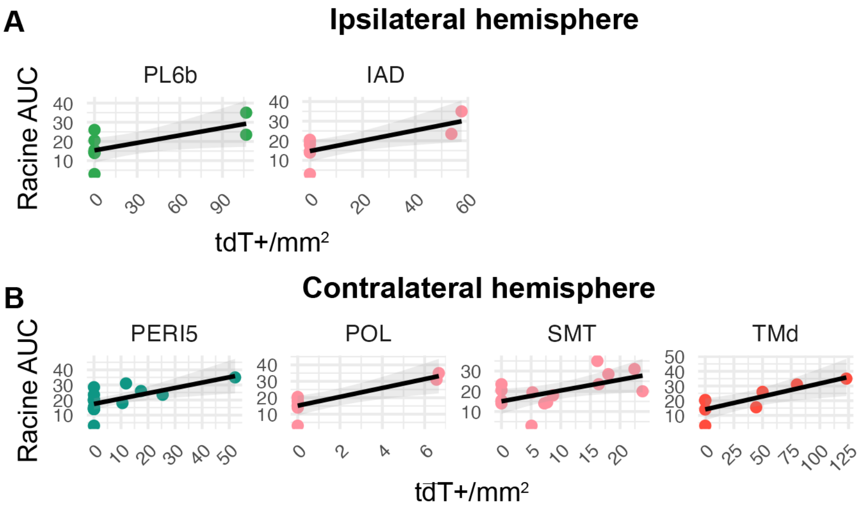

**Figure S7. Seizure severity is correlated with activity dependent tdTomato labeling in WT-TRAP mice.** Pearson correlations were performed between area under the curve (AUC) for Racine score in the 15 min following the final PTZ administration and regional tdT+ counts in WT-TRAP mice. (**A**) We found significant positive correlations in the ipsilateral (to AD-tau injection) (right) isocortex: prelimbic area layer 6b (PL6b,) and thalamus: interanterodorsal nucleus (IAD, r=0.75, p=0.020, n=8). (**B**) Significant positive correlations were found in the contralateral (left) isocortex: perirhinal area, layer 5 (PERI5, r=0.64, p=0.0075, n=13), thalamus: posterior limiting nucleus (POL, r=0.83, p=0.011, n=8), submedial nucleus (SMT, r=0.56, p=0.038, n=13) and hypothalamus: tuberomammillary nucleus, dorsal part (TMd, r=0.80, p=0.017, n=8). Shaded areas display 95% confidence interval.

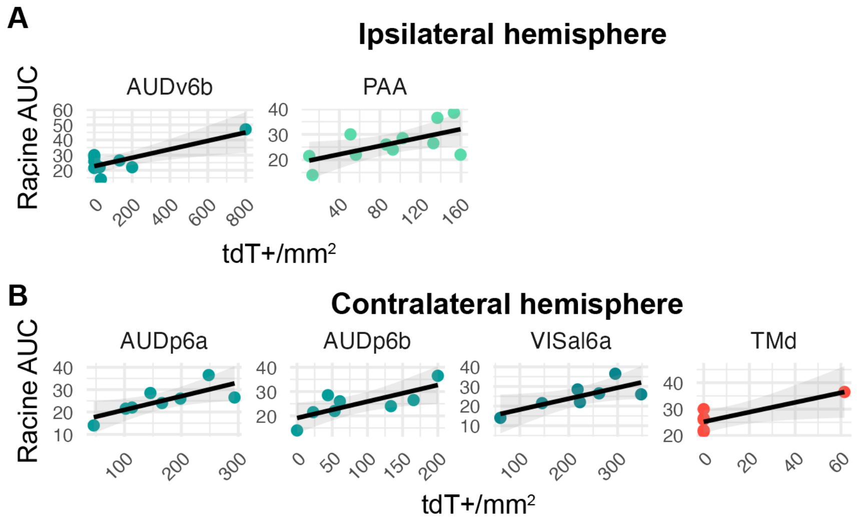

**Figure S8. Seizure severity is correlated with activity dependent tdTomato labeling in 5X-TRAP mice.** Pearson correlations were performed between area under the curve (AUC) for Racine score in the 15 min following the final PTZ administration and regional tdT+ counts in WT-TRAP mice. (**A**) We found significant positive correlations in the ipsilateral (to AD-tau injection) (right) isocortex: ventral auditory area, layer 6b (AUDv6b, r=0.80, p=0.0098, n=9), and olfactory areas: piriform amygdalar area (PAA, r=0.62, p=0.040, n=11). (**B**) Significant positive correlations were found in the contralateral (left) isocortex: primary auditory area layer 6a (AUDp6a, r=0.77, p=0.026, n=8) and 6b (AUDp6b, r=0.76, p=0.029, n=8), and anterolateral visual area layer 6b (VISal6a, r=0.77, p=0.44, n=7), and hypothalamus: tuberomammilary nucleus, dorsal part (TMd, r=0.83, p=0.42, n=6). Shaded areas display 95% confidence interval.

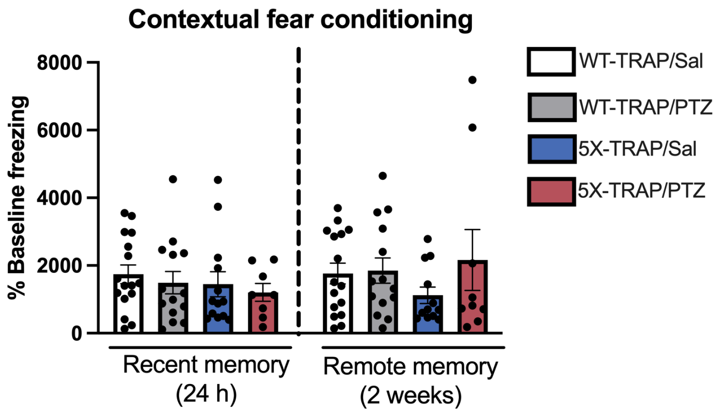

**Figure S9. Results from contextual fear conditioning.** Results are displayed as % change from baseline freezing levels to control for baseline differences in motor activity. Recent (1 day after conditioning) and remote (2 weeks after conditioning) recall were analyzed separately by 2-way ANOVA with genotype (WT-TRAP vs 5X-TRAP) and kindling (PTZ vs Sal) as independent variables. No significant effects were found. n=9-16/group. Histograms display group mean ± SEM.

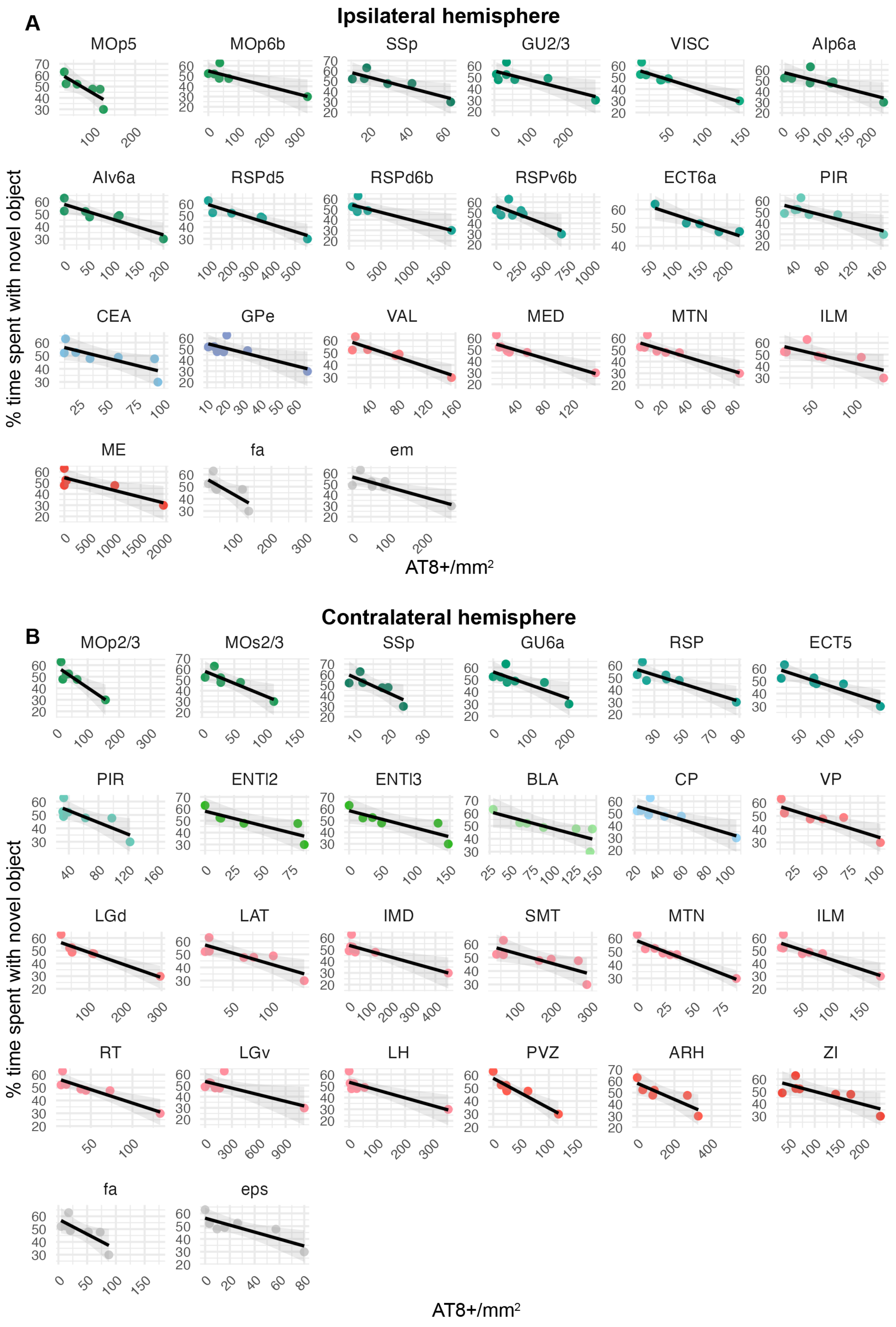
A
Ipsilateral hemisphere
MOp5
MOp6b
SSp
GU2/3
VISC
AIp6a
AIv6a
RSPd5
RSPd6b
RSPv6b
ECT6a
PIR
CEA
GPe
VAL
MED
MTN
ILM
ME
fa
em
% time spent with novel object
AT8+/mm2
B
Contralateral hemisphere
MOp2/3
MOs2/3
SSp
GU6a
RSP
ECT5
PIR
ENTl2
ENTl3
BLA
CP
VP
LGd
LAT
IMD
SMT
MTN
ILM
RT
LGv
LH
PVZ
ARH
ZI
fa
eps
% time spent with novel object
AT8+/mm2

**Figure S10. Memory deficits are correlated with tau spread in 5X-TRAP mice.** Pearson correlations were performed between % time spent with novel object during the novel object recognition (NOR) task and regional AT8+ levels in 5X-TRAP mice. (**A**) We found significant negative correlations in the ipsilateral (to AD-tau injection) (right) isocortex: primary motor area, layers 5 (MOp5, r=-0.82, p=0.044, n=6),and 6b (MOp6b, r=-0.86, p=0.027, n=6) primary somatosensory area, overall (SSp, r=-0.87, p=0.027, n=6), gustatory area, layer 2/3 (GU2/3, r=-0.81, p=0.029, n=7) visceral area, overall, (VISC, r=-0.93, p=0.0028, n=7), posterior and ventral agranular insular area, layer 6a (AIp6a, r=-0.79, p=0.031, n=7) (AIv6a, r=-0.90, p=0.0061, n=7), dorsal retrosplenial area, layers 5 and 6b (RSPd5, r=-0.93, p=0.0068, n=7) (RSPd6b, r=-0.87, p=0.022, n=7), ectorhinal area, layer 6a (ECT6a, r=-93, p=0.020, n=6), olfactory area: piriform area (PIR, r=-0.81, p=0.027, n=7), striatum: central amygdalar nucleus (CEA, r=-0.77, p=0.042, n=7), pallidum: external globus pallidus (GPe, r=-0.80, p=0.032, n=7), thalamus: ventral anterior lateral complex (VAL, r=-0.93, p=0.0030, n=7), medial group of the dorsal thalamus (MED, r=-0.91, p=0.0034, n=7), midline group of the dorsal thalamus (MTN, r=-0.90, p=0.0057, n=7), intralaminar nuclei of the dorsal thalamus (ILM, r=-0.76, p=0.045, n=7), hypothalamus: median eminence (ME, r=-0.86, p=0.026, n=6), and fiber tracts: corpus callosum anterior forceps (fa, r=-0.76, p=0047, n=7), external medullary lamina of the thalamus (em, r=-0.85, p=0.016, n=7). (**B**) Significant negative correlations were found in contralateral (left) isocortex: MOP2/3 (r=-0.91, p=0.012, n=6) secondary motor cortex layers 2/3 (MOs2/3, r=-0.88, p=0.022, n=6), SSp r=-0.82, p=0.047, n=6), GU6a (r=-0.81, p=0.028, n=6), RSP, overall (r=-0.88, p=0.0091, n=7)), ECT5 (r=-0.90, p=0.006, n=7), PIR (r=-0.82, p=0.023, n=7), entorhinal area, lateral part layer 2 and 4 (ENTl2, r=-0.83, p=0.039, n=6) (ENTl3, r=-0.85, p=0.034, n=6), cortical subplate: basolateral amygdala (BLA, r=-0.80, p=0.029, n=7), striatum: caudoputamen (CP, r=-0.85, p=0.013, n=7), thalamus: ventral posterior complex (VP, r=-0.88, p=0.0087, 6), dorsal lateral geniculate complex (LGd, r=-0.93, p=0.0030) lateral group of the dorsal thalamus (LAT, r=-0.85, p=0.015, n=6), intermediodorsal nucleus (IMD, r=-0.87, p=0.011, n=7), submedial nucleus (SMT, r=-0.89, p=.037, n=7), MTN (r=-0.97, p=0.00050, n=7), ILM (r=-0.91, p=0.0038, n=7), reticular nucleus (RT, r=-0.91, p=0.0043, n=7), ventral lateral geniculate complex (LGv, r=-0.78, p=0.039, n=7), lateral habenula (LH, r=-0.88, p=0.0091, n=7), hypothalamus: periventricular zone (PVZ, r=-0.93, p=0.0024, n=7), arcuate hypothalamic nucleus (ARH, r=-0.86, p=0.029, n=6), zona incerta (ZI, r=-0.79, p=0.034, n=7), and fiber tracts: fa (r=-0.77, p=0.042, n=7), extrapyramidal fiber system (eps, r=-0.83, p=0.020, n=7). Shaded areas display 95% confidence interval.

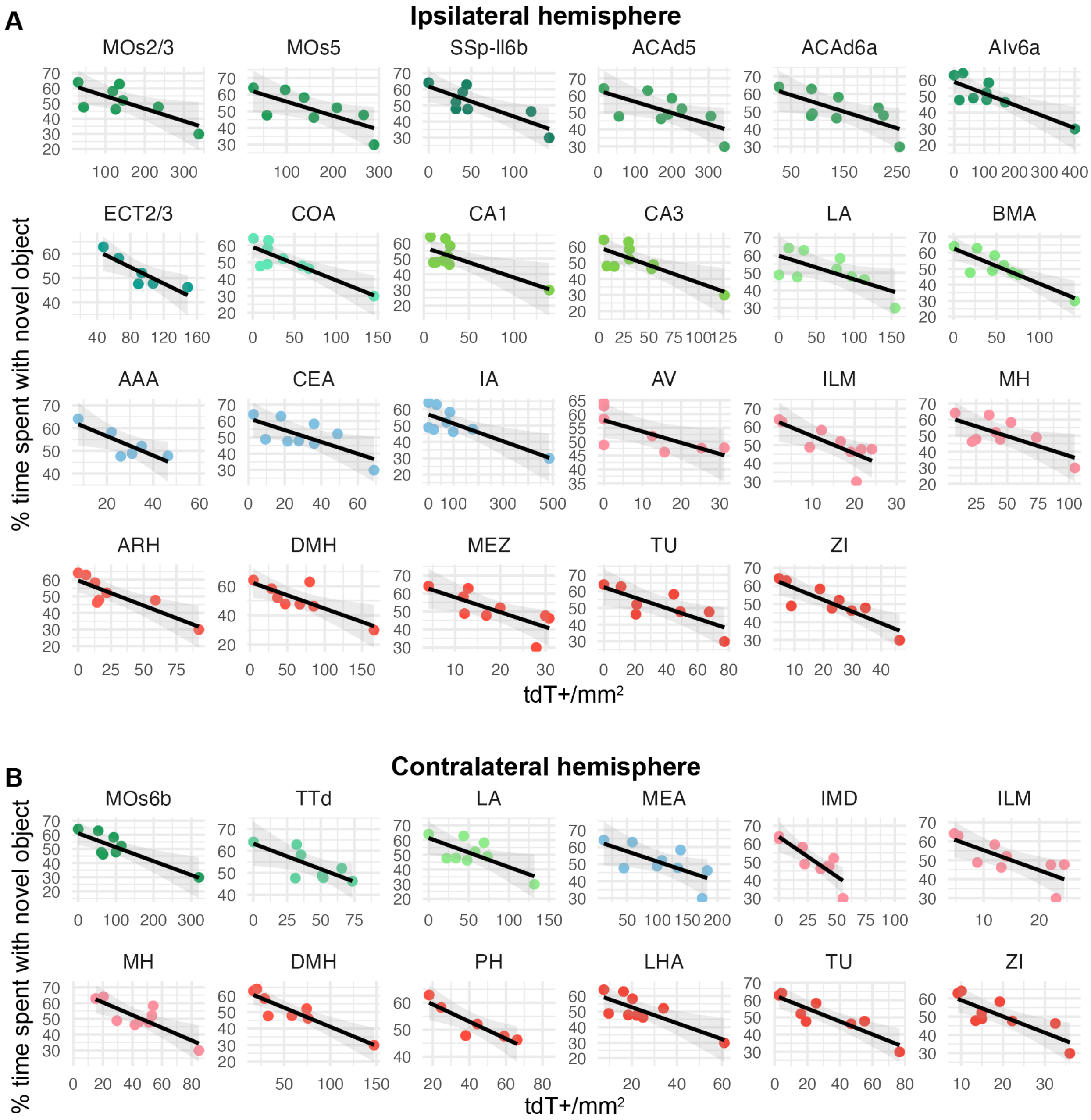

**Figure S11. Memory deficits are correlated with activity dependent tdTomato labeling in 5X-TRAP mice.** Pearson correlations were performed between % time spent with novel object during the novel object recognition (NOR) task and regional tdT+ counts in 5X-TRAP mice. We found significant negative correlations in the ipsilateral (to AD-tau injection) (right) (**A**) isocortex:

secondary motor cortex layers 2/3 (MOs2/3, r=-0.74, p=0.035, n=8) and 5 (MOs5, r=-0.72, p=0.043, n=8), primary somatosensory area, lower limb, layer 6b (SSp-ll6b, r=-0.81, p=0.015, n=8), dorsal anterior cingulate area layer 5 (ACAd5, r=-0.67, p=0.047, n=9) and 6a (ACAd6a, r=-0.69, p=0.037, n=9), ventral agranular insular area, layer 6a (AIv6a, r=-0.83, p=0.0057, n=9), ectorhinal area, layer 2/3 (ECT2/3, r=-0.87, p=0.023, n=6), cortical amygdalar area (COA, r=-0.86, p=0.0032, n=9), CA1 (r=-0.76, p=0.019, n=9), CA3 (r=-0.78, p=0.014, n=9), cortical subplate: lateral amygdala (LA, r=-0.67, p=0048, n=9), basomedial amygdala (BMA, r=-0.86, p=0.0027, n=9), striatum: anterior amygdalar area (AAA, r=-0.83, p=0.042, n=6), central amygdalar area (CEA, r=-0.71, p=0.031, n=9), intercalated amygdalar nucleus (IA, r=-0.80, p=0.0093, n=9), thalamus: anteroventral nucleus of the thalamus (AV, r=-0.71, p=0.048, n=8), intralaminar nuclei of the dorsal thalamus (ILM, r=-0.75, p=0.019, n=9), medial habenula (MH, r=-0.69, p=0.034, n=9), hypothalamus: arcuate nucleus (ARH, r=-0.86, p=0.0060, n=9), dorsal medial nucleus (DMH, r=-0.81, p=0.015, n=9), medial zone (MEZ, r=-0.75, p=0.021, n=9), tuberal nucleus (TU, r=-0.78, p=0.0022, n=8), zona incerta (ZI, r=-0.86, p=0.0027, n=9). (**B**) Significant negative correlations were found in the contralateral isocortex: MOs6b (r=-085, p=0.0079, n=8), olfactory area: dorsal taenia tecta (TTd, r=-0.74, p=0.036, n=8), cortical subplate: lateral amygdala (LA, r=-0.71, p=0.031, n=9), striatum: medial amygdalar nucleus (MEA, r=-0.67, p=0.050, n=9), thalamus: intermediodorsal nucleus (IMD, r=-0.85, p=0.0034, n=9), intralaminar nuclei of the dorsal thalamus (ILM, r=-0.76, p=0.17, n=9), MH (r=-0.81, p=0.0076, n=9), and hypothalamus: DMH (r=-0.91, p=0.0016, n=8), posterior hypothalamic nucleus (PH, r=-0.89, p=0.017, n=6), lateral hypothalamic area (LHA, r=-0.79, p=0.011, n=9), TU (r=-0.89, p=0.0026, n=8), ZI (r=-0.82, p=0.0073, n=9). Shaded areas display 95% confidence interval.

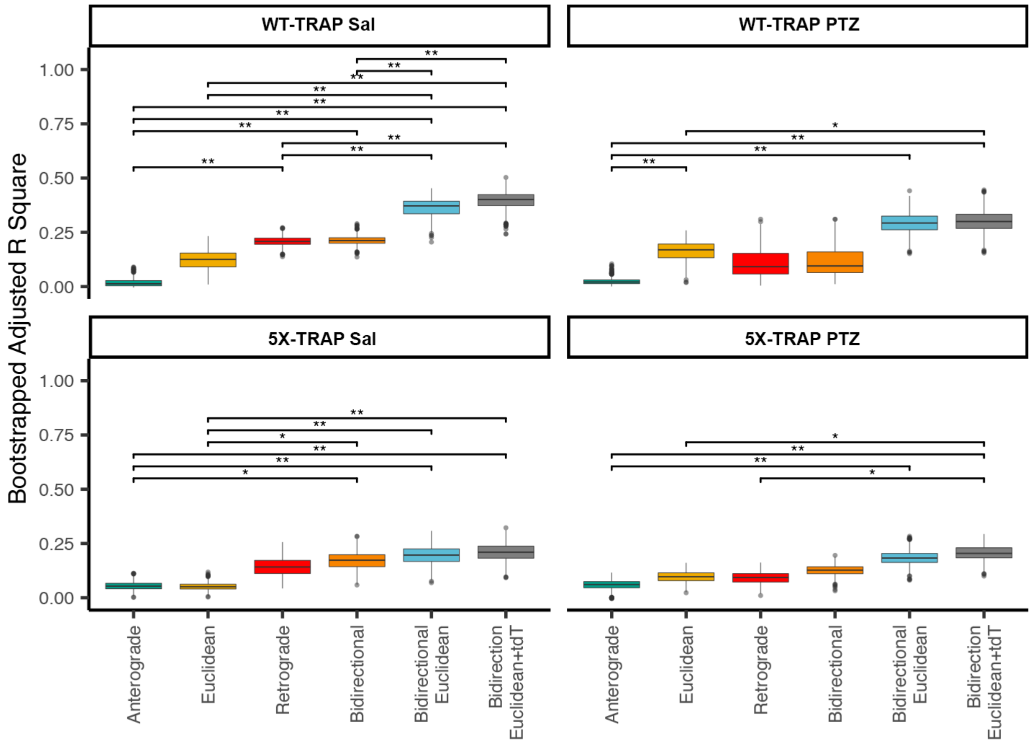

**Figure S12. Between model comparisons of tau spread model fit for each experimental group.** We fit spread models using a network defined by either anterograde connectivity, retrograde connectivity or Euclidean distance between regions. We also generated "bidirectional" models that combined anterograde and retrograde connections ("Bidirectional"), with the addition of Euclidean distance ("Bidirectional Euclidean") and a vector of tdT positive cell counts ("Bidirectional Euclidean + tdT"). We bootstrapped adjusted R square values for each of these model architectures to compare the model fit while accounting for the fact that increased model complexity will always improve training set performance. We compared bootstrapped adjusted R square models between groups for each model architecture. p-values were adjusted for multiple comparisons in each panel using FDR correction. *, $p<0.05$; **, $p<0.01$. Box plots boxes display the median, 25$^{th}$ and 75$^{th}$ percentiles, and whiskers extend to the data point that is at most 1.5 x interquartile range, and data beyond this point are plotted individually.

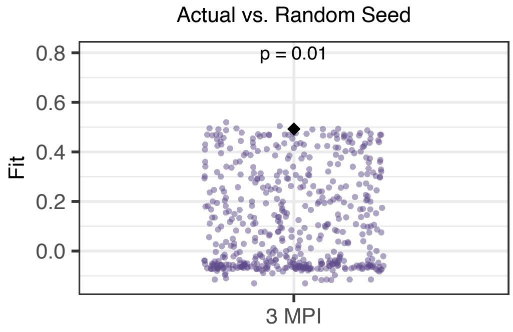

**Figure S13. Null model demonstrating specificity of computational model to experimental seed site.** We repeated the retrograde spread modeling with WT-TRAP mice using 500 random seed sites comprised of 4 seed sites with similar spatial clustering (mean interregional Euclidean distance) to the experimental seed site, which was the right dentate gyrus, CA1, CA3, and posterior parietal association area. Model fits when using random seeds are shown in light purple and model fit when using the experimental seed site is shown as a black diamond. When implementing the experimental seed site into the model in silico, the model performance in predicting regional AT8+ levels was better than 99% of random sites.

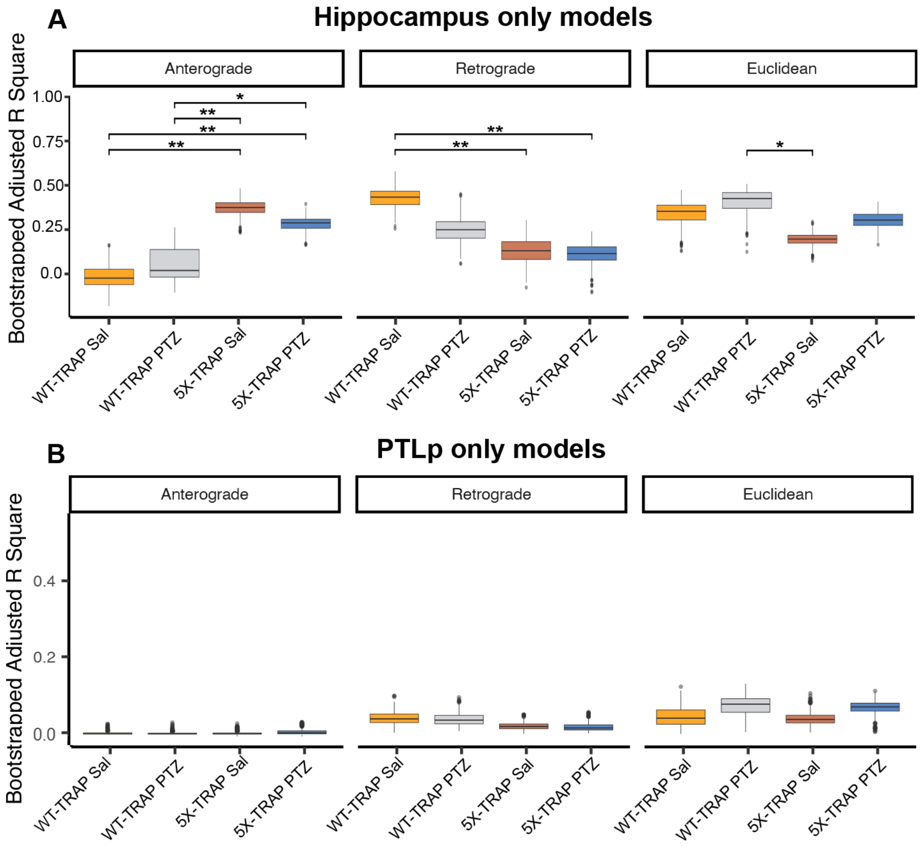

**Figure S14.** Between group comparisons of single site models of tau spread. (**A**) Bootstrapped adjusted R square values of predictions of tau spread based on anterograde, retrograde, and Euclidean models of spread models incorporating a single hippocampal seed site recapitulate results from the dual site models (Fig 4). (**B**) Bootstrapped adjusted R square values of predictions of tau spread based on anterograde, retrograde, and Euclidean models of spread models incorporating a single posterior parietal association cortex (PTLp) seed site perform poorly across all experimental groups. p-values were adjusted for multiple comparisons in each panel using FDR correction: *$p<0.05$, **$p<0.01$. Box plots boxes display the median, 25th and 75th percentiles, and whiskers extend to the data point that is at most 1.5 x interquartile range, and data beyond this point are plotted individually.

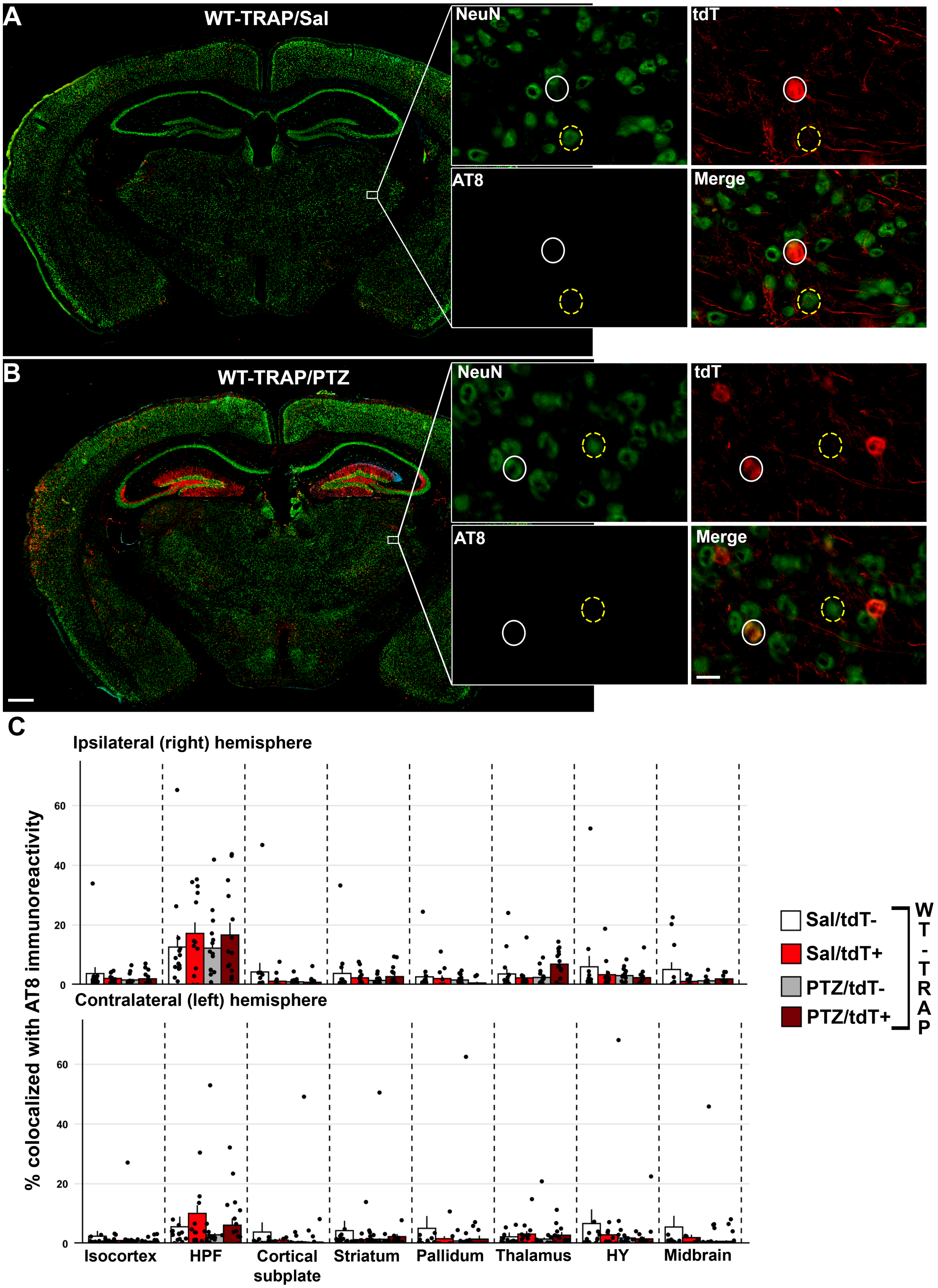

**Figure S15. tdT+ and tdT- neurons colocalization with tau pathology in WT-TRAP mice.**

Representative slide scanned coronal sections from (**A**) saline and (**B**) PTZ treated WT-TRAP mice. Inset displays thalamic tdT+ (white circles) and tdT- (NeuN+) (yellow dashed circles) neurons colocalization with AT8+ immunoreactive aggregates. Scale bars = 500 μm, 20 μm (inset). (**C**) tdT+, tdT-, and colocalized AT8+ immunoreactive aggregates were mapped to the Allen Brain Atlas and % of detected tdT+/tdT- cells were quantified. Two-way ANOVA with treatment (Sal vs PTZ) and tdT labelling (tdT+ vs tdT-) as independent variables were performed. No significant differences were found, n=8-15/group/brain region. Histograms display group mean ± SEM.

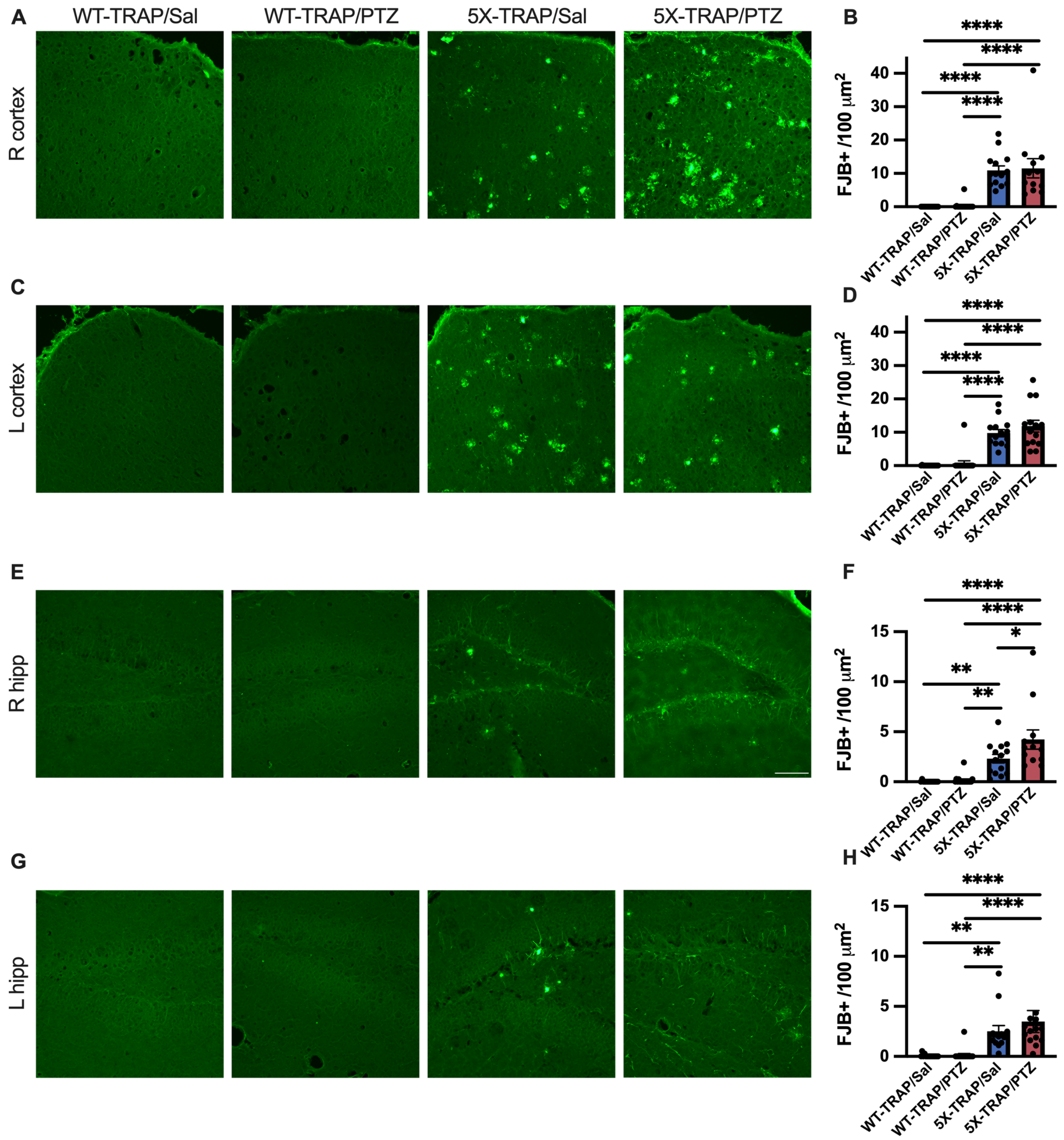

**Figure S16. Neurodegeneration in AD-tau seeded 5X-TRAP mice is exacerbated by PTZ kindling.** Neurodegeneration was assessed by Fluorojade-B (FJB) stain. Representative images of FJB stain from the (**A**) ipsilateral to injection site (right, R) cortex and (**C**) contralateral (left, L) cortex, (**E**) ipsilateral hippocampus, and (**G**) contralateral hippocampus. Scale bar = 50 µm. FJB+ counts were compared by two-way ANOVA with genotype (WT-TRAP vs 5X-TRAP) and kindling (Sal vs PTZ) as independent variables. Increased FJB+ counts were found in Sal and PTZ treated 5X-TRAP mice compared to Sal and PTZ treated WT-TRAP mice in all sampled brain regions ((**B**) R cortex genotype: $F_{1,53}$=62.19, p<0.0001; (**D**) L cortex genotype: $F_{1,56}$=99.55; p<0.0001; (**F**) R hippocampus genotype: $F_{1,53}$=47.40, p<0.0001; (**H**) L hippocampus genotype: $F_{1,56}$=24.78, p<0.0001), and in PTZ kindled 5X-TRAP mice compared to Sal treated 5X-TRAP mice in the R hippocampus (kindling: $F_{1,53}$=4.94, p<0.05). Tukey's post hoc: *,**, ****=p<0.05, 0.01, 0.0001. n=13-17/group. Histograms display group mean ± SEM.

| ID | Parent.ID | Region | WT.Sal | WT.Sal n | WT.PTZ | WT.PTZ n | 5X.Sal | 5X.Sal n | 5X.PTZ | 5X.PTZ n | Genotype effect | Treatment effect | Interaction | WT Sal vs WT PTZ | WT Sal vs WT 5XFAD | WT Sal vs 5XFAD PTZ | 5XFAD Sal vs WT PTZ | WT PTZ VS 5XFAD PT | 5XFAD Sal vs 5XFAD PTZ |
|---|---|---|---|---|---|---|---|---|---|---|---|---|---|---|---|---|---|---|---|
| R 315 | R 695 | R Isocortex | 10.28 ± 2.23 | 15 | 14.04 ± 4.14 | 15 | 23.64 ± 4.35 | 10 | 35.01 ± 3.64 | 10 | 0.94 [-2.29, 4.17] | 0.95 [-2.61, 4.41] | -1.88 [-6.54, 2.86] | | | | | | |
| L 315 | L 695 | L Isocortex | 4.57 ± 1.26 | 15 | 4.93 ± 1.28 | 15 | 11.53 ± 1.94 | 10 | 21.57 ± 4.37 | 10 | 2.53 [-0.78, 5.83] | 2.03 [-1.55, 5.53] | -3.25 [-8.13, 1.45] | | | | | | |
| R 500 | R 315 | R Somatomotor areas | 3.75 ± 0.79 | 15 | 12.15 ± 7.23 | 15 | 18.43 ± 3.96 | 10 | 24.66 ± 5.73 | 10 | 2.21 [-0.98, 5.41] | 1.52 [-2.05, 4.99] | -0.57 [-5.27, 4.02] | | | | | | |
| L 500 | L 315 | L Somatomotor areas | 3.27 ± 0.81 | 15 | 6.12 ± 2.22 | 15 | 17.63 ± 3.46 | 10 | 23.49 ± 6.20 | 10 | 4.20 [0.93, 7.53]* | 1.61 [-1.96, 5.18] | -2.03 [-6.8, 2.69] | -1.04 [-3.97, 1.67] | -4.64 [-7.87, -1.57]* | -6.22 [-9.35, -3.02]* | 3.60 [0.57, 6.8]* | -5.17 [-8.36, -2.03]* | -1.58 [-5.2, 1.77] |
| R 985 | R 500 | R Primary motor area | 6.44 ± 1.29 | 13 | 14.88 ± 7.80 | 15 | 23.84 ± 5.28 | 10 | 35.46 ± 7.70 | 9 | 2.70 [-0.64, 6.1] | 2.26 [-1.35, 5.93] | -2.27 [-7.15, 2.54] | | | | | | |
| L 985 | L 500 | L Primary motor area | 5.22 ± 1.33 | 15 | 7.08 ± 2.35 | 15 | 24.37 ± 5.65 | 10 | 37.62 ± 9.84 | 9 | 5.36 [2, 8.66]* | 2.99 [-0.7, 6.62] | -3.94 [-8.75, 0.92] | -0.52 [-3.34, 2.27] | -3.87 [-7.05, -0.67]* | -6.87 [-9.99, -3.54]* | 3.34 [0.24, 6.45]* | -6.34 [-9.45, -2.99]* | -2.97 [-6.44, 0.72] |
| R 993 | R 500 | R Secondary motor area | 5.80 ± 1.65 | 11 | 23.38 ± 15.05 | 15 | 36.65 ± 7.58 | 10 | 53.69 ± 11.17 | 9 | 4.74 [1.38, 8.08]* | 3.40 [-0.16, 7.02] | -1.73 [-6.62, 3.12] | -3.11 [-6.03, -0.14]* | -5.47 [-8.77, -2.18]* | -8.84 [-12.26, -5.55]* | 2.34 [-0.81, 5.43] | -5.73 [-8.85, -2.36]* | -3.38 [-6.96, 0.03] |
| L 993 | L 500 | L Secondary motor area | 4.74 ± 1.34 | 13 | 15.44 ± 7.16 | 13 | 31.73 ± 5.20 | 10 | 44.49 ± 10.09 | 9 | 5.41 [1.98, 8.82]* | 3.13 [-0.41, 6.67] | -1.98 [-6.79, 2.9] | -2.62 [-5.63, 0.32] | -5.87 [-9.12, -2.72]* | -9.00 [-12.28, -5.76]* | 3.27 [0.12, 6.51]* | -6.39 [-9.5, -2.89]* | -3.11 [-6.54, 0.39] |
| R 453 | R 315 | R Somatosensory areas | 7.98 ± 2.04 | 15 | 10.58 ± 3.13 | 15 | 16.86 ± 2.61 | 10 | 33.54 ± 5.92 | 10 | 1.78 [-1.46, 4.97] | 1.94 [-1.57, 5.46] | -2.90 [-7.58, 1.72] | | | | | | |
| L 453 | L 315 | L Somatosensory areas | 3.77 ± 1.03 | 15 | 4.88 ± 1.41 | 15 | 10.63 ± 1.88 | 10 | 17.38 ± 2.93 | 9 | 2.81 [-0.54, 6.15] | 2.24 [-1.34, 5.89] | -3.24 [-8, 1.53] | | | | | | |
| R 322 | R 453 | R Primary somatosensory area | 13.09 ± 3.48 | 14 | 14.55 ± 4.78 | 15 | 22.31 ± 4.10 | 9 | 39.50 ± 7.03 | 9 | 1.40 [-1.94, 4.7] | 1.82 [-1.87, 5.51] | -3.02 [-7.87, 1.85] | | | | | | |
| L 322 | L 453 | L Primary somatosensory area | 4.49 ± 1.24 | 15 | 5.16 ± 1.33 | 15 | 13.17 ± 2.48 | 10 | 21.91 ± 3.69 | 9 | 3.24 [-0.08, 6.63] | 2.40 [-1.24, 5.98] | -3.56 [-8.25, 1.25] | | | | | | |
| R 353 | R 322 | R Primary somatosensory area, nose | 5.00 ± 1.42 | 10 | 2.96 ± 0.91 | 9 | 34.14 ± 17.04 | 8 | 130.19 ± 82.08 | 8 | 24.08 [20.26, 27.85]* | 19.52 [15.64, 23.4]* | -20.80 [-26.16, -15.4]* | -0.18 [-3.65, 3.31] | -5.74 [-9.36, -2.12]* | -25.24 [-28.83, -21.61]* | 5.54 [1.92, 9.3]* | -25.05 [-28.78, -21.45]* | -19.50 [-23.24, -15.64]* |
| L 353 | L 322 | L Primary somatosensory area, nose | 2.53 ± 1.22 | 9 | 10.19 ± 6.34 | 7 | 14.55 ± 3.92 | 8 | 40.70 ± 22.49 | 7 | 11.25 [7.17, 15.3]* | 10.38 [6.47, 14.28]* | -8.62 [-14.27, -2.97]* | -3.22 [-7.05, 0.36] | -5.05 [-8.69, -1.36]* | -15.43 [-19.21, -11.75]* | 1.88 [-2.13, 5.47] | -12.22 [-16.11, -8.09]* | -10.35 [-14.1, -6.49]* |
| R 329 | R 322 | R Primary somatosensory area, barrel field | 21.31 ± 6.52 | 13 | 25.20 ± 7.66 | 15 | 36.52 ± 7.51 | 10 | 60.61 ± 13.56 | 9 | 1.16 [-2.11, 4.43] | 1.59 [-1.97, 5.21] | -2.71 [-7.41, 2.12] | | | | | | |
| L 329 | L 322 | L Primary somatosensory area, barrel field | 6.90 ± 2.48 | 14 | 6.63 ± 2.17 | 15 | 19.37 ± 3.84 | 10 | 34.91 ± 5.68 | 9 | 3.61 [0.31, 6.98]* | 2.70 [-0.93, 6.36] | -4.06 [-8.94, 0.75] | -0.10 [-2.94, 2.88] | -1.98 [-5.26, 1.19] | -4.68 [-8.02, -1.45]* | 1.89 [-1.25, 5.07] | -4.57 [-7.64, -1.1]* | -2.66 [-6.25, 0.81] |
| R 337 | R 322 | R Primary somatosensory area, lower limb | 20.10 ± 9.07 | 13 | 14.48 ± 5.11 | 11 | 49.70 ± 10.78 | 10 | 96.53 ± 21.81 | 9 | 3.35 [-0.17, 6.85] | 2.80 [-0.84, 6.39] | -4.17 [-9.06, 0.78] | | | | | | |
| L 337 | L 322 | L Primary somatosensory area, lower limb | 6.74 ± 3.03 | 13 | 3.01 ± 1.18 | 10 | 30.68 ± 5.73 | 10 | 58.81 ± 15.27 | 9 | 7.59 [3.94, 11.16]* | 4.62 [0.99, 8.22]* | -6.30 [-11.32, -1.32]* | 0.21 [-2.92, 3.46] | -3.72 [-7.21, -0.69]* | -8.33 [-11.55, -4.89]* | 3.98 [0.53, 7.37]* | -8.58 [-11.88, -4.94]* | -4.60 [-8.1, -1.09]* |
| R 345 | R 322 | R Primary somatosensory area, mouth | 7.15 ± 1.86 | 13 | 11.45 ± 4.21 | 13 | 17.03 ± 4.05 | 9 | 20.59 ± 5.08 | 9 | 0.74 [-2.69, 4.15] | 1.00 [-2.64, 4.72] | -1.62 [-6.5, 3.49] | | | | | | |
| L 345 | L 322 | L Primary somatosensory area, mouth | 9.63 ± 3.91 | 13 | 6.09 ± 1.03 | 12 | 15.55 ± 4.92 | 10 | 18.33 ± 4.13 | 8 | 0.52 [-3.05, 4.22] | 0.81 [-3.01, 4.56] | -2.12 [-7.21, 2.97] | | | | | | |
| R 369 | R 322 | R Primary somatosensory area, upper limb | 13.70 ± 4.35 | 14 | 11.60 ± 4.83 | 13 | 24.59 ± 4.76 | 10 | 67.75 ± 19.75 | 9 | 3.52 [0.12, 6.93]* | 3.59 [-0.03, 7.17] | -5.00 [-9.85, -0.27]* | -0.06 [-2.99, 2.85] | -0.97 [-4.07, 2.25] | -4.52 [-7.7, -1.1]* | 0.94 [-2.28, 4.17] | -4.46 [-7.78, -1.19]* | -3.57 [-6.91, 0.11] |
| L 369 | L 322 | L Primary somatosensory area, upper limb | 9.66 ± 3.36 | 14 | 6.89 ± 2.87 | 13 | 20.72 ± 4.37 | 10 | 34.75 ± 8.80 | 9 | 2.32 [-1.11, 5.72] | 1.92 [-1.68, 5.45] | -3.48 [-8.31, 1.43] | | | | | | |
| R 361 | R 322 | R Primary somatosensory area, trunk | 51.38 ± 26.74 | 14 | 37.77 ± 26.23 | 12 | 95.76 ± 30.65 | 10 | 145.56 ± 41.58 | 9 | 1.62 [-1.9, 5.07] | 1.42 [-2.18, 5.04] | -3.00 [-7.83, 1.93] | | | | | | |
| L 361 | L 322 | L Primary somatosensory area, trunk | 6.44 ± 1.56 | 13 | 6.60 ± 2.40 | 13 | 35.13 ± 5.87 | 10 | 46.55 ± 11.90 | 9 | 5.66 [2.26, 9.07]* | 2.22 [-1.42, 5.76] | -3.45 [-8.23, 1.51] | -0.23 [-3.33, 2.74] | -4.64 [-7.98, -1.42]* | -6.85 [-10.18, -3.52]* | 4.41 [1.07, 7.51]* | -6.62 [-9.92, -3.24]* | -2.23 [-5.59, 1.43] |
| R 182305689 | R 322 | R Primary somatosensory area, unassigned | 72.29 ± 39.72 | 14 | 8.27 ± 3.64 | 11 | 25.05 ± 5.66 | 10 | 83.67 ± 24.57 | 9 | 0.35 [-3.18, 3.9] | 1.28 [-2.35, 4.93] | -3.28 [-8.28, 1.59] | | | | | | |
| L 182305689 | L 322 | L Primary somatosensory area, unassigned | 7.64 ± 4.28 | 14 | 3.45 ± 1.32 | 11 | 14.41 ± 2.64 | 10 | 41.83 ± 12.39 | 9 | 4.58 [1.09, 8.13]* | 4.04 [0.43, 7.67]* | -5.95 [-10.94, -0.95]* | 0.46 [-2.61, 3.52] | -1.07 [-4.38, 2] | -5.07 [-8.21, -1.79]* | 1.53 [-1.87, 4.75] | -5.53 [-8.96, -2.2]* | -4.01 [-7.46, -0.49]* |
| R 378 | R 453 | R Supplemental somatosensory area | 8.32 ± 2.89 | 11 | 18.36 ± 8.34 | 15 | 29.30 ± 4.90 | 9 | 60.38 ± 14.11 | 10 | 3.96 [0.66, 7.15]* | 3.58 [-0.02, 7.21] | -3.84 [-8.76, 1.06] | -1.20 [-4.28, 1.81] | -2.59 [-6.01, 0.98] | -6.13 [-9.54, -2.85]* | 1.39 [-1.75, 4.69] | -4.94 [-8.04, -1.8]* | -3.56 [-7.15, -0.07]* |
| L 378 | L 453 | L Supplemental somatosensory area | 8.59 ± 2.35 | 14 | 12.11 ± 4.05 | 15 | 20.82 ± 3.37 | 10 | 50.86 ± 15.16 | 9 | 3.14 [-0.13, 6.46] | 3.07 [-0.54, 6.7] | -3.98 [-8.85, 0.81] | | | | | | |
| R 1057 | R 315 | R Gustatory areas | 13.02 ± 3.89 | 15 | 11.76 ± 3.43 | 13 | 18.97 ± 5.00 | 10 | 25.50 ± 6.51 | 10 | -0.09 [-3.47, 3.27] | 0.34 [-3.16, 3.88] | -1.69 [-6.42, 2.98] | | | | | | |
| L 1057 | L 315 | L Gustatory areas | 6.25 ± 2.92 | 15 | 8.05 ± 3.73 | 13 | 14.53 ± 4.51 | 10 | 35.97 ± 14.13 | 10 | 2.99 [-0.32, 6.31] | 3.28 [-0.29, 6.79] | -3.91 [-8.63, 0.88] | | | | | | |
| R 677 | R 315 | R Visceral area | 15.05 ± 5.31 | 13 | 29.68 ± 17.13 | 13 | 31.77 ± 9.44 | 9 | 44.54 ± 11.79 | 10 | -0.20 [-3.53, 3.13] | 0.70 [-2.9, 4.32] | -0.94 [-5.88, 3.87] | | | | | | |
| L 677 | L 315 | L Visceral area | 9.44 ± 2.88 | 14 | 10.52 ± 5.44 | 13 | 25.27 ± 3.71 | 10 | 35.68 ± 6.46 | 9 | 1.21 [-2.18, 4.63] | 0.67 [-3.03, 4.34] | -1.81 [-6.66, 3.1] | | | | | | |
| R 247 | R 315 | R Auditory areas | 17.78 ± 5.75 | 14 | 8.29 ± 2.05 | 15 | 20.08 ± 2.64 | 10 | 39.43 ± 8.95 | 9 | 0.63 [-2.71, 4.01] | 0.92 [-2.67, 4.54] | -2.68 [-7.51, 2.2] | | | | | | |
| L 247 | L 315 | L Auditory areas | 10.83 ± 2.92 | 14 | 6.04 ± 1.94 | 14 | 11.06 ± 2.13 | 10 | 54.02 ± 32.01 | 10 | 3.09 [-0.2, 6.29] | 3.80 [0.26, 7.28]* | -5.24 [-9.94, -0.55]* | -0.01 [-2.87, 2.89] | -0.30 [-3.58, 2.77] | -4.06 [-7.34, -0.98]* | 0.28 [-2.73, 3.58] | -4.06 [-7.19, -0.85]* | -3.77 [-7.3, -0.44]* |
| R 1011 | R 247 | R Dorsal auditory area | 27.64 ± 11.66 | 12 | 10.47 ± 2.59 | 15 | 36.61 ± 4.73 | 10 | 56.42 ± 11.20 | 8 | 0.55 [-2.92, 3.93] | 0.55 [-3.12, 4.22] | -2.37 [-7.31, 2.52] | | | | | | |
| L 1011 | L 247 | L Dorsal auditory area | 17.69 ± 8.89 | 12 | 8.52 ± 3.05 | 14 | 21.26 ± 3.62 | 10 | 83.71 ± 49.15 | 10 | 2.90 [-0.42, 6.2] | 3.38 [-0.13, 6.91] | -4.93 [-9.84, -0.25]* | 0.10 [-2.89, 3.11] | -0.39 [-3.73, 2.85] | -3.78 [-7, -0.46]* | 0.53 [-2.71, 3.69] | -3.87 [-6.98, -0.7]* | -3.35 [-6.86, 0.01] |
| R 1002 | R 247 | R Primary auditory area | 32.54 ± 13.40 | 13 | 18.16 ± 6.26 | 11 | 29.12 ± 6.48 | 10 | 63.99 ± 24.47 | 7 | 0.55 [-3.2, 4.22] | 1.25 [-2.54, 5.04] | -2.77 [-7.79, 2.22] | | | | | | |
| L 1002 | L 247 | L Primary auditory area | 16.81 ± 4.93 | 13 | 8.79 ± 3.94 | 13 | 16.43 ± 4.11 | 10 | 74.83 ± 52.16 | 9 | 2.93 [-0.46, 6.32] | 3.66 [0.07, 7.21]* | -5.15 [-10.02, -0.33]* | 0.04 [-3.01, 3.12] | -0.25 [-3.43, 3] | -3.88 [-7.14, -0.62]* | 0.26 [-2.97, 3.48] | -3.91 [-7.17, -0.6]* | -3.64 [-7.28, -0.31]* |
| L 1027 | L 247 | L Posterior auditory area | 20.38 ± 6.63 | 7 | 3.38 ± 2.15 | 7 | 19.22 ± 10.50 | 8 | 33.38 ± 14.04 | 6 | | | | | | | | | |
| R 1018 | R 247 | R Ventral auditory area | 33.66 ± 9.68 | 14 | 13.65 ± 3.35 | 15 | 27.87 ± 4.88 | 10 | 51.41 ± 15.55 | 8 | -0.01 [-3.46, 3.39] | 0.50 [-3.17, 4.23] | -2.33 [-7.25, 2.56] | | | | | | |
| L 1018 | L 247 | L Ventral auditory area | 15.58 ± 5.15 | 14 | 10.66 ± 3.62 | 14 | 14.93 ± 3.20 | 10 | 89.32 ± 55.95 | 10 | 3.70 [0.45, 7.03]* | 4.62 [1.12, 8.07]* | -5.92 [-10.63, -1.15]* | -0.16 [-3.14, 2.73] | -0.23 [-3.38, 2.93] | -4.82 [-7.93, -1.58]* | 0.08 [-3.15, 3.16] | -4.66 [-7.87, -1.52]* | -4.60 [-8.12, -1.27]* |
| R 669 | R 315 | R Visual areas | 54.72 ± 25.32 | 14 | 48.36 ± 17.02 | 15 | 57.83 ± 19.53 | 10 | 110.80 ± 29.95 | 9 | 0.04 [-3.3, 3.43] | 0.82 [-2.79, 4.5] | -2.16 [-7.04, 2.66] | | | | | | |
| L 669 | L 315 | L Visual areas | 8.42 ± 2.63 | 14 | 3.54 ± 1.08 | 14 | 13.11 ± 3.21 | 10 | 37.91 ± 18.18 | 8 | 3.10 [-0.41, 6.53] | 3.16 [-0.48, 6.79] | -4.75 [-9.66, 0.18] | | | | | | |
| L 402 | L 669 | L Anterolateral visual area | 15.34 ± 6.99 | 8 | 4.77 ± 1.90 | 7 | 16.94 ± 6.70 | 7 | 117.60 ± 92.92 | 7 | 6.16 [2.12, 10.25]* | 6.24 [2.22, 10.31]* | -7.82 [-13.53, -2.22]* | 0.14 [-3.66, 3.99] | -0.79 [-4.54, 3.07] | -6.99 [-10.93, -3.09]* | 0.93 [-3.25, 4.73] | -7.08 [-11.05, -3.11]* | -6.19 [-10.31, -2.32]* |
| R 394 | R 669 | R Anteromedial visual area | 44.33 ± 13.76 | 12 | 95.21 ± 36.19 | 14 | 103.91 ± 29.82 | 10 | 215.19 ± 64.92 | 9 | 1.65 [-1.68, 5.05] | 2.36 [-1.25, 5.9] | -2.49 [-7.45, 2.42] | | | | | | |
| L 394 | L 669 | L Anteromedial visual area | 11.54 ± 5.55 | 13 | 8.21 ± 4.02 | 13 | 28.16 ± 5.57 | 10 | 53.97 ± 14.75 | 8 | 3.01 [-0.45, 6.47] | 2.44 [-1.22, 6.07] | -3.78 [-8.69, 1.16] | | | | | | |
| R 385 | R 669 | R Primary visual area | 78.58 ± 24.18 | 11 | 164.43 ± 89.74 | 9 | 82.68 ± 36.28 | 9 | 230.76 ± 68.78 | 6 | -0.70 [-4.75, 3.42] | 1.55 [-2.58, 5.64] | -1.46 [-6.9, 3.99] | | | | | | |
| L 385 | L 669 | L Primary visual area | 14.06 ± 3.55 | 11 | 3.49 ± 1.28 | 10 | 13.67 ± 5.21 | 8 | 52.83 ± 27.42 | 8 | 2.52 [-1.11, 6.2] | 3.23 [-0.63, 7.1] | -4.90 [-10.15, 0.24] | | | | | | |
| R 533 | R 669 | R posteromedial visual area | 237.86 ± 167.43 | 10 | 153.57 ± 107.41 | 7 | 81.10 ± 31.50 | 9 | 152.00 ± 30.36 | 6 | -1.46 [-5.66, 2.76] | -0.03 [-3.98, 4] | -1.35 [-6.95, 4.12] | | | | | | |
| L 533 | L 669 | L posteromedial visual area | 26.40 ± 7.87 | 10 | 13.53 ± 7.23 | 9 | 18.53 ± 5.23 | 9 | 43.13 ± 10.06 | 7 | -0.22 [-4.18, 3.74] | 0.82 [-3.1, 4.74] | -2.24 [-7.59, 3.06] | | | | | | |
| R 31 | R 315 | R Anterior cingulate area | 8.29 ± 2.44 | 15 | 13.78 ± 5.10 | 15 | 32.48 ± 10.49 | 10 | 38.50 ± 12.30 | 10 | 1.87 [-1.45, 5.13] | 0.57 [-3, 4.11] | -1.20 [-5.96, 3.41] | | | | | | |
| L 31 | L 315 | L Anterior cingulate area | 3.94 ± 0.80 | 15 | 11.65 ± 5.33 | 14 | 21.63 ± 5.61 | 10 | 34.37 ± 11.04 | 10 | 4.70 [1.39, 8]* | 3.07 [-0.44, 6.6] | -2.46 [-7.27, 2.12] | -2.06 [-5, 0.66] | -4.71 [-7.85, -1.46]* | -7.75 [-10.91, -4.63]* | 2.63 [-0.61, 5.79] | -5.68 [-8.92, -2.57]* | -3.04 [-6.51, 0.36] |
| R 39 | R 31 | R Anterior cingulate area, dorsal part | 6.93 ± 1.86 | 13 | 17.59 ± 10.04 | 14 | 42.33 ± 9.84 | 10 | 54.62 ± 16.08 | 10 | 4.08 [0.73, 7.4]* | 1.59 [-2.04, 5.14] | -1.15 [-6.05, 3.68] | -1.88 [-4.96, 1.06] | -5.39 [-8.6, -2.03]* | -6.95 [-10.12, -3.6]* | 3.50 [0.35, 6.64]* | -5.07 [-8.41, -1.99]* | -1.57 [-4.99, 1.92] |
| L 39 | L 31 | L Anterior cingulate area, dorsal part | 4.57 ± 1.05 | 15 | 20.00 ± 10.28 | 14 | 33.95 ± 8.46 | 10 | 41.62 ± 8.83 | 10 | 3.65 [0.33, 6.94]* | 1.50 [-2.08, 5.07] | 0.54 [-4.18, 5.35] | -3.51 [-6.38, -0.7]* | -6.63 [-9.96, -3.58]* | -8.11 [-11.33, -4.98]* | 3.15 [0.03, 6.34]* | -4.63 [-7.71, -1.32]* | -1.49 [-4.83, 1.98] |
| R 48 | R 31 | R Anterior cingulate area, ventral part | 21.21 ± 6.18 | 15 | 20.35 ± 6.30 | 14 | 57.09 ± 22.12 | 10 | 66.57 ± 21.94 | 10 | 0.94 [-2.37, 4.23] | 0.28 [-3.28, 3.72] | -1.51 [-6.16, 3.13] | | | | | | |
| L 48 | L 31 | L Anterior cingulate area, ventral part | 7.54 ± 2.32 | 15 | 14.40 ± 5.09 | 13 | 32.99 ± 9.06 | 10 | 66.33 ± 26.42 | 10 | 5.69 [2.41, 9.02]* | 4.25 [0.8, 7.72]* | -4.57 [-9.27, 0.2] | -1.14 [-4.07, 1.75] | -3.57 [-6.73, -0.42]* | -7.81 [-10.95, -4.72]* | 2.42 [-0.79, 5.62] | -6.66 [-9.81, -3.44]* | -4.22 [-7.48, -0.69]* |
| R 95 | R 315 | R Agranular insular area | 6.38 ± 2.48 | 14 | 12.03 ± 4.65 | 15 | 17.93 ± 3.66 | 9 | 20.94 ± 3.65 | 10 | 0.29 [-2.98, 3.55] | 0.31 [-3.33, 3.87] | -0.72 [-5.54, 4.11] | | | | | | |
| L 95 | L 315 | L Agranular insular area | 5.38 ± 1.72 | 15 | 5.03 ± 0.89 | 15 | 14.86 ± 3.54 | 10 | 20.88 ± 4.34 | 10 | 1.84 [-1.39, 5.1] | 0.95 [-2.52, 4.4] | -2.31 [-6.92, 2.3] | | | | | | |
| R 104 | R 95 | R Agranular insular area, dorsal part | 6.52 ± 2.24 | 11 | 15.32 ± 5.72 | 14 | 26.78 ± 9.26 | 9 | 28.42 ± 7.20 | 10 | 0.94 [-2.4, 4.21] | 0.11 [-3.46, 3.72] | -0.29 [-5.2, 4.67] | | | | | | |
| L 104 | L 95 | L Agranular insular area, dorsal part | 6.25 ± 1.72 | 14 | 7.36 ± 2.40 | 15 | 24.94 ± 7.85 | 10 | 38.77 ± 12.65 | 9 | 3.91 [0.58, 7.27]* | 2.05 [-1.54, 5.57] | -3.12 [-8.01, 1.64] | -0.39 [-3.31, 2.47] | -3.23 [-6.64, -0.17]* | -5.26 [-8.54, -1.94]* | 2.84 [-0.24, 6.04] | -4.86 [-8.33, -1.88]* | -2.01 [-5.39, 1.58] |
| R 111 | R 95 | R Agranular insular area, posterior part | 11.70 ± 4.74 | 12 | 26.16 ± 17.79 | 12 | 35.56 ± 10.45 | 9 | 33.17 ± 6.35 | 10 | -0.50 [-3.88, 2.93] | -0.35 [-3.93, 3.32] | 0.20 [-4.76, 5.06] | | | | | | |
| L 111 | L 95 | L Agranular insular area, posterior part | 11.55 ± 4.78 | 15 | 8.13 ± 2.55 | 13 | 24.42 ± 5.26 | 10 | 32.43 ± 6.24 | 10 | 0.94 [-2.34, 4.22] | 0.53 [-3, 4.14] | -2.06 [-6.88, 2.71] | | | | | | |
| R 119 | R 95 | R Agranular insular area, ventral part | 11.71 ± 5.05 | 14 | 19.54 ± 10.15 | 14 | 26.73 ± 5.86 | 9 | 31.41 ± 8.69 | 10 | -0.05 [-3.37, 3.24] | 0.24 [-3.32, 3.87] | -0.92 [-5.71, 3.84] | | | | | | |
| L 119 | L 95 | L Agranular insular area, ventral part | 6.59 ± 1.79 | 14 | 5.92 ± 1.40 | 15 | 47.52 ± 24.82 | 10 | 40.86 ± 14.79 | 9 | 4.20 [0.79, 7.53]* | -1.19 [-4.8, 2.49] | -0.14 [-5.05, 4.72] | -0.12 [-2.9, 2.86] | -6.51 [-9.49, -3.22]* | -5.30 [-8.51, -1.99]* | 6.37 [3.26, 9.51]* | -5.17 [-8.39, -1.85]* | 1.21 [-2.27, 4.75] |
| R 254 | R 315 | R Retrosplenial area | 22.27 ± 9.52 | 14 | 24.48 ± 11.17 | 15 | 98.79 ± 32.94 | 10 | 88.68 ± 13.82 | 9 | 1.78 [-1.57, 5.07] | -0.63 [-4.22, 2.95] | -0.55 [-5.37, 4.25] | | | | | | |
| L 254 | L 315 | L Retrosplenial area | 10.49 ± 4.30 | 14 | 11.16 ± 4.02 | 15 | 30.99 ± 6.29 | 10 | 42.24 ± 6.72 | 10 | 1.85 [-1.42, 5.18] | 0.90 [-2.6, 4.44] | -2.13 [-6.84, 2.62] | | | | | | |
| R 894 | R 254 | R Retrosplenial area, lateral agranular part | 28.55 ± 7.17 | 14 | 28.25 ± 12.27 | 15 | 78.36 ± 13.86 | 10 | 111.40 ± 19.00 | 9 | 1.82 [-1.52, 5.12] | 1.01 [-2.63, 4.58] | -2.33 [-7.06, 2.37] | | | | | | |
| L 894 | L 254 | L Retrosplenial area, lateral agranular part | 11.77 ± 3.81 | 13 | 6.90 ± 2.30 | 15 | 41.80 ± 8.89 | 10 | 56.81 ± 6.75 | 10 | 3.11 [-0.17, 6.38] | 1.11 [-2.4, 4.61] | -2.81 [-7.57, 1.91] | | | | | | |
| R 879 | R 254 | R Retrosplenial area, dorsal part | 15.21 ± 3.59 | 15 | 12.53 ± 3.13 | 15 | 85.87 ± 21.90 | 9 | 110.52 ± 17.06 | 9 | 5.14 [1.83, 8.46]* | 1.43 [-2.17, 5.16] | -2.89 [-7.71, 1.9] | 0.03 [-2.76, 2.88] | -4.71 [-7.93, -1.36]* | -6.11 [-9.41, -2.91]* | 4.72 [1.43, 7.89]* | -6.10 [-9.3, -2.94]* | -1.39 [-5.06, 2.1] |
| L 879 | L 254 | L Retrosplenial area, dorsal part | 11.05 ± 2.70 | 14 | 9.64 ± 3.10 | 15 | 33.57 ± 5.62 | 10 | 61.98 ± 12.01 | 9 | 3.44 [0.1, 6.79]* | 2.23 [-1.42, 5.85] | -3.65 [-8.39, 1.1] | -0.03 [-2.86, 2.85] | -2.25 [-5.46, 1] | -4.45 [-7.79, -1.24]* | 2.21 [-0.89, 5.36] | -4.40 [-7.64, -1.18]* | -2.19[-5.71, -0.21]* |
| R 886 | R 254 | R Retrosplenial area, ventral part | 47.86 ± 24.36 | 15 | 60.11 ± 29.98 | 15 | 236.46 ± 88.67 | 10 | 177.67 ± 34.74 | 10 | 1.34 [-1.97, 4.67] | -1.40 [-4.9, 2.2] | 0.33 [-4.49, 5.04] | | | | | | |
| L 886 | L 254 | L Retrosplenial area, ventral part | 22.31 ± 11.30 | 14 | 27.87 ± 11.52 | 15 | 61.71 ± 14.92 | 10 | 76.84 ± 14.47 | 10 | 1.11 [-2.12, 4.33] | 0.53 [-2.99, 4.02] | -1.60 [-6.35, 3.14] | | | | | | |
| R 22 | R 315 | R Posterior parietal association areas | 39.18 ± 12.52 | 15 | 50.76 ± 23.16 | 15 | 62.21 ± 23.09 | 10 | 158.60 ± 46.08 | 9 | 1.64 [-1.77, 5.05] | 2.30 [-1.27, 5.99] | -3.29 [-8.11, 1.49] | | | | | | |
| L 22 | L 315 | L Posterior parietal association areas | 5.44 ± 1.80 | 14 | 3.93 ± 1.48 | 15 | 14.75 ± 2.94 | 10 | 45.64 ± 27.99 | 10 | 6.57 [3.4, 9.86]* | 5.52 [2.01, 9.05]* | -7.08 [-11.8, -2.48]* | 0.11 [-2.73, 2.96] | -1.93 [-5.2, 1.26] | -7.43 [-10.48, -4.24]* | 2.04 [-1.01, 5.3] | -7.52 [-10.76, -4.6]* | -5.47 [-8.97, -2.13]* |
| R 312782546 | R 22 | R Anterior area | 46.77 ± 13.33 | 15 | 92.78 ± 40.73 | 15 | 110.63 ± 45.89 | 10 | 266.65 ± 97.38 | 9 | 2.61 [-0.71, 5.95] | 3.18 [-0.44, 6.76] | -3.49 [-8.3, 1.26] | | | | | | |
| L 312782546 | L 22 | L Anterior area | 6.81 ± 2.56 | 15 | 5.89 ± 1.98 | 15 | 23.24 ± 4.54 | 10 | 28.46 ± 6.20 | 9 | 2.82 [-0.5, 6.26] | 1.22 [-2.44, 4.81] | -2.67 [-7.48, 2.11] | | | | | | |
| R 417 | R 22 | R Rostrolateral visual area | 50.15 ± 21.36 | 14 | 25.85 ± 10.34 | 15 | 48.28 ± 12.49 | 10 | 196.09 ± 73.10 | 9 | 2.29 [-1, 5.59] | 2.78 [-0.82, 6.43] | -4.50 [-9.21, 0.23] | | | | | | |
| L 417 | L 22 | L Rostrolateral visual area | 7.56 ± 2.93 | 14 | 6.39 ± 2.96 | 14 | 20.30 ± 5.14 | 10 | 81.81 ± 52.96 | 8 | 9.00 [5.53, 12.47]* | 8.35 [4.58, 12.02]* | -9.51 [-14.43, -4.65]* | -0.30 [-3.21, 2.58] | -1.95 [-5.14, 1.21] | -10.24 [-13.59, -6.83]* | 1.64 [-1.59, 4.73] | -9.95 [-13.24, -6.52]* | -8.32 [-12.01, -4.79]* |
| R 541 | R 315 | R Temporal association areas | 23.43 ± 6.28 | 13 | 19.91 ± 7.06 | 12 | 26.26 ± 4.11 | 10 | 34.78 ± 4.41 | 7 | 0.13 [-3.59, 3.84] | 0.98 [-2.91, 4.87] | -2.21 [-7.3, 2.93] | | | | | | |
| L 541 | L 315 | L Temporal association areas | 12.21 ± 5.50 | 13 | 7.10 ± 2.46 | 15 | 19.34 ± 2.76 | 10 | 53.15 ± 21.86 | 10 | 2.67 [-0.6, 5.95] | 2.61 [-0.82, 6.16] | -4.25 [-9.02, 0.41] | | | | | | |
| R 922 | R 315 | R Perirhinal area | 49.67 ± 29.68 | 14 | 28.17 ± 10.75 | 13 | 24.22 ± 8.63 | 10 | 81.15 ± 23.26 | 8 | 0.73 [-2.8, 4.27] | 1.67 [-2.04, 5.36] | -3.42 [-8.31, 1.41] | | | | | | |
| L 922 | L 315 | L Perirhinal area | 11.83 ± 4.85 | 14 | 4.56 ± 1.70 | 15 | 22.29 ± 6.38 | 10 | 19.37 ± 4.45 | 10 | 0.17 [-3.08, 3.46] | -0.41 [-3.94, 3.18] | -1.41 [-6.13, 3.36] | | | | | | |
| R 895 | R 315 | R Ectorhinal area | 22.00 ± 8.20 | 14 | 19.99 ± 6.73 | 13 | 28.60 ± 4.77 | 10 | 51.25 ± 8.81 | 8 | 1.10 [-2.47, 4.66] | 1.54 [-2.22, 5.3] | -2.97 [-7.92, 1.98] | | | | | | |
| L 895 | L 315 | L Ectorhinal area | 14.63 ± 8.02 | 14 | 3.28 ± 1.03 | 15 | 18.47 ± 5.31 | 10 | 30.13 ± 6.17 | 10 | 0.72 [-2.52, 3.92] | 0.63 [-2.86, 4.16] | -2.65 [-7.32, 2.01] | | | | | | |
| R 698 | R 695 | R Olfactory areas | 18.34 ± 7.02 | 14 | 10.86 ± 1.78 | 14 | 23.93 ± 5.68 | 12 | 35.38 ± 5.97 | 10 | -0.39 [-2.97, 2.16] | -0.07 [-2.49, 2.39] | 0.13 [-3.65, 3.89] | | | | | | |
| L 698 | L 695 | L Olfactory areas | 13.76 ± 4.55 | 14 | 6.69 ± 0.99 | 14 | 15.84 ± 2.56 | 12 | 31.83 ± 5.50 | 10 | -0.56 [-3.1, 2.03] | -0.19 [-2.6, 2.26] | 0.8 [-3.02, 4.48] | | | | | | |
| L 159 | L 698 | L Anterior olfactory nucleus | 18.65 ± 8.79 | 8 | 8.35 ± 2.50 | 7 | 89.98 ± 22.41 | 6 | 138.62 ± 27.32 | 6 | 2.63 [-0.72, 5.96] | -0.46 [-3.59, 2.8] | 3.57 [-1.27, 8.4] | | | | | | |
| R 589 | R 698 | R Taenia tecta | 9.34 ± 2.50 | 12 | 9.79 ± 2.38 | 9 | 62.86 ± 22.79 | 7 | 68.89 ± 34.04 | 8 | 4.16 [1.16, 7.14]* | -0.13 [-2.91, 2.62] | 1.04 [-3.3, 5.36] | 0.31 [-2.37, 2.98] | -4.95 [-7.94, -2.13]* | -6.75 [-9.47, -3.73]* | 5.27 [2.33, 8.34]* | -7.06 [-10.06, -4.05]* | -1.78 [-4.86, 1.53] |
| L 589 | L 698 | L Taenia tecta | 5.38 ± 1.42 | 12 | 5.09 ± 1.71 | 9 | 27.37 ± 7.68 | 8 | 64.79 ± 30.95 | 8 | 2.65 [-0.19, 5.48] | -0.25 [-3.03, 2.54] | 7.3 [3.06, 11.48]* | 0.44 [-2.29, 3.1] | -3.43 [-6.18, -0.64]* | -11.37 [-14.18, -8.49]* | 3.87 [0.96, 6.93]* | -11.81 [-14.78, -8.71]* | -7.9 [-11.02, -4.85]* |
| R 597 | R 589 | R Taenia tecta, dorsal part | 15.21 ± 4.11 | 12 | 25.67 ± 6.64 | 8 | 140.24 ± 59.50 | 7 | 137.62 ± 68.10 | 8 | 6.62 [3.64, 9.59]* | 0.55 [-2.25, 3.38] | -0.47 [-4.85, 3.87] | -0.35 [-3.08, 2.42] | -7.44 [-10.26, -4.49]* | -8.38 [-11.15, -5.39]* | 7.08 [4.05, 10.23]* | -8.02 [-11.01, -4.92]* | -0.96 [-4.28, 2.11] |
| L 597 | L 589 | L Taenia tecta, dorsal part | 10.92 ± 2.91 | 12 | 11.59 ± 3.97 | 8 | 59.20 ± 16.77 | 8 | 129.11 ± 62.04 | 8 | 2.98 [0.04, 5.93]* | -0.08 [-2.98, 2.87] | 6.58 [2.18, 10.99]* | 0.27 [-2.5, 3.13] | -3.8 [-6.59, -0.93]* | -11.15 [-14.15, -8.35]* | 4.05 [1.04, 7.15]* | -11.41 [-14.59, -8.46]* | -7.38 [-10.48, -4.12]* |
| R 814 | R 698 | R Dorsal peduncular area | 20.10 ± 8.89 | 14 | 22.98 ± 6.95 | 8 | 54.90 ± 18.17 | 6 | 90.51 ± 31.69 | 8 | 1.1 [-1.97, 4.17] | 0.15 [-2.61, 2.99] | 1.05 [-3.39, 5.46] | | | | | | |
| L 814 | L 698 | L Dorsal peduncular area | 12.17 ± 2.80 | 13 | 29.03 ± 10.53 | 8 | 34.47 ± 13.42 | 6 | 103.29 ± 50.58 | 8 | 1.22 [-1.79, 4.3] | 1.41 [-1.46, 4.2] | 3.69 [-0.79, 8.21] | | | | | | |
| R 961 | R 698 | R Piriform area | 51.78 ± 23.45 | 10 | 36.69 ± 5.87 | 10 | 65.44 ± 19.47 | 7 | 78.16 ± 14.62 | 10 | -0.07 [-3.14, 3.04] | 0.01 [-2.79, 2.9] | -0.71 [-4.95, 3.57] | | | | | | |
| L 961 | L 698 | L Piriform area | 41.73 ± 11.14 | 12 | 36.06 ± 13.48 | 10 | 41.49 ± 7.31 | 8 | 66.44 ± 14.62 | 10 | -0.33 [-3.19, 2.61] | 0.19 [-2.58, 2.93] | -0.46 [-4.59, 3.66] | | | | | | |
| R 631 | R 698 | R Cortical amygdalar area | 20.26 ± 7.06 | 14 | 7.22 ± 1.45 | 11 | 19.74 ± 7.48 | 9 | 23.57 ± 3.80 | 10 | -0.25 [-3.02, 2.51] | -0.27 [-2.81, 2.31] | -0.54 [-4.49, 3.38] | | | | | | |
| L 631 | L 698 | L Cortical amygdalar area | 9.08 ± 4.90 | 14 | 6.53 ± 1.94 | 12 | 9.45 ± 1.94 | 9 | 22.45 ± 5.46 | 10 | 0 [-2.73, 2.72] | -0.03 [-2.61, 2.46] | 0.2 [-3.7, 4.08] | | | | | | |
| R 639 | R 631 | R Cortical amygdalar area, anterior part | 46.91 ± 19.68 | 14 | 24.76 ± 6.14 | 10 | 74.29 ± 32.14 | 9 | 45.71 ± 8.80 | 10 | 0.35 [-2.38, 3.07] | -0.12 [-2.79, 2.53] | -1.48 [-5.49, 2.45] | | | | | | |
| L 639 | L 631 | L Cortical amygdalar area, anterior part | 29.24 ± 16.45 | 14 | 12.46 ± 2.80 | 11 | 45.04 ± 9.23 | 9 | 86.95 ± 23.30 | 10 | 0.49 [-2.21, 3.29] | -0.29 [-2.86, 2.31] | 0.47 [-3.57, 4.37] | | | | | | |
| R 647 | R 631 | R Cortical amygdalar area, posterior part | 29.81 ± 11.85 | 12 | 10.08 ± 2.54 | 12 | 22.09 ± 9.94 | 9 | 30.12 ± 7.01 | 8 | -0.46 [-3.27, 2.32] | -0.35 [-2.91, 2.25] | -0.55 [-4.58, 3.56] | | | | | | |
| L 647 | L 631 | L Cortical amygdalar area, posterior part | 12.20 ± 6.80 | 13 | 8.41 ± 3.13 | 12 | 5.50 ± 1.52 | 8 | 26.65 ± 7.97 | 9 | -0.53 [-3.34, 2.34] | -0.05 [-2.69, 2.52] | 0.22 [-3.95, 4.3] | | | | | | |
| R 655 | R 647 | R Cortical amygdalar area, posterior part, lateral zone | 64.74 ± 25.38 | 13 | 24.92 ± 7.26 | 12 | 46.82 ± 17.49 | 9 | 69.03 ± 17.24 | 8 | -0.51 [-3.29, 2.26] | -0.35 [-2.93, 2.22] | -0.5 [-4.52, 3.61] | | | | | | |
| L 655 | L 647 | L Cortical amygdalar area, posterior part, lateral zone | 22.05 ± 11.43 | 13 | 15.43 ± 7.18 | 12 | 13.23 ± 3.21 | 8 | 62.89 ± 19.48 | 9 | -0.37 [-3.24, 2.53] | -0.03 [-2.54, 2.56] | 0.71 [-3.4, 4.72] | | | | | | |
| R 663 | R 647 | R Cortical amygdalar area, posterior part, medial zone | 50.29 ± 22.20 | 11 | 14.53 ± 3.42 | 12 | 47.42 ± 26.24 | 8 | 49.42 ± 12.75 | 8 | -0.3 [-3.19, 2.7] | -0.38 [-3.05, 2.3] | -0.7 [-4.87, 3.51] | | | | | | |
| L 663 | L 647 | L Cortical amygdalar area, posterior part, medial zone | 28.58 ± 19.10 | 11 | 17.71 ± 9.68 | 12 | 8.65 ± 3.66 | 8 | 38.29 ± 15.09 | 9 | -0.62 [-3.61, 2.32] | -0.05 [-2.72, 2.54] | -0.48 [-4.62, 3.65] | | | | | | |
| R 788 | R 698 | R Piriform-amygdalar area | 50.03 ± 17.37 | 15 | 30.51 ± 8.44 | 11 | 54.78 ± 13.06 | 10 | 61.20 ± 12.39 | 10 | | | | | | | | | |
| L 788 | L 698 | L Piriform-amygdalar area | 24.69 ± 10.40 | 15 | 16.54 ± 6.42 | 14 | 33.19 ± 7.27 | 10 | 59.63 ± 18.51 | 10 | 0.07 [-2.6, 2.7] | -0.04 [-2.49, 2.39] | 0.1 [-3.7, 3.93] | | | | | | |
| R 1089 | R 695 | R Hippocampal formation | 76.17 ± 21.82 | 15 | 113.86 ± 24.92 | 15 | 307.68 ± 54.66 | 12 | 288.09 ± 66.01 | 10 | 2.14 [-0.34, 4.63] | 0.69 [-1.59, 2.99] | -1.51 [-5.09, 2.05] | | | | | | |
| L 1089 | L 695 | L Hippocampal formation | 13.31 ± 3.98 | 15 | 16.43 ± 3.07 | 15 | 57.00 ± 13.93 | 12 | 60.94 ± 10.31 | 10 | 2.38 [-0.05, 4.83] | 0.43 [-1.85, 2.74] | -0.69 [-4.29, 2.9] | | | | | | |
| R 1080 | R 1089 | R Hippocampal region | 102.19 ± 30.45 | 15 | 169.93 ± 33.74 | 15 | 413.26 ± 72.46 | 12 | 388.38 ± 95.07 | 10 | 2.14 [-0.32, 4.59] | 0.85 [-1.48, 3.16] | -1.65 [-5.18, 1.93] | | | | | | |
| L 1080 | L 1089 | L Hippocampal region | 17.95 ± 5.52 | 15 | 24.50 ± 4.58 | 15 | 76.30 ± 19.02 | 12 | 79.64 ± 14.85 | 10 | 2.35 [-0.08, 4.84] | 0.57 [-1.68, 2.9] | -0.92 [-4.5, 2.58] | | | | | | |
| R 375 | R 1080 | R Ammon's horn | 141.91 ± 46.22 | 15 | 218.54 ± 45.48 | 15 | 489.25 ± 85.73 | 12 | 503.46 ± 115.49 | 10 | 1.54 [-0.89, 3.95] | 0.72 [-1.52, 2.98] | -1.17 [-4.64, 2.34] | | | | | | |
| L 375 | L 1080 | L Ammon's horn | 22.27 ± 6.57 | 15 | 35.81 ± 7.65 | 15 | 97.57 ± 21.92 | 12 | 106.24 ± 19.19 | 10 | 2.49 [0.07, 4.93]* | 0.82 [-1.51, 3.14] | -0.97 [-4.52, 2.57] | -0.65 [-2.89, 1.58] | -3.37 [-5.71, -0.95]* | -4.23 [-6.65, -1.6]* | 2.68 [0.22, 5.04]* | -3.57 [-6.07, -0.99]* | -0.89 [-3.48, 1.83] |
| R 382 | R 375 | R Field CA1 | 254.69 ± 91.37 | 15 | 361.42 ± 72.10 | 15 | 856.87 ± 149.04 | 12 | 811.05 ± 164.63 | 10 | 1.47 [-0.98, 3.9] | 0.62 [-1.65, 2.89] | -1.35 [-4.98, 2.24] | | | | | | |
| L 382 | L 375 | L Field CA1 | 39.81 ± 13.45 | 15 | 74.95 ± 18.36 | 15 | 236.04 ± 46.87 | 12 | 232.86 ± 38.55 | 10 | 4.04 [1.64, 6.47]* | 1.07 [-1.19, 3.36] | -1.71 [-5.3, 1.82] | -0.92 [-3.07, 1.4] | -4.91 [-7.35, -2.62]* | -5.31 [-7.84, -2.71]* | 3.99 [1.73, 6.51]* | -4.39 [-7, -1.87]* | -0.39 [-3.03, 2.33] |
| R 423 | R 375 | R Field CA2 | 415.99 ± 119.59 | 15 | 508.83 ± 108.52 | 15 | 1121.75 ± 168.86 | 12 | 1077.28 ± 296.20 | 10 | 0.81 [-1.66, 3.29] | 0.42 [-1.93, 2.74] | -1.08 [-4.64, 2.52] | | | | | | |
| L 423 | L 375 | L Field CA2 | 42.46 ± 14.34 | 15 | 47.93 ± 13.61 | 15 | 148.84 ± 44.92 | 12 | 189.78 ± 56.18 | 9 | 1.61 [-0.89, 4.04] | 0.34 [-2, 2.66] | -0.06 [-3.66, 3.57] | | | | | | |
| R 463 | R 375 | R Field CA3 | 316.25 ± 107.59 | 14 | 547.25 ± 133.79 | 15 | 1116.28 ± 213.95 | 12 | 1130.61 ± 306.88 | 10 | 1.47 [-0.96, 3.95] | 0.76 [-1.56, 3.07] | -1.28 [-4.87, 2.3] | | | | | | |
| L 463 | L 375 | L Field CA3 | 49.49 ± 15.67 | 15 | 70.68 ± 14.52 | 15 | 158.04 ± 42.60 | 12 | 206.46 ± 41.21 | 9 | 1.3 [-1.14, 3.76] | 0.63 [-1.67, 2.9] | -0.35 [-3.97, 3.26] | | | | | | |
| R 726 | R 1080 | R Dentate gyrus | 189.20 ± 55.75 | 15 | 362.60 ± 77.45 | 15 | 1016.85 ± 191.95 | 12 | 849.39 ± 251.92 | 10 | 3.48 [1.02, 5.93]* | 1.1 [-1.2, 3.39] | -2.56 [-6.11, 0.98] | -0.94 [-3.2, 1.34] | -4.34 [-6.69, -1.97]* | -3.92 [-6.37, -1.28]* | 3.4 [1.07, 5.84]* | -2.98 [-5.64, -0.52]* | 0.43 [-2.2, 3.08] |
| L 726 | L 1080 | L Dentate gyrus | 38.56 ± 13.38 | 15 | 39.91 ± 9.02 | 15 | 172.39 ± 57.24 | 11 | 156.31 ± 32.65 | 10 | 2.62 [0.14, 5.13]* | 0.22 [-2.08, 2.52] | -1.25 [-4.89, 2.41] | -0.06 [-2.26, 2.23] | -3.49 [-5.79, -0.92]* | -3.49 [-6.03, -0.87]* | 3.42 [1, 5.86]* | -3.43 [-5.99, -0.84]* | 0 [-2.64, 2.83] |
| R 10703 | R 726 | R Dentate gyrus, molecular layer | 367.17 ± 114.58 | 15 | 636.27 ± 152.08 | 15 | 1744.62 ± 320.17 | 12 | 1496.69 ± 417.39 | 10 | 2.86 [0.44, 5.37]* | 0.93 [-1.33, 3.18] | -2.17 [-5.71, 1.37] | -0.78 [-3.02, 1.41] | -3.71 [-6.14, -1.36]* | -3.53 [-6.01, -0.89]* | 2.96 [0.49, 5.24]* | -2.75 [-5.17, -0.13]* | 0.22 [-2.46, 2.86] |
| L 10703 | L 726 | L Dentate gyrus, molecular layer | 77.49 ± 25.82 | 15 | 66.72 ± 15.46 | 15 | 313.35 ± 96.93 | 11 | 271.15 ± 53.22 | 10 | 2.19 [-0.27, 4.68] | 0.05 [-2.22, 2.31] | -1.2 [-4.8, 2.33] | | | | | | |
| R 10704 | R 726 | R Dentate gyrus, polymorph layer | 478.71 ± 137.74 | 15 | 1153.91 ± 259.60 | 15 | 2797.94 ± 545.16 | 12 | 2472.65 ± 823.91 | 10 | 3.96 [1.58, 6.45]* | 1.62 [-0.68, 3.92] | -2.86 [-6.42, 0.69] | -1.46 [-3.72, 0.73] | -4.83 [-7.24, -2.46]* | -4.62 [-7.31, -2.11]* | 3.36 [1.12, 5.82]* | -3.16 [-5.8, -0.69]* | 0.2 [-2.59, 2.79] |
| L 10704 | L 726 | L Dentate gyrus, polymorph layer | 98.11 ± 46.41 | 15 | 142.45 ± 34.99 | 15 | 332.78 ± 133.74 | 11 | 465.18 ± 110.73 | 10 | 1.53 [-0.95, 4.03] | 0.63 [-1.67, 2.9] | 0.12 [-3.51, 3.7] | | | | | | |
| R 632 | R 726 | R Dentate gyrus, granule cell layer | 373.29 ± 107.26 | 15 | 793.64 ± 159.74 | 15 | 2510.59 ± 551.19 | 12 | 1950.43 ± 627.18 | 10 | 4.84 [2.4, 7.28]* | 1.32 [-0.94, 3.66] | -3.39 [-6.98, 0.1] | -1.16 [-3.36, 1.15] | -5.71 [-8.17, -3.41]* | -4.66 [-7.18, -2.07]* | 4.54 [2.16, 6.91]* | -3.5 [-5.98, -0.97]* | 1.04 [-1.7, 3.57] |
| L 632 | L 726 | L Dentate gyrus, granule cell layer | 69.34 ± 25.97 | 15 | 91.38 ± 25.63 | 15 | 426.23 ± 160.88 | 11 | 374.11 ± 90.58 | 10 | 4.28 [1.82, 6.78]* | 0.51 [-1.77, 2.8] | -1.86 [-5.43, 1.72] | -0.35 [-2.73, 1.77] | -5.15 [-7.56, -2.73]* | -4.83 [-7.38, -2.23]* | 4.81 [2.3, 7.24]* | -4.47 [-7.07, -1.92]* | 0.3 [-2.41, 2.96] |
| R 982 | R 1080 | R Fasciola cinerea | 484.18 ± 230.22 | 13 | 692.89 ± 216.02 | 15 | 2064.50 ± 503.44 | 11 | 1579.94 ± 651.17 | 8 | 2.44 [-0.08, 4.99] | 0.72 [-1.62, 3.07] | -2.14 [-5.88, 1.6] | | | | | | |
| L 982 | L 1080 | L Fasciola cinerea | 201.26 ± 60.95 | 13 | 235.22 ± 60.88 | 15 | 665.18 ± 283.96 | 11 | 658.64 ± 241.10 | 9 | 1.45 [-1.06, 3.95] | 0.46 [-1.9, 2.82] | -0.94 [-4.57, 2.68] | | | | | | |
| R 19 | R 1080 | R Induseum griseum | 66.55 ± 20.23 | 15 | 78.42 ± 25.34 | 12 | 235.81 ± 80.92 | 12 | 159.08 ± 57.06 | 10 | 1.66 [-0.77, 4.11] | 0.35 [-2.04, 2.71] | -2.07 [-5.69, 1.61] | | | | | | |
| L 19 | L 1080 | L Induseum griseum | 40.36 ± 16.37 | 15 | 88.73 ± 38.72 | 12 | 54.06 ± 14.72 | 12 | 142.48 ± 37.23 | 10 | -0.55 [-3, 1.91] | 1.36 [-0.99, 3.74] | 0.27 [-3.43, 3.83] | | | | | | |
| R 822 | R 1089 | R Retrohippocampal region | 76.27 ± 31.20 | 14 | 71.30 ± 34.44 | 13 | 414.84 ± 184.98 | 12 | 306.12 ± 78.27 | 8 | 3.61 [1.17, 6.02]* | 0.21 [-2.16, 2.62] | -2.21 [-5.87, 1.43] | -0.03 [-2.31, 2.36] | -4.49 [-6.81, -2.11]* | -3.52 [-6.3, -0.87]* | 4.42 [1.97, 6.9]* | -3.47 [-6.25, -0.83]* | 0.95 [-1.73, 3.8] |
| L 822 | L 1089 | L Retrohippocampal region | 10.48 ± 3.52 | 14 | 8.80 ± 2.88 | 15 | 96.91 ± 53.50 | 12 | 68.76 ± 14.20 | 9 | 7.42 [4.97, 9.95]* | 0.1 [-2.23, 2.41] | -3.41 [-7.08, 0.23] | 0.06 [-2.16, 2.46] | -8.27 [-10.81, -5.92]* | -6.03 [-8.6, -3.36]* | 8.34 [5.93, 10.72]* | -6.07 [-8.67, -3.53]* | 2.25 [-0.61, 4.89] |
| R 909 | R 822 | R Entorhinal area | 33.81 ± 14.20 | 13 | 23.78 ± 7.10 | 12 | 39.52 ± 23.14 | 10 | 50.79 ± 14.55 | 8 | | | | | | | | | |
| L 909 | L 822 | L Entorhinal area | 7.12 ± 3.36 | 14 | 5.27 ± 2.28 | 15 | 10.23 ± 3.05 | 10 | 16.21 ± 4.33 | 9 | 0.25 [-2.3, 2.81] | 0 [-2.37, 2.34] | -0.42 [-4.1, 3.32] | | | | | | |
| R 918 | R 909 | R Entorhinal area, lateral part | 52.33 ± 21.96 | 13 | 38.34 ± 12.27 | 12 | 60.87 ± 35.61 | 10 | 78.32 ± 22.42 | 8 | 0 [-2.59, 2.57] | -0.02 [-2.48, 2.45] | -0.85 [-4.67, 2.98] | | | | | | |
| L 918 | L 909 | L Entorhinal area, lateral part | 11.03 ± 5.20 | 14 | 7.66 ± 3.52 | 15 | 15.78 ± 4.71 | 10 | 25.05 ± 6.70 | 9 | 0.24 [-2.29, 2.84] | -0.05 [-2.36, 2.29] | -0.4 [-4.08, 3.28] | | | | | | |
| R 502 | R 822 | R Subiculum | 457.85 ± 173.38 | 12 | 463.21 ± 212.07 | 10 | 2475.96 ± 853.31 | 10 | 2339.07 ± 608.71 | 7 | 3.55 [0.97, 6.15]* | 0.19 [-2.4, 2.72] | -0.72 [-4.71, 3.25] | -0.04 [-2.48, 2.54] | -4.42 [-7, -1.95]* | -4.92 [-7.79, -2.06]* | 4.38 [1.72, 7.02]* | -4.89 [-7.94, -2.11]* | -0.49 [-3.49, 2.46] |
| L 502 | L 822 | L Subiculum | 105.07 ± 47.75 | 12 | 49.79 ± 29.39 | 11 | 546.74 ± 158.41 | 10 | 651.54 ± 87.13 | 7 | 3.33 [0.7, 5.93]* | -0.26 [-2.79, 2.23] | 1.02 [-2.96, 4.91] | 0.42 [-2.13, 2.79] | -4.2 [-6.78, -1.7]* | -5.98 [-8.84, -3.04]* | 4.62 [2.06, 7.27]* | -6.42 [-9.25, -3.33]* | -1.79 [-4.72, 1.22] |
| R 484682470 | R 822 | R Prosubiculum | 1001.71 ± 472.42 | 10 | 806.08 ± 417.98 | 9 | 2890.09 ± 912.18 | 10 | 2521.16 ± 673.63 | 7 | 1.15 [-1.5, 3.9] | 0.06 [-2.63, 2.82] | -0.66 [-4.67, 3.36] | | | | | | |
| L 484682470 | L 822 | L Prosubiculum | 25.39 ± 11.06 | 10 | 86.19 ± 58.03 | 10 | 527.25 ± 149.10 | 10 | 669.50 ± 104.13 | 7 | 18.72 [16, 21.45]* | 2.39 [-0.34, 5.06] | 2.97 [-1.01, 6.95] | -2.24 [-4.81, 0.44] | -19.59 [-22.19, -16.86]* | -25.98 [-29.05, -23.12]* | 17.36 [14.69, 20.04]* | -23.74 [-26.69, -20.9]* | -6.38 [-9.28, -3.37]* |
| R 703 | R 688 | R Cortical subplate | 11.67 ± 5.27 | 15 | 7.44 ± 2.25 | 15 | 33.72 ± 6.72 | 12 | 44.85 ± 5.86 | 10 | 1.78 [0.03, 3.62]* | -0.14 [-1.8, 1.53] | -0.03 [-2.62, 2.57] | 0.21 [-1.42, 1.91] | -2.36 [-4.17, -0.64]* | -3.33 [-5.3, -1.42]* | 2.59 [0.8, 4.36]* | -3.54 [-5.56, -1.69]* | -0.97 [-2.88, 1.11] |
| L 703 | L 688 | L Cortical subplate | 9.13 ± 6.04 | 15 | 5.66 ± 1.49 | 15 | 23.45 ± 3.00 | 12 | 40.15 ± 6.94 | 10 | 1.46 [-0.28, 3.22] | -0.17 [-1.81, 1.49] | 0.87 [-1.76, 3.43] | | | | | | |
| R 583 | R 703 | R Claustrum | 31.89 ± 7.27 | 15 | 17.22 ± 5.61 | 14 | 93.93 ± 30.71 | 12 | 69.55 ± 13.71 | 10 | 1.84 [0.09, 3.61]* | -0.16 [-1.84, 1.48] | -1.73 [-4.29, 0.89] | 0.23 [-1.43, 1.89] | -2.43 [-4.13, -0.65]* | -1.68 [-3.69, 0.14] | 2.66 [0.86, 4.4]* | -1.91 [-3.85, 0] | 0.76 [-1.32, 2.67] |
| L 583 | L 703 | L Claustrum | 7.28 ± 3.41 | 15 | 24.59 ± 11.25 | 14 | 71.94 ± 18.02 | 12 | 74.03 ± 21.49 | 10 | 8.78 [7.02, 10.53]* | 2.68 [1.01, 4.37]* | -3.52 [-6.1, -0.89]* | -2.61 [-4.31, -0.97]* | -9.36 [-11.08, -7.64]* | -9.64 [-11.67, -7.81]* | 6.75 [4.96, 8.52]* | -7.05 [-9.03, -5.12]* | -0.3 [-2.28, 1.69] |
| R 942 | R 703 | R Endopiriform nucleus | 8.90 ± 1.87 | 13 | 8.70 ± 1.80 | 13 | 31.80 ± 8.11 | 12 | 46.38 ± 8.63 | 10 | 1.98 [0.15, 3.8]* | -0.17 [-1.89, 1.56] | 0.67 [-1.95, 3.34] | 0.23 [-1.48, 1.97] | -2.58 [-4.33, -0.76]* | -4.2 [-6.18, -2.29]* | 2.8 [1, 4.69]* | -4.45 [-6.41, -2.5]* | -1.65 [-3.69, 0.33] |
| L 942 | L 703 | L Endopiriform nucleus | 9.26 ± 5.30 | 14 | 12.28 ± 5.18 | 15 | 23.68 ± 4.72 | 12 | 36.81 ± 6.88 | 10 | 1.5 [-0.3, 3.31] | 0.58 [-1.06, 2.3] | -0.3 [-2.89, 2.3] | | | | | | |
| R 952 | R 942 | R Endopiriform nucleus, dorsal part | 18.61 ± 5.22 | 13 | 21.35 ± 6.25 | 14 | 59.39 ± 13.09 | 12 | 79.66 ± 18.40 | 10 | 1.59 [-0.18, 3.43] | -0.16 [-1.87, 1.58] | 0.11 [-2.52, 2.69] | | | | | | |
| L 952 | L 942 | L Endopiriform nucleus, dorsal part | 16.66 ± 8.93 | 14 | 21.67 ± 5.76 | 15 | 38.13 ± 6.83 | 12 | 60.91 ± 13.99 | 10 | 1.23 [-0.59, 3.06] | 0.55 [-1.17, 2.25] | -0.32 [-2.95, 2.35] | | | | | | |
| R 966 | R 942 | R Endopiriform nucleus, ventral part | 16.14 ± 3.39 | 13 | 35.54 ± 15.88 | 14 | 70.02 ± 22.92 | 12 | 114.22 ± 17.44 | 10 | 2.74 [0.91, 4.6]* | 0.89 [-0.86, 2.62] | 0.72 [-1.93, 3.36] | -0.82 [-2.64, 0.82] | -3.32 [-5.15, -1.5]* | -6.09 [-8.06, -4.13]* | 2.51 [0.72, 4.29]* | -5.26 [-7.16, -3.28]* | -2.75 [-4.81, -0.78]* |
| L 966 | L 942 | L Endopiriform nucleus, ventral part | 21.66 ± 13.40 | 14 | 28.32 ± 20.09 | 15 | 63.03 ± 14.33 | 12 | 95.95 ± 16.36 | 10 | 1.85 [0.09, 3.62]* | 0.56 [-1.08, 2.22] | -0.17 [-2.72, 2.34] | -0.49 [-2.09, 1.15] | -2.44 [-4.19, -0.69]* | -3.96 [-5.95, -2.04]* | 1.94 [0.17, 3.71]* | -3.46 [-5.44, -1.58]* | -1.53 [-3.53, 0.46] |
| R 131 | R 703 | R Lateral amygdalar nucleus | 35.39 ± 15.44 | 13 | 20.01 ± 8.78 | 15 | 70.01 ± 14.09 | 12 | 56.87 ± 10.71 | 10 | 0.99 [-0.78, 2.81] | -0.11 [-1.8, 1.59] | -1.38 [-4.01, 1.22] | | | | | | |
| L 131 | L 703 | L Lateral amygdalar nucleus | 17.70 ± 9.49 | 14 | 15.93 ± 5.01 | 15 | 46.27 ± 14.90 | 12 | 54.01 ± 12.70 | 10 | 1.58 [-0.21, 3.38] | 0.19 [-1.51, 1.85] | -0.87 [-3.5, 1.7] | | | | | | |
| R 295 | R 703 | R Basolateral amygdalar nucleus | 17.99 ± 11.30 | 15 | 5.41 ± 1.01 | 13 | 71.25 ± 11.46 | 10 | 98.08 ± 12.73 | 10 | 3.16 [1.36, 4.97]* | -0.34 [-2, 1.33] | 0.4 [-2.19, 3.06] | 0.41 [-1.25, 2.09] | -3.76 [-5.48, -1.89]* | -4.96 [-6.77, -2.89]* | 4.16 [2.25, 6.02]* | -5.37 [-7.31, -3.38]* | -1.2 [-3.25, 0.86] |
| L 295 | L 703 | L Basolateral amygdalar nucleus | 14.02 ± 10.08 | 14 | 5.39 ± 1.44 | 13 | 67.14 ± 5.24 | 10 | 90.75 ± 12.86 | 10 | 4.09 [2.25, 5.94]* | -0.35 [-2.06, 1.35] | 0.54 [-2.14, 3.17] | 0.43 [-1.33, 2.08] | -4.66 [-6.53, -2.91]* | -6.01 [-7.98, -4.01]* | 5.1 [3.14, 6.91]* | -6.42 [-8.4, -4.43]* | -1.34 [-3.39, 0.8] |
| R 303 | R 295 | R Basolateral amygdalar nucleus, anterior part | 27.41 ± 12.59 | 15 | 10.53 ± 3.20 | 12 | 182.11 ± 29.04 | 10 | 232.34 ± 33.97 | 10 | 5.86 [4.06, 7.72]* | -0.23 [-2.01, 1.53] | 0.61 [-2.17, 3.31] | 0.3 [-1.44, 2.06] | -6.45 [-8.27, -4.63]* | -7.98 [-9.95, -6.06]* | 6.75 [4.9, 8.65]* | -8.27 [-10.2, -6.25]* | -1.53 [-3.64, 0.51] |
| L 303 | L 295 | L Basolateral amygdalar nucleus, anterior part | 27.70 ± 17.97 | 14 | 15.37 ± 4.75 | 13 | 161.03 ± 13.94 | 10 | 230.26 ± 35.55 | 10 | 5.11 [3.26, 6.99]* | -0.18 [-1.89, 1.51] | 1.19 [-1.51, 3.82] | 0.25 [-1.37, 2] | -5.7 [-7.49, -3.8]* | -7.85 [-9.79, -5.87]* | 5.94 [4.03, 7.8]* | -8.1 [-10.17, -6.16]* | -2.16 [-4.22, -0.04]* |
| R 311 | R 295 | R Basolateral amygdalar nucleus, posterior part | 33.62 ± 26.51 | 14 | 11.39 ± 3.17 | 12 | 116.90 ± 37.54 | 10 | 184.86 ± 17.21 | 8 | 2.76 [0.9, 4.62]* | -0.25 [-2, 1.47] | 0.85 [-1.87, 3.62] | 0.3 [-1.43, 2.05] | -3.37 [-5.22, -1.55]* | -5.09 [-7.2, -3.1]* | 3.68 [1.83, 5.61]* | -5.42 [-7.52, -3.33]* | -1.73 [-3.87, 0.4] |
| L 311 | L 295 | L Basolateral amygdalar nucleus, posterior part | 17.26 ± 9.77 | 13 | 8.51 ± 3.80 | 13 | 103.56 ± 22.26 | 9 | 161.51 ± 32.43 | 9 | 5.26 [3.36, 7.17]* | -0.16 [-1.87, 1.56] | 2.02 [-0.69, 4.73] | 0.24 [-1.49, 1.89] | -5.86 [-7.64, -3.89]* | -8.85 [-10.81, -6.85]* | 6.08 [4.15, 7.97]* | -9.08 [-11.09, -7.15]* | -3 [-5.14, -0.9]* |
| R 451 | R 295 | R Basolateral amygdalar nucleus, ventral part | 71.38 ± 58.75 | 14 | 8.95 ± 3.05 | 11 | 93.49 ± 13.87 | 9 | 137.15 ± 29.84 | 8 | 0.6 [-1.29, 2.45] | -0.44 [-2.21, 1.31] | -0.39 [-3.08, 2.39] | | | | | | |
| L 451 | L 295 | L Basolateral amygdalar nucleus, ventral part | 51.17 ± 47.51 | 12 | 4.39 ± 2.48 | 12 | 74.52 ± 19.59 | 8 | 114.95 ± 23.46 | 9 | 0.79 [-1.17, 2.76] | -0.5 [-2.29, 1.28] | -0.23 [-3.06, 2.55] | | | | | | |
| R 319 | R 703 | R Basomedial amygdalar nucleus | 16.79 ± 8.23 | 14 | 5.79 ± 1.38 | 13 | 58.64 ± 12.08 | 10 | 55.45 ± 10.92 | 10 | 2.62 [0.76, 4.45]* | -0.14 [-1.87, 1.56] | -1.39 [-4.06, 1.27] | 0.2 [-1.49, 1.89] | -3.2 [-5.07, -1.42]* | -2.83 [-4.69, -0.73]* | 3.41 [1.55, 5.34]* | -3.05 [-5.03, -1.11]* | 0.38 [-1.73, 2.41] |
| L 319 | L 703 | L Basomedial amygdalar nucleus | 17.86 ± 13.73 | 15 | 4.98 ± 1.81 | 13 | 30.35 ± 6.79 | 11 | 61.05 ± 13.40 | 10 | 0.95 [-0.82, 2.75] | -0.48 [-2.17, 1.23] | 0.71 [-1.87, 3.31] | | | | | | |

| | | | | | | | | | | | | | | | | | | | |
|---|---|---|---|---|---|---|---|---|---|---|---|---|---|---|---|---|---|---|---|
| R 327 | R 319 | R Basomedial amygdalar nucleus, anterior part | 28.67 ± 9.27 | 14 | 15.31 ± 5.53 | 12 | 80.67 ± 27.17 | 10 | 68.09 ± 11.54 | 10 | 1.94 [0.1, 3.75]* | 0.08 [-1.67, 1.81] | -1.84 [-4.45, 0.86] | -0.01 [-1.73, 1.66] | -2.53 [-4.35, -0.76]* | -1.91 [-3.78, 0.09] | 2.52 [0.62, 4.37]* | -1.89 [-3.96, 0.03] | 0.63 [-1.49, 2.63] |
| L 327 | L 319 | L Basomedial amygdalar nucleus, anterior part | 23.09 ± 16.70 | 15 | 7.29 ± 2.46 | 11 | 71.46 ± 22.20 | 11 | 86.26 ± 21.42 | 10 | 2.35 [0.59, 4.15]* | -0.31 [-2.07, 1.43] | -0.55 [-3.19, 2.08] | 0.37 [-1.36, 2.08] | -2.95 [-4.7, -1.2]* | -3.23 [-5.1, -1.23]* | 3.31 [1.54, 5.25]* | -3.59 [-5.59, -1.62]* | -0.28 [-2.25, 1.75] |
| R 334 | R 319 | R Basomedial amygdalar nucleus, posterior part | 38.57 ± 27.60 | 13 | 8.97 ± 1.99 | 11 | 163.37 ± 31.18 | 9 | 158.56 ± 38.99 | 8 | 3.3 [1.41, 5.24]* | -0.2 [-1.97, 1.57] | -1.1 [-3.92, 1.68] | 0.27 [-1.48, 2.08] | -3.88 [-5.77, -2.03]* | -3.73 [-5.69, -1.6]* | 4.16 [2.14, 6.05]* | -3.99 [-6.03, -1.86]* | 0.16 [-1.98, 2.37] |
| L 334 | L 319 | L Basomedial amygdalar nucleus, posterior part | 50.46 ± 39.91 | 14 | 8.44 ± 3.82 | 13 | 56.75 ± 12.04 | 9 | 160.46 ± 34.00 | 9 | 0.15 [-1.71, 2.05] | -0.52 [-2.22, 1.21] | 1.25 [-1.48, 3.98] | | | | | | |
| R 780 | R 703 | R Posterior amygdalar nucleus | 29.70 ± 16.76 | 12 | 29.28 ± 11.34 | 12 | 73.91 ± 27.78 | 10 | 124.88 ± 27.95 | 8 | 1.17 [-0.71, 3.03] | 0.48 [-1.31, 2.24] | 0.48 [-2.25, 3.26] | | | | | | |
| L 780 | L 703 | L Posterior amygdalar nucleus | 32.66 ± 22.10 | 12 | 17.03 ± 9.75 | 13 | 22.03 ± 10.62 | 10 | 53.02 ± 16.70 | 9 | | | | | | | | | |
| R 477 | R 623 | R Striatum | 11.03 ± 2.39 | 15 | 10.88 ± 1.47 | 15 | 36.84 ± 5.41 | 12 | 40.83 ± 3.92 | 10 | 1.58 [-2.06, 5.12] | 0.05 [-3.3, 3.41] | -0.88 [-6.1, 4.55] | | | | | | |
| L 477 | L 623 | L Striatum | 9.90 ± 3.18 | 15 | 13.25 ± 2.64 | 15 | 25.06 ± 3.17 | 12 | 32.84 ± 2.44 | 10 | 0.76 [-2.85, 4.36] | 0.42 [-2.99, 3.8] | -0.79 [-6.18, 4.53] | | | | | | |
| R 485 | R 477 | R Striatum dorsal region | 11.71 ± 1.98 | 15 | 11.62 ± 1.82 | 15 | 30.11 ± 5.23 | 12 | 28.73 ± 5.48 | 10 | 0.79 [-2.69, 4.34] | 0.06 [-3.28, 3.46] | -1.32 [-6.66, 3.88] | | | | | | |
| L 485 | L 477 | L Striatum dorsal region | 9.98 ± 1.91 | 15 | 20.59 ± 6.41 | 15 | 18.39 ± 3.30 | 12 | 23.70 ± 5.47 | 10 | 0.07 [-3.53, 3.67] | 1.14 [-2.25, 4.51] | -1.76 [-6.98, 3.61] | | | | | | |
| R 672 | R 485 | R Caudoputamen | 25.08 ± 3.86 | 14 | 24.89 ± 3.49 | 14 | 60.22 ± 10.46 | 12 | 63.83 ± 9.96 | 9 | 0.52 [-3.09, 4.18] | 0.08 [-3.49, 3.55] | -0.94 [-6.42, 4.52] | | | | | | |
| L 672 | L 485 | L Caudoputamen | 23.02 ± 3.71 | 13 | 41.18 ± 12.82 | 15 | 36.77 ± 6.59 | 12 | 52.66 ± 10.72 | 9 | -0.1 [-3.75, 3.52] | 0.94 [-2.5, 4.4] | -1.25 [-6.6, 4.02] | | | | | | |
| R 493 | R 477 | R Striatum ventral region | 18.55 ± 4.39 | 15 | 16.00 ± 2.36 | 14 | 58.79 ± 6.41 | 12 | 62.97 ± 9.01 | 10 | 1.39 [-2.19, 4.9] | -0.04 [-3.5, 3.41] | -0.89 [-6.14, 4.33] | | | | | | |
| L 493 | L 477 | L Striatum ventral region | 17.36 ± 4.85 | 15 | 18.10 ± 3.50 | 14 | 60.48 ± 6.20 | 12 | 67.55 ± 13.23 | 10 | 1.7 [-1.81, 5.23] | 0.14 [-3.28, 3.58] | -0.89 [-6.24, 4.34] | | | | | | |
| R 56 | R 493 | R Nucleus accumbens | 28.64 ± 6.26 | 12 | 22.27 ± 1.91 | 12 | 149.21 ± 19.73 | 12 | 179.24 ± 16.54 | 8 | 3.54 [-0.18, 7.28] | -0.01 [-3.67, 3.77] | -0.25 [-5.96, 5.51] | | | | | | |
| L 56 | L 493 | L Nucleus accumbens | 21.39 ± 3.87 | 13 | 17.66 ± 2.92 | 13 | 147.04 ± 18.04 | 12 | 159.72 ± 16.88 | 8 | 5.16 [1.49, 8.74]* | 0.04 [-3.55, 3.64] | -0.78 [-6.38, 4.78] | 0.05 [-3.3, 3.7] | -6.03 [-9.54, -2.46]* | -6.57 [-10.53, -2.46]* | 6.03 [2.46, 9.46]* | -6.58 [-10.63, -2.67]* | -0.55 [-4.5, 3.57] |
| R 754 | R 493 | R Olfactory tubercle | 72.10 ± 19.61 | 13 | 61.79 ± 10.43 | 11 | 82.80 ± 11.83 | 9 | 91.05 ± 14.31 | 9 | -0.67 [-4.55, 3.22] | -0.06 [-3.74, 3.62] | -0.71 [-6.42, 5.06] | | | | | | |
| L 754 | L 493 | L Olfactory tubercle | 73.85 ± 23.46 | 13 | 68.94 ± 12.81 | 12 | 87.05 ± 10.73 | 12 | 78.94 ± 6.82 | 8 | | | | | | | | | |
| R 275 | R 477 | R Lateral septal complex | 13.31 ± 2.76 | 14 | 27.97 ± 8.03 | 12 | 217.10 ± 33.91 | 11 | 242.29 ± 31.38 | 10 | 14.58 [10.88, 18.19]* | 1.35 [-2.21, 5.02] | -0.6 [-6.07, 4.94] | -1.28 [-4.9, 2.17] | -15.42 [-18.94, -11.82]* | -17.45 [-21.22, -13.82]* | 14.15 [10.41, 17.89]* | -16.16 [-20.06, -12.43]* | -2.04 [-5.96, 1.91] |
| L 275 | L 477 | L Lateral septal complex | 6.44 ± 1.83 | 14 | 11.89 ± 2.78 | 12 | 124.10 ± 14.85 | 11 | 171.78 ± 17.69 | 10 | 17.5 [13.72, 21.14]* | 1.08 [-2.49, 4.75] | 5.16 [-0.29, 10.61] | -1 [-4.47, 2.55] | -18.38 [-21.78, -14.55]* | -25.89 [-29.58, -22.19]* | 17.33 [13.45, 20.96]* | -24.88 [-28.71, -21.18]* | -7.56 [-11.36, -3.54]* |
| R 242 | R 275 | R Lateral septal nucleus | 18.94 ± 3.90 | 14 | 44.23 ± 14.98 | 12 | 302.46 ± 44.31 | 11 | 358.21 ± 51.01 | 10 | 14.22 [10.58, 17.88]* | 1.58 [-2.03, 5.1] | 0.2 [-5.2, 5.65] | -1.51 [-4.98, 1.99] | -15.04 [-18.68, -11.56]* | -18.12 [-21.78, -14.42]* | 13.55 [9.84, 17.3]* | -16.65 [-20.27, -12.71]* | -3.08 [-7, 0.8] |
| L 242 | L 275 | L Lateral septal nucleus | 7.92 ± 2.28 | 14 | 14.16 ± 3.65 | 12 | 177.79 ± 19.45 | 11 | 259.15 ± 28.15 | 10 | 20.69 [16.97, 24.39]* | 1.04 [-2.57, 4.72] | 8.12 [2.56, 13.63]* | -0.97 [-4.42, 2.61] | -21.53 [-25.1, -17.86]* | -31.97 [-35.62, -28.2]* | 20.56 [16.93, 24.26]* | -31.05 [-34.73, -27.15]* | -10.44 [-14.21, -6.5]* |
| R 258 | R 242 | R Lateral septal nucleus, rostral (rostroventral) part | 39.71 ± 7.94 | 14 | 78.51 ± 33.06 | 11 | 682.80 ± 93.04 | 11 | 785.07 ± 123.41 | 10 | 15.42 [11.7, 19.05]* | 1.3 [-2.37, 4.98] | 0.15 [-5.38, 5.63] | -1.2 [-4.9, 2.23] | -16.26 [-19.74, -12.56]* | -19 [-22.69, -15.35]* | 15.05 [11.2, 18.96]* | -17.77 [-21.52, -13.84]* | -2.74 [-6.58, 1.18] |
| L 258 | L 242 | L Lateral septal nucleus, rostral (rostroventral) part | 16.28 ± 4.63 | 14 | 31.78 ± 7.94 | 11 | 388.50 ± 46.88 | 11 | 540.04 ± 55.75 | 10 | 22.14 [18.49, 25.82]* | 1.27 [-2.37, 4.93] | 6.89 [1.5, 12.33]* | -1.19 [-4.67, 2.39] | -22.96 [-26.51, -19.44]* | -32.44 [-36.14, -28.68]* | 21.77 [18.02, 25.51]* | -31.23 [-34.98, -27.22]* | -9.47 [-13.39, -5.72]* |
| R 266 | R 242 | R Lateral septal nucleus, ventral part | 25.33 ± 12.20 | 11 | 86.68 ± 44.01 | 6 | 331.69 ± 77.68 | 10 | 492.62 ± 98.12 | 8 | 11.24 [7.18, 15.34]* | 2.63 [-1.94, 7.26] | 2.67 [-3.82, 9.14] | -2.54 [-7.08, 1.95] | -12.08 [-16.22, -8.18]* | -18.68 [-22.89, -14.61]* | 9.51 [4.88, 14.08]* | -16.11 [-20.72, -11.09]* | -6.62 [-10.78, -2.3]* |
| L 266 | L 242 | L Lateral septal nucleus, ventral part | 7.76 ± 6.51 | 10 | 12.13 ± 5.67 | 8 | 285.96 ± 35.68 | 11 | 438.18 ± 100.78 | 8 | 35.16 [31.15, 39.14]* | 0.87 [-3.44, 5.12] | 17.31 [11.06, 23.33]* | -0.78 [-5.11, 3.27] | -36 [-39.81, -32.09]* | -55.44 [-59.63, -51.16]* | 35.23 [31.03, 39.36]* | -54.7 [-59.03, -50.13]* | -19.46 [-23.76, -15.5]* |
| R 333 | R 275 | R Septohippocampal nucleus | 52.76 ± 15.81 | 14 | 149.02 ± 46.55 | 7 | 477.08 ± 131.21 | 9 | 374.90 ± 137.25 | 6 | 7.2 [3.33, 11.11]* | 2.19 [-1.99, 6.37] | -5.05 [-11.44, 1.35] | -2.1 [-6.14, 2.04] | -8.03 [-11.83, -4.19]* | -6.49 [-10.64, -1.89]* | 5.92 [1.32, 10.2]* | -4.34 [-9.24, 0.61] | 1.57 [-3.23, 6.14] |
| L 333 | L 275 | L Septohippocampal nucleus | 22.66 ± 11.46 | 13 | 132.15 ± 49.38 | 7 | 339.67 ± 116.13 | 9 | 357.09 ± 141.35 | 6 | 13.18 [9.3, 17.12]* | 5.22 [0.97, 9.52]* | -5.42 [-11.84, 0.98] | -5.16 [-9.19, -0.83]* | -14 [-17.78, -10.16]* | -15.12 [-19.41, -10.65]* | 8.87 [4.44, 13.26]* | -9.94 [-14.87, -5.19]* | -1.08 [-5.79, 3.53] |
| R 278 | R 477 | R Striatum-like amygdalar nuclei | 19.13 ± 8.08 | 15 | 15.29 ± 5.72 | 14 | 27.18 ± 5.21 | 12 | 32.90 ± 6.42 | 10 | -0.36 [-3.94, 3.24] | -0.12 [-3.55, 3.35] | -0.73 [-6.07, 4.6] | | | | | | |
| L 278 | L 477 | L Striatum-like amygdalar nuclei | 21.37 ± 12.64 | 15 | 8.97 ± 2.41 | 14 | 16.01 ± 3.23 | 12 | 29.88 ± 7.32 | 10 | -1.01 [-4.64, 2.59] | -0.49 [-3.98, 2.98] | -0.02 [-5.4, 5.22] | | | | | | |
| R 23 | R 278 | R Anterior amygdalar area | 45.65 ± 30.06 | 13 | 10.06 ± 4.43 | 9 | 50.15 ± 21.99 | 10 | 80.85 ± 28.71 | 6 | -0.27 [-4.13, 3.52] | -0.39 [-4.39, 3.52] | -0.74 [-6.81, 5.5] | | | | | | |
| R 536 | R 278 | R Central amygdalar nucleus | 13.55 ± 4.78 | 15 | 8.79 ± 3.61 | 14 | 28.15 ± 8.17 | 11 | 48.53 ± 11.11 | 10 | 0.12 [-3.61, 3.75] | -0.27 [-3.74, 3.21] | 0.8 [-4.48, 6.12] | | | | | | |
| L 536 | L 278 | L Central amygdalar nucleus | 25.99 ± 17.15 | 14 | 8.86 ± 3.22 | 13 | 23.94 ± 6.53 | 12 | 39.82 ± 10.38 | 10 | -0.85 [-4.44, 2.76] | -0.57 [-4.1, 2.89] | 0.01 [-5.3, 5.38] | | | | | | |
| R 544 | R 536 | R Central amygdalar nucleus, capsular part | 20.64 ± 4.01 | 14 | 19.51 ± 7.53 | 14 | 87.67 ± 27.05 | 10 | 179.49 ± 43.15 | 10 | 2.24 [-1.51, 6.05] | 0.08 [-3.38, 3.54] | 3.48 [-2.04, 8.91] | | | | | | |
| L 544 | L 536 | L Central amygdalar nucleus, capsular part | 29.55 ± 9.83 | 13 | 26.91 ± 8.25 | 13 | 78.56 ± 9.38 | 12 | 121.58 ± 29.46 | 10 | 0.94 [-2.79, 4.67] | 0.09 [-3.56, 3.74] | 0.2 [-5.25, 5.53] | | | | | | |
| R 551 | R 536 | R Central amygdalar nucleus, lateral part | 34.65 ± 9.90 | 14 | 17.59 ± 7.38 | 14 | 57.95 ± 25.49 | 10 | 82.58 ± 29.32 | 10 | -0.35 [-4.12, 3.36] | -0.34 [-3.81, 3.17] | 0.21 [-5.29, 5.71] | | | | | | |
| L 551 | L 536 | L Central amygdalar nucleus, lateral part | 49.60 ± 29.61 | 13 | 15.11 ± 7.72 | 13 | 37.64 ± 12.09 | 12 | 56.78 ± 22.88 | 10 | -0.95 [-4.63, 2.71] | -0.51 [-4.14, 3.16] | -0.28 [-5.85, 5.27] | | | | | | |
| R 559 | R 536 | R Central amygdalar nucleus, medial part | 28.99 ± 13.37 | 15 | 17.74 ± 7.59 | 13 | 45.41 ± 11.56 | 11 | 67.77 ± 16.11 | 10 | -0.4 [-4.08, 3.25] | -0.27 [-3.8, 3.25] | 0.09 [-5.32, 5.67] | | | | | | |
| L 559 | L 536 | L Central amygdalar nucleus, medial part | 61.36 ± 45.46 | 14 | 14.59 ± 6.72 | 13 | 38.71 ± 18.03 | 12 | 74.22 ± 21.46 | 10 | -1.15 [-4.69, 2.53] | -0.65 [-4.15, 2.91] | 0.06 [-5.38, 5.37] | | | | | | |
| R 1105 | R 278 | R Intercalated amygdalar nucleus | 24.56 ± 11.42 | 15 | 27.85 ± 15.48 | 13 | 77.50 ± 24.56 | 12 | 118.48 ± 58.08 | 10 | 1.38 [-2.21, 5] | 0.25 [-3.28, 3.79] | 0.26 [-5.25, 5.72] | | | | | | |
| L 1105 | L 278 | L Intercalated amygdalar nucleus | 9.47 ± 4.97 | 15 | 15.42 ± 7.54 | 13 | 48.35 ± 18.71 | 12 | 145.47 ± 51.42 | 10 | 3.35 [-0.2, 6.86] | 0.75 [-2.74, 4.2] | 8.35 [2.98, 13.71]* | -0.67 [-4.11, 2.62] | -4.17 [-7.6, -0.73]* | -14.59 [-18.14, -10.92]* | 3.51 [-0.21, 6.91] | -13.93 [-17.51, -10.04]* | -10.4 [-14.07, -6.41]* |
| R 403 | R 278 | R Medial amygdalar nucleus | 61.02 ± 30.40 | 15 | 42.56 ± 15.57 | 14 | 70.43 ± 12.83 | 12 | 62.93 ± 13.26 | 10 | -0.62 [-4.21, 3.05] | -0.22 [-3.63, 3.2] | -1.06 [-6.36, 4.26] | | | | | | |
| L 403 | L 278 | L Medial amygdalar nucleus | 54.00 ± 35.75 | 15 | 23.55 ± 6.66 | 14 | 26.18 ± 4.56 | 12 | 45.30 ± 8.65 | 10 | -1.28 [-4.83, 2.28] | -0.47 [-3.88, 2.87] | -0.34 [-5.78, 4.95] | | | | | | |
| R 803 | R 623 | R Pallidum | 10.95 ± 3.33 | 15 | 13.69 ± 3.79 | 14 | 24.96 ± 8.77 | 11 | 31.76 ± 6.76 | 10 | 1.37 [-0.02, 2.77] | 0.02 [-1.3, 1.33] | -0.23 [-2.28, 1.78] | | | | | | |
| L 803 | L 623 | L Pallidum | 7.46 ± 2.51 | 15 | 12.27 ± 4.28 | 14 | 16.37 ± 4.73 | 12 | 25.01 ± 4.72 | 10 | 1.4 [0, 2.78] | 0.44 [-0.89, 1.77] | -0.21 [-2.27, 1.87] | -0.55 [-1.88, 0.83] | -1.71 [-3.14, -0.33]* | -2.83 [-4.41, -1.19]* | 1.15 [-0.35, 2.63] | -2.27 [-3.91, -0.69]* | -1.12 [-2.89, 0.43] |
| R 818 | R 803 | R Pallidum, dorsal region | 8.80 ± 4.46 | 15 | 10.34 ± 6.62 | 13 | 3.51 ± 0.84 | 12 | 11.72 ± 2.12 | 10 | | | | | | | | | |
| L 818 | L 803 | L Pallidum, dorsal region | 5.72 ± 3.18 | 15 | 8.29 ± 6.27 | 13 | 3.81 ± 0.92 | 12 | 9.21 ± 1.79 | 10 | -0.14 [-1.52, 1.24] | 0.37 [-0.99, 1.71] | -0.34 [-2.4, 1.67] | | | | | | |
| R 1022 | R 818 | R Globus pallidus, external segment | 21.06 ± 11.51 | 15 | 25.02 ± 17.10 | 13 | 7.78 ± 1.60 | 11 | 24.15 ± 5.38 | 10 | -0.53 [-1.96, 0.86] | 0.1 [-1.25, 1.44] | -0.15 [-2.21, 1.89] | | | | | | |
| L 1022 | L 818 | L Globus pallidus, external segment | 16.80 ± 11.74 | 14 | 19.60 ± 16.92 | 13 | 6.70 ± 1.68 | 12 | 18.06 ± 4.68 | 10 | -0.39 [-1.8, 1.01] | 0.09 [-1.27, 1.45] | -0.35 [-2.38, 1.73] | | | | | | |
| R 1031 | R 818 | R Globus pallidus, internal segment | 16.00 ± 5.51 | 14 | 19.15 ± 11.89 | 12 | 7.70 ± 2.40 | 10 | 21.75 ± 4.34 | 10 | -0.37 [-1.76, 1.06] | 0.12 [-1.27, 1.49] | -0.06 [-2.15, 2.04] | | | | | | |
| L 1031 | L 818 | L Globus pallidus, internal segment | 10.01 ± 2.81 | 13 | 11.39 ± 5.68 | 13 | 7.65 ± 2.56 | 12 | 16.85 ± 6.38 | 9 | 0.05 [-1.37, 1.45] | 0.14 [-1.22, 1.5] | 0.02 [-2.04, 2.1] | | | | | | |
| R 835 | R 803 | R Pallidum, ventral region | 14.28 ± 4.00 | 15 | 8.45 ± 2.10 | 14 | 21.11 ± 5.53 | 12 | 27.03 ± 5.61 | 10 | 0.67 [-0.73, 2.06] | -0.63 [-2, 0.69] | 0.13 [-1.91, 2.19] | | | | | | |
| L 835 | L 803 | L Pallidum, ventral region | 11.95 ± 3.31 | 15 | 8.17 ± 1.88 | 14 | 18.32 ± 4.56 | 12 | 29.25 ± 5.98 | 10 | 0.73 [-0.65, 2.1] | -0.53 [-1.86, 0.81] | 0.53 [-1.5, 2.54] | | | | | | |
| R 342 | R 835 | R Substantia innominata | 29.16 ± 8.25 | 15 | 16.54 ± 4.23 | 14 | 40.64 ± 11.01 | 12 | 58.74 ± 9.68 | 9 | 0.58 [-0.76, 1.98] | -0.65 [-2, 0.66] | 0.43 [-1.61, 2.44] | | | | | | |
| L 342 | L 835 | L Substantia innominata | 24.62 ± 6.84 | 15 | 16.99 ± 3.89 | 13 | 36.13 ± 9.26 | 12 | 60.74 ± 8.49 | 9 | 0.66 [-0.73, 2.06] | -0.57 [-1.93, 0.74] | 0.73 [-1.33, 2.85] | | | | | | |
| R 826 | R 803 | R Pallidum, medial region | 32.88 ± 8.04 | 14 | 54.80 ± 10.86 | 11 | 129.94 ± 45.83 | 12 | 140.21 ± 38.18 | 9 | 3.14 [1.76, 4.55]* | 0.82 [-0.58, 2.2] | -1.49 [-3.6, 0.58] | -0.93 [-2.29, 0.51] | -3.44 [-4.82, -2.02]* | -3.67 [-5.35, -2.08]* | 2.52 [0.96, 4.01]* | -2.74 [-4.38, -1.06]* | -0.22 [-1.87, 1.42] |
| L 826 | L 803 | L Pallidum, medial region | 23.24 ± 7.98 | 14 | 50.37 ± 15.84 | 10 | 75.50 ± 24.77 | 12 | 97.42 ± 24.45 | 9 | 2.45 [1.07, 3.86]* | 1.31 [-0.13, 2.74] | -1.34 [-3.46, 0.77] | -1.42 [-2.84, 0.02] | -2.75 [-4.27, -1.42]* | -3.6 [-5.29, -1.95]* | 1.35 [-0.27, 2.88] | -2.19 [-3.89, -0.45]* | -0.85 [-2.53, 0.81] |
| R 904 | R 826 | R Medial septal complex | 49.32 ± 12.06 | 14 | 82.91 ± 16.02 | 11 | 175.64 ± 58.33 | 12 | 188.96 ± 52.37 | 9 | 2.77 [1.36, 4.18]* | 0.85 [-0.55, 2.27] | -1.56 [-3.65, 0.59] | -0.96 [-2.41, 0.46] | -3.08 [-4.47, -1.62]* | -3.25 [-4.85, -1.58]* | 2.11 [0.54, 3.64]* | -2.29 [-3.92, -0.54]* | -0.18 [-1.85, 1.46] |
| L 904 | L 826 | L Medial septal complex | 34.86 ± 11.97 | 14 | 75.92 ± 23.68 | 10 | 107.54 ± 35.01 | 12 | 142.33 ± 35.54 | 9 | 2.28 [0.9, 3.69]* | 1.32 [-0.08, 2.72] | -1.3 [-3.45, 0.76] | -1.43 [-2.8, 0.03] | -2.59 [-4.01, -1.19]* | -3.5 [-5.1, -1.83]* | 1.16 [-0.48, 2.66] | -2.06 [-3.73, -0.39]* | -0.91 [-2.53, 0.81] |
| R 596 | R 904 | R Diagonal band nucleus | 100.10 ± 24.59 | 14 | 165.86 ± 34.16 | 11 | 314.18 ± 108.74 | 12 | 306.71 ± 102.76 | 9 | 2.34 [0.95, 3.77]* | 0.82 [-0.59, 2.2] | -1.87 [-3.99, 0.27] | -0.93 [-2.33, 0.5] | -2.64 [-4.1, -1.25]* | -2.48 [-4.17, -0.89]* | 1.71 [0.19, 3.29]* | -1.54 [-3.26, 0.09] | 0.16 [-1.55, 1.81] |
| L 596 | L 904 | L Diagonal band nucleus | 69.79 ± 24.71 | 14 | 150.60 ± 47.34 | 10 | 208.20 ± 67.34 | 12 | 250.01 ± 65.77 | 9 | 2.18 [0.8, 3.57]* | 1.3 [-0.08, 2.71] | -1.68 [-3.79, 0.41] | -1.41 [-2.8, 0.05] | -2.49 [-3.88, -1.09]* | -3 [-4.59, -1.32]* | 1.08 [-0.47, 2.59] | -1.58 [-3.27, 0.12] | -0.51 [-2.24, 1.1] |
| R 549 | R 1129 | R Thalamus | 20.10 ± 11.44 | 15 | 18.26 ± 6.07 | 15 | 25.35 ± 3.49 | 12 | 43.46 ± 10.29 | 10 | -0.33 [-4.08, 3.46] | -0.03 [-3.52, 3.58] | -0.23 [-5.81, 5.33] | | | | | | |
| L 549 | L 1129 | L Thalamus | 5.43 ± 1.80 | 14 | 6.69 ± 2.00 | 15 | 14.90 ± 2.66 | 12 | 25.71 ± 5.24 | 10 | 1.19 [-2.68, 5.01] | 0.31 [-3.25, 3.84] | 0.53 [-4.98, 6.21] | | | | | | |
| R 864 | R 549 | R Thalamus, sensory-motor cortex related | 17.48 ± 11.66 | 15 | 14.61 ± 4.45 | 15 | 25.15 ± 4.45 | 12 | 42.37 ± 10.76 | 10 | -0.12 [-3.92, 3.73] | -0.09 [-3.63, 3.44] | -0.07 [-5.73, 5.61] | | | | | | |
| L 864 | L 549 | L Thalamus, sensory-motor cortex related | 2.79 ± 0.92 | 14 | 4.57 ± 1.40 | 15 | 15.30 ± 3.16 | 12 | 26.39 ± 5.49 | 10 | 3.9 [0.09, 7.69]* | 0.72 [-2.84, 4.34] | 2.14 [-3.34, 7.68] | -0.67 [-4.18, 2.84] | -4.73 [-8.47, -0.94]* | -9 [-13.07, -5.13]* | 4.05 [0.34, 7.79]* | -8.32 [-12.39, -4.58]* | -4.27 [-8.42, -0.33]* |
| R 637 | R 864 | R Ventral group of the dorsal thalamus | 15.51 ± 10.86 | 15 | 11.38 ± 6.16 | 15 | 23.34 ± 3.89 | 12 | 34.53 ± 6.14 | 10 | -0.1 [-3.87, 3.67] | -0.23 [-3.77, 3.26] | -0.2 [-5.71, 5.44] | | | | | | |
| L 637 | L 864 | L Ventral group of the dorsal thalamus | 3.22 ± 1.12 | 14 | 4.05 ± 1.49 | 15 | 18.55 ± 3.68 | 12 | 31.14 ± 6.94 | 10 | 4.21 [0.42, 8.07]* | 0.34 [-3.13, 4.02] | 2.44 [-3.13, 8.01] | -0.29 [-3.9, 3.14] | -5.04 [-9.05, -1.55]* | -9.25 [-13.39, -5.25]* | 4.75 [1.23, 8.55]* | -8.96 [-12.94, -4.91]* | -4.19 [-8.41, -0.06]* |
| R 629 | R 637 | R Ventral anterior-lateral complex of the thalamus | 9.12 ± 3.13 | 15 | 17.55 ± 12.40 | 13 | 65.36 ± 18.51 | 12 | 69.37 ± 14.29 | 10 | 5.59 [1.76, 9.31]* | 0.93 [-2.69, 4.61] | -1.65 [-7.25, 3.95] | -0.86 [-4.6, 2.51] | -6.4 [-10.03, -2.57]* | -7.11 [-11.25, -3.32]* | 5.53 [1.58, 9.16]* | -6.2 [-10.2, -2.19]* | -0.71 [-4.72, 3.49] |
| L 629 | L 637 | L Ventral anterior-lateral complex of the thalamus | 5.87 ± 2.40 | 14 | 13.22 ± 6.49 | 13 | 39.66 ± 9.69 | 12 | 78.12 ± 25.98 | 10 | 5.19 [1.27, 9.01]* | 1.29 [-2.5, 5.04] | 4.1 [-1.59, 9.9] | -1.24 [-4.99, 2.38] | -6.02 [-9.79, -2.18]* | -12.8 [-16.85, -8.86]* | 4.78 [1.03, 8.59]* | -11.56 [-15.72, -7.64]* | -6.82 [-10.9, -2.57]* |
| R 685 | R 637 | R Ventral medial nucleus of the thalamus | 9.14 ± 3.17 | 15 | 37.64 ± 21.93 | 14 | 53.02 ± 8.69 | 12 | 77.09 ± 12.81 | 10 | 4.22 [0.38, 7.95]* | 3.12 [-0.41, 6.59] | -1.66 [-7.13, 3.98] | -3.06 [-6.54, 0.39] | -5.07 [-8.78, -1.34]* | -7.9 [-11.96, -4.08]* | 1.95 [-1.82, 5.88] | -4.85 [-8.77, -0.85]* | -2.88 [-7.22, 1.17] |
| L 685 | L 637 | L Ventral medial nucleus of the thalamus | 9.89 ± 5.38 | 14 | 12.47 ± 5.40 | 14 | 41.80 ± 8.97 | 12 | 69.33 ± 20.04 | 10 | 2.68 [-1.14, 6.49] | 0.29 [-3.38, 3.93] | 1.34 [-4.23, 6.96] | | | | | | |
| R 709 | R 637 | R Ventral posterior complex of the thalamus | 31.99 ± 24.77 | 15 | 16.48 ± 8.81 | 15 | 31.36 ± 5.32 | 12 | 47.66 ± 10.18 | 10 | -0.63 [-4.33, 3.07] | -0.46 [-3.97, 3.07] | -0.2 [-5.74, 5.34] | | | | | | |
| L 709 | L 637 | L Ventral posterior complex of the thalamus | 4.75 ± 1.64 | 13 | 4.26 ± 2.01 | 15 | 25.97 ± 5.19 | 12 | 43.42 ± 9.24 | 10 | 3.99 [0.19, 7.88]* | 0.05 [-3.56, 3.68] | 2.45 [-3.13, 8.09] | -0.01 [-3.56, 3.5] | -4.83 [-8.68, -1.14]* | -8.72 [-12.77, -4.66]* | 4.82 [1.03, 8.36]* | -8.74 [-12.71, -4.88]* | -3.91 [-7.93, 0.29] |
| R 718 | R 709 | R Ventral posterolateral nucleus of the thalamus | 128.35 ± 93.72 | 15 | 48.32 ± 25.65 | 15 | 56.55 ± 8.03 | 12 | 85.67 ± 13.99 | 10 | -1.13 [-4.89, 2.62] | -0.56 [-4.1, 2.98] | -0.39 [-5.89, 5.18] | | | | | | |
| L 718 | L 709 | L Ventral posterolateral nucleus of the thalamus | 16.85 ± 7.04 | 13 | 10.88 ± 4.50 | 15 | 49.65 ± 10.89 | 12 | 98.11 ± 28.94 | 10 | 1.45 [-2.43, 5.25] | -0.19 [-3.76, 3.47] | 1.92 [-3.7, 7.58] | | | | | | |
| R 725 | R 709 | R Ventral posterolateral nucleus of the thalamus, parvicellular part | 38.35 ± 24.97 | 11 | 6.53 ± 2.14 | 12 | 135.09 ± 57.56 | 9 | 88.12 ± 19.81 | 7 | 2.59 [-1.79, 6.88] | -0.22 [-4.29, 3.79] | -1.79 [-8.07, 4.47] | | | | | | |
| L 725 | L 709 | L Ventral posterolateral nucleus of the thalamus, parvicellular part | 4.87 ± 3.79 | 11 | 3.51 ± 1.92 | 11 | 57.63 ± 16.45 | 8 | 63.30 ± 21.19 | 7 | 10.25 [5.9, 14.58]* | -0.08 [-4.13, 3.86] | 0.93 [-5.44, 7.41] | 0.13 [-3.75, 4.13] | -11.06 [-15.38, -6.84]* | -13.34 [-17.93, -8.76]* | 11.19 [6.82, 15.43]* | -13.46 [-18.28, -8.92]* | -2.26 [-6.98, 2.67] |
| R 733 | R 709 | R Ventral posteromedial nucleus of the thalamus | 25.91 ± 20.76 | 14 | 26.36 ± 12.92 | 15 | 64.74 ± 13.91 | 12 | 107.63 ± 28.63 | 10 | 0.96 [-2.88, 4.87] | 0.14 [-3.47, 3.8] | 0.38 [-5.25, 6] | | | | | | |
| L 733 | L 709 | L Ventral posteromedial nucleus of the thalamus | 5.90 ± 2.79 | 13 | 7.65 ± 4.50 | 15 | 52.26 ± 9.56 | 12 | 86.59 ± 19.14 | 10 | 7.34 [3.48, 11.19]* | 0.44 [-3.12, 4.1] | 4.25 [-1.32, 9.76] | -0.39 [-3.94, 3.22] | -8.16 [-12.19, -4.57]* | -14.24 [-18.37, -10.31]* | 7.78 [4.12, 11.5]* | -13.86 [-17.86, -10.03]* | -6.13 [-10.39, -2.25]* |
| R 741 | R 709 | R Ventral posteromedial nucleus of the thalamus, parvicellular part | 6.20 ± 4.23 | 9 | 4.38 ± 1.70 | 11 | 62.10 ± 16.54 | 12 | 52.47 ± 14.93 | 7 | 8.06 [3.85, 12.26]* | -0.27 [-4.43, 3.98] | -1.84 [-8.13, 4.38] | 0.31 [-3.83, 4.51] | -8.89 [-13.01, -4.79]* | -8.23 [-12.7, -3.25]* | 9.19 [5.22, 13.17]* | -8.53 [-12.96, -4.03]* | 0.69 [-3.81, 5.25] |
| L 741 | L 709 | L Ventral posteromedial nucleus of the thalamus, parvicellular part | 2.07 ± 1.07 | 8 | 3.56 ± 2.66 | 13 | 58.91 ± 23.08 | 10 | 54.65 ± 17.29 | 7 | 26.43 [21.93, 30.86]* | 0.18 [-4.16, 4.45] | -2.75 [-9.24, 3.64] | -0.13 [-4.24, 4.22] | -27.26 [-31.61, -22.77]* | -26.08 [-31.23, -21.59]* | 27.13 [22.99, 30.99]* | -25.99 [-30.36, -21.57]* | 1.16 [-3.36, 5.92] |
| R 563807435 | R 637 | R Posterior triangular thalamic nucleus | 69.49 ± 46.29 | 9 | 11.19 ± 7.21 | 9 | 32.61 ± 13.28 | 9 | 96.76 ± 21.06 | 7 | -0.97 [-5.5, 3.5] | -0.73 [-5.16, 3.7] | 2 [-4.67, 8.62] | | | | | | |
| L 563807435 | L 637 | L Posterior triangular thalamic nucleus | 4.36 ± 2.35 | 9 | 6.77 ± 2.87 | 7 | 47.28 ± 16.60 | 9 | 58.52 ± 17.80 | 7 | 9.43 [5.05, 13.94]* | 1.5 [-3.12, 6.22] | 1.55 [-5.17, 8.24] | -1.45 [-6, 3.23] | -10.24 [-14.55, -5.87]* | -14.7 [-19.48, -10.07]* | 8.81 [4.12, 13.55]* | -13.25 [-18.28, -8.26]* | -4.44 [-9.3, -0.02]* |
| R 406 | R 864 | R Subparafascicular nucleus | 44.16 ± 28.24 | 11 | 6.49 ± 5.38 | 13 | 29.21 ± 9.23 | 10 | 52.60 ± 8.26 | 8 | -0.85 [-5.01, 3.4] | -0.84 [-4.78, 3.11] | 1.32 [-4.7, 7.39] | | | | | | |
| L 406 | L 864 | L Subparafascicular nucleus | 8.44 ± 5.00 | 11 | 5.69 ± 2.85 | 12 | 24.15 ± 9.92 | 10 | 32.98 ± 7.14 | 8 | 1.38 [-2.85, 5.54] | -0.08 [-4.04, 3.83] | 1.07 [-5.16, 7.14] | | | | | | |
| R 414 | R 406 | R Subparafascicular nucleus, magnocellular part | 25.76 ± 18.91 | 6 | 1.32 ± 1.32 | 8 | 56.26 ± 19.56 | 7 | 58.22 ± 11.25 | 6 | 0.86 [-4.22, 5.91] | -0.8 [-5.77, 4.13] | -0.2 [-7.33, 6.97] | | | | | | |
| R 422 | R 406 | R Subparafascicular nucleus, parvicellular part | 107.27 ± 68.83 | 9 | 19.60 ± 15.39 | 9 | 52.50 ± 25.20 | 7 | 118.87 ± 29.15 | 7 | -1.02 [-5.73, 3.62] | -0.72 [-5.12, 3.68] | 1.76 [-4.99, 8.49] | | | | | | |
| L 422 | L 406 | L Subparafascicular nucleus, parvicellular part | 19.72 ± 13.16 | 9 | 4.39 ± 2.28 | 8 | 65.84 ± 29.07 | 8 | 94.85 ± 29.39 | 7 | 1.82 [-2.68, 6.2] | -0.76 [-5.2, 3.72] | 2.79 [-3.75, 9.42] | | | | | | |
| R 1008 | R 864 | R Geniculate group, dorsal thalamus | 97.46 ± 64.51 | 14 | 65.02 ± 23.10 | 15 | 94.50 ± 24.48 | 12 | 169.97 ± 83.25 | 9 | -0.55 [-4.44, 3.28] | -0.21 [-3.86, 3.37] | 0.85 [-4.79, 6.56] | | | | | | |
| L 1008 | L 864 | L Geniculate group, dorsal thalamus | 7.04 ± 2.94 | 14 | 19.72 ± 11.41 | 15 | 33.54 ± 7.36 | 12 | 65.35 ± 17.46 | 10 | 3.21 [-0.6, 7.12] | 1.91 [-1.68, 5.46] | 1.47 [-4.02, 6.98] | | | | | | |
| R 475 | R 1008 | R Medial geniculate complex | 6.88 ± 4.39 | 8 | 19.54 ± 9.22 | 8 | 32.87 ± 15.76 | 8 | 74.48 ± 24.96 | 7 | 2.94 [-1.59, 7.55] | 1.38 [-3.19, 5.9] | 4.93 [-1.81, 11.66] | | | | | | |
| L 475 | L 1008 | L Medial geniculate complex | 7.45 ± 2.53 | 8 | 13.83 ± 7.30 | 8 | 29.40 ± 12.34 | 9 | 52.94 ± 13.33 | 7 | 2.08 [-2.45, 6.58] | 1.37 [-3.18, 5.95] | 2.25 [-4.41, 8.87] | | | | | | |
| L 1072 | L 475 | L Medial geniculate complex, dorsal part | 12.41 ± 5.03 | 7 | 91.06 ± 54.05 | 6 | 40.99 ± 20.17 | 8 | 95.63 ± 17.73 | 7 | 2.12 [-2.74, 6.85] | 6.76 [1.7, 11.84]* | -2.57 [-9.54, 4.6] | -6.7 [-11.83, -1.83]* | -2.96 [-7.65, 1.85] | -8.56 [-13.31, -3.49]* | -3.77 [-8.86, 1.14] | -1.8 [-7.16, 3.25] | -5.59 [-10.43, -0.71]* |
| L 1079 | L 475 | L Medial geniculate complex, ventral part | 30.94 ± 12.78 | 6 | 31.94 ± 19.23 | 6 | 29.31 ± 18.12 | 8 | 90.43 ± 36.02 | 6 | -0.13 [-5.03, 4.88] | 0.52 [-4.73, 5.85] | 0.92 [-6.32, 8.27] | | | | | | |
| R 1088 | R 475 | R Medial geniculate complex, medial part | 15.67 ± 15.67 | 8 | 12.74 ± 5.66 | 8 | 69.02 ± 34.05 | 8 | 91.69 ± 37.99 | 7 | 2.59 [-2.04, 7.2] | -0.65 [-5.26, 3.89] | 2.34 [-4.23, 9.04] | | | | | | |
| L 1088 | L 475 | L Medial geniculate complex, medial part | 6.25 ± 2.52 | 8 | 10.63 ± 5.64 | 8 | 68.13 ± 26.35 | 9 | 120.21 ± 33.45 | 7 | 9.02 [4.48, 13.61]* | 1.2 [-3.35, 5.79] | 7.59 [0.9, 14.35]* | -1.13 [-5.63, 3.45] | -9.83 [-14.43, -5.4]* | -20.04 [-24.76, -15.05]* | 8.73 [4.14, 13.14]* | -18.89 [-23.58, -13.93]* | -10.18 [-14.95, -5.43]* |
| R 170 | R 1008 | R Dorsal part of the lateral geniculate complex | 150.07 ± 96.41 | 14 | 113.69 ± 38.51 | 15 | 169.40 ± 39.77 | 12 | 278.64 ± 123.91 | 9 | -0.36 [-4.17, 3.4] | -0.1 [-3.68, 3.46] | 0.69 [-4.92, 6.36] | | | | | | |
| L 170 | L 1008 | L Dorsal part of the lateral geniculate complex | 9.60 ± 4.67 | 14 | 31.23 ± 17.32 | 15 | 58.96 ± 10.55 | 12 | 104.54 ± 26.58 | 10 | 4.59 [0.82, 8.39]* | 2.36 [-1.24, 5.94] | 1.23 [-4.38, 6.81] | -2.31 [-5.86, 1.23] | -5.41 [-9.07, -1.6]* | -10.42 [-14.53, -6.6]* | 3.11 [-0.73, 6.61] | -8.14 [-12, -4.14]* | -5.02 [-9.13, -0.92]* |
| R 496345664 | R 170 | R Dorsal part of the lateral geniculate complex, shell | 388.57 ± 233.96 | 13 | 405.49 ± 168.04 | 14 | 432.40 ± 119.55 | 12 | 523.24 ± 263.57 | 9 | -0.38 [-4.27, 3.53] | 0.12 [-3.6, 3.84] | -0.04 [-5.91, 5.58] | | | | | | |
| L 496345664 | L 170 | L Dorsal part of the lateral geniculate complex, shell | 42.96 ± 22.57 | 14 | 139.70 ± 109.01 | 14 | 113.37 ± 33.45 | 11 | 171.97 ± 43.09 | 9 | 1.04 [-2.82, 4.96] | 2.86 [-0.79, 6.53] | -1.57 [-7.36, 4.14] | | | | | | |
| R 496345668 | R 170 | R Dorsal part of the lateral geniculate complex, core | 374.58 ± 226.87 | 14 | 184.68 ± 62.13 | 15 | 373.06 ± 113.51 | 12 | 513.57 ± 212.84 | 9 | -0.51 [-4.35, 3.22] | -0.4 [-4.04, 3.2] | 0.61 [-5.09, 6.3] | | | | | | |
| L 496345668 | L 170 | L Dorsal part of the lateral geniculate complex, core | 12.38 ± 5.65 | 14 | 62.29 ± 31.82 | 15 | 116.80 ± 21.13 | 12 | 208.32 ± 49.98 | 10 | 7.91 [4.11, 11.78]* | 4.15 [0.62, 7.69]* | 2.06 [-3.46, 7.68] | -4.11 [-7.74, -0.76]* | -8.7 [-12.54, -5.02]* | -16.37 [-20.27, -12.33]* | 4.62 [0.96, 8.43]* | -12.24 [-16.17, -8.29]* | -7.61 [-11.73, -3.52]* |
| R 496345672 | R 170 | R Dorsal part of the lateral geniculate complex, ipsilateral zone | 99.05 ± 65.13 | 12 | 255.88 ± 138.88 | 14 | 200.82 ± 74.45 | 12 | 805.96 ± 391.03 | 9 | 0.58 [-3.38, 4.58] | 1.67 [-2.13, 5.4] | 4.3 [-1.4, 10.21] | | | | | | |
| L 496345672 | L 170 | L Dorsal part of the lateral geniculate complex, ipsilateral zone | 17.71 ± 10.20 | 12 | 13.18 ± 7.24 | 15 | 128.28 ± 46.07 | 10 | 176.01 ± 64.80 | 9 | 5.81 [1.54, 9.95]* | -0.04 [-3.76, 3.79] | 2.61 [-3.3, 8.51] | 0.09 [-3.67, 3.78] | -6.65 [-10.78, -2.55]* | -10.62 [-14.99, -6.51]* | 6.7 [2.88, 10.73]* | -10.7 [-14.78, -6.53]* | -3.99 [-8.44, 0.36] |
| R 856 | R 549 | R Thalamus, polymodal association cortex related | 30.76 ± 16.73 | 15 | 29.17 ± 10.87 | 15 | 37.05 ± 4.79 | 12 | 64.38 ± 15.46 | 10 | -0.39 [-4.17, 3.32] | 0 [-3.56, 3.58] | -0.27 [-5.84, 5.3] | | | | | | |
| L 856 | L 549 | L Thalamus, polymodal association cortex related | 9.44 ± 3.11 | 15 | 11.07 ± 3.28 | 15 | 21.20 ± 3.68 | 12 | 37.45 ± 8.22 | 10 | 0.64 [-3.08, 4.46] | 0.23 [-3.34, 3.8] | 0.36 [-5.27, 5.86] | | | | | | |
| R 138 | R 856 | R Lateral group of the dorsal thalamus | 31.62 ± 14.89 | 14 | 38.92 ± 13.83 | 15 | 59.08 ± 8.95 | 12 | 118.55 ± 34.82 | 10 | 0.36 [-3.52, 4.2] | 0.35 [-3.31, 3.98] | 0.36 [-5.13, 5.93] | | | | | | |
| L 138 | L 856 | L Lateral group of the dorsal thalamus | 11.30 ± 5.98 | 14 | 17.34 ± 7.24 | 15 | 31.05 ± 4.71 | 12 | 64.58 ± 12.36 | 10 | 1.23 [-2.61, 5.03] | 0.67 [-2.95, 4.26] | 1.12 [-4.47, 6.84] | | | | | | |
| R 218 | R 138 | R Lateral posterior nucleus of the thalamus | 143.56 ± 81.82 | 14 | 166.37 ± 63.85 | 15 | 185.02 ± 37.15 | 12 | 441.34 ± 163.92 | 10 | -0.23 [-3.96, 3.48] | 0.28 [-3.25, 3.81] | 0.34 [-5.18, 5.87] | | | | | | |
| L 218 | L 138 | L Lateral posterior nucleus of the thalamus | 51.91 ± 33.73 | 14 | 67.20 ± 29.40 | 15 | 61.87 ± 13.06 | 12 | 147.33 ± 29.46 | 10 | -0.35 [-4.18, 3.41] | 0.4 [-3.15, 3.99] | 0.08 [-5.54, 5.64] | | | | | | |
| R 1020 | R 138 | R Posterior complex of the thalamus | 16.51 ± 6.34 | 14 | 8.77 ± 4.29 | 15 | 69.99 ± 12.18 | 12 | 102.37 ± 20.01 | 10 | 2.71 [-1.15, 6.52] | -0.36 [-3.96, 3.22] | 1.15 [-4.45, 6.71] | | | | | | |
| L 1020 | L 138 | L Posterior complex of the thalamus | 7.07 ± 2.46 | 14 | 8.70 ± 3.78 | 15 | 50.76 ± 7.97 | 12 | 106.25 ± 25.05 | 10 | 5.66 [1.82, 9.46]* | 0.35 [-3.17, 3.9] | 6.35 [0.79, 11.92]* | -0.31 [-3.89, 3.1] | -6.47 [-10.21, -2.68]* | -14.61 [-18.48, -10.53]* | 6.2 [2.3, 9.66]* | -14.31 [-18.33, -10.49]* | -8.12 [-12.31, -4.02]* |
| R 1029 | R 138 | R Posterior limiting nucleus of the thalamus | 96.89 ± 78.90 | 10 | 77.22 ± 47.96 | 9 | 87.13 ± 42.16 | 10 | 224.90 ± 61.40 | 8 | -0.53 [-4.78, 3.63] | -0.1 [-4.29, 4.2] | 1.44 [-4.86, 7.85] | | | | | | |
| L 1029 | L 138 | L Posterior limiting nucleus of the thalamus | 23.63 ± 12.52 | 11 | 13.06 ± 7.59 | 7 | 86.03 ± 20.70 | 10 | 111.88 ± 30.34 | 7 | 2.21 [-2.04, 6.39] | 0.43 [-4, 4.81] | 1.11 [-5.41, 7.77] | | | | | | |
| R 560581551 | R 138 | R Ethmoid nucleus of the thalamus | 9.78 ± 5.55 | 13 | 11.67 ± 4.13 | 15 | 97.74 ± 30.46 | 12 | 75.05 ± 18.10 | 10 | 8.5 [4.62, 12.35]* | 0.35 [-3.25, 4.06] | -3.83 [-9.39, 1.77] | -0.29 [-3.96, 3.18] | -9.31 [-13.15, -5.58]* | -7.25 [-11.36, -3.28]* | 9.03 [5.33, 12.69]* | -6.96 [-10.9, -2.96]* | 2.09 [-2.03, 6.17] |
| L 560581551 | L 138 | L Ethmoid nucleus of the thalamus | 4.91 ± 2.08 | 13 | 5.56 ± 3.49 | 15 | 87.72 ± 22.92 | 12 | 108.25 ± 24.31 | 10 | 16.36 [12.41, 20.28]* | 0.28 [-3.36, 3.95] | 2.75 [-2.86, 8.3] | -0.22 [-3.78, 3.4] | -17.17 [-21.01, -13.36]* | -21.61 [-25.83, -17.71]* | 16.96 [13.24, 20.77]* | -21.38 [-25.39, -17.51]* | -4.42 [-8.55, -0.33]* |
| R 239 | R 856 | R Anterior group of the dorsal thalamus | 70.40 ± 36.10 | 15 | 61.45 ± 23.79 | 13 | 77.17 ± 12.97 | 12 | 131.21 ± 37.69 | 10 | -0.48 [-4.23, 3.26] | -0.13 [-3.78, 3.48] | -0.26 [-5.88, 5.37] | | | | | | |
| L 239 | L 856 | L Anterior group of the dorsal thalamus | 11.54 ± 3.57 | 15 | 31.14 ± 12.66 | 13 | 40.59 ± 8.80 | 12 | 54.07 ± 11.32 | 10 | 1.92 [-1.81, 5.64] | 1.69 [-2, 5.31] | -1.68 [-7.27, 3.87] | | | | | | |
| R 255 | R 239 | R Anteroventral nucleus of thalamus | 30.60 ± 13.55 | 13 | 41.79 ± 26.63 | 9 | 121.32 ± 72.85 | 12 | 57.63 ± 20.20 | 8 | 2.42 [-1.51, 6.26] | 0.27 [-3.75, 4.3] | -2.03 [-8.11, 4.02] | | | | | | |
| L 255 | L 239 | L Anteroventral nucleus of thalamus | 10.43 ± 3.49 | 12 | 52.75 ± 47.70 | 9 | 40.02 ± 14.51 | 12 | 57.69 ± 23.59 | 8 | 2.15 [-1.96, 6.18] | 3.78 [-0.49, 7.95] | -1.9 [-8.21, 4.38] | | | | | | |
| R 127 | R 239 | R Anteromedial nucleus | 4.99 ± 1.51 | 13 | 16.19 ± 13.89 | 8 | 19.12 ± 7.66 | 12 | 18.18 ± 6.40 | 8 | 2.05 [-1.87, 5.94] | 2.23 [-2.02, 6.41] | -2.08 [-8.14, 3.99] | | | | | | |
| L 127 | L 239 | L Anteromedial nucleus | 5.54 ± 2.21 | 13 | 26.15 ± 23.18 | 9 | 12.91 ± 4.37 | 12 | 11.98 ± 4.64 | 8 | 0.54 [-3.3, 4.41] | 3.37 [-0.66, 7.4] | -3.23 [-9.19, 2.78] | | | | | | |
| R 1096 | R 127 | R Anteromedial nucleus, dorsal part | 9.49 ± 4.06 | 13 | 31.83 ± 26.10 | 8 | 34.95 ± 12.18 | 12 | 37.11 ± 15.84 | 8 | 1.93 [-1.97, 5.78] | 2.36 [-1.85, 6.66] | -1.81 [-7.93, 4.47] | | | | | | |
| L 1096 | L 127 | L Anteromedial nucleus, dorsal part | 14.59 ± 6.50 | 13 | 52.74 ± 48.72 | 9 | 28.19 ± 11.40 | 12 | 28.46 ± 10.75 | 8 | 0.17 [-3.67, 4.02] | 2.28 [-1.75, 6.33] | -1.95 [-8.08, 4.07] | | | | | | |
| R 1104 | R 127 | R Anteromedial nucleus, ventral part | 10.44 ± 3.94 | 13 | 34.16 ± 32.01 | 8 | 45.82 ± 23.27 | 12 | 37.44 ± 13.30 | 8 | 2.62 [-1.25, 6.44] | 2.26 [-1.99, 6.43] | -2.72 [-8.81, 3.39] | | | | | | |
| L 1104 | L 127 | L Anteromedial nucleus, ventral part | 5.50 ± 2.81 | 13 | 51.87 ± 42.97 | 9 | 20.30 ± 10.20 | 12 | 21.32 ± 10.55 | 7 | 1.93 [-1.95, 5.78] | 8.09 [4.05, 12.04]* | -7.1 [-13.17, -0.9]* | -8.02 [-11.86, -3.92]* | -2.77 [-6.46, 1.11] | -5.15 [-9.39, -0.63]* | -5.31 [-9.31, -0.96]* | 2.93 [-1.72, 7.56] | -2.4 [-6.73, 2.16] |
| R 64 | R 239 | R Anterodorsal nucleus | 119.85 ± 60.23 | 12 | 176.82 ± 109.31 | 9 | 202.86 ± 94.83 | 11 | 161.16 ± 49.59 | 8 | -0.01 [-4.08, 4.02] | 0.2 [-4.01, 4.36] | -0.23 [-6.48, 6.04] | | | | | | |
| L 64 | L 239 | L Anterodorsal nucleus | 81.92 ± 36.74 | 11 | 100.42 ± 77.65 | 9 | 127.10 ± 64.34 | 10 | 110.63 ± 66.99 | 7 | -0.01 [-4.16, 4.2] | 0 [-4.33, 4.21] | 0.11 [-6.32, 6.44] | | | | | | |
| R 1120 | R 239 | R Interanteromedial nucleus of the thalamus | 3.17 ± 3.17 | 12 | 32.88 ± 27.73 | 8 | 8.08 ± 3.75 | 11 | 40.26 ± 21.36 | 7 | 0.63 [-3.35, 4.7] | 8.89 [4.61, 13.14]* | 1.79 [-4.54, 8.26] | -8.83 [-13.05, -4.65]* | -1.45 [-5.32, 2.65] | -13.53 [-18.04, -9.21]* | -7.4 [-11.48, -2.89]* | -4.7 [-9.34, 0.17] | -12.06 [-16.7, -7.59]* |
| L 1120 | L 239 | L Interanteromedial nucleus of the thalamus | 6.10 ± 3.21 | 11 | 54.43 ± 51.37 | 8 | 12.43 ± 4.82 | 11 | 39.37 ± 15.72 | 7 | 0.18 [-3.83, 4.21] | 7.49 [3.15, 11.83]* | -2.55 [-8.85, 3.8] | -7.44 [-11.57, -2.98]* | -1.01 [-4.86, 3.1] | -7.38 [-11.76, -2.75]* | -6.4 [-10.8, -2.25]* | 0.08 [-4.8, 4.8] | -6.35 [-11, -1.91]* |
| R 1113 | R 239 | R Interanterodorsal nucleus of the thalamus | 14.57 ± 7.21 | 12 | 45.64 ± 40.34 | 9 | 49.80 ± 15.76 | 12 | 71.45 ± 24.84 | 8 | 1.71 [-2.17, 5.66] | 1.85 [-2.3, 5.97] | -0.04 [-6.13, 5.96] | | | | | | |
| L 1113 | L 239 | L Interanterodorsal nucleus of the thalamus | 5.32 ± 2.42 | 12 | 52.34 ± 49.21 | 9 | 28.93 ± 11.22 | 12 | 35.77 ± 14.05 | 7 | 3.73 [-0.22, 7.68] | 8.57 [4.41, 12.69]* | -6.49 [-12.73, -0.2]* | -8.51 [-12.66, -4.53]* | -4.55 [-8.41, -0.57]* | -8.05 [-12.27, -3.44]* | -3.98 [-8.04, 0.19] | 0.47 [-4.14, 5.04] | -3.48 [-7.84, 1.08] |
| R 155 | R 239 | R Lateral dorsal nucleus of thalamus | 244.15 ± 128.66 | 15 | 148.97 ± 51.10 | 13 | 253.47 ± 43.50 | 12 | 424.32 ± 152.63 | 10 | -0.57 [-4.28, 3.14] | -0.42 [-4.01, 3.21] | -0.01 [-5.57, 5.55] | | | | | | |
| L 155 | L 239 | L Lateral dorsal nucleus of thalamus | 24.13 ± 8.21 | 15 | 48.65 ± 18.96 | 13 | 130.27 ± 33.97 | 12 | 151.08 ± 32.49 | 10 | 3.78 [-0.02, 7.52] | 1 [-2.62, 4.7] | -1.28 [-6.9, 4.46] | | | | | | |
| R 444 | R 856 | R Medial group of the dorsal thalamus | 11.02 ± 5.45 | 15 | 14.62 ± 7.00 | 14 | 32.40 ± 9.11 | 12 | 45.19 ± 13.31 | 10 | 1.37 [-2.37, 5.15] | 0.33 [-3.19, 3.93] | -0.35 [-5.85, 5.27] | | | | | | |
| L 444 | L 856 | L Medial group of the dorsal thalamus | 11.57 ± 5.44 | 15 | 8.70 ± 4.65 | 14 | 26.24 ± 6.51 | 12 | 52.67 ± 12.71 | 10 | 0.7 [-3.1, 4.38] | -0.24 [-3.8, 3.34] | 1.37 [-4.16, 6.91] | | | | | | |
| R 59 | R 444 | R Intermediodorsal nucleus of the thalamus | 13.77 ± 10.14 | 14 | 12.77 ± 6.75 | 14 | 34.49 ± 12.01 | 12 | 68.80 ± 37.54 | 10 | 0.99 [-2.82, 4.86] | -0.01 [-3.73, 3.7] | 1.32 [-4.29, 6.95] | | | | | | |
| L 59 | L 444 | L Intermediodorsal nucleus of the thalamus | 14.24 ± 4.84 | 14 | 17.17 ± 9.43 | 14 | 40.24 ± 20.12 | 12 | 75.28 ± 44.24 | 10 | 1.3 [-2.51, 5.1] | 0.28 [-3.41, 3.91] | 1.04 [-4.42, 6.59] | | | | | | |
| R 362 | R 444 | R Mediodorsal nucleus of thalamus | 17.50 ± 7.24 | 15 | 37.72 ± 17.02 | 14 | 68.41 ± 23.15 | 12 | 88.23 ± 31.48 | 10 | 2.3 [-1.47, 6.11] | 1.15 [-2.43, 4.74] | -1.16 [-6.81, 4.38] | | | | | | |
| L 362 | L 444 | L Mediodorsal nucleus of thalamus | 26.59 ± 12.87 | 15 | 20.76 ± 13.15 | 13 | 55.36 ± 12.84 | 12 | 103.54 ± 27.18 | 10 | 0.47 [-3.27, 4.23] | -0.24 [-3.87, 3.44] | 0.88 [-4.59, 6.51] | | | | | | |
| R 366 | R 444 | R Submedial nucleus of the thalamus | 24.80 ± 17.01 | 14 | 15.06 ± 8.70 | 13 | 70.19 ± 20.68 | 11 | 139.98 ± 36.44 | 9 | 1.35 [-2.57, 5.31] | -0.32 [-4.02, 3.34] | 2.1 [-3.64, 7.95] | | | | | | |
| L 366 | L 444 | L Submedial nucleus of the thalamus | 11.08 ± 4.89 | 14 | 10.99 ± 3.13 | 13 | 51.37 ± 14.37 | 11 | 159.86 ± 34.10 | 10 | 3.6 [-0.36, 7.51] | 0.07 [-3.6, 3.74] | 8.09 [2.37, 13.9]* | -0.03 [-3.54, 3.72] | -4.41 [-8.38, -0.53]* | -14.01 [-17.99, -10.02]* | 4.41 [0.54, 8.3]* | -14 [-17.77, -9.66]* | -9.56 [-13.9, -5.55]* |
| R 1077 | R 444 | R Perireunensis nucleus | 61.28 ± 32.99 | 15 | 65.79 ± 36.80 | 13 | 33.97 ± 11.08 | 12 | 39.51 ± 12.06 | 10 | -1.05 [-4.81, 2.74] | 0.06 [-3.59, 3.81] | -1.11 [-6.75, 4.57] | | | | | | |
| L 1077 | L 444 | L Perireunensis nucleus | 33.14 ± 16.12 | 15 | 24.84 ± 9.80 | 13 | 21.60 ± 10.12 | 12 | 68.25 ± 35.99 | 10 | -0.94 [-4.63, 2.82] | -0.24 [-3.97, 3.36] | 0.5 [-4.97, 5.98] | | | | | | |
| R 571 | R 856 | R Midline group of the dorsal thalamus | 29.85 ± 14.81 | 15 | 26.67 ± 10.35 | 15 | 30.75 ± 7.40 | 12 | 31.47 ± 8.98 | 10 | -0.59 [-4.32, 3.24] | -0.09 [-3.63, 3.51] | -1.06 [-6.63, 4.39] | | | | | | |
| L 571 | L 856 | L Midline group of the dorsal thalamus | 23.42 ± 14.89 | 15 | 20.03 ± 8.12 | 15 | 16.76 ± 4.35 | 12 | 27.25 ± 7.77 | 10 | | | | | | | | | |
| R 149 | R 571 | R Paraventricular nucleus of the thalamus | 92.30 ± 54.15 | 15 | 81.41 ± 36.71 | 15 | 63.02 ± 27.99 | 12 | 59.50 ± 27.61 | 10 | -0.9 [-4.75, 3.01] | -0.06 [-3.6, 3.51] | -1.14 [-6.77, 4.31] | | | | | | |
| L 149 | L 571 | L Paraventricular nucleus of the thalamus | 95.98 ± 67.14 | 15 | 50.63 ± 21.28 | 15 | 28.83 ± 13.40 | 12 | 75.55 ± 27.74 | 10 | -1.29 [-5.05, 2.5] | -0.42 [-3.99, 3.08] | -0.23 [-5.78, 5.34] | | | | | | |
| R 181 | R 571 | R Nucleus of reuniens | 44.35 ± 17.17 | 15 | 36.03 ± 14.26 | 13 | 52.47 ± 16.04 | 12 | 77.39 ± 33.57 | 10 | -0.43 [-4.19, 3.38] | -0.2 [-3.84, 3.44] | -0.39 [-6.04, 5.27] | | | | | | |
| L 181 | L 571 | L Nucleus of reuniens | 25.61 ± 11.13 | 15 | 33.55 ± 14.45 | 13 | 34.01 ± 9.60 | 12 | 47.80 ± 11.97 | 10 | -0.28 [-4.07, 3.51] | 0.32 [-3.38, 3.89] | -0.92 [-6.44, 4.74] | | | | | | |
| R 560581559 | R 571 | R Xiphoid thalamic nucleus | 47.39 ± 15.76 | 15 | 59.45 ± 22.53 | 13 | 74.23 ± 24.04 | 12 | 103.66 ± 26.07 | 10 | -0.04 [-3.8, 3.75] | 0.24 [-3.46, 3.85] | -0.77 [-6.42, 4.94] | | | | | | |
| L 560581559 | L 571 | L Xiphoid thalamic nucleus | 22.47 ± 10.35 | 15 | 48.35 ± 20.48 | 13 | 40.36 ± 25.74 | 12 | 43.45 ± 12.85 | 10 | 0.18 [-3.56, 3.92] | 1.13 [-2.46, 4.75] | -2.16 [-7.66, 3.45] | | | | | | |
| R 51 | R 856 | R Intralaminar nuclei of the dorsal thalamus | 8.53 ± 3.34 | 15 | 12.83 ± 5.35 | 15 | 35.21 ± 5.39 | 12 | 64.34 ± 12.04 | 10 | 2.56 [-1.25, 6.36] | 0.55 [-3, 4.12] | 1.67 [-3.92, 7.27] | | | | | | |
| L 51 | L 856 | L Intralaminar nuclei of the dorsal thalamus | 6.70 ± 2.76 | 15 | 6.42 ± 1.77 | 15 | 32.21 ± 5.03 | 12 | 56.75 ± 15.41 | 10 | 3.22 [-0.52, 7.07] | 0.02 [-3.5, 3.61] | 2.47 [-3.06, 7.89] | | | | | | |
| R 189 | R 51 | R Rhomboid nucleus | 9.13 ± 3.94 | 15 | 30.63 ± 23.71 | 13 | 66.87 ± 19.53 | 12 | 154.40 ± 54.69 | 10 | 5.74 [1.96, 9.54]* | 2.35 [-1.29, 6.02] | 6.07 [0.4, 11.66]* | -2.3 [-5.92, 1.31] | -6.55 [-10.26, -2.76]* | -16.42 [-20.08, -12.11]* | 4.24 [0.51, 8.08]* | -14.08 [-18.22, -10.11]* | -9.87 [-14.02, -5.74]* |
| L 189 | L 51 | L Rhomboid nucleus | 14.29 ± 5.63 | 15 | 15.07 ± 7.73 | 13 | 83.37 ± 42.10 | 12 | 145.38 ± 46.47 | 10 | 4.26 [0.55, 7.98]* | 0.03 [-3.59, 3.59] | 3.14 [-2.37, 8.72] | 0.03 [-3.39, 3.7] | -5.07 [-8.77, -1.41]* | -9.69 [-13.59, -5.71]* | 5.08 [1.31, 8.94]* | -9.68 [-13.78, -5.76]* | -4.59 [-8.68, -0.43]* |
| R 599 | R 51 | R Central medial nucleus of the thalamus | 5.54 ± 1.87 | 15 | 23.87 ± 10.81 | 14 | 41.27 ± 11.24 | 12 | 78.72 ± 30.24 | 10 | 5.85 [2.12, 9.67]* | 3.3 [-0.31, 6.96] | 2.29 [-3.38, 7.88] | -3.25 [-6.74, 0.29] | -6.69 [-10.36, -2.99]* | -13.67 [-17.6, -9.73]* | 3.42 [-0.34, 7.04] | -10.44 [-14.26, -6.24]* | -7.03 [-11.16, -2.99]* |
| L 599 | L 51 | L Central medial nucleus of the thalamus | 7.23 ± 3.19 | 15 | 21.54 ± 12.23 | 14 | 33.04 ± 10.89 | 12 | 76.12 ± 26.92 | 10 | 2.99 [-0.82, 6.64] | 2.02 [-1.55, 5.54] | 2.78 [-2.63, 8.25] | | | | | | |
| R 907 | R 51 | R Paracentral nucleus | 6.53 ± 2.92 | 14 | 16.57 ± 9.33 | 13 | 87.34 ± 20.90 | 11 | 95.51 ± 15.27 | 10 | 11.93 [8.09, 15.85]* | 1.63 [-2.07, 5.33] | -1.62 [-7.47, 4.06] | -1.59 [-5.22, 2.06] | -12.76 [-16.65, -9.03]* | -14.17 [-18.07, -10.06]* | 11.16 [7.57, 15.15]* | -12.61 [-16.51, -8.4]* | -1.41 [-5.63, 2.83] |
| L 907 | L 51 | L Paracentral nucleus | 5.75 ± 2.52 | 14 | 3.91 ± 2.03 | 13 | 63.98 ± 20.37 | 12 | 118.79 ± 36.69 | 10 | 9.61 [5.85, 13.42]* | -0.22 [-3.88, 3.41] | 8.6 [2.87, 14.21]* | 0.27 [-3.36, 3.79] | -10.46 [-14.28, -6.76]* | -20.23 [-24.21, -16.22]* | 10.71 [6.9, 14.44]* | -20.51 [-24.5, -16.52]* | -9.8 [-14, -5.64]* |
| R 575 | R 51 | R Central lateral nucleus of the thalamus | 42.14 ± 22.93 | 14 | 53.59 ± 20.49 | 14 | 46.98 ± 12.85 | 12 | 144.27 ± 27.95 | 10 | -0.42 [-4.18, 3.41] | 0.39 [-3.17, 4.06] | 0.75 [-4.84, 6.27] | | | | | | |
| L 575 | L 51 | L Central lateral nucleus of the thalamus | 30.41 ± 16.11 | 14 | 20.61 ± 7.83 | 14 | 48.24 ± 8.63 | 12 | 100.20 ± 33.71 | 10 | 0.07 [-3.72, 3.92] | -0.2 [-3.81, 3.52] | 0.76 [-4.82, 6.35] | | | | | | |
| R 930 | R 51 | R Parafascicular nucleus | 9.96 ± 3.89 | 11 | 13.73 ± 8.31 | 10 | 90.52 ± 13.59 | 10 | 186.55 ± 35.66 | 7 | 8.13 [3.97, 12.31]* | 0.85 [-3.26, 4.84] | 8.04 [1.75, 14.34]* | -0.81 [-4.84, 3.15] | -8.94 [-13.06, -4.85]* | -19.27 [-23.71, -14.53]* | 8.13 [3.89, 12.33]* | -18.46 [-23.06, -13.83]* | -10.3 [-15.1, -5.79]* |
| L 930 | L 51 | L Parafascicular nucleus | 11.84 ± 5.51 | 10 | 3.78 ± 1.21 | 10 | 88.11 ± 14.82 | 8 | 116.19 ± 30.95 | 7 | 6.6 [2.16, 11]* | -0.91 [-5.13, 3.29] | 2.44 [-4.01, 8.94] | 0.94 [-3.28, 5.01] | -7.46 [-11.89, -3.18]* | -10.37 [-14.88, -5.89]* | 8.39 [4, 12.74]* | -11.38 [-15.81, -6.48]* | -2.95 [-7.66, 1.94] |

| | | | | | | | | | | | | | | | | | | |
|---|---|---|---|---|---|---|---|---|---|---|---|---|---|---|---|---|---|---|
| R 262 | R 856 | R Reticular nucleus of the thalamus | 147.28 ± 98.89 | 15 | 150.34 ± 69.44 | 14 | 67.19 ± 9.27 | 12 | 88.68 ± 12.28 | 10 | -1.14 [-4.95, 2.63] | 0.03 [-3.51, 3.65] | -1.03 [-6.64, 4.55] | | | | | |
| L 262 | L 856 | L Reticular nucleus of the thalamus | 38.38 ± 20.08 | 15 | 20.33 ± 9.57 | 14 | 35.30 ± 9.20 | 12 | 54.39 ± 11.66 | 10 | -0.67 [-4.46, 3.11] | -0.48 [-4.07, 3.12] | -0.19 [-5.79, 5.51] | | | | | |
| R 1014 | R 856 | R Geniculate group, ventral thalamus | 204.26 ± 149.45 | 13 | 131.01 ± 85.05 | 15 | 131.50 ± 42.87 | 12 | 147.24 ± 56.70 | 9 | -0.83 [-4.66, 3.09] | -0.17 [-3.82, 3.43] | 0.09 [-5.72, 5.79] | | | | | |
| L 1014 | L 856 | L Geniculate group, ventral thalamus | 24.28 ± 10.71 | 14 | 61.35 ± 45.68 | 15 | 37.65 ± 9.63 | 12 | 97.66 ± 51.54 | 10 | -0.01 [-3.83, 3.83] | 1.65 [-2.05, 5.13] | -0.32 [-5.84, 5.34] | | | | | |
| R 27 | R 1014 | R Intergeniculate leaflet of the lateral geniculate complex | 699.24 ± 651.07 | 9 | 337.85 ± 170.18 | 12 | 152.71 ± 67.90 | 10 | 634.83 ± 297.17 | 8 | -1.16 [-5.55, 3.19] | -0.49 [-4.68, 3.68] | 1.12 [-5.14, 7.3] | | | | | |
| L 27 | L 1014 | L Intergeniculate leaflet of the lateral geniculate complex | 162.58 ± 132.15 | 12 | 37.19 ± 24.92 | 10 | 87.57 ± 46.12 | 10 | 66.59 ± 19.88 | 8 | -0.96 [-5.07, 3.25] | 0.06 [-4.02, 4.08] | 0.16 [-6, 6.37] | | | | | |
| R 178 | R 1014 | R Ventral part of the lateral geniculate complex | 422.46 ± 314.26 | 13 | 247.26 ± 178.69 | 15 | 264.47 ± 87.31 | 12 | 272.06 ± 109.51 | 9 | -0.84 [-4.69, 3.07] | -0.25 [-3.81, 3.43] | 0.11 [-5.44, 5.76] | | | | | |
| L 178 | L 1014 | L Ventral part of the lateral geniculate complex | 48.55 ± 23.98 | 14 | 122.35 ± 91.96 | 15 | 78.23 ± 19.60 | 12 | 202.64 ± 102.79 | 10 | 0.07 [-3.75, 3.83] | 1.63 [-1.94, 5.24] | -0.2 [-5.79, 5.4] | | | | | |
| L 321 | L 1014 | L Subgeniculate nucleus | 85.30 ± 55.75 | 8 | 16.92 ± 9.46 | 10 | 39.26 ± 24.73 | 9 | 61.06 ± 28.13 | 8 | -0.98 [-5.57, 3.58] | -0.48 [-4.83, 3.86] | 0.85 [-5.41, 7.31] | | | | | |
| R 958 | R 856 | R Epithalamus | 49.64 ± 21.98 | 15 | 69.94 ± 33.29 | 15 | 46.02 ± 15.49 | 12 | 76.87 ± 16.73 | 10 | -0.68 [-4.31, 2.98] | 0.48 [-3.03, 4.07] | -1 [-6.55, 4.59] | | | | | |
| L 958 | L 856 | L Epithalamus | 25.39 ± 6.95 | 15 | 17.04 ± 4.36 | 15 | 36.30 ± 15.33 | 12 | 67.75 ± 30.83 | 10 | -0.16 [-3.91, 3.65] | -0.28 [-3.78, 3.27] | 0.36 [-5.17, 5.83] | | | | | |
| R 483 | R 958 | R Medial habenula | 92.77 ± 51.22 | 15 | 129.91 ± 50.93 | 15 | 113.12 ± 37.60 | 12 | 170.17 ± 41.54 | 10 | -0.36 [-4.14, 3.42] | 0.45 [-3.03, 4.06] | -1.01 [-6.5, 4.44] | | | | | |
| L 483 | L 958 | L Medial habenula | 50.41 ± 13.56 | 15 | 36.77 ± 10.69 | 15 | 122.13 ± 55.55 | 12 | 204.29 ± 98.23 | 10 | 0.83 [-2.93, 4.59] | -0.25 [-3.73, 3.35] | 0.7 [-4.92, 6.22] | | | | | |
| R 186 | R 958 | R Lateral habenula | 99.49 ± 41.89 | 15 | 187.00 ± 133.69 | 14 | 68.20 ± 31.88 | 12 | 135.10 ± 34.78 | 10 | -0.91 [-4.65, 2.87] | 0.93 [-2.59, 4.55] | -1.42 [-7.03, 4.1] | | | | | |
| L 186 | L 958 | L Lateral habenula | 47.05 ± 18.22 | 15 | 26.02 ± 7.44 | 14 | 17.33 ± 4.61 | 12 | 70.46 ± 35.80 | 10 | -1.24 [-4.98, 2.5] | -0.39 [-3.97, 3.24] | 0.38 [-5.03, 5.89] | | | | | |
| R 1097 | R 1129 | R Hypothalamus | 28.16 ± 12.18 | 14 | 25.05 ± 5.54 | 14 | 32.63 ± 8.05 | 12 | 44.52 ± 8.01 | 10 | | | | | | | | |
| L 1097 | L 1129 | L Hypothalamus | 25.06 ± 14.00 | 14 | 11.80 ± 2.92 | 13 | 18.49 ± 4.27 | 12 | 29.52 ± 4.80 | 10 | -0.77 [-2.52, 1.05] | -0.32 [-2.13, 1.42] | 0.05 [-2.58, 2.72] | | | | | |
| R 157 | R 1097 | R Periventricular zone | 72.81 ± 35.11 | 14 | 27.51 ± 8.36 | 14 | 27.19 ± 13.82 | 12 | 76.25 ± 22.27 | 10 | -1.13 [-2.88, 0.64] | -0.48 [-2.21, 1.21] | 0.44 [-2.22, 3.06] | | | | | |
| L 157 | L 1097 | L Periventricular zone | 66.69 ± 39.27 | 15 | 25.96 ± 7.32 | 14 | 19.00 ± 6.70 | 12 | 55.92 ± 17.27 | 10 | -1.25 [-3.03, 0.52] | -0.5 [-2.25, 1.24] | 0.35 [-2.29, 3] | | | | | |
| R 390 | R 157 | R Supraoptic nucleus | 47.81 ± 16.96 | 10 | 35.43 ± 21.53 | 7 | 97.29 ± 36.14 | 11 | 44.67 ± 33.88 | 6 | 0.11 [-1.81, 2.02] | -0.33 [-2.45, 1.75] | -1.49 [-4.61, 1.64] | | | | | |
| R 38 | R 157 | R Paraventricular hypothalamic nucleus | 35.09 ± 15.33 | 13 | 30.70 ± 15.32 | 8 | 23.50 ± 5.32 | 12 | 44.11 ± 12.91 | 8 | -0.81 [-2.61, 0.99] | 0.31 [-1.66, 2.29] | -0.18 [-3.08, 2.66] | | | | | |
| L 38 | L 157 | L Paraventricular hypothalamic nucleus | 41.95 ± 17.23 | 13 | 25.10 ± 18.09 | 8 | 18.40 ± 5.21 | 12 | 46.33 ± 13.36 | 8 | -1.04 [-2.84, 0.79] | 0.04 [-1.93, 2.04] | 0.15 [-2.71, 2.99] | | | | | |
| R 118 | R 157 | R Periventricular hypothalamic nucleus, intermediate part | 38.99 ± 21.20 | 13 | 22.20 ± 8.93 | 14 | 10.03 ± 2.78 | 12 | 41.25 ± 18.36 | 10 | -1.18 [-2.97, 0.65] | -0.22 [-1.99, 1.53] | 0.31 [-2.33, 2.96] | | | | | |
| L 118 | L 157 | L Periventricular hypothalamic nucleus, intermediate part | 43.35 ± 23.12 | 13 | 20.81 ± 8.37 | 14 | 7.00 ± 2.03 | 12 | 32.33 ± 10.36 | 10 | -1.26 [-3.07, 0.58] | -0.31 [-2.06, 1.46] | 0.17 [-2.5, 2.8] | | | | | |
| R 223 | R 157 | R Arcuate hypothalamic nucleus | 302.37 ± 132.61 | 13 | 85.80 ± 26.59 | 13 | 251.83 ± 207.61 | 11 | 279.64 ± 84.26 | 9 | -0.57 [-2.44, 1.25] | -0.46 [-2.23, 1.31] | -0.21 [-2.87, 2.51] | | | | | |
| L 223 | L 157 | L Arcuate hypothalamic nucleus | 318.68 ± 171.74 | 13 | 92.06 ± 29.42 | 14 | 189.43 ± 124.48 | 10 | 193.24 ± 56.31 | 9 | -0.84 [-2.71, 1.03] | -0.52 [-2.3, 1.25] | -0.22 [-2.98, 2.54] | | | | | |
| R 141 | R 1097 | R Periventricular region | 63.80 ± 32.39 | 15 | 44.58 ± 24.72 | 15 | 23.24 ± 6.33 | 12 | 40.48 ± 12.00 | 10 | -1.16 [-2.87, 0.6] | -0.21 [-1.89, 1.5] | -0.25 [-2.85, 2.36] | | | | | |
| L 141 | L 1097 | L Periventricular region | 86.06 ± 40.49 | 14 | 30.77 ± 14.28 | 15 | 15.53 ± 6.08 | 12 | 35.96 ± 8.50 | 10 | -1.32 [-3.1, 0.43] | -0.53 [-2.26, 1.14] | 0.05 [-2.55, 2.7] | | | | | |
| R 830 | R 141 | R Dorsomedial nucleus of the hypothalamus | 98.32 ± 60.62 | 12 | 57.27 ± 21.03 | 13 | 23.46 ± 4.79 | 11 | 41.48 ± 13.08 | 9 | -1.12 [-2.97, 0.77] | -0.21 [-2.03, 1.64] | -0.38 [-3.15, 2.4] | | | | | |
| L 830 | L 141 | L Dorsomedial nucleus of the hypothalamus | 70.98 ± 53.60 | 12 | 37.20 ± 20.76 | 13 | 14.88 ± 4.13 | 10 | 30.42 ± 11.27 | 9 | -1.15 [-3.06, 0.76] | -0.27 [-2.09, 1.56] | -0.29 [-3.09, 2.51] | | | | | |
| R 126 | R 141 | R Periventricular hypothalamic nucleus, posterior part | 312.33 ± 199.14 | 9 | 148.59 ± 112.61 | 9 | 97.40 ± 49.60 | 6 | 147.55 ± 75.43 | 8 | -1.22 [-3.48, 1] | -0.08 [-2.14, 1.97] | -0.12 [-3.23, 3.03] | | | | | |
| L 126 | L 141 | L Periventricular hypothalamic nucleus, posterior part | 378.70 ± 196.82 | 9 | 152.59 ± 98.81 | 9 | 115.81 ± 59.39 | 6 | 131.20 ± 44.61 | 8 | -1.25 [-3.5, 1.01] | -0.16 [-2.23, 1.86] | -0.14 [-3.25, 3.03] | | | | | |
| R 347 | R 141 | R Subparaventricular zone | 33.91 ± 23.07 | 13 | 26.96 ± 14.59 | 8 | 10.00 ± 3.96 | 12 | 28.46 ± 12.82 | 8 | -1.16 [-2.95, 0.66] | 0.25 [-1.7, 2.25] | -0.18 [-3.05, 2.68] | | | | | |
| L 347 | L 141 | L Subparaventricular zone | 28.36 ± 21.05 | 13 | 20.04 ± 8.81 | 8 | 18.88 ± 6.59 | 11 | 39.34 ± 21.22 | 7 | -0.84 [-2.67, 0.99] | 0.17 [-1.81, 2.14] | 0.36 [-2.59, 3.36] | | | | | |
| R 467 | R 1097 | R Hypothalamic medial zone | 40.12 ± 18.14 | 15 | 36.63 ± 9.43 | 15 | 50.24 ± 17.65 | 12 | 50.79 ± 8.62 | 10 | -0.26 [-2.02, 1.56] | 0.02 [-1.67, 1.73] | -0.73 [-3.37, 1.9] | | | | | |
| L 467 | L 1097 | L Hypothalamic medial zone | 35.50 ± 22.22 | 15 | 20.25 ± 5.68 | 15 | 23.48 ± 8.93 | 12 | 34.73 ± 6.82 | 10 | -0.87 [-2.63, 0.91] | -0.35 [-2.06, 1.35] | -0.04 [-2.69, 2.58] | | | | | |
| R 88 | R 467 | R Anterior hypothalamic nucleus | 32.01 ± 13.11 | 15 | 27.06 ± 8.29 | 11 | 46.07 ± 10.36 | 12 | 46.34 ± 17.25 | 8 | -0.08 [-1.82, 1.72] | 0.02 [-1.8, 1.86] | -0.48 [-3.22, 2.28] | | | | | |
| L 88 | L 467 | L Anterior hypothalamic nucleus | 16.06 ± 6.92 | 14 | 28.22 ± 13.25 | 11 | 20.19 ± 4.94 | 12 | 25.68 ± 7.62 | 9 | -0.21 [-1.99, 1.6] | 0.89 [-0.92, 2.76] | -1.19 [-3.94, 1.53] | | | | | |
| R 331 | R 467 | R Mammillary body | 72.82 ± 26.42 | 9 | 73.02 ± 18.50 | 12 | 101.54 ± 52.67 | 10 | 107.91 ± 27.22 | 7 | 0.16 [-1.86, 2.16] | 0.27 [-1.68, 2.24] | -0.69 [-3.65, 2.28] | | | | | |
| L 331 | L 467 | L Mammillary body | 61.08 ± 36.25 | 10 | 22.45 ± 5.47 | 11 | 53.35 ± 28.51 | 10 | 79.23 ± 23.24 | 8 | -0.37 [-2.33, 1.61] | -0.22 [-2.17, 1.69] | -0.04 [-2.97, 2.85] | | | | | |
| R 210 | R 331 | R Lateral mammillary nucleus | 108.63 ± 23.21 | 6 | 69.04 ± 40.90 | 7 | 46.56 ± 27.81 | 7 | 116.18 ± 20.62 | 7 | -0.6 [-2.93, 1.77] | 0.43 [-1.96, 2.8] | -0.12 [-3.46, 3.21] | | | | | |
| L 210 | L 331 | L Lateral mammillary nucleus | 175.64 ± 131.22 | 7 | 32.86 ± 17.39 | 8 | 65.32 ± 49.93 | 7 | 58.14 ± 16.55 | 8 | -0.85 [-3.11, 1.41] | -0.26 [-2.46, 1.92] | -0.3 [-3.5, 2.84] | | | | | |
| R 491 | R 331 | R Medial mammillary nucleus | 57.56 ± 15.88 | 7 | 25.01 ± 10.45 | 9 | 105.50 ± 70.20 | 6 | 147.44 ± 30.48 | 7 | 0.55 [-1.85, 2.92] | 0.05 [-2.15, 2.19] | 0.41 [-2.84, 3.68] | | | | | |
| L 491 | L 331 | L Medial mammillary nucleus | 39.65 ± 16.02 | 7 | 12.27 ± 9.40 | 8 | 81.71 ± 46.24 | 6 | 100.44 ± 13.74 | 7 | 0.78 [-1.61, 3.16] | -0.13 [-2.34, 2.11] | 0.34 [-3, 3.69] | | | | | |
| R 732 | R 491 | R Medial mammillary nucleus, median part | 80.71 ± 41.99 | 7 | 67.75 ± 43.24 | 8 | 186.65 ± 131.35 | 6 | 169.94 ± 63.25 | 7 | 1.04 [-1.31, 3.41] | 0.42 [-1.86, 2.65] | -0.89 [-4.22, 2.47] | | | | | |
| L 732 | L 491 | L Medial mammillary nucleus, median part | 80.55 ± 42.06 | 7 | 7.02 ± 7.02 | 8 | 142.80 ± 86.57 | 6 | 100.13 ± 33.49 | 7 | 0.49 [-1.79, 2.81] | -0.34 [-2.56, 1.85] | -0.45 [-3.7, 2.81] | | | | | |
| R 606826647 | R 491 | R Medial mammillary nucleus, lateral part | 154.69 ± 40.16 | 7 | 77.25 ± 40.46 | 9 | 201.72 ± 127.50 | 6 | 257.59 ± 71.80 | 7 | 0.01 [-2.33, 2.37] | 0.11 [-2.05, 2.32] | -0.01 [-3.33, 3.21] | | | | | |
| L 606826647 | L 491 | L Medial mammillary nucleus, lateral part | 75.70 ± 30.84 | 7 | 18.15 ± 11.53 | 8 | 112.24 ± 63.78 | 6 | 177.88 ± 22.02 | 7 | 0.19 [-2.15, 2.54] | -0.19 [-2.43, 2.05] | 0.8 [-2.5, 4.15] | | | | | |
| R 606826651 | R 491 | R Medial mammillary nucleus, medial part | 94.27 ± 42.52 | 7 | 26.23 ± 21.29 | 9 | 215.63 ± 140.85 | 6 | 414.54 ± 102.92 | 6 | 1 [-1.33, 3.33] | -0.1 [-2.29, 2.04] | 2.2 [-1.1, 5.55] | | | | | |
| L 606826651 | L 491 | L Medial mammillary nucleus, medial part | 79.06 ± 36.07 | 7 | 47.19 ± 36.38 | 8 | 219.77 ± 131.63 | 6 | 309.25 ± 61.29 | 6 | 1.5 [-0.84, 3.81] | 0.17 [-2.02, 2.35] | 0.96 [-2.39, 4.25] | | | | | |
| R 525 | R 331 | R Supramammillary nucleus | 428.91 ± 196.29 | 8 | 406.35 ± 130.90 | 9 | 807.88 ± 424.70 | 8 | 588.50 ± 212.44 | 7 | 0.64 [-1.53, 2.77] | 0.44 [-1.7, 2.58] | -1.37 [-4.45, 1.77] | | | | | |
| L 525 | L 331 | L Supramammillary nucleus | 397.32 ± 248.17 | 9 | 133.36 ± 35.24 | 9 | 386.22 ± 229.90 | 9 | 344.77 ± 184.63 | 8 | -0.32 [-2.34, 1.72] | -0.22 [-2.23, 1.84] | -0.5 [-3.44, 2.5] | | | | | |
| R 557 | R 331 | R Tuberomammillary nucleus | 35.13 ± 24.15 | 8 | 68.58 ± 25.03 | 11 | 18.54 ± 6.66 | 8 | 13.85 ± 2.31 | 6 | -0.77 [-2.9, 1.41] | 1.28 [-0.77, 3.28] | -1.7 [-4.79, 1.4] | | | | | |
| L 557 | L 331 | L Tuberomammillary nucleus | 48.60 ± 40.88 | 10 | 44.08 ± 22.29 | 9 | 7.85 ± 3.50 | 8 | 17.46 ± 8.24 | 7 | -1.18 [-3.23, 0.89] | 0.31 [-1.7, 2.3] | -0.64 [-3.69, 2.42] | | | | | |
| R 1 | R 557 | R Tuberomammillary nucleus, ventral part | 28.87 ± 14.55 | 7 | 134.14 ± 62.53 | 9 | 40.66 ± 15.37 | 8 | 32.81 ± 5.69 | 6 | -0.43 [-2.65, 1.78] | 3.45 [1.25, 5.62]* | -3.98 [-7.19, -0.74]* | -3.35 [-5.45, -1.3]* | -0.17 [-2.25, 2] | -0.44 [-2.75, 1.86] | -3.18 [-5.21, -1.25]* | 2.92 [0.66, 5.07]* | -0.27 [-2.5, 1.96] |
| L 1 | L 557 | L Tuberomammillary nucleus, ventral part | 103.48 ± 84.74 | 10 | 25.98 ± 19.07 | 7 | 12.90 ± 4.04 | 8 | 19.98 ± 16.55 | 7 | -1.2 [-3.28, 0.82] | -0.21 [-2.37, 1.91] | -0.25 [-3.31, 2.87] | | | | | |
| L 980 | L 467 | L Dorsal premammillary nucleus | 213.87 ± 180.59 | 10 | 61.05 ± 34.13 | 8 | 23.91 ± 12.21 | 7 | 70.39 ± 15.60 | 8 | -1.33 [-3.45, 0.81] | -0.23 [-2.27, 1.86] | -0.01 [-3.09, 3.08] | | | | | |
| R 1004 | R 467 | R Ventral premammillary nucleus | 266.08 ± 241.83 | 8 | 120.94 ± 50.97 | 10 | 40.36 ± 13.60 | 8 | 76.88 ± 33.43 | 7 | -1.15 [-3.29, 1.03] | -0.27 [-2.35, 1.81] | -0.09 [-3.17, 2.98] | | | | | |
| L 1004 | L 467 | L Ventral premammillary nucleus | 245.89 ± 184.72 | 10 | 68.42 ± 26.38 | 9 | 23.91 ± 9.66 | 9 | 74.02 ± 20.23 | 8 | -1.17 [-3.18, 0.82] | -0.31 [-2.3, 1.74] | -0.14 [-3.15, 2.83] | | | | | |
| R 63 | R 467 | R Paraventricular hypothalamic nucleus, descending division | 62.09 ± 32.35 | 13 | 55.39 ± 26.26 | 11 | 18.38 ± 8.33 | 10 | 48.67 ± 17.75 | 9 | -1.17 [-3.01, 0.71] | 0.14 [-1.71, 1.98] | -0.27 [-3.03, 2.48] | | | | | |
| L 63 | L 467 | L Paraventricular hypothalamic nucleus, descending division | 33.70 ± 21.58 | 12 | 16.24 ± 8.64 | 11 | 28.75 ± 14.78 | 10 | 21.69 ± 12.73 | 8 | -0.72 [-2.59, 1.18] | -0.37 [-2.24, 1.53] | -0.35 [-3.24, 2.52] | | | | | |
| R 693 | R 467 | R Ventromedial hypothalamic nucleus | 73.58 ± 35.94 | 14 | 26.33 ± 13.13 | 12 | 69.23 ± 42.28 | 11 | 55.33 ± 11.99 | 9 | -0.55 [-2.37, 1.29] | -0.39 [-2.2, 1.4] | -0.51 [-3.25, 2.24] | | | | | |
| L 693 | L 467 | L Ventromedial hypothalamic nucleus | 64.66 ± 32.39 | 14 | 31.33 ± 15.55 | 12 | 33.32 ± 23.51 | 10 | 21.61 ± 5.55 | 9 | -0.93 [-2.76, 0.97] | -0.26 [-2.03, 1.52] | -0.68 [-3.46, 2.09] | | | | | |
| R 946 | R 467 | R Posterior hypothalamic nucleus | 126.67 ± 72.67 | 14 | 106.33 ± 57.44 | 14 | 97.26 ± 32.76 | 10 | 92.86 ± 29.33 | 8 | -0.68 [-2.54, 1.16] | -0.06 [-1.76, 1.65] | -0.65 [-3.43, 2.13] | | | | | |
| L 946 | L 467 | L Posterior hypothalamic nucleus | 109.07 ± 75.76 | 14 | 51.35 ± 22.42 | 14 | 42.09 ± 11.66 | 10 | 52.51 ± 9.50 | 8 | -1.04 [-2.93, 0.82] | -0.42 [-2.15, 1.32] | -0.17 [-2.91, 2.57] | | | | | |
| R 290 | R 1097 | R Hypothalamic lateral zone | 23.87 ± 8.50 | 15 | 23.20 ± 5.92 | 15 | 44.06 ± 8.71 | 12 | 61.49 ± 12.83 | 10 | 0.32 [-1.46, 2.07] | 0.06 [-1.64, 1.75] | -0.05 [-2.68, 2.54] | | | | | |
| L 290 | L 1097 | L Hypothalamic lateral zone | 16.48 ± 8.91 | 15 | 11.98 ± 2.79 | 15 | 28.02 ± 4.98 | 12 | 40.68 ± 7.00 | 10 | 0.17 [-1.63, 1.94] | -0.19 [-1.91, 1.49] | 0.25 [-2.38, 2.88] | | | | | |
| R 194 | R 290 | R Lateral hypothalamic area | 50.26 ± 16.54 | 15 | 69.22 ± 13.30 | 15 | 93.28 ± 21.54 | 10 | 105.23 ± 24.49 | 10 | 0.28 [-1.52, 2.11] | 0.47 [-1.23, 2.19] | -0.88 [-3.55, 1.79] | | | | | |
| L 194 | L 290 | L Lateral hypothalamic area | 37.01 ± 21.70 | 15 | 40.30 ± 9.14 | 15 | 44.74 ± 11.43 | 11 | 61.46 ± 17.65 | 10 | -0.39 [-2.18, 1.41] | 0.16 [-1.53, 1.84] | -0.36 [-2.96, 2.31] | | | | | |
| R 364 | R 290 | R Parasubthalamic nucleus | 58.49 ± 36.83 | 12 | 34.61 ± 11.23 | 12 | 54.60 ± 16.58 | 10 | 110.18 ± 46.03 | 8 | -0.42 [-2.32, 1.49] | -0.22 [-2.05, 1.62] | 0.48 [-2.39, 3.32] | | | | | |
| L 364 | L 290 | L Parasubthalamic nucleus | 66.15 ± 49.11 | 12 | 15.26 ± 4.77 | 12 | 37.78 ± 14.03 | 10 | 51.80 ± 21.80 | 8 | -0.75 [-2.66, 1.14] | -0.38 [-2.22, 1.43] | -0.08 [-2.89, 2.73] | | | | | |
| R 576073704 | R 290 | R Perifornical nucleus | 35.07 ± 10.31 | 15 | 41.01 ± 12.34 | 13 | 125.40 ± 37.00 | 12 | 75.59 ± 21.68 | 9 | 2.06 [0.31, 3.8]* | 0.33 [-1.43, 2.08] | -2.43 [-5.11, 0.23] | -0.25 [-1.89, 1.35] | -2.67 [-4.32, -1.01]* | -1.37 [-3.15, 0.49] | 2.42 [0.71, 4.12]* | -1.12 [-3, 0.73] | 1.32 [-0.59, 3.15] |
| L 576073704 | L 290 | L Perifornical nucleus | 22.84 ± 10.08 | 15 | 13.52 ± 5.12 | 13 | 41.68 ± 11.09 | 12 | 42.27 ± 10.53 | 8 | 0.3 [-1.46, 2] | -0.24 [-2.01, 1.5] | -0.31 [-2.97, 2.47] | | | | | |
| R 470 | R 290 | R Subthalamic nucleus | 3.38 ± 2.00 | 12 | 7.28 ± 2.38 | 12 | 51.64 ± 13.72 | 12 | 94.00 ± 27.40 | 8 | 13.89 [12.08, 15.73]* | 1.39 [-0.49, 3.24] | 10.49 [7.71, 13.23]* | -1.3 [-3.06, 0.4] | -14.5 [-16.21, -12.76]* | -27.16 [-29.03, -25.19]* | 13.2 [11.38, 14.9]* | -25.86 [-27.88, -24.01]* | -12.67 [-14.55, -10.7]* |
| L 470 | L 290 | L Subthalamic nucleus | 27.44 ± 24.44 | 10 | 8.09 ± 3.51 | 15 | 44.92 ± 10.07 | 11 | 88.79 ± 27.16 | 9 | 0.13 [-1.81, 2.07] | -0.59 [-2.43, 1.25] | 1.45 [-1.31, 4.23] | | | | | |
| R 614 | R 290 | R Tuberal nucleus | 78.43 ± 41.56 | 14 | 39.17 ± 15.23 | 13 | 48.96 ± 18.53 | 10 | 86.11 ± 21.49 | 9 | -0.81 [-2.66, 1.04] | -0.27 [-2.05, 1.48] | -0.02 [-2.71, 2.75] | | | | | |
| L 614 | L 290 | L Tuberal nucleus | 78.77 ± 50.12 | 14 | 24.59 ± 9.62 | 13 | 32.32 ± 8.79 | 10 | 41.42 ± 7.67 | 9 | -1.04 [-2.91, 0.86] | -0.47 [-2.24, 1.31] | -0.18 [-2.94, 2.55] | | | | | |
| R 797 | R 290 | R Zona incerta | 35.59 ± 12.19 | 15 | 43.43 ± 17.94 | 15 | 110.49 ± 20.98 | 12 | 142.45 ± 33.21 | 10 | 1.58 [-0.23, 3.38] | 0.3 [-1.41, 2.01] | -0.12 [-2.7, 2.49] | | | | | |
| L 797 | L 290 | L Zona incerta | 17.43 ± 6.44 | 15 | 13.00 ± 3.81 | 15 | 82.26 ± 12.95 | 12 | 119.90 ± 24.67 | 10 | 3.18 [1.41, 4.98]* | -0.17 [-1.89, 1.53] | 1.62 [-1, 4.25] | 0.25 [-1.3, 1.86] | -3.79 [-5.44, -2.12]* | -6.03 [-7.75, -4.29]* | 4.05 [2.34, 5.69]* | -6.29 [-8.05, -4.54]* | -2.25 [-4.06, -0.36]* |
| R 804 | R 797 | R Fields of Forel | 21.56 ± 14.47 | 10 | 6.13 ± 2.17 | 10 | 116.28 ± 27.14 | 8 | 159.17 ± 40.25 | 8 | 4.03 [1.98, 6.08]* | -0.27 [-2.25, 1.66] | 1.7 [-1.25, 4.59] | 0.35 [-1.42, 2.25] | -4.64 [-6.67, -2.76]* | -6.86 [-8.81, -4.89]* | 5 [3.03, 7]* | -7.23 [-9.16, -5.23]* | -2.22 [-4.28, -0.15]* |
| L 804 | L 797 | L Fields of Forel | 14.47 ± 9.86 | 10 | 6.76 ± 3.09 | 9 | 80.49 ± 22.78 | 9 | 147.48 ± 30.36 | 8 | 4.3 [2.33, 6.31]* | -0.11 [-2.11, 1.89] | 4.09 [1.11, 7.09]* | 0.21 [-1.69, 2.05] | -4.9 [-6.82, -3.04]* | -9.69 [-11.66, -7.63]* | 5.11 [3.1, 7.03]* | -9.87 [-11.91, -7.86]* | -4.78 [-6.79, -2.7]* |
| R 10671 | R 1097 | R Median eminence | 370.20 ± 175.50 | 12 | 123.49 ± 60.37 | 13 | 86.38 ± 37.80 | 9 | 527.81 ± 229.09 | 9 | -1.13 [-3.06, 0.81] | -0.5 [-2.32, 1.35] | 0.89 [-1.91, 3.69] | | | | | |
| L 10671 | L 1097 | L Median eminence | 388.51 ± 217.62 | 12 | 165.39 ± 117.44 | 13 | 166.01 ± 141.29 | 10 | 633.30 ± 276.05 | 9 | -0.91 [-2.82, 0.95] | -0.39 [-2.21, 1.41] | 0.78 [-1.93, 3.55] | | | | | |
| R 313 | R 343 | R Midbrain | 46.07 ± 33.32 | 13 | 14.19 ± 5.33 | 12 | 26.34 ± 7.51 | 12 | 70.78 ± 26.74 | 8 | -1.17 [-6.25, 3.98] | -0.24 [-5.42, 4.92] | -0.44 [-8.39, 7.35] | | | | | |
| L 313 | L 343 | L Midbrain | 17.42 ± 9.95 | 13 | 15.86 ± 10.31 | 14 | 19.87 ± 5.91 | 12 | 41.98 ± 10.21 | 9 | -0.61 [-5.79, 4.66] | 0.04 [-4.99, 5.15] | -0.42 [-8.19, 7.3] | | | | | |
| R 339 | R 313 | R Midbrain, sensory related | 7.20 ± 4.90 | 11 | 18.73 ± 6.44 | 10 | 55.32 ± 31.88 | 10 | 17.40 ± 7.05 | 7 | 6.05 [0.33, 11.8]* | 1.74 [-3.98, 7.35] | -8.49 [-17.07, 0.04] | -1.71 [-7.26, 3.89] | -6.98 [-12.24, -1]* | -1.92 [-8.18, 4.33] | 5.28 [-0.45, 11.23] | -0.27 [-6.74, 5.8] | 4.95 [-1.25, 11.54] |
| L 339 | L 313 | L Midbrain, sensory related | 8.74 ± 7.90 | 11 | 8.88 ± 3.23 | 10 | 119.58 ± 79.23 | 10 | 15.52 ± 5.10 | 7 | 11.99 [6.2, 17.72]* | 0.19 [-5.62, 5.86] | -13.5 [-22.14, -4.77]* | -0.17 [-5.77, 5.64] | -12.91 [-18.61, -7.33]* | -1.32 [-7.46, 4.8] | 12.77 [7.02, 18.41]* | -1.18 [-7.34, 5.15] | 11.57 [5.19, 17.89]* |
| R 599626923 | R 339 | R Subcommissural organ | 51.95 ± 51.95 | 11 | 0.00 ± 0.00 | 8 | 130.94 ± 62.43 | 10 | 0.00 ± 0.00 | 7 | 0.88 [-4.8, 6.49] | -0.42 [-6.43, 5.53] | -3.54 [-12.18, 5.03] | | | | | |
| L 599626923 | L 339 | L Subcommissural organ | 23.27 ± 23.27 | 11 | 0.00 ± 0.00 | 8 | 473.17 ± 332.90 | 10 | 31.32 ± 23.01 | 7 | 18.66 [12.92, 24.39]* | -0.42 [-6.43, 5.5] | -19.97 [-28.77, -11.26]* | 0.49 [-5.57, 6.24] | -19.54 [-25.34, -14.09]* | -0.94 [-7.21, 5.23] | 20.06 [13.81, 25.99]* | -1.37 [-7.93, 5.29] | 18.66 [12.36, 24.87]* |
| R 323 | R 313 | R Midbrain, motor related | 49.73 ± 43.16 | 13 | 20.43 ± 8.81 | 13 | 26.14 ± 6.98 | 12 | 74.73 ± 31.40 | 8 | | | | | | | | | |
| L 323 | L 313 | L Midbrain, motor related | 16.78 ± 10.94 | 13 | 3.79 ± 1.17 | 14 | 14.07 ± 4.83 | 12 | 23.96 ± 5.13 | 8 | -0.9 [-6.1, 4.22] | -0.67 [-5.64, 4.36] | -0.39 [-8.19, 7.55] | | | | | |
| L 381 | L 323 | L Substantia nigra, reticular part | 29.35 ± 27.22 | 6 | 3.91 ± 2.69 | 7 | 10.73 ± 4.94 | 7 | 22.81 ± 10.68 | 7 | -1.22 [-8.37, 5.95] | -0.58 [-7.58, 6.57] | -0.72 [-10.7, 9.41] | | | | | |
| R 128 | R 323 | R Midbrain reticular nucleus | 18.86 ± 14.81 | 8 | 13.13 ± 7.73 | 9 | 74.69 ± 34.09 | 9 | 97.43 ± 16.67 | 6 | 1.8 [-4.49, 8.09] | -0.61 [-6.88, 5.72] | 0.66 [-8.74, 10.02] | | | | | |
| L 128 | L 323 | L Midbrain reticular nucleus | 12.61 ± 7.67 | 8 | 6.14 ± 2.73 | 9 | 107.45 ± 49.14 | 9 | 115.01 ± 28.23 | 7 | 6.43 [0.21, 12.75]* | -0.75 [-6.98, 5.5] | 0.02 [-9.05, 8.88] | 0.79 [-5.39, 6.89] | -7.32 [-13.7, -1.44]* | -8.34 [-14.93, -1.87]* | 8.13 [2.12, 13.97]* | -9.1 [-15.73, -2.87]* | -0.99 [-7.34, 5.51] |
| R 795 | R 323 | R Periaqueductal gray | 60.32 ± 44.36 | 13 | 19.42 ± 10.33 | 13 | 25.90 ± 6.13 | 12 | 34.38 ± 6.72 | 8 | -1.34 [-6.53, 3.96] | -0.55 [-5.7, 4.67] | -0.92 [-8.75, 6.94] | | | | | |
| L 795 | L 323 | L Periaqueductal gray | 32.38 ± 17.10 | 13 | 9.55 ± 3.64 | 13 | 15.82 ± 6.06 | 12 | 24.34 ± 4.73 | 8 | -1.25 [-6.43, 4.05] | -0.59 [-5.79, 4.57] | -0.83 [-8.75, 7.23] | | | | | |
| R 50 | R 795 | R Precommissural nucleus | 129.02 ± 101.46 | 10 | 54.64 ± 41.27 | 12 | 49.27 ± 14.22 | 12 | 44.49 ± 16.18 | 8 | -1.2 [-6.81, 4.35] | -0.22 [-5.77, 5.23] | -1.46 [-9.66, 6.88] | | | | | |
| L 50 | L 795 | L Precommissural nucleus | 49.97 ± 29.18 | 10 | 12.55 ± 6.94 | 12 | 25.40 ± 14.83 | 12 | 109.77 ± 88.66 | 8 | -1.11 [-6.73, 4.54] | -0.45 [-6.11, 5.18] | 0.52 [-7.71, 8.76] | | | | | |
| R 1100 | R 323 | R Pretectal region | 107.69 ± 100.48 | 11 | 388.61 ± 349.69 | 11 | 53.19 ± 18.65 | 11 | 266.43 ± 193.53 | 8 | -1.19 [-6.82, 4.35] | 2.8 [-2.61, 8.36] | -2.46 [-10.57, 5.76] | | | | | |
| L 1100 | L 323 | L Pretectal region | 28.89 ± 21.88 | 12 | 4.88 ± 1.61 | 10 | 22.25 ± 6.76 | 10 | 37.75 ± 8.15 | 7 | -0.92 [-6.48, 4.68] | -0.4 [-5.88, 5.08] | -0.52 [-9.07, 8.03] | | | | | |
| R 215 | R 1100 | R Anterior pretectal nucleus | 235.65 ± 224.06 | 11 | 753.95 ± 700.98 | 11 | 109.58 ± 37.15 | 11 | 559.69 ± 385.94 | 8 | -1.24 [-6.85, 4.26] | 2.38 [-3.1, 7.94] | -2.08 [-10.35, 6.15] | | | | | |
| L 215 | L 1100 | L Anterior pretectal nucleus | 54.45 ± 44.47 | 12 | 9.32 ± 3.19 | 10 | 54.50 ± 17.16 | 10 | 86.13 ± 19.55 | 7 | -0.71 [-6.33, 4.85] | -0.4 [-5.92, 5.13] | -0.45 [-9.02, 8.18] | | | | | |
| R 628 | R 1100 | R Nucleus of the optic tract | 37.68 ± 19.92 | 8 | 531.48 ± 325.14 | 7 | 242.80 ± 128.48 | 8 | 48.75 ± 27.27 | 7 | 3.95 [-2.45, 10.48] | 13.29 [6.64, 20.02]* | -19.37 [-28.83, -9.78]* | -13.23 [-19.88, -6.63]* | -4.85 [-11.5, 1.35] | -0.49 [-7.21, 5.94] | -8.4 [-15.02, -1.61]* | 12.72 [5.58, 19.3]* | 4.37 [-2.31, 11.02] |
| L 628 | L 1100 | L Nucleus of the optic tract | 24.98 ± 9.95 | 8 | 8.54 ± 6.16 | 7 | 27.67 ± 12.32 | 9 | 44.98 ± 22.42 | 7 | -1.07 [-7.48, 5.18] | -0.42 [-7.12, 6.27] | -0.11 [-9.39, 9.54] | | | | | |
| R 634 | R 1100 | R Nucleus of the posterior commissure | 169.70 ± 152.90 | 9 | 17.55 ± 5.16 | 7 | 25.81 ± 16.08 | 8 | 50.26 ± 17.86 | 7 | -1.91 [-8.2, 4.36] | -0.26 [-6.83, 6.33] | -0.51 [-9.93, 8.99] | | | | | |
| L 634 | L 1100 | L Nucleus of the posterior commissure | 83.32 ± 56.59 | 9 | 10.91 ± 5.09 | 7 | 27.19 ± 10.37 | 8 | 46.40 ± 16.32 | 7 | -1.75 [-8.07, 4.7] | -0.23 [-6.72, 6.25] | -0.46 [-9.77, 8.88] | | | | | |
| R 706 | R 1100 | R Olivary pretectal nucleus | 169.57 ± 123.91 | 8 | 67.76 ± 30.35 | 7 | 52.99 ± 24.74 | 7 | 175.73 ± 133.90 | 7 | -2.08 [-8.76, 4.56] | 0.05 [-6.55, 6.49] | 0.11 [-9.5, 9.73] | | | | | |
| L 706 | L 1100 | L Olivary pretectal nucleus | 51.72 ± 46.14 | 6 | 21.61 ± 13.07 | 7 | 23.63 ± 9.81 | 6 | 42.37 ± 21.85 | 7 | -2.59 [-10.03, 4.84] | -0.35 [-7.51, 6.76] | 0.34 [-9.97, 10.44] | | | | | |
| L 1061 | L 1100 | L Posterior pretectal nucleus | 10.80 ± 4.26 | 8 | 6.82 ± 3.66 | 7 | 22.34 ± 11.49 | 8 | 41.74 ± 22.01 | 7 | -0.33 [-6.57, 5.99] | -0.17 [-6.71, 6.42] | 0.93 [-8.39, 10.27] | | | | | |
| R 1009 | R 997 | R fiber tracts | 49.43 ± 11.92 | 15 | 44.88 ± 8.45 | 15 | 143.96 ± 21.77 | 12 | 179.27 ± 25.61 | 10 | 1.7 [-0.33, 3.66] | -0.03 [-1.91, 1.8] | 0.15 [-2.71, 3.12] | | | | | |
| L 1009 | L 997 | L fiber tracts | 12.93 ± 4.34 | 15 | 11.90 ± 2.84 | 15 | 24.10 ± 5.18 | 12 | 52.17 ± 9.51 | 10 | 0.65 [-1.32, 2.59] | -0.01 [-1.9, 1.86] | 1.6 [-1.31, 4.5] | | | | | |
| R 967 | R 1009 | R cranial nerves | 26.62 ± 11.38 | 15 | 25.26 ± 7.47 | 15 | 17.63 ± 6.33 | 12 | 57.56 ± 17.62 | 10 | -0.54 [-2.51, 1.41] | 0.02 [-1.86, 1.9] | 0.88 [-2.02, 3.83] | | | | | |
| L 967 | L 1009 | L cranial nerves | 18.92 ± 8.80 | 15 | 8.96 ± 3.61 | 15 | 13.68 ± 4.13 | 12 | 86.71 ± 39.32 | 10 | -0.49 [-2.46, 1.49] | -0.45 [-2.25, 1.41] | 3.73 [0.8, 6.58]* | 0.54 [-1.16, 2.26] | 0.1 [-1.83, 1.91] | -3.82 [-5.76, -1.91]* | 0.44 [-1.44, 2.31] | -4.36 [-6.35, -2.47]* | -3.91 [-6.04, -1.89]* |
| R 840 | R 967 | R olfactory nerve | 24.71 ± 15.28 | 15 | 10.31 ± 3.73 | 15 | 6.65 ± 1.18 | 12 | 26.08 ± 12.27 | 10 | -0.94 [-2.93, 1] | -0.5 [-2.36, 1.36] | 0.7 [-2.22, 3.65] | | | | | |
| L 840 | L 967 | L olfactory nerve | 11.90 ± 4.32 | 15 | 9.64 ± 2.28 | 15 | 7.23 ± 2.47 | 12 | 25.95 ± 11.60 | 10 | -0.6 [-2.59, 1.39] | -0.11 [-2.03, 1.77] | 1.1 [-1.86, 4.03] | | | | | |
| R 21 | R 840 | R lateral olfactory tract, general | 52.27 ± 36.18 | 15 | 18.84 ± 7.78 | 15 | 7.62 ± 2.19 | 12 | 29.99 ± 18.63 | 10 | -1.05 [-3.01, 0.9] | -0.55 [-2.43, 1.33] | 0.38 [-2.6, 3.28] | | | | | |
| L 21 | L 840 | L lateral olfactory tract, general | 18.61 ± 7.10 | 15 | 16.48 ± 6.18 | 15 | 7.58 ± 3.95 | 12 | 13.55 ± 6.23 | 10 | | | | | | | | | |
| R 665 | R 21 | R lateral olfactory tract, body | 104.53 ± 72.36 | 15 | 37.68 ± 15.56 | 15 | 15.23 ± 4.37 | 12 | 59.97 ± 37.26 | 10 | -1.07 [-3.03, 0.89] | -0.57 [-2.46, 1.28] | 0.43 [-2.57, 3.37] | | | | | |
| L 665 | L 21 | L lateral olfactory tract, body | 37.23 ± 14.20 | 15 | 32.97 ± 12.36 | 15 | 15.15 ± 7.91 | 12 | 27.09 ± 12.47 | 10 | -0.82 [-2.81, 1.17] | -0.05 [-1.9, 1.81] | -0.2 [-3.07, 2.71] | | | | | |
| R 900 | R 840 | R anterior commissure, olfactory limb | 16.92 ± 8.35 | 15 | 13.80 ± 4.48 | 14 | 17.62 ± 3.17 | 12 | 52.60 ± 19.85 | 9 | -0.17 [-2.13, 1.79] | -0.09 [-1.95, 1.81] | 1.57 [-1.37, 4.55] | | | | | |
| L 900 | L 840 | L anterior commissure, olfactory limb | 21.04 ± 8.80 | 15 | 19.06 ± 8.53 | 14 | 22.75 ± 8.82 | 12 | 118.79 ± 56.55 | 9 | -0.13 [-2.11, 1.83] | 0 [-1.96, 1.91] | 3.98 [1.01, 6.97]* | 0.08 [-1.72, 1.86] | -0.26 [-2.12, 1.55] | -4.88 [-6.9, -2.86]* | 0.36 [-1.57, 2.19] | -4.96 [-7.06, -2.92]* | -4.62 [-6.8, -2.64]* |
| R 848 | R 967 | R optic nerve | 64.81 ± 30.68 | 15 | 80.74 ± 25.95 | 15 | 39.40 ± 17.31 | 12 | 137.09 ± 48.33 | 10 | -0.6 [-2.57, 1.36] | 0.31 [-1.52, 2.13] | 0.62 [-2.31, 3.54] | | | | | |
| L 848 | L 967 | L optic nerve | 47.68 ± 26.63 | 15 | 21.23 ± 9.93 | 15 | 29.74 ± 10.32 | 12 | 203.66 ± 96.87 | 10 | -0.59 [-2.58, 1.36] | -0.49 [-2.35, 1.34] | 3.54 [0.62, 6.5]* | 0.56 [-1.08, 2.35] | 0.17 [-1.59, 2.13] | -3.5 [-5.5, -1.56]* | 0.38 [-1.58, 2.2] | -4.09 [-6.06, -2.09]* | -3.69 [-5.75, -1.61]* |
| R 916 | R 848 | R brachium of the superior colliculus | 617.44 ± 558.13 | 10 | 652.16 ± 284.02 | 10 | 336.43 ± 103.58 | 10 | 641.74 ± 297.80 | 8 | -0.44 [-2.63, 1.72] | 0.46 [-1.75, 2.64] | -0.36 [-3.55, 2.93] | | | | | |
| L 916 | L 848 | L brachium of the superior colliculus | 66.80 ± 43.16 | 11 | 62.85 ± 31.42 | 9 | 125.08 ± 82.93 | 10 | 154.96 ± 58.14 | 7 | 0.8 [-1.39, 2.92] | 0.33 [-1.91, 2.55] | -0.13 [-3.49, 3.17] | | | | | |
| R 125 | R 848 | R optic tract | 138.35 ± 66.54 | 15 | 131.67 ± 52.32 | 15 | 54.57 ± 26.46 | 12 | 282.88 ± 116.67 | 10 | -0.81 [-2.76, 1.14] | 0.02 [-1.82, 1.9] | 1.03 [-1.91, 3.9] | | | | | |
| L 125 | L 848 | L optic tract | 112.59 ± 64.93 | 15 | 48.15 ± 29.04 | 15 | 61.62 ± 30.12 | 12 | 525.56 ± 297.51 | 10 | -0.65 [-2.71, 1.33] | -0.49 [-2.42, 1.4] | 4.03 [1.14, 7.03]* | 0.57 [-1.15, 2.4] | 0.24 [-1.64, 2.16] | -3.91 [-5.88, -1.92]* | 0.32 [-1.56, 2.21] | -4.48 [-6.48, -2.56]* | -4.16 [-6.26, -2.12]* |
| R 832 | R 967 | R oculomotor nerve | 36.05 ± 24.45 | 10 | 8.85 ± 5.08 | 9 | 10.21 ± 8.43 | 9 | 18.24 ± 9.27 | 7 | -0.77 [-3.02, 1.47] | -0.41 [-2.64, 1.83] | 0.48 [-2.88, 3.8] | | | | | |
| L 832 | L 967 | L oculomotor nerve | 12.44 ± 11.02 | 9 | 8.38 ± 6.47 | 9 | 3.78 ± 3.78 | 9 | 17.77 ± 15.56 | 7 | -0.74 [-3.03, 1.53] | 0.05 [-2.24, 2.33] | 0.91 [-2.44, 4.33] | | | | | |
| R 158 | R 832 | R posterior commissure | 72.11 ± 48.91 | 10 | 13.61 ± 10.77 | 7 | 20.42 ± 16.85 | 9 | 35.67 ± 19.29 | 7 | -0.78 [-3.06, 1.41] | -0.33 [-2.69, 2.05] | 0.39 [-3.17, 3.83] | | | | | |
| L 158 | L 832 | L posterior commissure | 24.88 ± 22.05 | 9 | 4.62 ± 3.10 | 7 | 7.57 ± 7.57 | 9 | 35.54 ± 31.11 | 7 | -0.75 [-3.07, 1.57] | -0.33 [-2.75, 2.12] | 1.31 [-2.15, 4.88] | | | | | |
| R 792 | R 967 | R dorsal roots | 6.76 ± 3.21 | 10 | 2.84 ± 1.45 | 11 | 20.62 ± 7.65 | 10 | 28.57 ± 9.56 | 8 | 2.23 [0.07, 4.41]* | -0.24 [-2.36, 1.92] | 0.86 [-2.34, 4.02] | 0.32 [-1.71, 2.38] | -2.63 [-4.69, -0.54]* | -3.89 [-6.17, -1.68]* | 2.95 [0.81, 4.99]* | -4.21 [-6.44, -2.11]* | -1.25 [-3.48, 1.09] |
| L 792 | L 967 | L dorsal roots | 4.94 ± 2.79 | 10 | 1.68 ± 0.83 | 11 | 13.44 ± 5.01 | 10 | 24.10 ± 7.17 | 8 | 1.91 [-0.31, 4.09] | -0.38 [-2.56, 1.78] | 1.97 [-1.25, 5.17] | | | | | |
| R 932 | R 792 | R cervicothalamic tract | 10.15 ± 4.81 | 10 | 4.27 ± 2.17 | 11 | 30.94 ± 11.48 | 10 | 42.86 ± 14.34 | 8 | 2.23 [0.04, 4.4]* | -0.24 [-2.42, 1.95] | 0.86 [-2.39, 4.08] | 0.32 [-1.81, 2.32] | -2.64 [-4.7, -0.57]* | -3.9 [-6.09, -1.59]* | 2.96 [0.89, 5.05]* | -4.22 [-6.43, -1.99]* | -1.27 [-3.55, 0.98] |
| L 932 | L 792 | L cervicothalamic tract | 7.42 ± 4.18 | 10 | 2.51 ± 1.24 | 11 | 20.15 ± 7.52 | 10 | 36.15 ± 10.76 | 8 | 1.91 [-0.26, 4.09] | -0.37 [-2.48, 1.74] | 1.97 [-1.26, 5.16] | | | | | |
| R 697 | R 932 | R medial lemniscus | 20.29 ± 9.63 | 10 | 8.53 ± 4.34 | 11 | 61.87 ± 22.96 | 10 | 85.71 ± 28.68 | 8 | 2.23 [0.05, 4.39]* | -0.26 [-2.36, 1.91] | 0.86 [-2.33, 4.12] | 0.34 [-1.77, 2.3] | -2.64 [-4.81, -0.65]* | -3.89 [-6.19, -1.74]* | 2.96 [0.92, 4.95]* | -4.21 [-6.38, -2.01]* | -1.25 [-3.51, 0.91] |
| L 697 | L 932 | L medial lemniscus | 14.83 ± 8.37 | 10 | 5.03 ± 2.48 | 11 | 40.31 ± 15.03 | 10 | 72.29 ± 21.52 | 8 | 1.89 [-0.3, 4.08] | -0.39 [-2.51, 1.78] | 1.99 [-1.28, 5.26] | | | | | |
| R 484682512 | R 1009 | R supra-callosal cerebral white matter | 184.43 ± 69.10 | 15 | 170.87 ± 80.90 | 15 | 319.85 ± 75.99 | 12 | 557.55 ± 87.73 | 10 | 0.52 [-1.44, 2.5] | 0 [-1.88, 1.83] | 0.71 [-2.23, 3.62] | | | | | |
| L 484682512 | L 1009 | L supra-callosal cerebral white matter | 27.23 ± 12.53 | 15 | 38.47 ± 12.02 | 15 | 32.88 ± 9.56 | 12 | 129.40 ± 65.28 | 10 | -0.01 [-2.03, 1.97] | 0.47 [-1.42, 2.36] | 2.48 [-0.47, 5.42] | | | | | |
| R 983 | R 1009 | R lateral forebrain bundle system | 63.46 ± 15.03 | 15 | 55.80 ± 13.68 | 15 | 141.84 ± 21.70 | 12 | 179.96 ± 22.64 | 10 | 1.04 [-0.94, 3.07] | -0.03 [-1.94, 1.84] | 0.04 [-2.85, 2.98] | | | | | |
| L 983 | L 1009 | L lateral forebrain bundle system | 13.41 ± 5.08 | 15 | 12.36 ± 2.96 | 15 | 19.98 ± 3.62 | 12 | 51.01 ± 12.53 | 10 | 0.29 [-1.69, 2.28] | 0 [-1.88, 1.86] | 1.71 [-1.27, 4.7] | | | | | |
| R 776 | R 983 | R corpus callosum | 106.41 ± 25.89 | 15 | 109.07 ± 30.58 | 15 | 289.64 ± 47.01 | 12 | 365.30 ± 49.34 | 10 | 1.52 [-0.44, 3.44] | 0.1 [-1.75, 1.95] | 0.02 [-2.96, 2.95] | | | | | |
| L 776 | L 983 | L corpus callosum | 21.15 ± 7.45 | 15 | 26.97 ± 7.25 | 15 | 43.56 ± 7.93 | 12 | 98.42 ± 24.90 | 10 | 0.84 [-1.14, 2.81] | 0.33 [-1.56, 2.18] | 1.68 [-1.24, 4.61] | | | | | |
| R 956 | R 776 | R corpus callosum, anterior forceps | 16.65 ± 4.03 | 15 | 22.87 ± 6.44 | 15 | 34.18 ± 8.74 | 12 | 88.50 ± 27.18 | 10 | 0.84 [-1.13, 2.82] | 0.45 [-1.44, 2.33] | 2.23 [-0.73, 5.16] | | | | | |
| L 956 | L 776 | L corpus callosum, anterior forceps | 15.06 ± 6.73 | 15 | 14.81 ± 5.89 | 15 | 25.98 ± 4.83 | 12 | 54.57 ± 16.23 | 10 | 0.52 [-1.46, 2.48] | 0.05 [-1.78, 1.9] | 1.25 [-1.64, 4.17] | | | | | |
| R 579 | R 956 | R external capsule | 33.99 ± 8.40 | 15 | 32.65 ± 8.09 | 15 | 84.35 ± 19.36 | 12 | 200.01 ± 53.34 | 10 | 1.29 [-0.68, 3.24] | 0.05 [-1.8, 1.88] | 2.75 [-0.16, 5.63] | | | | | |
| L 579 | L 956 | L external capsule | 31.70 ± 14.00 | 15 | 17.61 ± 3.97 | 15 | 58.10 ± 9.37 | 12 | 89.02 ± 18.35 | 10 | 0.63 [-1.32, 2.56] | -0.37 [-2.23, 1.48] | 0.76 [-2.16, 3.66] | | | | | |
| R 964 | R 776 | R corpus callosum, extreme capsule | 31.57 ± 12.45 | 15 | 34.44 ± 15.81 | 15 | 109.01 ± 37.37 | 12 | 135.05 ± 39.98 | 10 | 2.25 [0.28, 4.21]* | 0.17 [-1.69, 2.05] | 0.06 [-2.84, 2.94] | -0.09 [-1.85, 1.66] | -2.64 [-4.45, -0.78]* | -3.51 [-5.49, -1.56]* | 2.56 [0.69, 4.37]* | -3.44 [-5.39, -1.49]* | -0.89 [-2.92, 1.2] |
| L 964 | L 776 | L corpus callosum, extreme capsule | 14.66 ± 9.68 | 15 | 15.82 ± 6.94 | 15 | 51.23 ± 15.66 | 12 | 59.83 ± 16.44 | 10 | 2.3 [0.35, 4.24]* | 0.16 [-1.71, 2.02] | -0.16 [-3.08, 2.74] | -0.09 [-1.81, 1.68] | -2.7 [-4.5, -0.85]* | -3.34 [-5.28, -1.34]* | 2.62 [0.73, 4.42]* | -3.25 [-5.25, -1.36]* | -0.64 [-2.72, 1.45] |
| R 1108 | R 776 | R genu of corpus callosum | 25.53 ± 11.81 | 14 | 35.87 ± 19.28 | 11 | 34.70 ± 12.04 | 11 | 63.60 ± 20.90 | 8 | 0.3 [-1.75, 2.33] | 0.58 [-1.43, 2.57] | -0.04 [-3.2, 3.04] | | | | | |
| L 1108 | L 776 | L genu of corpus callosum | 18.60 ± 11.16 | 14 | 27.92 ± 10.22 | 10 | 30.33 ± 13.05 | 10 | 92.22 ± 47.55 | 9 | 0.55 [-1.52, 2.63] | 0.76 [-1.31, 2.81] | 1.89 [-1.24, 4.99] | | | | | |
| R 971 | R 776 | R corpus callosum, posterior forceps | 705.24 ± 147.11 | 11 | 522.06 ± 188.66 | 12 | 1441.98 ± 443.04 | 10 | 1858.05 ± 399.63 | 9 | 0.99 [-1.14, 3.06] | 0.04 [-2.01, 2.08] | 0.07 [-3.05, 3.2] | | | | | |
| L 971 | L 776 | L corpus callosum, posterior forceps | 87.24 ± 36.94 | 12 | 70.89 ± 24.71 | 10 | 138.38 ± 35.45 | 10 | 372.80 ± 125.50 | 8 | 0.47 [-1.68, 2.55] | 0.18 [-1.97, 2.31] | 2.13 [-1.05, 5.31] | | | | | |
| R 484682516 | R 776 | R corpus callosum, body | 189.01 ± 49.52 | 15 | 195.44 ± 54.88 | 13 | 570.95 ± 115.50 | 12 | 802.53 ± 201.16 | 10 | 1.8 [-0.13, 3.74] | 0.13 [-1.77, 2.04] | 0.52 [-2.41, 3.46] | | | | | |
| L 484682516 | L 776 | L corpus callosum, body | 46.99 ± 16.65 | 15 | 68.39 ± 31.10 | 13 | 80.13 ± 22.82 | 12 | 304.41 ± 125.95 | 10 | 0.49 [-1.52, 2.43] | 0.56 [-1.37, 2.51] | 3.62 [0.65, 6.58]* | -0.48 [-2.36, 1.24] | -0.91 [-2.72, 0.95] | -5.71 [-7.61, -3.67]* | 0.41 [-1.47, 2.33] | -5.24 [-7.27, -3.2]* | -4.81 [-6.9, -2.82]* |
| R 986 | R 776 | R corpus callosum, splenium | 644.85 ± 189.46 | 13 | 513.17 ± 173.47 | 15 | 1340.09 ± 249.82 | 12 | 1382.18 ± 338.32 | 9 | 0.97 [-1, 2.96] | -0.03 [-1.92, 1.88] | -0.5 [-3.54, 2.4] | | | | | |
| L 986 | L 776 | L corpus callosum, splenium | 86.26 ± 33.95 | 13 | 94.78 ± 35.24 | 15 | 207.78 ± 49.78 | 12 | 356.71 ± 118.77 | 10 | 1.3 [-0.7, 3.33] | 0.27 [-1.7, 2.21] | 0.86 [-2.04, 3.79] | | | | | |
| R 784 | R 983 | R corticospinal tract | 41.70 ± 19.71 | 15 | 13.93 ± 4.73 | 15 | 22.53 ± 7.83 | 12 | 58.88 ± 16.61 | 10 | -0.67 [-2.63, 1.32] | -0.61 [-2.51, 1.26] | 0.88 [-2.04, 3.81] | | | | | |
| L 784 | L 983 | L corticospinal tract | 11.90 ± 6.49 | 15 | 5.10 ± 1.65 | 15 | 6.88 ± 1.87 | 12 | 36.20 ± 12.74 | 10 | -0.63 [-2.59, 1.26] | -0.49 [-2.35, 1.38] | 2.36 [-0.51, 5.25] | | | | | |
| R 6 | R 784 | R internal capsule | 77.42 ± 37.50 | 15 | 16.14 ± 5.50 | 14 | 23.52 ± 4.81 | 12 | 84.33 ± 21.52 | 10 | -0.9 [-2.87, 1.05] | -0.68 [-2.56, 1.22] | 0.88 [-2.1, 3.83] | | | | | |
| L 6 | L 784 | L internal capsule | 20.14 ± 9.44 | 15 | 9.56 ± 3.17 | 15 | 14.12 ± 3.55 | 12 | 76.21 ± 33.40 | 10 | -0.51 [-2.5, 1.43] | -0.44 [-2.34, 1.44] | 2.95 [0.04, 5.91]* | 0.52 [-1.32, 2.22] | 0.1 [-1.78, 1.99] | -3.05 [-5.02, -1.06]* | 0.41 [-1.5, 2.26] | -3.56 [-5.56, -1.65]* | -3.15 [-5.22, -1.04]* |
| R 924 | R 784 | R cerebal peduncle | 107.17 ± 101.58 | 7 | 69.17 ± 31.55 | 9 | 173.06 ± 83.99 | 8 | 286.44 ± 132.42 | 7 | 0.51 [-1.97, 2.96] | -0.24 [-2.67, 2.26] | 0.88 [-2.77, 4.47] | | | | | |
| L 924 | L 784 | L cerebal peduncle | 60.91 ± 52.75 | 9 | 29.21 ± 24.12 | 9 | 11.75 ± 7.14 | 9 | 66.50 ± 15.26 | 7 | -0.6 [-2.83, 1.68] | -0.13 [-2.34, 2.15] | 0.62 [-2.73, 4.02] | | | | | |

| | | | | | | | | | | | | | | | | | | | |
|---|---|---|---|---|---|---|---|---|---|---|---|---|---|---|---|---|---|---|---|
| R 896 | R 983 | R thalamus related | 152.34 ± 50.45 | 15 | 92.01 ± 20.61 | 15 | 212.24 ± 41.23 | 12 | 238.98 ± 44.53 | 10 | 0.18 [-1.76, 2.12] | -0.33 [-2.21, 1.55] | -0.08 [-3.02, 2.81] | | | | | | |
| L 896 | L 983 | L thalamus related | 25.07 ± 11.40 | 15 | 13.88 ± 3.67 | 15 | 17.98 ± 3.75 | 12 | 55.80 ± 16.35 | 10 | -0.48 [-2.44, 1.45] | -0.36 [-2.22, 1.51] | 1.28 [-1.61, 4.22] | | | | | | |
| R 1092 | R 896 | R external medullary lamina of the thalamus | 125.34 ± 102.16 | 15 | 76.67 ± 37.57 | 14 | 58.85 ± 15.71 | 12 | 72.31 ± 23.55 | 10 | -0.74 [-2.71, 1.21] | -0.3 [-2.22, 1.6] | -0.17 [-3.12, 2.8] | | | | | | |
| L 1092 | L 896 | L external medullary lamina of the thalamus | 28.89 ± 14.01 | 15 | 21.00 ± 15.18 | 13 | 26.46 ± 5.41 | 12 | 55.76 ± 14.99 | 10 | -0.28 [-2.22, 1.68] | -0.15 [-2.06, 1.76] | 0.57 [-2.34, 3.52] | | | | | | |
| R 484682520 | R 896 | R optic radiation | 357.52 ± 111.50 | 15 | 208.58 ± 51.65 | 15 | 523.66 ± 101.82 | 12 | 609.47 ± 112.86 | 9 | 0.26 [-1.71, 2.2] | -0.35 [-2.23, 1.56] | 0 [-3.05, 2.95] | | | | | | |
| L 484682520 | L 896 | L optic radiation | 58.17 ± 27.36 | 14 | 31.72 ± 8.64 | 15 | 36.20 ± 9.82 | 12 | 128.42 ± 40.85 | 10 | -0.52 [-2.54, 1.44] | -0.31 [-2.19, 1.57] | 1.3 [-1.65, 4.24] | | | | | | |
| R 484682524 | R 896 | R auditory radiation | 131.47 ± 79.58 | 14 | 109.44 ± 66.44 | 15 | 123.15 ± 55.00 | 12 | 174.76 ± 45.51 | 8 | -0.21 [-2.2, 1.76] | -0.03 [-1.91, 1.9] | -0.08 [-3.1, 2.99] | | | | | | |
| L 484682524 | L 896 | L auditory radiation | 44.18 ± 23.30 | 14 | 11.58 ± 5.39 | 15 | 37.57 ± 8.32 | 12 | 66.87 ± 16.29 | 10 | -0.29 [-2.27, 1.72] | -0.6 [-2.52, 1.24] | 0.68 [-2.26, 3.63] | | | | | | |
| R 1000 | R 1009 | R extrapyramidal fiber systems | 5.06 ± 2.92 | 15 | 9.62 ± 4.37 | 15 | 26.24 ± 5.83 | 12 | 25.07 ± 7.52 | 10 | 3.98 [2, 5.92]* | 0.97 [-0.92, 2.82] | -1.78 [-4.7, 1.12] | -0.9 [-2.7, 0.82] | -4.38 [-6.29, -2.6]* | -4.2 [-6.14, -2.24]* | 3.48 [1.69, 5.44]* | -3.31 [-5.36, -1.4]* | 0.2 [-1.83, 2.25] |
| L 1000 | L 1009 | L extrapyramidal fiber systems | 7.66 ± 4.21 | 15 | 8.01 ± 2.78 | 15 | 26.57 ± 6.63 | 12 | 28.58 ± 8.46 | 10 | 2.26 [0.26, 4.21]* | 0.12 [-1.76, 2.01] | -0.45 [-3.4, 2.51] | -0.04 [-1.77, 1.71] | -2.67 [-4.5, -0.77]* | -2.96 [-4.89, -0.96]* | 2.63 [0.61, 4.34]* | -2.93 [-4.82, -0.88]* | -0.3 [-2.32, 1.84] |
| R 760 | R 1000 | R cerebral nuclei related | 7.60 ± 4.38 | 15 | 14.72 ± 6.82 | 14 | 39.36 ± 8.75 | 12 | 38.60 ± 11.06 | 10 | 3.99 [1.98, 5.91]* | 1.04 [-0.84, 2.94] | -1.74 [-4.64, 1.16] | -0.97 [-2.76, 0.77] | -4.39 [-6.27, -2.55]* | -4.31 [-6.26, -2.31]* | 3.41 [1.55, 5.29]* | -3.37 [-5.37, -1.41]* | 0.06 [-2.01, 2.07] |
| L 760 | L 1000 | L cerebral nuclei related | 11.49 ± 6.32 | 15 | 14.17 ± 4.95 | 15 | 39.86 ± 9.95 | 12 | 43.81 ± 12.49 | 10 | 2.27 [0.21, 4.26]* | 0.31 [-1.59, 2.16] | -0.56 [-3.47, 2.35] | -0.23 [-2.01, 1.55] | -2.69 [-4.59, -0.77]* | -3.08 [-5.01, -1.11]* | 2.44 [0.62, 4.3]* | -2.83 [-4.76, -0.91]* | -0.39 [-2.5, 1.7] |
| R 102 | R 760 | R nigrostriatal tract | 15.19 ± 8.77 | 15 | 29.43 ± 13.65 | 14 | 78.72 ± 17.50 | 12 | 77.21 ± 22.12 | 10 | 3.98 [2.02, 5.91]* | 1.02 [-0.87, 2.87] | -1.71 [-4.63, 1.2] | -0.94 [-2.6, 0.9] | -4.38 [-6.18, -2.54]* | -4.33 [-6.34, -2.37]* | 3.45 [1.56, 5.26]* | -3.39 [-5.41, -1.41]* | 0.06 [-2.03, 2.06] |
| L 102 | L 760 | L nigrostriatal tract | 22.97 ± 12.64 | 15 | 28.33 ± 9.91 | 15 | 79.72 ± 19.90 | 12 | 87.62 ± 24.98 | 10 | 2.27 [0.33, 4.22]* | 0.31 [-1.52, 2.15] | -0.56 [-3.49, 2.29] | -0.23 [-1.98, 1.48] | -2.67 [-4.59, -0.88]* | -3.04 [-5.01, -1.11]* | 2.45 [0.56, 4.3]* | -2.83 [-4.79, -0.89]* | -0.39 [-2.47, 1.59] |
| R 991 | R 1009 | R medial forebrain bundle system | 62.75 ± 17.51 | 15 | 66.32 ± 12.53 | 15 | 287.98 ± 55.23 | 12 | 330.40 ± 70.81 | 10 | 3.39 [1.4, 5.36]* | 0.13 [-1.74, 2] | -0.04 [-2.99, 2.91] | -0.05 [-1.77, 1.68] | -3.79 [-5.61, -1.9]* | -4.51 [-6.43, -2.48]* | 3.75 [1.95, 5.63]* | -4.46 [-6.46, -2.55]* | -0.71 [-2.82, 1.32] |
| L 991 | L 1009 | L medial forebrain bundle system | 15.44 ± 4.66 | 15 | 19.01 ± 5.58 | 15 | 46.67 ± 11.15 | 12 | 78.22 ± 14.20 | 10 | 1.81 [-0.13, 3.77] | 0.3 [-1.56, 2.18] | 1.15 [-1.77, 4] | | | | | | |
| R 768 | R 991 | R cerebrum related | 92.93 ± 25.51 | 15 | 99.39 ± 19.63 | 15 | 434.75 ± 83.75 | 12 | 496.60 ± 108.00 | 10 | 3.46 [1.47, 5.4]* | 0.14 [-1.72, 2.03] | -0.05 [-3, 2.87] | -0.06 [-1.78, 1.7] | -3.86 [-5.69, -1.99]* | -4.6 [-6.54, -2.62]* | 3.8 [1.99, 5.65]* | -4.53 [-6.37, -2.49]* | -0.72 [-2.89, 1.29] |
| L 768 | L 991 | L cerebrum related | 21.26 ± 6.17 | 15 | 27.71 ± 8.87 | 15 | 67.15 ± 16.18 | 12 | 114.04 ± 20.53 | 10 | 1.95 [-0.05, 3.9] | 0.39 [-1.52, 2.28] | 1.23 [-1.7, 4.12] | | | | | | |
| R 884 | R 768 | R amygdalar capsule | 32.40 ± 17.76 | 15 | 25.53 ± 10.91 | 15 | 60.16 ± 14.57 | 12 | 104.35 ± 18.80 | 10 | 0.65 [-1.28, 2.56] | -0.14 [-2.01, 1.71] | 0.92 [-1.98, 3.86] | | | | | | |
| L 884 | L 768 | L amygdalar capsule | 22.97 ± 16.16 | 15 | 17.95 ± 7.37 | 15 | 50.42 ± 8.88 | 12 | 52.42 ± 14.74 | 10 | 0.99 [-0.99, 2.95] | -0.14 [-2.01, 1.73] | -0.37 [-3.36, 2.5] | | | | | | |
| R 940 | R 768 | R cingulum bundle | 141.13 ± 37.11 | 15 | 180.21 ± 86.67 | 15 | 273.44 ± 61.72 | 12 | 453.27 ± 71.35 | 10 | 0.75 [-1.2, 2.66] | 0.36 [-1.47, 2.21] | 0.31 [-2.63, 3.27] | | | | | | |
| L 940 | L 768 | L cingulum bundle | 44.69 ± 17.01 | 15 | 32.57 ± 11.61 | 15 | 89.45 ± 35.80 | 12 | 192.93 ± 73.04 | 10 | 0.78 [-1.2, 2.79] | -0.21 [-2.1, 1.65] | 1.95 [-0.98, 4.91] | | | | | | |
| R 1099 | R 768 | R fornix system | 152.91 ± 47.98 | 15 | 160.32 ± 29.76 | 15 | 757.87 ± 147.25 | 12 | 848.15 ± 188.38 | 10 | 3.75 [1.82, 5.72]* | 0.13 [-1.71, 1.99] | -0.13 [-3.01, 2.78] | -0.06 [-1.84, 1.64] | -4.16 [-6.03, -2.33]* | -4.8 [-6.73, -2.8]* | 4.11 [2.27, 5.99]* | -4.75 [-6.61, -2.7]* | -0.65 [-2.76, 1.29] |
| L 1099 | L 768 | L fornix system | 29.15 ± 8.47 | 15 | 44.57 ± 15.10 | 15 | 108.91 ± 26.25 | 12 | 178.56 ± 33.94 | 10 | 2.53 [0.55, 4.49]* | 0.6 [-1.3, 2.48] | 1.19 [-1.75, 4.13] | -0.52 [-2.27, 1.23] | -2.93 [-4.79, -1.11]* | -5.36 [-7.35, -3.4]* | 2.41 [0.5, 4.25]* | -4.84 [-6.81, -2.88]* | -2.43 [-4.47, -0.29]* |
| R 466 | R 1099 | R alveus | 326.61 ± 95.12 | 15 | 424.47 ± 62.45 | 15 | 1571.15 ± 216.91 | 12 | 1842.55 ± 369.51 | 10 | 3.62 [1.63, 5.54]* | 0.38 [-1.49, 2.25] | -0.16 [-3.03, 2.81] | -0.31 [-2.03, 1.42] | -4.04 [-5.83, -2.19]* | -4.88 [-6.85, -2.91]* | 3.7 [1.89, 5.54]* | -4.58 [-6.56, -2.59]* | -0.86 [-2.93, 1.16] |
| L 466 | L 1099 | L alveus | 42.32 ± 7.57 | 15 | 99.47 ± 27.97 | 15 | 225.63 ± 41.40 | 12 | 391.64 ± 71.87 | 10 | 4.11 [2.1, 6.07]* | 1.42 [-0.46, 3.33] | 1.93 [-0.98, 4.89] | -1.33 [-3.08, 0.43] | -4.51 [-6.33, -2.58]* | -8.51 [-10.45, -6.56]* | 3.18 [1.28, 5.02]* | -7.15 [-8.99, -5.11]* | -4 [-6.1, -1.96]* |
| R 603 | R 1099 | R fimbria | 356.68 ± 142.39 | 15 | 363.15 ± 89.01 | 15 | 2032.74 ± 475.59 | 12 | 2165.30 ± 620.41 | 10 | 4.5 [2.53, 6.44]* | 0.09 [-1.78, 1.98] | -0.32 [-3.21, 2.59] | -0.02 [-1.77, 1.72] | -4.91 [-6.71, -3.05]* | -5.31 [-7.22, -3.29]* | 4.89 [2.99, 6.64]* | -5.29 [-7.16, -3.31]* | -0.41 [-2.4, 1.69] |
| L 603 | L 1099 | L fimbria | 73.25 ± 26.47 | 15 | 99.33 ± 59.27 | 15 | 228.80 ± 70.79 | 12 | 357.27 ± 95.99 | 10 | 1.89 [-0.01, 3.82] | 0.42 [-1.43, 2.3] | 0.76 [-2.19, 3.68] | | | | | | |
| R 737 | R 1099 | R postcommissural fornix | 24.29 ± 11.52 | 15 | 18.10 ± 5.48 | 14 | 143.57 ± 51.11 | 12 | 118.40 ± 26.93 | 10 | 4.68 [2.7, 6.59]* | -0.19 [-2.1, 1.7] | -1.42 [-4.35, 1.47] | 0.27 [-1.48, 2.06] | -5.08 [-6.88, -3.21]* | -4.11 [-6.11, -2.13]* | 5.35 [3.53, 7.34]* | -4.38 [-6.41, -2.4]* | 0.98 [-1.17, 2.99] |
| L 737 | L 1099 | L postcommissural fornix | 12.99 ± 7.05 | 15 | 14.58 ± 5.18 | 14 | 32.46 ± 6.00 | 12 | 42.60 ± 8.47 | 10 | 1.31 [-0.71, 3.27] | 0.22 [-1.67, 2.09] | -0.03 [-2.95, 2.89] | | | | | | |
| R 436 | R 737 | R columns of the fornix | 48.58 ± 23.04 | 15 | 36.19 ± 10.95 | 14 | 287.14 ± 102.22 | 12 | 236.79 ± 53.87 | 10 | 4.71 [2.74, 6.61]* | -0.17 [-2, 1.77] | -1.46 [-4.37, 1.51] | 0.25 [-1.55, 1.98] | -5.12 [-6.89, -3.15]* | -4.12 [-6.2, -2.27]* | 5.35 [3.41, 7.12]* | -4.36 [-6.44, -2.46]* | 0.99 [-1.07, 3.02] |
| L 436 | L 737 | L columns of the fornix | 25.98 ± 14.09 | 15 | 29.16 ± 10.37 | 14 | 64.92 ± 11.99 | 12 | 86.63 ± 16.91 | 10 | 1.29 [-0.65, 3.23] | 0.21 [-1.69, 2.1] | 0.04 [-2.9, 2.94] | | | | | | |
| R 618 | R 1099 | R hippocampal commissures | 182.91 ± 42.44 | 15 | 162.04 ± 51.79 | 15 | 502.90 ± 123.86 | 12 | 830.63 ± 103.58 | 10 | 1.53 [-0.45, 3.52] | -0.05 [-1.96, 1.84] | 1.26 [-1.64, 4.16] | | | | | | |
| L 618 | L 1099 | L hippocampal commissures | 46.11 ± 16.09 | 15 | 59.70 ± 15.35 | 15 | 195.81 ± 48.76 | 12 | 339.58 ± 111.27 | 10 | 3.04 [1.12, 4.97]* | 0.37 [-1.46, 2.21] | 2.15 [-0.75, 5.08] | -0.3 [-2.08, 1.41] | -3.45 [-5.24, -1.56]* | -6.6 [-8.53, -4.6]* | 3.15 [1.37, 5.02]* | -6.31 [-8.16, -4.27]* | -3.16 [-5.16, -1.08]* |
| R 443 | R 618 | R dorsal hippocampal commissure | 359.91 ± 85.84 | 15 | 325.07 ± 103.37 | 15 | 1009.13 ± 246.88 | 12 | 1661.26 ± 207.17 | 10 | 1.59 [-0.41, 3.55] | -0.03 [-1.93, 1.83] | 1.27 [-1.64, 4.17] | | | | | | |
| L 443 | L 618 | L dorsal hippocampal commissure | 92.15 ± 32.21 | 15 | 118.47 ± 30.95 | 15 | 391.90 ± 97.45 | 12 | 679.15 ± 222.55 | 10 | 3.06 [1.09, 5.03]* | 0.37 [-1.48, 2.22] | 2.16 [-0.73, 5.09] | -0.28 [-2.07, 1.41] | -3.46 [-5.27, -1.58]* | -6.62 [-8.59, -4.78]* | 3.16 [1.32, 5.01]* | -6.34 [-8.33, -4.42]* | -3.18 [-5.13, -1.09]* |
| R 301 | R 768 | R stria terminalis | 68.75 ± 21.35 | 15 | 50.88 ± 20.47 | 15 | 284.35 ± 162.65 | 12 | 300.84 ± 130.15 | 10 | 2.92 [0.95, 4.84]* | -0.19 [-2.07, 1.7] | -0.17 [-3.13, 2.71] | 0.28 [-1.52, 1.99] | -3.33 [-5.21, -1.53]* | -3.61 [-5.64, -1.66]* | 3.6 [1.67, 5.39]* | -3.86 [-5.8, -1.85]* | -0.28 [-2.33, 1.81] |
| L 301 | L 768 | L stria terminalis | 107.60 ± 58.08 | 15 | 21.81 ± 14.12 | 15 | 61.58 ± 29.67 | 12 | 120.46 ± 36.42 | 10 | -0.64 [-2.62, 1.36] | -0.74 [-2.63, 1.16] | 0.69 [-2.24, 3.58] | | | | | | |
| R 484682528 | R 301 | R commissural branch of stria terminalis | 11.08 ± 8.12 | 14 | 37.22 ± 21.98 | 12 | 59.80 ± 20.95 | 12 | 76.82 ± 36.39 | 10 | 4.26 [2.3, 6.21]* | 2.52 [0.53, 4.5]* | -1.58 [-4.56, 1.41] | -2.45 [-4.29, -0.55]* | -4.66 [-6.55, -2.81]* | -6.24 [-8.26, -4.3]* | 2.21 [0.2, 4.13]* | -3.8 [-5.84, -1.65]* | -1.59 [-3.56, 0.56] |
| L 484682528 | L 301 | L commissural branch of stria terminalis | 219.71 ± 202.36 | 14 | 5.67 ± 4.01 | 13 | 67.28 ± 28.84 | 12 | 106.33 ± 39.64 | 9 | -0.85 [-2.85, 1.18] | -0.81 [-2.8, 1.15] | 0.41 [-2.66, 3.42] | | | | | | |
| R 824 | R 991 | R hypothalamus related | 33.22 ± 15.61 | 15 | 27.79 ± 6.72 | 15 | 43.15 ± 11.09 | 12 | 82.10 ± 28.26 | 10 | 0.09 [-1.88, 2.02] | -0.09 [-2, 1.77] | 0.68 [-2.32, 3.63] | | | | | | |
| L 824 | L 991 | L hypothalamus related | 19.37 ± 8.94 | 15 | 14.95 ± 3.21 | 14 | 31.66 ± 7.10 | 12 | 50.29 ± 18.98 | 10 | 0.4 [-1.59, 2.4] | -0.21 [-2.13, 1.67] | 0.61 [-2.35, 3.52] | | | | | | |
| R 54 | R 824 | R medial forebrain bundle | 160.47 ± 81.63 | 8 | 43.51 ± 20.58 | 10 | 235.56 ± 124.46 | 9 | 573.40 ± 276.38 | 7 | 0.36 [-1.93, 2.7] | -0.39 [-2.67, 1.89] | 2.29 [-1.04, 5.65] | | | | | | |
| L 54 | L 824 | L medial forebrain bundle | 87.09 ± 59.44 | 9 | 107.97 ± 69.07 | 10 | 144.28 ± 43.15 | 8 | 84.58 ± 25.11 | 8 | 0.74 [-1.6, 2.98] | 0.54 [-1.7, 2.79] | -1.68 [-5.01, 1.69] | | | | | | |
| R 349 | R 824 | R supraoptic commissures | 99.82 ± 47.49 | 14 | 84.73 ± 63.76 | 12 | 28.64 ± 17.10 | 12 | 51.04 ± 21.70 | 10 | -0.84 [-2.84, 1.15] | 0.01 [-1.98, 1.99] | -0.39 [-3.37, 2.63] | | | | | | |
| L 349 | L 824 | L supraoptic commissures | 50.45 ± 36.05 | 14 | 28.14 ± 15.77 | 12 | 33.28 ± 22.92 | 12 | 8.01 ± 5.34 | 10 | -0.49 [-2.47, 1.48] | -0.29 [-2.21, 1.68] | -0.8 [-3.81, 2.15] | | | | | | |
| R 46 | R 824 | R mammillary related | 20.62 ± 8.13 | 14 | 63.65 ± 19.68 | 15 | 33.87 ± 11.53 | 12 | 52.08 ± 11.87 | 10 | 0.51 [-1.46, 2.48] | 2.24 [0.33, 4.13]* | -1.95 [-4.89, 0.96] | -2.17 [-3.96, -0.41]* | -0.92 [-2.74, 1.05] | -1.84 [-3.91, 0.06] | -1.25 [-3.13, 0.57] | 0.32 [-1.63, 2.33] | -0.93 [-3.07, 1.02] |
| L 46 | L 824 | L mammillary related | 18.49 ± 10.50 | 14 | 15.97 ± 4.82 | 15 | 41.00 ± 9.79 | 12 | 46.52 ± 15.87 | 10 | 1.08 [-0.87, 3.05] | 0.01 [-1.88, 1.85] | -0.31 [-3.24, 2.54] | | | | | | |
| R 753 | R 46 | R principal mammillary tract | 166.83 ± 88.13 | 9 | 580.09 ± 166.03 | 9 | 164.50 ± 96.77 | 9 | 246.45 ± 83.95 | 7 | 0.16 [-2.14, 2.46] | 2.83 [0.55, 5.13]* | -2.79 [-6.16, 0.65] | -2.75 [-4.94, -0.56]* | -0.55 [-2.79, 1.63] | -1.23 [-3.55, 1.15] | -2.2 [-4.41, 0.03] | 1.52 [-0.81, 3.91] | -0.67 [-2.98, 1.75] |
| L 753 | L 46 | L principal mammillary tract | 74.67 ± 36.86 | 10 | 171.35 ± 65.76 | 9 | 258.27 ± 94.48 | 9 | 278.72 ± 136.47 | 8 | 2.63 [0.4, 4.85]* | 1.67 [-0.55, 3.89] | -1.95 [-5.23, 1.34] | -1.59 [-3.7, 0.51] | -3.05 [-5.27, -1.02]* | -3.39 [-5.68, -1.19]* | 1.44 [-0.7, 3.65] | -1.79 [-4.1, 0.46] | -0.34 [-2.71, 1.8] |
| R 690 | R 46 | R mammillothalamic tract | 18.60 ± 7.41 | 14 | 55.72 ± 36.17 | 14 | 45.87 ± 13.52 | 12 | 77.29 ± 19.33 | 10 | 1.33 [-0.64, 3.27] | 2.15 [0.26, 4.04]* | -1.06 [-3.99, 1.85] | -2.06 [-3.9, -0.34]* | -1.74 [-3.59, 0.11] | -3.47 [-5.42, -1.46]* | -0.35 [-2.25, 1.49] | -1.39 [-3.33, 0.6] | -1.74 [-3.74, 0.35] |
| L 690 | L 46 | L mammillothalamic tract | 31.97 ± 21.49 | 14 | 7.91 ± 3.26 | 14 | 45.85 ± 13.98 | 12 | 62.31 ± 14.13 | 10 | 0.3 [-1.71, 2.28] | -0.59 [-2.5, 1.32] | 0.51 [-2.43, 3.51] | | | | | | |
| R 681 | R 46 | R mammillotegmental tract | 58.53 ± 34.35 | 9 | 23.20 ± 19.90 | 10 | 142.86 ± 95.98 | 8 | 32.43 ± 14.68 | 6 | 1.57 [-0.76, 3.9] | -0.32 [-2.58, 1.93] | -1.76 [-5.29, 1.74] | | | | | | |
| L 681 | L 46 | L mammillotegmental tract | 72.63 ± 43.47 | 9 | 81.62 ± 38.40 | 10 | 37.08 ± 24.78 | 9 | 158.73 ± 99.79 | 7 | -0.31 [-2.6, 1.98] | 0.43 [-1.84, 2.64] | 0.84 [-2.56, 4.17] | | | | | | |
| R 1083 | R 824 | R epithalamus related | 59.71 ± 32.08 | 15 | 27.21 ± 10.86 | 15 | 70.30 ± 22.40 | 12 | 120.70 ± 50.88 | 10 | -0.03 [-1.98, 1.88] | -0.46 [-2.35, 1.46] | 0.73 [-2.14, 3.62] | | | | | | |
| L 1083 | L 824 | L epithalamus related | 30.32 ± 14.68 | 15 | 16.47 ± 4.22 | 15 | 44.81 ± 15.84 | 12 | 83.95 ± 39.89 | 10 | 0.28 [-1.67, 2.2] | -0.38 [-2.22, 1.45] | 1.08 [-1.83, 4.05] | | | | | | |
| R 802 | R 1083 | R stria medullaris | 167.95 ± 105.45 | 15 | 83.99 ± 49.61 | 13 | 222.00 ± 77.80 | 12 | 394.23 ± 155.43 | 10 | 0.12 [-1.83, 2.08] | -0.37 [-2.27, 1.53] | 0.81 [-2.2, 3.74] | | | | | | |
| L 802 | L 1083 | L stria medullaris | 90.06 ± 63.96 | 15 | 52.67 ± 18.18 | 13 | 130.89 ± 53.44 | 12 | 272.04 ± 149.28 | 10 | 0.24 [-1.74, 2.16] | -0.3 [-2.24, 1.62] | 1.28 [-1.71, 4.26] | | | | | | |
| R 595 | R 1083 | R fasciculus retroflexus | 48.10 ± 27.68 | 14 | 21.16 ± 11.59 | 15 | 63.72 ± 27.60 | 12 | 34.49 ± 14.44 | 10 | 0.21 [-1.81, 2.19] | -0.4 [-2.3, 1.48] | -0.81 [-3.73, 2.14] | | | | | | |
| L 595 | L 1083 | L fasciculus retroflexus | 24.38 ± 17.63 | 14 | 11.13 ± 3.92 | 15 | 29.97 ± 14.24 | 12 | 37.38 ± 13.66 | 10 | 0.08 [-1.94, 2.08] | -0.41 [-2.3, 1.49] | 0.13 [-2.85, 3.03] | | | | | | |
| R 611 | R 1083 | R habenular commissure | 103.13 ± 58.96 | 13 | 67.05 ± 34.43 | 15 | 108.52 ± 30.46 | 11 | 241.58 ± 147.82 | 8 | -0.11 [-2.16, 1.93] | -0.19 [-2.14, 1.71] | 0.9 [-2.17, 3.97] | | | | | | |
| L 611 | L 1083 | L habenular commissure | 61.83 ± 42.92 | 13 | 40.92 ± 23.96 | 15 | 208.57 ± 126.02 | 11 | 91.46 ± 38.69 | 9 | 2.25 [0.23, 4.31]* | -0.15 [-2.05, 1.75] | -2.32 [-5.34, 0.69] | 0.24 [-1.56, 2.01] | -2.65 [-4.67, -0.77]* | -0.82 [-2.8, 1.29] | 2.88 [0.99, 4.75]* | -1.05 [-3.03, 1.01] | 1.83 [-0.3, 3.95] |

**Table S1**. **Results from parent and subregion level AT8+ immunoreactive aggregate mapping.** Mean AT8+ immunoreactive aggregates/$mm^2$ +/- standard error of the mean and sample size for each experimental group (n) for each sampled brain region. Parent ID indicates the hierarchical “parent” area for each region. Results from Bayesian statistical models (genotype, treatment, and brain region as fixed effects, animal as random effect) show the estimated relative change (normalized to regional WT-Sal) and 95% credible intervals (CI) in AT8+ levels due to interactions between brain region and 5XFAD genotype (genotype effect), PTZ treatment (treatment effect), and 5XFAD genotype and PTZ (Interaction). Estimated marginal means and 95% CI were used for group contrasts at each brain region, where credible genotype, treatment, or interaction effects were found. Separate models were performed for the isocortex, olfactory areas, hippocampal formation, cortical subplate, striatum, pallidum, thalamus, hypothalamus, midbrain, and fiber tracts. *=results are considered credible when the 95% CI does not cross 0.

| ID | Parent | Region | WT Sal | n | WT PTZ | n | 5X Sal | n | 5X PTZ | n | Genotype effect | Treatment effect | Interaction | WT Sal vs WT PTZ | WT Sal vs 5XFAD Sal | WT Sal vs 5XFAD PTZ | 5XFAD Sal vs WT PTZ | WT PTZ VS 5XFAD PT | 5XFAD Sal vs 5XFAD PTZ |
|---|---|---|---|---|---|---|---|---|---|---|---|---|---|---|---|---|---|---|---|
| R 315 | R 695 | R Isocortex | 5.04 ± 1.12 | 15 | 14.57 ± 3.57 | 16 | 6.16 ± 0.89 | 11 | 26.77 ± 4.49 | 12 | -0.02 [-2.03, 2.02] | 1.72 [-0.16, 3.61] | 2.18 [-0.62, 4.9] | | | | | | |
| L 315 | L 695 | L Isocortex | 4.95 ± 0.88 | 15 | 14.91 ± 3.41 | 16 | 6.65 ± 0.97 | 11 | 27.62 ± 5.00 | 12 | 0.11 [-1.86, 2.15] | 1.84 [0.01, 3.69]* | 2.19 [-0.62, 4.94] | -1.97 [-3.99, -0.06]* | -0.43 [-2.51, 1.66] | -4.83 [-7.02, -2.74]* | -1.57 [-3.69, 0.6] | -2.84 [-5, -0.67]* | -4.42 [-6.73, -2.14]* |
| R 500 | R 315 | R Somatomotor areas | 3.71 ± 1.00 | 15 | 18.48 ± 4.95 | 16 | 6.04 ± 0.87 | 11 | 37.68 ± 7.56 | 12 | 0.38 [-1.6, 2.37] | 3.81 [1.95, 5.66]* | 4.51 [1.71, 7.28]* | -3.95 [-5.92, -1.99]* | -0.69 [-2.74, 1.38] | -9.39 [-11.51, -7.23]* | -3.26 [-5.38, -1.08]* | -5.45 [-7.52, -3.25]* | -8.71 [-11.03, -6.44]* |
| L 500 | L 315 | L Somatomotor areas | 3.69 ± 0.69 | 15 | 17.85 ± 4.81 | 16 | 6.42 ± 0.95 | 11 | 33.58 ± 7.00 | 12 | 0.5 [-1.51, 2.52] | 3.67 [1.84, 5.54]* | 3.49 [0.67, 6.29]* | -3.8 [-5.71, -1.8]* | -0.8 [-2.84, 1.33] | -8.35 [-10.58, -6.28]* | -2.99 [-5.02, -0.7]* | -4.55 [-6.63, -2.43]* | -7.54 [-9.81, -5.19]* |
| R 985 | R 500 | R Primary motor area | 5.74 ± 1.30 | 13 | 28.86 ± 8.51 | 16 | 8.09 ± 1.35 | 11 | 63.27 ± 11.88 | 11 | 0.23 [-1.86, 2.28] | 3.93 [2.04, 5.87]* | 5.47 [2.66, 8.28]* | -4.07 [-6.02, -2]* | -0.54 [-2.62, 1.67] | -10.33 [-12.52, -8.08]* | -3.52 [-5.6, -1.31]* | -6.25 [-8.41, -4.09]* | -9.77 [-12.14, -7.48]* |
| L 985 | L 500 | L Primary motor area | 4.95 ± 0.85 | 15 | 27.45 ± 7.77 | 16 | 7.95 ± 1.28 | 11 | 56.97 ± 10.37 | 11 | 0.37 [-1.61, 2.41] | 4.38 [2.52, 6.3]* | 5.31 [2.45, 8.09]* | -4.52 [-6.59, -2.6]* | -0.69 [-2.72, 1.46] | -10.75 [-12.96, -8.55]* | -3.84 [-6, -1.71]* | -6.23 [-8.36, -4.07]* | -10.08 [-12.37, -7.79]* |
| R 993 | R 500 | R Secondary motor area | 8.40 ± 2.76 | 11 | 27.21 ± 6.25 | 16 | 11.40 ± 1.65 | 11 | 62.35 ± 11.15 | 11 | 0.14 [-1.92, 2.28] | 2.11 [0.19, 4.07]* | 3.73 [0.87, 6.54]* | -2.24 [-4.37, -0.28]* | -0.45 [-2.63, 1.64] | -6.67 [-8.92, -4.37]* | -1.8 [-3.95, 0.38] | -4.44 [-6.69, -2.36]* | -6.22 [-8.51, -3.87]* |
| L 993 | L 500 | L Secondary motor area | 7.66 ± 1.62 | 13 | 27.83 ± 7.77 | 14 | 13.26 ± 2.13 | 11 | 54.46 ± 11.00 | 11 | 0.48 [-1.56, 2.55] | 2.64 [0.71, 4.58]* | 2.53 [-0.32, 5.38] | -2.78 [-4.81, -0.74]* | -0.79 [-2.97, 1.3] | -6.35 [-8.49, -4.06]* | -1.99 [-4.19, 0.2] | -3.56 [-5.76, -1.33]* | -5.55 [-7.93, -3.23]* |
| R 453 | R 315 | R Somatosensory areas | 5.73 ± 1.41 | 15 | 20.09 ± 5.36 | 16 | 7.81 ± 1.09 | 11 | 33.35 ± 6.03 | 12 | 0.13 [-1.86, 2.19] | 2.34 [0.49, 4.19]* | 1.91 [-0.88, 4.64] | -2.46 [-4.42, -0.51]* | -0.44 [-2.53, 1.66] | -5.07 [-7.22, -2.85]* | -2.04 [-4.22, 0.1] | -2.6 [-4.75, -0.49]* | -4.63 [-6.87, -2.22]* |
| L 453 | L 315 | L Somatosensory areas | 5.48 ± 1.07 | 15 | 19.32 ± 4.66 | 16 | 7.83 ± 1.04 | 11 | 34.68 ± 7.16 | 11 | 0.19 [-1.84, 2.25] | 2.36 [0.48, 4.27]* | 2.32 [-0.53, 5.15] | -2.51 [-4.48, -0.52]* | -0.5 [-2.62, 1.62] | -5.56 [-7.78, -3.4]* | -2.01 [-4.15, 0.14] | -3.07 [-5.21, -0.92]* | -5.07 [-7.37, -2.66]* |
| R 322 | R 453 | R Primary somatosensory area | 8.57 ± 1.99 | 14 | 26.65 ± 7.19 | 16 | 10.42 ± 1.58 | 10 | 44.78 ± 8.12 | 11 | -0.06 [-2.09, 2.07] | 1.96 [0.12, 3.84]* | 1.88 [-1.01, 4.7] | -2.1 [-4.13, -0.18]* | -0.25 [-2.41, 1.88] | -4.46 [-6.61, -2.23]* | -1.84 [-3.98, 0.4] | -2.36 [-4.54, -0.24]* | -4.21 [-6.6, -1.89]* |
| L 322 | L 453 | L Primary somatosensory area | 7.20 ± 1.39 | 15 | 25.53 ± 6.13 | 16 | 10.42 ± 1.37 | 11 | 45.48 ± 8.94 | 11 | 0.21 [-1.8, 2.24] | 2.39 [0.55, 4.26]* | 2.27 [-0.54, 5.03] | -2.52 [-4.57, -0.63]* | -0.53 [-2.63, 1.57] | -5.57 [-7.81, -3.48]* | -2.01 [-4.12, 0.15] | -3.04 [-5.27, -0.94]* | -5.06 [-7.3, -2.68]* |
| R 353 | R 322 | R Primary somatosensory area, nose | 12.22 ± 3.12 | 10 | 28.02 ± 11.77 | 10 | 11.46 ± 3.12 | 8 | 46.69 ± 13.05 | 11 | -0.17 [-2.39, 2.04] | 1.02 [-1.08, 3.1] | 1.57 [-1.44, 4.6] | | | | | | |
| L 353 | L 322 | L Primary somatosensory area, nose | 12.72 ± 2.94 | 9 | 15.42 ± 8.08 | 8 | 12.32 ± 3.30 | 9 | 115.37 ± 76.99 | 10 | 0.04 [-2.14, 2.27] | 0.7 [-1.51, 2.9] | 7.43 [4.31, 10.55]* | -0.85 [-3.08, 1.54] | -0.34 [-2.69, 1.91] | -8.87 [-11.11, -6.45]* | -0.5 [-2.81, 1.95] | -8.02 [-10.38, -5.67]* | -8.51 [-10.89, -6.05]* |
| R 329 | R 322 | R Primary somatosensory area, barrel fiel | 15.56 ± 3.64 | 14 | 45.92 ± 12.98 | 16 | 19.13 ± 2.63 | 11 | 70.87 ± 13.14 | 11 | -0.01 [-2.05, 2.06] | 1.79 [-0.1, 3.71] | 1.31 [-1.47, 4.07] | | | | | | |
| L 329 | L 322 | L Primary somatosensory area, barrel fiel | 14.96 ± 2.83 | 14 | 42.41 ± 9.72 | 16 | 18.73 ± 2.21 | 11 | 70.64 ± 14.03 | 11 | 0 [-2.02, 2.02] | 1.67 [-0.17, 3.57] | 1.59 [-1.26, 4.38] | | | | | | |
| R 337 | R 322 | R Primary somatosensory area, lower limb | 13.30 ± 4.49 | 13 | 47.05 ± 18.71 | 12 | 14.62 ± 2.37 | 11 | 67.35 ± 13.58 | 11 | -0.14 [-2.16, 1.89] | 2.63 [0.67, 4.61]* | 1.12 [-1.74, 3.97] | -2.76 [-4.79, -0.72]* | -0.18 [-2.34, 1.94] | -4.32 [-6.57, -2.12]* | -2.6 [-4.8, -0.36]* | -1.54 [-3.76, 0.67] | -4.13 [-6.38, -1.76]* |
| L 337 | L 322 | L Primary somatosensory area, lower limb | 12.63 ± 2.71 | 13 | 35.55 ± 11.17 | 11 | 16.21 ± 2.84 | 11 | 68.79 ± 14.96 | 11 | 0.04 [-1.99, 2.14] | 1.68 [-0.31, 3.7] | 2.27 [-0.67, 5.12] | | | | | | |
| R 345 | R 322 | R Primary somatosensory area, mouth | 7.18 ± 1.45 | 13 | 33.31 ± 9.18 | 14 | 11.11 ± 1.94 | 10 | 66.08 ± 11.69 | 11 | 0.27 [-1.77, 2.36] | 3.24 [1.3, 5.22]* | 4.25 [1.4, 7.08]* | -3.38 [-5.4, -1.33]* | -0.58 [-2.71, 1.57] | -8.46 [-10.73, -6.36]* | -2.8 [-4.99, -0.58]* | -5.07 [-7.28, -2.93]* | -7.86 [-10.16, -5.5]* |
| L 345 | L 322 | L Primary somatosensory area, mouth | 6.53 ± 1.21 | 13 | 36.43 ± 11.13 | 13 | 9.95 ± 1.64 | 11 | 71.73 ± 14.97 | 10 | 0.3 [-1.75, 2.41] | 3.99 [2.1, 5.95]* | 5.2 [2.29, 8.02]* | -4.11 [-6.24, -2.16]* | -0.61 [-2.7, 1.51] | -10.18 [-12.33, -7.89]* | -3.54 [-5.71, -1.27]* | -6.04 [-8.29, -3.86]* | -9.57 [-12.01, -7.29]* |
| R 369 | R 322 | R Primary somatosensory area, upper limb | 9.66 ± 2.71 | 14 | 37.98 ± 12.85 | 14 | 13.75 ± 3.06 | 11 | 65.58 ± 12.62 | 11 | 0.18 [-1.81, 2.21] | 2.92 [1.04, 4.89]* | 2.22 [-0.63, 5.07] | -3.05 [-5.18, -1.13]* | -0.49 [-2.61, 1.61] | -6.01 [-8.22, -3.75]* | -2.56 [-4.76, -0.42]* | -2.95 [-5.25, -0.86]* | -5.52 [-7.85, -3.2]* |
| L 369 | L 322 | L Primary somatosensory area, upper limb | 8.48 ± 2.17 | 14 | 35.45 ± 12.31 | 14 | 13.05 ± 2.34 | 11 | 62.32 ± 12.52 | 11 | 0.3 [-1.73, 2.34] | 3.19 [1.31, 5.14]* | 2.41 [-0.47, 5.17] | -3.33 [-5.28, -1.3]* | -0.61 [-2.72, 1.51] | -6.59 [-8.81, -4.41]* | -2.72 [-4.84, -0.49]* | -3.27 [-5.45, -1.05]* | -5.98 [-8.4, -3.72]* |
| R 361 | R 322 | R Primary somatosensory area, trunk | 14.14 ± 3.24 | 14 | 45.63 ± 15.40 | 13 | 17.15 ± 2.49 | 11 | 71.97 ± 14.77 | 11 | -0.03 [-2.04, 2.02] | 2.04 [0.12, 4.01]* | 1.63 [-1.28, 4.44] | -2.16 [-4.23, -0.19]* | -0.26 [-2.46, 1.74] | -4.33 [-6.57, -2.15]* | -1.9 [-4.07, 0.34] | -2.17 [-4.48, -0.05]* | -4.05 [-6.33, -1.66]* |
| L 361 | L 322 | L Primary somatosensory area, trunk | 10.41 ± 2.03 | 13 | 41.51 ± 12.37 | 14 | 18.44 ± 3.96 | 11 | 72.25 ± 15.06 | 11 | 0.53 [-1.53, 2.6] | 2.99 [1.05, 4.97]* | 1.96 [-0.93, 4.81] | -3.12 [-5.14, -1.04]* | -0.86 [-2.88, 1.34] | -6.17 [-8.47, -4.08]* | -2.28 [-4.41, -0.04]* | -3.05 [-5.29, -0.94]* | -5.34 [-7.75, -3.11]* |
| R 182305 | R 322 | R Primary somatosensory area, unassigne | 14.46 ± 4.12 | 14 | 45.79 ± 22.24 | 12 | 17.82 ± 1.97 | 11 | 67.34 ± 15.21 | 11 | 0 [-2.05, 2.06] | 2.26 [0.29, 4.23]* | 0.94 [-1.96, 3.76] | -2.41 [-4.46, -0.36]* | -0.3 [-2.29, 1.92] | -3.88 [-6.08, -1.7]* | -2.09 [-4.23, 0.17] | -1.5 [-3.68, 0.7] | -3.58 [-5.8, -1.14]* |
| L 182305 | L 322 | L Primary somatosensory area, unassigne | 14.27 ± 4.33 | 14 | 39.66 ± 14.25 | 12 | 18.93 ± 2.60 | 11 | 66.71 ± 16.11 | 11 | 0.09 [-1.95, 2.16] | 1.54 [-0.43, 3.52] | 1.59 [-1.32, 4.48] | | | | | | |
| R 378 | R 453 | R Supplemental somatosensory area | 9.47 ± 2.42 | 12 | 32.43 ± 8.57 | 16 | 12.91 ± 1.91 | 10 | 59.62 ± 10.69 | 12 | 0.07 [-1.98, 2.21] | 2.28 [0.39, 4.25]* | 2.53 [-0.4, 5.31] | -2.41 [-4.46, -0.37]* | -0.38 [-2.54, 1.79] | -5.57 [-7.77, -3.35]* | -2.04 [-4.25, 0.15] | -3.15 [-5.3, -0.99]* | -5.2 [-7.46, -2.76]* |
| L 378 | L 453 | L Supplemental somatosensory area | 9.73 ± 1.87 | 14 | 32.44 ± 8.20 | 16 | 12.62 ± 1.78 | 11 | 67.50 ± 13.81 | 10 | 0.06 [-1.98, 2.16] | 2.17 [0.33, 4.05]* | 2.83 [-0.02, 5.62] | -2.31 [-4.3, -0.33]* | -0.36 [-2.56, 1.72] | -5.75 [-8.06, -3.61]* | -1.95 [-4.12, 0.26] | -3.44 [-5.68, -1.27]* | -5.39 [-7.76, -3.03]* |
| R 1057 | R 315 | R Gustatory areas | 7.49 ± 2.30 | 15 | 24.01 ± 5.81 | 14 | 5.31 ± 1.04 | 11 | 53.26 ± 9.53 | 12 | -0.52 [-2.5, 1.47] | 1.8 [-0.04, 3.69] | 4.41 [1.56, 7.15]* | -1.92 [-3.88, 0.07] | 0.22 [-1.86, 2.26] | -6.37 [-8.53, -4.21]* | -2.15 [-4.29, 0.07] | -4.45 [-6.59, -2.32]* | -6.59 [-8.85, -4.23]* |
| L 1057 | L 315 | L Gustatory areas | 7.49 ± 1.86 | 15 | 29.04 ± 8.91 | 14 | 4.95 ± 1.10 | 11 | 61.45 ± 12.76 | 12 | -0.58 [-2.58, 1.44] | 2.51 [0.65, 4.4]* | 4.83 [2.04, 7.64]* | -2.64 [-4.69, -0.71]* | 0.27 [-1.81, 2.41] | -7.46 [-9.66, -5.32]* | -2.93 [-5.12, -0.76]* | -4.81 [-6.91, -2.62]* | -7.73 [-10.02, -5.44]* |
| R 677 | R 315 | R Visceral area | 4.79 ± 0.99 | 14 | 20.61 ± 6.73 | 14 | 6.58 ± 1.11 | 10 | 38.16 ± 9.44 | 12 | 0.06 [-1.98, 2.13] | 3.29 [1.39, 5.26]* | 3.17 [0.31, 6.03]* | -3.43 [-5.52, -1.46]* | -0.36 [-2.59, 1.68] | -7.2 [-9.39, -5.02]* | -3.07 [-5.3, -0.83]* | -3.77 [-6.03, -1.7]* | -6.86 [-9.11, -4.39]* |
| L 677 | L 315 | L Visceral area | 5.66 ± 1.40 | 14 | 25.20 ± 7.86 | 14 | 6.18 ± 1.26 | 11 | 54.27 ± 10.63 | 10 | -0.15 [-2.16, 1.93] | 3.45 [1.55, 5.37]* | 4.41 [1.52, 7.24]* | -3.59 [-5.68, -1.64]* | -0.16 [-2.3, 1.88] | -8.41 [-10.61, -6.19]* | -3.43 [-5.57, -1.21]* | -4.82 [-7.07, -2.67]* | -8.24 [-10.6, -5.91]* |
| R 247 | R 315 | R Auditory areas | 9.40 ± 1.98 | 14 | 20.44 ± 5.19 | 16 | 11.01 ± 1.91 | 11 | 39.70 ± 7.94 | 11 | 0.01 [-1.98, 2.05] | 1.09 [-0.77, 3.02] | 1.71 [-1.1, 4.46] | | | | | | |
| L 247 | L 315 | L Auditory areas | 8.55 ± 1.86 | 14 | 22.16 ± 5.06 | 15 | 12.46 ± 2.16 | 11 | 49.35 ± 11.63 | 12 | 0.3 [-1.76, 2.35] | 1.38 [-0.53, 3.31] | 2.74 [-0.11, 5.54] | | | | | | |
| R 1011 | R 247 | R Dorsal auditory area | 16.71 ± 4.34 | 12 | 32.89 ± 9.09 | 16 | 17.00 ± 3.10 | 11 | 62.03 ± 13.74 | 10 | -0.15 [-2.2, 1.93] | 0.89 [-1.04, 2.85] | 1.91 [-1.02, 4.79] | | | | | | |
| L 1011 | L 247 | L Dorsal auditory area | 11.87 ± 3.46 | 12 | 38.01 ± 9.52 | 15 | 17.05 ± 2.61 | 11 | 77.59 ± 17.50 | 12 | 0.27 [-1.81, 2.38] | 1.97 [0.04, 3.94]* | 2.93 [0.01, 5.73]* | -2.11 [-4.21, -0.15]* | -0.58 [-2.76, 1.51] | -5.86 [-8.16, -3.69]* | -1.53 [-3.8, 0.6] | -3.75 [-5.87, -1.54]* | -5.27 [-7.59, -2.95]* |
| R 1002 | R 247 | R Primary auditory area | 16.72 ± 2.80 | 13 | 35.50 ± 9.00 | 12 | 18.33 ± 3.79 | 11 | 56.58 ± 20.14 | 9 | -0.06 [-2.1, 2.03] | 1 [-0.93, 3.01] | 1.44 [-1.49, 4.31] | | | | | | |
| L 1002 | L 247 | L Primary auditory area | 14.94 ± 3.26 | 13 | 36.62 ± 7.95 | 14 | 22.03 ± 4.04 | 11 | 82.65 ± 22.99 | 11 | 0.34 [-1.7, 2.42] | 1.07 [-0.85, 3.03] | 2.68 [-0.17, 5.51] | | | | | | |
| R 1027 | R 247 | R Posterior auditory area | 14.02 ± 4.30 | 6 | 30.85 ± 16.99 | 7 | 13.43 ± 4.18 | 10 | 95.27 ± 21.89 | 6 | -0.33 [-2.67, 2.08] | 0.27 [-2.15, 2.68] | 3.92 [0.52, 7.28]* | -0.42 [-2.87, 2.18] | 0.01 [-2.48, 2.39] | -4.55 [-7.27, -1.9]* | -0.43 [-2.88, 1.88] | -4.15 [-6.91, -1.7]* | -4.56 [-7.16, -2.06]* |
| L 1027 | L 247 | L Posterior auditory area | 18.86 ± 4.26 | 7 | 39.66 ± 16.56 | 7 | 17.83 ± 7.22 | 8 | 86.33 ± 28.91 | 7 | -0.05 [-2.37, 2.32] | -1.16 [-3.49, 1.26] | 3.67 [0.27, 6.99]* | 1.03 [-1.52, 3.4] | -0.27 [-2.56, 2.31] | -3.14 [-5.67, -0.59]* | 1.27 [-1.06, 3.86] | -4.19 [-6.68, -1.62]* | -2.89 [-5.44, -0.3]* |
| R 1018 | R 247 | R Ventral auditory area | 11.28 ± 3.13 | 14 | 28.60 ± 7.11 | 16 | 15.39 ± 2.52 | 11 | 50.72 ± 13.45 | 10 | 0.2 [-1.87, 2.22] | 1.44 [-0.45, 3.36] | 1.79 [-0.98, 4.64] | | | | | | |
| L 1018 | L 247 | L Ventral auditory area | 11.47 ± 2.65 | 14 | 32.13 ± 7.77 | 15 | 16.43 ± 3.09 | 11 | 63.53 ± 15.22 | 12 | 0.26 [-1.73, 2.36] | 1.56 [-0.32, 3.48] | 2.33 [-0.48, 5.07] | | | | | | |
| R 669 | R 315 | R Visual areas | 15.86 ± 3.60 | 14 | 19.17 ± 5.72 | 16 | 13.57 ± 3.11 | 11 | 44.94 ± 8.20 | 11 | -0.32 [-2.36, 1.74] | 0.11 [-1.76, 1.98] | 1.63 [-1.27, 4.44] | | | | | | |
| L 669 | L 315 | L Visual areas | 15.15 ± 3.23 | 14 | 23.35 ± 6.16 | 15 | 17.20 ± 3.77 | 11 | 51.66 ± 10.33 | 10 | -0.04 [-2.06, 2] | 0.3 [-1.57, 2.23] | 1.59 [-1.32, 4.4] | | | | | | |
| R 402 | R 669 | R Anterolateral visual area | 23.40 ± 8.47 | 7 | 37.45 ± 13.13 | 8 | 23.00 ± 7.31 | 9 | 87.20 ± 17.94 | 6 | -0.2 [-2.51, 2.13] | 0 [-2.29, 2.33] | 1.16 [-2.2, 4.45] | | | | | | |
| L 402 | L 669 | L Anterolateral visual area | 24.28 ± 6.23 | 8 | 43.41 ± 12.87 | 8 | 20.47 ± 9.24 | 8 | 103.15 ± 21.45 | 8 | -0.15 [-2.42, 2.14] | -0.89 [-3.15, 1.37] | 3.47 [0.24, 6.65]* | 0.75 [-1.63, 3.12] | -0.16 [-2.57, 2.19] | -3.14 [-5.56, -0.66]* | 0.9 [-1.51, 3.39] | -3.87 [-6.29, -1.49]* | -2.99 [-5.42, -0.36]* |
| R 394 | R 669 | R Anteromedial visual area | 16.65 ± 5.36 | 12 | 30.98 ± 8.55 | 15 | 17.60 ± 2.26 | 11 | 65.10 ± 10.90 | 11 | -0.09 [-2.15, 1.99] | 0.69 [-1.23, 2.6] | 1.91 [-0.96, 4.73] | | | | | | |
| L 394 | L 669 | L Anteromedial visual area | 15.82 ± 3.33 | 13 | 41.29 ± 10.53 | 14 | 24.33 ± 5.35 | 11 | 64.18 ± 13.72 | 10 | 0.39 [-1.64, 2.49] | 1.6 [-0.3, 3.55] | 0.53 [-2.39, 3.37] | | | | | | |
| R 385 | R 669 | R Primary visual area | 34.16 ± 8.77 | 11 | 47.47 ± 14.41 | 10 | 25.30 ± 8.76 | 9 | 88.51 ± 17.72 | 7 | -0.16 [-2.35, 2.06] | -0.67 [-2.75, 1.43] | 1.01 [-2.14, 4.16] | | | | | | |
| L 385 | L 669 | L Primary visual area | 41.30 ± 7.94 | 11 | 48.38 ± 14.51 | 11 | 34.11 ± 9.21 | 9 | 87.50 ± 19.41 | 9 | -0.26 [-2.38, 1.89] | -0.55 [-2.56, 1.48] | 1.62 [-1.37, 4.56] | | | | | | |
| R 533 | R 669 | R posteromedial visual area | 21.31 ± 6.70 | 10 | 47.17 ± 14.83 | 8 | 15.39 ± 4.15 | 10 | 60.61 ± 11.41 | 7 | -0.24 [-2.39, 1.93] | -0.51 [-2.67, 1.69] | 1.2 [-1.97, 4.33] | | | | | | |
| L 533 | L 669 | L posteromedial visual area | 16.93 ± 3.76 | 10 | 36.99 ± 11.74 | 10 | 22.56 ± 6.05 | 10 | 62.11 ± 13.03 | 9 | 0.36 [-1.77, 2.53] | 0.14 [-1.93, 2.27] | 1.86 [-1.18, 4.85] | | | | | | |
| R 31 | R 315 | R Anterior cingulate area | 9.34 ± 2.83 | 15 | 20.34 ± 3.87 | 16 | 11.85 ± 2.07 | 11 | 41.89 ± 5.69 | 12 | 0.04 [-1.96, 2.07] | 1.02 [-0.82, 2.9] | 1.99 [-0.77, 4.76] | | | | | | |
| L 31 | L 315 | L Anterior cingulate area | 8.45 ± 1.34 | 15 | 20.36 ± 3.87 | 15 | 12.22 ± 2.01 | 11 | 38.11 ± 5.63 | 12 | 0.2 [-1.75, 2.2] | 1.55 [-0.3, 3.42] | 1.32 [-1.47, 4.07] | | | | | | |
| R 39 | R 31 | R Anterior cingulate area, dorsal part | 15.65 ± 4.83 | 13 | 32.65 ± 7.35 | 15 | 19.02 ± 3.47 | 11 | 68.64 ± 10.38 | 12 | 0.04 [-1.99, 2.11] | 1.08 [-0.82, 3.03] | 1.88 [-0.99, 4.69] | | | | | | |
| L 39 | L 31 | L Anterior cingulate area, dorsal part | 13.99 ± 2.40 | 15 | 34.05 ± 7.14 | 15 | 18.60 ± 3.57 | 11 | 58.46 ± 8.97 | 12 | 0.09 [-1.92, 2.09] | 1.57 [-0.33, 3.48] | 1.07 [-1.72, 3.84] | | | | | | |
| R 48 | R 31 | R Anterior cingulate area, ventral part | 14.80 ± 4.61 | 15 | 30.02 ± 5.04 | 15 | 17.81 ± 3.18 | 11 | 61.48 ± 8.25 | 12 | -0.02 [-2.01, 2.02] | 0.82 [-1.02, 2.72] | 1.92 [-0.89, 4.69] | | | | | | |
| L 48 | L 31 | L Anterior cingulate area, ventral part | 11.41 ± 2.17 | 15 | 28.69 ± 4.83 | 14 | 19.41 ± 3.04 | 11 | 59.06 ± 8.76 | 12 | 0.46 [-1.55, 2.5] | 1.62 [-0.26, 3.58] | 1.65 [-1.19, 4.43] | | | | | | |
| R 972 | R 315 | R Prelimbic area | 6.52 ± 2.51 | 10 | 33.54 ± 11.10 | 10 | 23.03 ± 8.75 | 6 | 49.99 ± 12.92 | 8 | 2.34 [0, 4.7] | 3.64 [1.56, 5.75]* | -0.39 [-3.6, 2.78] | -3.77 [-5.91, -1.51]* | -2.65 [-5.1, -0.25]* | -6.29 [-8.66, -3.9]* | -1.12 [-3.57, 1.29] | -2.51 [-4.92, -0.21]* | -3.63 [-6.18, -0.94]* |
| L 972 | L 315 | L Prelimbic area | 15.87 ± 2.35 | 9 | 34.74 ± 14.44 | 8 | 15.72 ± 8.68 | 6 | 48.82 ± 17.46 | 7 | -0.35 [-2.72, 2.02] | 0.26 [-2, 2.51] | 0.36 [-2.94, 3.69] | | | | | | |
| R 95 | R 315 | R Agranular insular area | 4.11 ± 0.80 | 14 | 11.83 ± 3.38 | 16 | 4.27 ± 0.61 | 10 | 23.24 ± 3.91 | 12 | -0.2 [-2.27, 1.89] | 1.78 [-0.1, 3.73] | 2.7 [-0.15, 5.51] | | | | | | |
| L 95 | L 315 | L Agranular insular area | 3.64 ± 0.52 | 15 | 12.75 ± 3.56 | 16 | 4.46 ± 0.81 | 11 | 26.90 ± 4.31 | 11 | -0.01 [-2, 1.98] | 2.34 [0.51, 4.19]* | 3.23 [0.41, 5.98]* | -2.47 [-4.42, -0.49]* | -0.29 [-2.41, 1.73] | -6.25 [-8.51, -4.11]* | -2.18 [-4.29, 0.01] | -3.77 [-5.84, -1.49]* | -5.96 [-8.24, -3.57]* |
| R 104 | R 95 | R Agranular insular area, dorsal part | 6.94 ± 1.59 | 11 | 20.72 ± 5.96 | 15 | 6.04 ± 0.97 | 10 | 38.58 ± 7.27 | 12 | -0.38 [-2.48, 1.74] | 1.69 [-0.26, 3.68] | 2.86 [-0.03, 5.72] | | | | | | |
| L 104 | L 95 | L Agranular insular area, dorsal part | 5.99 ± 0.62 | 14 | 19.66 ± 5.90 | 16 | 6.27 ± 1.28 | 11 | 43.61 ± 7.90 | 10 | -0.13 [-2.14, 1.95] | 2.19 [0.32, 4.11]* | 3.36 [0.45, 6.17]* | -2.31 [-4.26, -0.31]* | -0.17 [-2.3, 1.86] | -6.1 [-8.42, -3.92]* | -2.16 [-4.24, 0.05] | -3.79 [-5.97, -1.64]* | -5.94 [-8.16, -3.43]* |
| R 111 | R 95 | R Agranular insular area, posterior part | 5.02 ± 1.12 | 13 | 13.42 ± 4.28 | 13 | 5.43 ± 1.20 | 10 | 30.83 ± 5.64 | 12 | -0.14 [-2.2, 1.94] | 1.68 [-0.27, 3.64] | 3.25 [0.36, 6.09]* | -1.81 [-3.86, 0.24] | -0.17 [-2.27, 2.04] | -5.48 [-7.72, -3.33]* | -1.65 [-3.81, 0.68] | -3.66 [-5.89, -1.56]* | -5.31 [-7.66, -2.97]* |
| L 111 | L 95 | L Agranular insular area, posterior part | 3.64 ± 0.65 | 15 | 13.43 ± 3.77 | 14 | 5.34 ± 1.00 | 11 | 35.13 ± 5.96 | 11 | 0.24 [-1.71, 2.3] | 2.7 [0.83, 4.62]* | 4.88 [2, 7.69]* | -2.82 [-4.86, -0.92]* | -0.54 [-2.65, 1.5] | -8.51 [-10.5, -6.15]* | -2.28 [-4.56, -0.11]* | -5.67 [-7.72, -3.42]* | -7.97 [-10.26, -5.58]* |
| R 119 | R 95 | R Agranular insular area, ventral part | 9.07 ± 1.89 | 14 | 22.11 ± 5.87 | 15 | 7.12 ± 1.78 | 10 | 53.62 ± 14.97 | 12 | -0.46 [-2.48, 1.58] | 1.17 [-0.71, 3.06] | 3.84 [1.02, 6.62]* | -1.3 [-3.27, 0.7] | 0.15 [-2.08, 2.2] | -5.24 [-7.36, -3.04]* | -1.45 [-3.63, 0.81] | -3.95 [-6.04, -1.71]* | -5.38 [-7.62, -2.91]* |
| L 119 | L 95 | L Agranular insular area, ventral part | 9.33 ± 1.29 | 14 | 21.88 ± 6.38 | 16 | 9.48 ± 2.44 | 11 | 49.42 ± 7.51 | 10 | -0.15 [-2.19, 1.9] | 1.26 [-0.59, 3.15] | 2.32 [-0.59, 5.12] | | | | | | |
| R 254 | R 315 | R Retrosplenial area | 8.07 ± 2.09 | 14 | 18.72 ± 4.35 | 16 | 8.69 ± 1.53 | 11 | 33.62 ± 6.24 | 11 | -0.16 [-2.17, 1.87] | 1.15 [-0.69, 3.06] | 2.03 [-0.78, 4.81] | | | | | | |
| L 254 | L 315 | L Retrosplenial area | 8.94 ± 1.54 | 14 | 18.96 ± 4.67 | 16 | 10.39 ± 2.21 | 11 | 31.38 ± 6.32 | 12 | 0 [-2.04, 2.04] | 1.02 [-0.87, 2.88] | 1.12 [-1.67, 3.93] | | | | | | |
| R 894 | R 254 | R Retrosplenial area, lateral agranular part | 14.96 ± 3.04 | 14 | 34.34 ± 9.33 | 16 | 16.52 ± 2.28 | 11 | 61.34 ± 10.97 | 11 | -0.1 [-2.11, 1.97] | 1.18 [-0.69, 3.09] | 1.94 [-0.93, 4.68] | | | | | | |
| L 894 | L 254 | L Retrosplenial area, lateral agranular part | 15.69 ± 2.53 | 13 | 36.33 ± 8.78 | 16 | 17.56 ± 2.99 | 11 | 58.68 ± 10.51 | 12 | -0.01 [-2.05, 2.1] | 1.26 [-0.56, 3.18] | 1.15 [-1.64, 3.95] | | | | | | |
| R 879 | R 254 | R Retrosplenial area, dorsal part | 13.43 ± 3.44 | 15 | 33.68 ± 8.98 | 16 | 17.39 ± 3.20 | 10 | 69.85 ± 11.82 | 11 | -0.04 [-2.04, 2.08] | 1.34 [-0.5, 3.22] | 2.18 [-0.67, 4.99] | | | | | | |
| L 879 | L 254 | L Retrosplenial area, dorsal part | 14.89 ± 2.66 | 14 | 34.03 ± 8.70 | 16 | 20.55 ± 4.50 | 11 | 62.34 ± 14.08 | 11 | 0.22 [-1.81, 2.25] | 1.2 [-0.67, 3.11] | 1.13 [-1.74, 3.9] | | | | | | |
| R 886 | R 254 | R Retrosplenial area, ventral part | 11.35 ± 2.91 | 15 | 23.47 ± 5.05 | 16 | 10.53 ± 2.29 | 11 | 40.57 ± 7.16 | 12 | -0.33 [-2.35, 1.7] | 0.89 [-0.94, 2.74] | 1.57 [-1.25, 4.28] | | | | | | |
| L 886 | L 254 | L Retrosplenial area, ventral part | 12.76 ± 2.38 | 14 | 22.66 ± 5.51 | 16 | 12.31 ± 3.00 | 11 | 36.93 ± 7.36 | 12 | -0.22 [-2.29, 1.81] | 0.67 [-1.19, 2.61] | 1.06 [-1.75, 3.9] | | | | | | |
| R 22 | R 315 | R Posterior parietal association areas | 11.53 ± 2.78 | 15 | 27.38 ± 7.29 | 16 | 14.09 ± 2.28 | 11 | 48.52 ± 7.76 | 11 | -0.02 [-2.01, 2.04] | 1.21 [-0.64, 3.08] | 1.54 [-1.22, 4.27] | | | | | | |
| L 22 | L 315 | L Posterior parietal association areas | 13.49 ± 2.82 | 14 | 30.90 ± 7.12 | 16 | 18.35 ± 3.07 | 11 | 49.98 ± 9.58 | 12 | 0.18 [-1.84, 2.22] | 1.19 [-0.67, 3.08] | 0.96 [-1.89, 3.72] | | | | | | |
| R 312782 | R 22 | R Anterior area | 14.76 ± 3.79 | 15 | 39.69 ± 11.04 | 16 | 17.06 ± 3.32 | 11 | 65.65 ± 11.15 | 11 | -0.09 [-2.04, 1.95] | 1.52 [-0.33, 3.4] | 1.54 [-1.31, 4.31] | | | | | | |
| L 312782 | L 22 | L Anterior area | 17.74 ± 4.47 | 15 | 47.39 ± 11.73 | 16 | 24.59 ± 5.71 | 11 | 70.61 ± 15.47 | 11 | 0.14 [-1.83, 2.19] | 1.51 [-0.31, 3.41] | 0.87 [-1.97, 3.66] | | | | | | |
| R 417 | R 22 | R Rostrolateral visual area | 21.82 ± 5.89 | 14 | 47.23 ± 11.78 | 16 | 24.51 ± 5.11 | 11 | 85.05 ± 13.90 | 11 | -0.05 [-2.09, 1.99] | 1.06 [-0.79, 2.93] | 1.47 [-1.38, 4.22] | | | | | | |
| L 417 | L 22 | L Rostrolateral visual area | 20.64 ± 5.79 | 14 | 47.69 ± 9.88 | 15 | 33.77 ± 6.07 | 11 | 98.49 ± 16.90 | 10 | 0.48 [-1.55, 2.53] | 1.09 [-0.77, 3.02] | 1.65 [-1.22, 4.42] | | | | | | |
| R 541 | R 315 | R Temporal association areas | 8.46 ± 2.76 | 13 | 20.62 ± 6.01 | 13 | 8.58 ± 1.88 | 11 | 44.76 ± 12.66 | 9 | -0.15 [-2.17, 1.97] | 1.45 [-0.49, 3.46] | 2.94 [-0.02, 5.88] | | | | | | |
| L 541 | L 315 | L Temporal association areas | 8.36 ± 1.77 | 13 | 20.77 ± 5.81 | 16 | 12.16 ± 2.09 | 11 | 45.41 ± 10.25 | 12 | 0.29 [-1.74, 2.37] | 1.39 [-0.52, 3.33] | 2.38 [-0.45, 5.19] | | | | | | |
| R 922 | R 315 | R Perirhinal area | 3.17 ± 1.00 | 14 | 8.07 ± 2.25 | 14 | 4.22 ± 1.46 | 11 | 26.00 ± 4.61 | 9 | 0.16 [-1.86, 2.21] | 1.7 [-0.16, 3.64] | 4.43 [1.54, 7.29]* | -1.82 [-3.93, 0.1] | -0.47 [-2.64, 1.54] | -6.97 [-9.28, -4.78]* | -1.35 [-3.49, 0.9] | -5.14 [-7.43, -2.91]* | -6.5 [-8.99, -4.25]* |
| L 922 | L 315 | L Perirhinal area | 1.61 ± 0.68 | 14 | 8.17 ± 2.23 | 16 | 3.21 ± 1.16 | 11 | 16.66 ± 6.16 | 12 | 0.82 [-1.16, 2.9] | 3.98 [2.12, 5.86]* | 4.19 [1.41, 6.94]* | -4.11 [-6.15, -2.19]* | -1.12 [-3.2, 1.04] | -9.68 [-11.88, -7.51]* | -3 [-5.08, -0.77]* | -5.56 [-7.68, -3.41]* | -8.55 [-10.92, -6.31]* |
| R 895 | R 315 | R Ectorhinal area | 7.15 ± 1.98 | 14 | 15.48 ± 4.02 | 14 | 6.91 ± 2.11 | 11 | 36.78 ± 8.29 | 9 | -0.2 [-2.25, 1.87] | 1.31 [-0.61, 3.23] | 2.1 [-0.79, 4.97] | | | | | | |
| L 895 | L 315 | L Ectorhinal area | 6.68 ± 1.54 | 14 | 17.96 ± 5.05 | 16 | 7.18 ± 1.77 | 11 | 30.29 ± 6.83 | 12 | -0.1 [-2.08, 1.96] | 1.59 [-0.25, 3.51] | 1.67 [-1.17, 4.41] | | | | | | |
| R 698 | R 695 | R Olfactory areas | 7.98 ± 2.04 | 14 | 17.33 ± 5.03 | 15 | 7.27 ± 1.29 | 12 | 38.37 ± 7.38 | 12 | -0.42 [-2.48, 1.64] | 0.9 [-1.05, 2.87] | 2.68 [-0.28, 5.62] | | | | | | |
| L 698 | L 695 | L Olfactory areas | 7.85 ± 1.38 | 14 | 14.38 ± 3.93 | 15 | 7.86 ± 1.39 | 12 | 33.13 ± 5.98 | 12 | -0.33 [-2.44, 1.77] | 0.57 [-1.45, 2.62] | 2.34 [-0.66, 5.27] | | | | | | |
| L 159 | L 698 | L Anterior olfactory nucleus | 14.53 ± 3.78 | 8 | 21.33 ± 6.60 | 7 | 13.43 ± 4.80 | 6 | 132.21 ± 58.22 | 6 | -0.34 [-2.83, 2.22] | 0.04 [-2.4, 2.44] | 6.93 [3.31, 10.59]* | -0.25 [-2.63, 1.96] | 0 [-2.33, 2.48] | -7.43 [-9.82, -5.06]* | -0.25 [-2.63, 2.26] | -7.19 [-9.63, -4.8]* | -7.43 [-9.97, -4.91]* |
| R 589 | R 698 | R Taenia tecta | 8.64 ± 1.99 | 12 | 24.39 ± 9.70 | 10 | 7.23 ± 2.12 | 8 | 25.80 ± 4.75 | 10 | -0.53 [-2.83, 1.71] | 1.46 [-0.73, 3.64] | 0.41 [-2.82, 3.65] | | | | | | |
| L 589 | L 698 | L Taenia tecta | 7.57 ± 1.68 | 12 | 17.36 ± 6.61 | 10 | 6.25 ± 0.92 | 9 | 23.18 ± 3.90 | 10 | -0.56 [-2.75, 1.63] | 0.93 [-1.21, 3.1] | 1.06 [-2.07, 4.21] | | | | | | |
| R 597 | R 589 | R Taenia tecta, dorsal part | 17.39 ± 3.87 | 12 | 54.08 ± 20.90 | 9 | 14.53 ± 4.23 | 8 | 51.25 ± 9.82 | 10 | -0.53 [-2.76, 1.73] | 1.62 [-0.56, 3.82] | 0.23 [-2.96, 3.43] | | | | | | |
| L 597 | L 589 | L Taenia tecta, dorsal part | 15.30 ± 3.43 | 12 | 39.39 ± 13.96 | 9 | 11.96 ± 1.85 | 9 | 46.45 ± 8.09 | 10 | -0.6 [-2.78, 1.61] | 1.08 [-1.1, 3.27] | 0.92 [-2.24, 4.09] | | | | | | |
| R 814 | R 698 | R Dorsal peduncular area | 18.58 ± 6.56 | 14 | 49.87 ± 17.89 | 9 | 20.14 ± 4.32 | 8 | 64.39 ± 11.32 | 10 | -0.38 [-2.59, 1.8] | 1.15 [-0.99, 3.31] | 1.01 [-2.2, 4.21] | | | | | | |
| L 814 | L 698 | L Dorsal peduncular area | 16.60 ± 4.15 | 13 | 50.30 ± 17.35 | 9 | 27.19 ± 9.69 | 8 | 69.07 ± 10.31 | 10 | 0.21 [-2.06, 2.44] | 1.53 [-0.64, 3.75] | 0.8 [-2.5, 4.07] | | | | | | |
| R 961 | R 698 | R Piriform area | 21.15 ± 5.31 | 10 | 65.41 ± 18.77 | 10 | 21.92 ± 3.36 | 8 | 94.77 ± 17.95 | 12 | -0.39 [-2.66, 1.92] | 1.28 [-0.94, 3.53] | 2.03 [-1.11, 5.25] | | | | | | |
| L 961 | L 698 | L Piriform area | 20.18 ± 2.77 | 12 | 32.56 ± 12.33 | 11 | 22.35 ± 3.81 | 9 | 95.32 ± 15.51 | 11 | -0.12 [-2.32, 2.1] | 1.02 [-1.1, 3.17] | 2.06 [-1.07, 5.27] | | | | | | |
| R 619 | R 698 | R Nucleus of the lateral olfactory tract | 15.24 ± 5.00 | 13 | 28.54 ± 11.13 | 6 | 12.29 ± 3.44 | 10 | 30.33 ± 8.99 | 6 | -0.41 [-2.58, 1.72] | -0.19 [-2.5, 2.22] | 1.02 [-2.45, 4.5] | | | | | | |
| L 619 | L 698 | L Nucleus of the lateral olfactory tract | 15.47 ± 6.13 | 10 | 40.34 ± 22.24 | 7 | 23.31 ± 8.47 | 8 | 41.71 ± 12.07 | 6 | 0.21 [-2.14, 2.55] | 0.74 [-1.63, 3.16] | -1.37 [-4.85, 2.2] | | | | | | |
| R 260 | R 619 | R Nucleus of the lateral olfactory tract, mo | 51.39 ± 23.28 | 13 | 44.57 ± 23.43 | 6 | 20.90 ± 7.25 | 9 | 41.71 ± 21.73 | 6 | -0.83 [-2.99, 1.35] | -1.18 [-3.58, 1.18] | 1.26 [-2.28, 4.67] | | | | | | |
| L 260 | L 619 | L Nucleus of the lateral olfactory tract, mo | 36.09 ± 14.87 | 10 | 31.02 ± 14.72 | 6 | 28.21 ± 10.92 | 8 | 43.82 ± 21.87 | 6 | -0.54 [-2.86, 1.79] | -1.42 [-3.86, 1.01] | 0.05 [-3.51, 3.61] | | | | | | |
| R 268 | R 619 | R Nucleus of the lateral olfactory tract, py | 27.31 ± 10.44 | 13 | 78.51 ± 32.74 | 6 | 25.07 ± 8.84 | 10 | 73.88 ± 24.95 | 6 | -0.3 [-2.43, 1.83] | 0.81 [-1.56, 3.17] | 0.62 [-2.73, 4.07] | | | | | | |
| L 268 | L 619 | L Nucleus of the lateral olfactory tract, pyr | 34.58 ± 14.31 | 10 | 110.60 ± 62.91 | 7 | 61.90 ± 21.29 | 8 | 104.47 ± 25.15 | 6 | 0.49 [-1.84, 2.83] | 1.33 [-1.03, 3.65] | -1.92 [-5.43, 1.62] | | | | | | |
| R 1139 | R 619 | R Nucleus of the lateral olfactory tract, lay | 19.95 ± 6.97 | 13 | 40.97 ± 20.19 | 6 | 19.55 ± 7.76 | 10 | 57.88 ± 24.48 | 6 | -0.23 [-2.38, 1.89] | 0.01 [-2.33, 2.37] | 1.55 [-1.86, 5.01] | | | | | | |
| L 1139 | L 619 | L Nucleus of the lateral olfactory tract, lay | 23.78 ± 11.81 | 10 | 114.79 ± 88.86 | 7 | 43.27 ± 18.66 | 8 | 206.06 ± 173.41 | 6 | 0.51 [-1.78, 2.85] | 2.95 [0.58, 5.35]* | 2.07 [-1.42, 5.58] | -3.17 [-5.31, -0.96]* | -0.84 [-3.01, 1.31] | -6.36 [-8.77, -4.11]* | -2.31 [-4.55, 0.04] | -3.18 [-5.49, -0.67]* | -5.51 [-7.91, -3.03]* |
| R 631 | R 698 | R Cortical amygdalar area | 11.47 ± 2.67 | 14 | 15.75 ± 3.61 | 11 | 8.81 ± 1.51 | 10 | 37.68 ± 11.97 | 12 | -0.47 [-2.58, 1.64] | -0.22 [-2.27, 1.85] | 2.36 [-0.64, 5.34] | | | | | | |
| L 631 | L 698 | L Cortical amygdalar area | 11.12 ± 2.75 | 14 | 20.61 ± 6.10 | 13 | 8.48 ± 2.27 | 10 | 26.91 ± 5.97 | 12 | -0.39 [-2.51, 1.73] | 0.5 [-1.5, 2.5] | 0.73 [-2.28, 3.74] | | | | | | |
| R 639 | R 631 | R Cortical amygdalar area, anterior part | 49.04 ± 10.48 | 15 | 75.23 ± 15.14 | 10 | 48.66 ± 8.13 | 10 | 112.09 ± 23.51 | 12 | -0.27 [-2.35, 1.86] | 0.01 [-2.08, 2.12] | 0.9 [-2.12, 3.97] | | | | | | |
| L 639 | L 631 | L Cortical amygdalar area, anterior part | 48.15 ± 10.58 | 14 | 91.56 ± 17.99 | 12 | 40.61 ± 9.81 | 10 | 89.02 ± 15.67 | 12 | -0.3 [-2.43, 1.81] | 0.7 [-1.33, 2.75] | -0.14 [-3.17, 2.92] | | | | | | |
| R 647 | R 631 | R Cortical amygdalar area, posterior part | 7.52 ± 2.84 | 13 | 11.28 ± 3.23 | 12 | 4.93 ± 1.70 | 10 | 37.48 ± 11.79 | 9 | -0.51 [-2.69, 1.67] | 0.55 [-1.49, 2.63] | 3.44 [0.33, 6.49]* | -0.75 [-2.59, 1.14] | 0.17 [-1.79, 2.25] | -4.29 [-6.34, -2.3]* | -0.93 [-2.96, 1.09] | -3.53 [-5.53, -1.48]* | -4.45 [-6.6, -2.24]* |
| L 647 | L 631 | L Cortical amygdalar area, posterior part | 5.78 ± 1.43 | 13 | 14.89 ± 3.43 | 13 | 4.45 ± 1.18 | 9 | 29.35 ± 7.48 | 11 | -0.26 [-2.43, 1.89] | 1.71 [-0.33, 3.73] | 2.26 [-0.71, 5.33] | | | | | | |
| R 655 | R 647 | R Cortical amygdalar area, posterior part, | 15.03 ± 5.08 | 13 | 19.09 ± 5.67 | 12 | 10.28 ± 4.09 | 10 | 74.75 ± 21.75 | 9 | -0.47 [-2.6, 1.73] | 0.33 [-1.72, 2.42] | 3.62 [0.51, 6.79]* | -0.54 [-2.41, 1.37] | 0.13 [-1.8, 2.1] | -4.27 [-6.25, -2.21]* | -0.67 [-2.64, 1.36] | -3.75 [-5.76, -1.7]* | -4.43 [-6.6, -2.3]* |
| L 655 | L 647 | L Cortical amygdalar area, posterior part, l | 10.20 ± 3.12 | 13 | 25.85 ± 8.75 | 13 | 10.61 ± 3.31 | 9 | 58.92 ± 14.25 | 11 | 0 [-2.21, 2.22] | 1.66 [-0.38, 3.71] | 2.76 [-0.32, 5.88] | | | | | | |
| R 663 | R 647 | R Cortical amygdalar area, posterior part, | 17.98 ± 8.08 | 11 | 26.92 ± 9.35 | 12 | 10.40 ± 3.13 | 9 | 65.03 ± 27.72 | 8 | -0.51 [-2.78, 1.68] | 0.57 [-1.57, 2.71] | 2.48 [-0.73, 5.69] | | | | | | |
| L 663 | L 647 | L Cortical amygdalar area, posterior part, r | 13.45 ± 4.01 | 11 | 32.59 ± 8.45 | 13 | 7.02 ± 1.80 | 9 | 53.96 ± 14.78 | 9 | -0.5 [-2.77, 1.71] | 1.56 [-0.54, 3.62] | 1.72 [-1.41, 4.93] | | | | | | |
| R 788 | R 698 | R Piriform-amygdalar area | 13.50 ± 2.82 | 15 | 35.75 ± 12.88 | 11 | 15.95 ± 2.13 | 10 | 90.37 ± 15.91 | 11 | -0.09 [-2.17, 2.05] | 0.94 [-1.13, 3.02] | 3.97 [0.9, 7.07]* | -1.17 [-2.98, 0.73] | -0.24 [-2.2, 1.68] | -5.64 [-7.49, -3.68]* | -0.91 [-2.9, 1.13] | -4.49 [-6.48, -2.51]* | -5.39 [-7.49, -3.32]* |
| L 788 | L 698 | L Piriform-amygdalar area | 12.94 ± 2.71 | 15 | 38.14 ± 15.13 | 15 | 16.46 ± 4.25 | 11 | 78.13 ± 12.67 | 11 | -0.07 [-2.21, 1.99] | 1.7 [-0.3, 3.69] | 2.53 [-0.43, 5.52] | | | | | | |
| R 1089 | R 695 | R Hippocampal formation | 1.21 ± 0.27 | 15 | 7.66 ± 2.32 | 16 | 1.81 ± 0.39 | 12 | 13.40 ± 3.20 | 12 | -0.78 [-7.77, 6.27] | -0.68 [-7.44, 6.23] | -0.98 [-9.23, 7.08] | | | | | | |
| L 1089 | L 695 | L Hippocampal formation | 0.98 ± 0.17 | 15 | 7.42 ± 2.28 | 16 | 1.38 ± 0.25 | 12 | 11.56 ± 2.31 | 12 | -0.89 [-7.78, 6.07] | -0.03 [-6.69, 6.68] | -0.95 [-9.08, 7.04] | | | | | | |
| R 1080 | R 1089 | R Hippocampal region | 1.53 ± 0.34 | 15 | 10.13 ± 3.15 | 16 | 2.27 ± 0.48 | 12 | 17.46 ± 4.33 | 12 | -0.8 [-7.64, 6.12] | -0.51 [-7.19, 6.26] | -0.91 [-8.87, 7.13] | | | | | | |
| L 1080 | L 1089 | L Hippocampal region | 1.23 ± 0.20 | 15 | 10.25 ± 3.16 | 16 | 1.80 ± 0.33 | 12 | 15.48 ± 3.19 | 12 | -0.9 [-8.01, 6.23] | 0.34 [-6.63, 7.17] | -0.85 [-8.96, 7.34] | | | | | | |
| R 375 | R 1080 | R Ammon's horn | 1.67 ± 0.39 | 15 | 7.60 ± 2.04 | 16 | 2.43 ± 0.45 | 12 | 15.87 ± 4.25 | 12 | -0.75 [-7.59, 6.13] | -1.56 [-8.35, 5.25] | -1.17 [-9.27, 6.9] | | | | | | |
| L 375 | L 1080 | L Ammon's horn | 1.28 ± 0.25 | 15 | 8.00 ± 2.30 | 16 | 2.23 ± 0.41 | 12 | 11.54 ± 2.26 | 12 | -1.07 [-8.15, 5.95] | -1.05 [-7.91, 5.84] | -1.71 [-9.78, 6.33] | | | | | | |
| R 382 | R 375 | R Field CA1 | 2.50 ± 0.55 | 15 | 10.49 ± 2.91 | 16 | 3.79 ± 0.83 | 12 | 31.40 ± 10.41 | 12 | -0.12 [-7.09, 6.8] | -1.23 [-7.98, 5.54] | -0.17 [-8.15, 7.88] | | | | | | |
| L 382 | L 375 | L Field CA1 | 1.69 ± 0.43 | 15 | 11.19 ± 3.70 | 16 | 3.82 ± 1.02 | 12 | 24.27 ± 4.65 | 12 | 0.01 [-7.02, 7.07] | -0.19 [-7.06, 6.6] | -0.11 [-8.1, 7.86] | | | | | | |
| R 423 | R 375 | R Field CA2 | 1.17 ± 0.59 | 15 | 11.48 ± 4.05 | 16 | 2.39 ± 1.18 | 12 | 25.80 ± 6.74 | 12 | 0.66 [-6.26, 7.59] | 2.18 [-4.48, 8.86] | 2.01 [-5.97, 9.86] | | | | | | |
| L 423 | L 375 | L Field CA2 | 2.06 ± 0.68 | 15 | 15.77 ± 5.94 | 16 | 2.98 ± 1.21 | 12 | 20.51 ± 6.74 | 11 | -1.03 [-7.95, 5.89] | -0.14 [-6.74, 6.4] | -1.24 [-9.36, 6.94] | | | | | | |
| R 463 | R 375 | R Field CA3 | 4.80 ± 1.33 | 14 | 21.51 ± 6.03 | 16 | 6.21 ± 1.04 | 12 | 35.27 ± 9.84 | 12 | -1.13 [-8.1, 5.92] | -1.89 [-8.74, 4.9] | -1.87 [-10.08, 6.3] | | | | | | |
| L 463 | L 375 | L Field CA3 | 3.57 ± 0.76 | 15 | 21.43 ± 6.24 | 16 | 5.17 ± 0.74 | 12 | 22.80 ± 5.27 | 11 | -1.31 [-8.24, 5.52] | -1.28 [-8.07, 5.47] | -2.11 [-10.26, 5.89] | | | | | | |
| R 726 | R 1080 | R Dentate gyrus | 4.04 ± 0.87 | 15 | 36.43 ± 12.86 | 16 | 6.23 ± 1.55 | 12 | 55.24 ± 12.89 | 12 | -0.75 [-7.6, 6.23] | 0.82 [-5.94, 7.49] | -0.68 [-8.71, 7.43] | | | | | | |
| L 726 | L 1080 | L Dentate gyrus | 3.51 ± 0.60 | 15 | 36.54 ± 12.74 | 16 | 4.13 ± 0.90 | 11 | 56.09 ± 12.35 | 12 | -0.75 [-7.96, 6.39] | 1.71 [-5.06, 8.53] | -0.05 [-8.31, 8.03] | | | | | | |
| R 10703 | R 726 | R Dentate gyrus, molecular layer | 5.73 ± 1.19 | 15 | 42.80 ± 13.80 | 16 | 9.46 ± 2.88 | 12 | 66.77 ± 14.13 | 12 | -0.83 [-7.74, 6.08] | -0.14 [-6.88, 6.55] | -1.05 [-9.22, 7.12] | | | | | | |
| L 10703 | L 726 | L Dentate gyrus, molecular layer | 5.54 ± 1.14 | 15 | 40.57 ± 14.01 | 16 | 5.66 ± 1.54 | 11 | 78.01 ± 19.84 | 12 | -0.59 [-7.54, 6.37] | 0.28 [-6.53, 7.07] | -0.09 [-8.07, 7.83] | | | | | | |
| R 10704 | R 726 | R Dentate gyrus, polymorph layer | 8.36 ± 2.79 | 15 | 120.11 ± 50.37 | 16 | 23.96 ± 9.26 | 12 | 157.82 ± 43.21 | 12 | 13.67 [-22.98, 50.16] | 05.86 [71.57, 140.05] | 10.1 [-41.49, 62.26] | -13.77 [-26.42, -0.71]* | -1.91 [-16.12, 11.84] | -19.22 [-33.62, -5.54]* | -11.92 [-25.82, 1.87] | -5.44 [-19.06, 8.48] | -17.37 [-31.96, -1.95]* |
| L 10704 | L 726 | L Dentate gyrus, polymorph layer | 7.13 ± 2.96 | 15 | 145.22 ± 50.71 | 16 | 7.85 ± 2.64 | 11 | 115.20 ± 29.64 | 12 | -0.91 [-38.11, 36.06] | 32.29 [98.53, 166.68] | -42.94 [-95.53, 9.68] | -19.62 [-32.65, -6.47]* | -0.05 [-13.77, 15.14] | -16.49 [-30.24, -2.19]* | -19.59 [-34.26, -5.71]* | 3.09 [-10.98, 16.95] | -16.46 [-31.75, -1.27]* |
| R 632 | R 726 | R Dentate gyrus, granule cell layer | 13.80 ± 3.95 | 15 | 131.62 ± 48.34 | 16 | 16.46 ± 3.96 | 12 | 193.94 ± 48.17 | 12 | 0.71 [-35.42, 36.8] | 11.84 [77.61, 146.09] | 47.89 [-3.95, 98.47] | -9 [-21.46, 4.29] | -0.27 [-14.53, 13.33] | -14.57 [-28.22, -0.14]* | -8.71 [-22.5, 5.08] | -5.54 [-18.99, 8.4] | -14.26 [-29.05, 0.7] |
| L 632 | L 726 | L Dentate gyrus, granule cell layer | 10.85 ± 2.02 | 15 | 131.25 ± 48.03 | 16 | 14.92 ± 3.23 | 11 | 198.30 ± 45.79 | 12 | 2.2 [-34.91, 39.45] | 114.4 [79.59, 148.99]* | 51.11 [-1.02, 103.75] | -11.71 [-24.74, 1.54] | -0.31 [-15.14, 13.3] | -18.68 [-32.63, -4.29]* | -11.14 [-25.5, 3.05] | -6.99 [-20.86, 6.46] | -18.22 [-33.77, -3.1]* |
| R 982 | R 1080 | R Fasciola cinerea | 7.75 ± 4.41 | 13 | 27.78 ± 10.36 | 16 | 7.65 ± 5.56 | 12 | 28.14 ± 12.89 | 10 | -1.53 [-8.59, 5.6] | -2.57 [-9.38, 4.13] | -2.49 [-10.58, 5.55] | | | | | | |
| L 982 | L 1080 | L Fasciola cinerea | 8.20 ± 6.37 | 13 | 15.61 ± 6.04 | 16 | 6.97 ± 3.79 | 12 | 22.34 ± 9.92 | 11 | -1.44 [-8.4, 5.55] | -3.32 [-10.11, 3.46] | -2.58 [-10.8, 5.6] | | | | | | |
| R 19 | R 1080 | R Induseum griseum | 0.46 ± 0.46 | 15 | 14.41 ± 8.49 | 13 | 7.99 ± 4.24 | 12 | 50.33 ± 11.89 | 12 | 16.79 [9.67, 23.9]* | 21.25 [14.29, 28.23]* | 24.63 [16.5, 32.65]* | -27.11 [-34.99, -19.38]* | -18.54 [-26.82, -10.35]* | -75.66 [-86.51, -64.81]* | -8.55 [-19.76, 1.99] | -48.58 [-59.1, -38.96]* | -57.15 [-67.86, -47.56]* |
| L 19 | L 1080 | L Induseum griseum | 0.51 ± 0.51 | 15 | 16.37 ± 12.13 | 13 | 11.87 ± 9.26 | 12 | 54.85 ± 19.84 | 12 | 18.51 [11.57, 25.54]* | 20.54 [13.6, 27.53]* | 23.08 [15.04, 31.11]* | -26.39 [-34.41, -18.61]* | -20.23 [-27.97, -12.15]* | -75.1 [-86.11, -64.61]* | -6.21 [-16.78, 4.74] | -48.68 [-58.91, -38.93]* | -54.88 [-64.84, -44.57]* |
| R 822 | R 1089 | R Retrohippocampal region | 2.34 ± 0.69 | 14 | 5.58 ± 2.14 | 14 | 2.92 ± 1.07 | 12 | 14.98 ± 3.14 | 9 | -1 [-7.96, 6.08] | -2.45 [-9.4, 4.58] | -1.7 [-9.93, 6.5] | | | | | | |
| L 822 | L 1089 | L Retrohippocampal region | 1.78 ± 0.60 | 14 | 4.99 ± 1.86 | 16 | 1.74 ± 0.50 | 12 | 9.33 ± 2.84 | 11 | -1.19 [-8.14, 5.9] | -2.68 [-9.51, 4.07] | -2.07 [-10.22, 6.12] | | | | | | |
| R 909 | R 822 | R Entorhinal area | 2.66 ± 0.88 | 14 | 5.33 ± 1.68 | 13 | 2.10 ± 0.77 | 11 | 18.22 ± 4.64 | 9 | -0.92 [-8.01, 6.19] | -2.52 [-9.7, 4.48] | -1.4 [-9.46, 6.82] | | | | | | |
| L 909 | L 822 | L Entorhinal area | 2.08 ± 0.69 | 14 | 5.68 ± 1.80 | 16 | 1.95 ± 0.59 | 11 | 11.93 ± 3.69 | 11 | -1.19 [-8.21, 5.88] | -2.64 [-9.43, 4.17] | -1.87 [-10.09, 6.17] | | | | | | |
| R 918 | R 909 | R Entorhinal area, lateral part | 4.11 ± 1.36 | 14 | 8.27 ± 2.61 | 13 | 3.24 ± 1.18 | 11 | 28.09 ± 7.15 | 9 | -0.96 [-8.1, 6.19] | -2.52 [-9.45, 4.47] | -1.42 [-9.65, 6.82] | | | | | | |
| L 918 | L 909 | L Entorhinal area, lateral part | 3.21 ± 1.07 | 14 | 9.29 ± 3.23 | 16 | 3.01 ± 0.91 | 11 | 18.42 ± 5.69 | 11 | -1.24 [-8.28, 5.67] | -2.56 [-9.5, 4.34] | -1.91 [-10.01, 6.15] | | | | | | |
| R 502 | R 822 | R Subiculum | 7.24 ± 2.88 | 12 | 21.06 ± 9.08 | 11 | 20.49 ± 10.80 | 10 | 31.91 ± 6.98 | 7 | -0.62 [-7.96, 6.68] | -2.82 [-10.04, 4.51] | -1.99 [-10.37, 6.47] | | | | | | |
| L 502 | L 822 | L Subiculum | 7.28 ± 2.58 | 12 | 19.18 ± 12.82 | 12 | 6.74 ± 2.08 | 10 | 24.45 ± 8.55 | 7 | -1.09 [-8.49, 6.29] | -3.28 [-10.39, 3.98] | -1.86 [-10.22, 6.39] | | | | | | |
| R 484682 | R 822 | R Prosubiculum | 1.51 ± 1.14 | 10 | 19.98 ± 8.23 | 10 | 22.49 ± 6.78 | 10 | 53.19 ± 13.12 | 7 | 6.14 [-1.18, 13.4] | 2.87 [-4.57, 10.18] | 3.33 [-5.03, 11.68] | | | | | | |
| L 484682 | L 822 | L Prosubiculum | 4.00 ± 2.67 | 10 | 14.14 ± 7.89 | 11 | 5.97 ± 2.93 | 10 | 31.01 ± 13.21 | 7 | -0.58 [-7.84, 6.67] | -2.01 [-9.24, 5.26] | -1.02 [-9.28, 7.39] | | | | | | |
| R 703 | R 688 | R Cortical subplate | 5.74 ± 1.15 | 15 | 12.77 ± 3.41 | 16 | 6.13 ± 1.12 | 12 | 29.73 ± 5.83 | 12 | -0.24 [-1.94, 1.48] | 1.12 [-0.5, 2.77] | 2.79 [0.37, 5.23]* | -1.24 [-3.26, 0.72] | -0.07 [-2.11, 1.95] | -4.55 [-6.59, -2.3]* | -1.16 [-3.31, 0.94] | -3.29 [-5.4, -1.17]* | -4.46 [-6.66, -2.19]* |
| L 703 | L 688 | L Cortical subplate | 4.78 ± 0.76 | 15 | 12.95 ± 3.38 | 16 | 6.81 ± 1.22 | 12 | 28.76 ± 5.36 | 12 | 0.12 [-1.56, 1.81] | 1.6 [-0.03, 3.21] | 2.78 [0.31, 5.16]* | -1.74 [-3.62, 0.24] | -0.44 [-2.43, 1.61] | -5.37 [-7.57, -3.29]* | -1.27 [-3.47, 0.73] | -3.64 [-5.85, -1.65]* | -4.92 [-7.2, -2.77]* |
| R 583 | R 703 | R Claustrum | 18.66 ± 5.42 | 15 | 41.93 ± 14.47 | 15 | 31.45 ± 8.47 | 12 | 106.07 ± 15.90 | 12 | 0.38 [-1.31, 2.06] | 1.14 [-0.5, 2.78] | 2.64 [0.13, 5.12]* | -1.26 [-3.28, 0.71] | -0.7 [-2.77, 1.28] | -5.03 [-7.26, -3]* | -0.56 [-2.68, 1.52] | -3.76 [-5.99, -1.73]* | -4.32 [-6.6, -2.13]* |
| L 583 | L 703 | L Claustrum | 18.06 ± 4.12 | 15 | 41.60 ± 13.14 | 15 | 27.92 ± 4.68 | 12 | 108.11 ± 19.85 | 12 | 0.24 [-1.45, 1.95] | 1.2 [-0.46, 2.85] | 3.04 [0.61, 5.47]* | -1.33 [-3.37, 0.57] | -0.56 [-2.54, 1.52] | -5.33 [-7.45, -3.16]* | -0.77 [-2.89, 1.29] | -4 [-6.1, -1.9]* | -4.76 [-6.93, -2.49]* |
| R 942 | R 703 | R Endopiriform nucleus | 6.55 ± 1.50 | 13 | 16.32 ± 4.66 | 14 | 7.14 ± 1.39 | 12 | 28.77 ± 5.41 | 12 | -0.2 [-1.95, 1.5] | 1.16 [-0.51, 2.86] | 1.94 [-0.52, 4.45] | | | | | | |
| L 942 | L 703 | L Endopiriform nucleus | 5.07 ± 0.83 | 14 | 13.44 ± 3.77 | 16 | 8.24 ± 1.53 | 12 | 34.08 ± 6.71 | 11 | 0.36 [-1.35, 2.05] | 1.58 [-0.08, 3.19] | 2.93 [0.46, 5.36]* | -1.72 [-3.67, 0.3] | -0.68 [-2.73, 1.35] | -5.74 [-7.91, -3.53]* | -1.04 [-3.18, 1] | -4.02 [-6.18, -1.9]* | -5.06 [-7.34, -2.86]* |
| R 952 | R 942 | R Endopiriform nucleus, dorsal part | 13.00 ± 3.30 | 13 | 36.72 ± 10.53 | 15 | 12.48 ± 2.04 | 12 | 55.72 ± 9.41 | 12 | -0.33 [-2.04, 1.39] | 1.65 [0, 3.31] | 1.48 [-0.98, 3.92] | -1.77 [-3.77, 0.26] | 0.02 [-1.98, 2.12] | -3.66 [-5.84, -1.52]* | -1.78 [-3.89, 0.31] | -1.88 [-4.03, 0.21] | -3.66 [-5.88, -1.49]* |
| L 952 | L 942 | L Endopiriform nucleus, dorsal part | 10.88 ± 1.85 | 14 | 27.75 ± 7.25 | 16 | 16.92 ± 3.05 | 12 | 62.27 ± 12.59 | 12 | 0.29 [-1.46, 2.03] | 1.48 [-0.22, 3.19] | 2.48 [0, 4.95] | | | | | | |
| R 966 | R 942 | R Endopiriform nucleus, ventral part | 13.07 ± 3.56 | 13 | 19.91 ± 5.96 | 15 | 17.62 ± 4.84 | 12 | 60.30 ± 13.57 | 12 | 0.06 [-1.69, 1.79] | 0.36 [-1.27, 2.05] | 2.69 [0.25, 5.11]* | -0.5 [-2.45, 1.55] | -0.39 [-2.54, 1.59] | -3.98 [-6.26, -1.89]* | -0.11 [-2.31, 1.9] | -3.5 [-5.66, -1.44]* | -3.61 [-5.83, -1.35]* |
| L 966 | L 942 | L Endopiriform nucleus, ventral part | 8.68 ± 1.81 | 14 | 25.82 ± 8.54 | 16 | 15.78 ± 3.45 | 12 | 63.55 ± 15.57 | 12 | 0.56 [-1.16, 2.24] | 1.91 [0.24, 3.54]* | 3.39 [0.94, 5.78]* | -2.04 [-4.05, -0.1]* | -0.87 [-2.91, 1.18] | -6.72 [-8.96, -4.59]* | -1.16 [-3.34, 0.9] | -4.69 [-6.8, -2.53]* | -5.83 [-8.1, -3.63]* |

| | | | | | | | | | | | | | | | | | | | |
|---|---|---|---|---|---|---|---|---|---|---|---|---|---|---|---|---|---|---|---|
| R 131 | R 703 | R Lateral amygdalar nucleus | 11.85 ± 3.78 | 14 | 33.91 ± 9.13 | 15 | 11.11 ± 2.27 | 12 | 69.13 ± 15.59 | 12 | -0.3 [-2.03, 1.44] | 1.75 [0.1, 3.38]* | 2.94 [0.48, 5.42]* | -1.88 [-3.83, 0.17] | -0.03 [-2.12, 1.93] | -5.26 [-7.37, -3.13]* | -1.86 [-3.99, 0.29] | -3.38 [-5.45, -1.21]* | -5.24 [-7.43, -2.96]* |
| L 131 | L 703 | L Lateral amygdalar nucleus | 6.60 ± 1.94 | 14 | 37.24 ± 10.12 | 16 | 12.61 ± 3.04 | 12 | 56.53 ± 13.12 | 12 | 0.67 [-1.04, 2.38] | 4.6 [2.93, 6.24]* | 1.85 [-0.59, 4.34] | -4.75 [-6.68, -2.72]* | -0.99 [-3.08, 0.98] | -7.99 [-10.26, -5.93]* | -3.75 [-5.84, -1.61]* | -3.25 [-5.38, -1.16]* | -6.99 [-9.28, -4.79]* |
| R 295 | R 703 | R Basolateral amygdalar nucleus | 7.80 ± 1.80 | 15 | 19.23 ± 5.51 | 14 | 9.71 ± 2.01 | 9 | 44.45 ± 9.72 | 12 | -0.08 [-1.89, 1.68] | 1.12 [-0.57, 2.76] | 3.14 [0.69, 5.68]* | -1.26 [-3.16, 0.78] | -0.24 [-2.38, 1.82] | -5.06 [-7.23, -2.93]* | -1.01 [-3.3, 1.1] | -3.8 [-6.01, -1.75]* | -4.81 [-7.16, -2.59]* |
| L 295 | L 703 | L Basolateral amygdalar nucleus | 6.04 ± 1.23 | 14 | 19.49 ± 5.65 | 14 | 9.68 ± 2.38 | 10 | 48.97 ± 8.65 | 11 | 0.41 [-1.35, 2.18] | 1.84 [0.19, 3.53]* | 4.01 [1.48, 6.51]* | -1.98 [-3.99, -0.02]* | -0.74 [-2.72, 1.43] | -7.13 [-9.32, -4.96]* | -1.23 [-3.43, 0.93] | -5.15 [-7.31, -3.07]* | -6.4 [-8.71, -4.1]* |
| R 303 | R 295 | R Basolateral amygdalar nucleus, anterior | 23.11 ± 5.94 | 15 | 41.34 ± 10.69 | 13 | 31.69 ± 6.81 | 9 | 111.23 ± 25.99 | 12 | 0.05 [-1.7, 1.85] | 0.88 [-0.81, 2.54] | 2.37 [-0.16, 4.88] | | | | | | |
| L 303 | L 295 | L Basolateral amygdalar nucleus, anterior | 16.63 ± 3.38 | 14 | 48.80 ± 11.50 | 14 | 28.29 ± 7.46 | 10 | 132.88 ± 22.42 | 11 | 0.51 [-1.23, 2.23] | 1.55 [-0.12, 3.23] | 4.08 [1.55, 6.58]* | -1.7 [-3.66, 0.32] | -0.84 [-2.85, 1.33] | -7 [-9.17, -4.84]* | -0.84 [-3.01, 1.31] | -5.33 [-7.61, -3.3]* | -6.17 [-8.48, -3.89]* |
| R 311 | R 295 | R Basolateral amygdalar nucleus, posterio | 9.43 ± 3.52 | 14 | 27.76 ± 10.45 | 13 | 8.08 ± 2.77 | 9 | 83.37 ± 22.15 | 9 | -0.4 [-2.22, 1.4] | 2.19 [0.5, 3.87]* | 5.28 [2.71, 7.88]* | -2.32 [-4.33, -0.27]* | 0.08 [-2.08, 2.21] | -7.94 [-10.21, -5.78]* | -2.41 [-4.63, -0.15]* | -5.61 [-7.92, -3.47]* | -8.01 [-10.48, -5.75]* |
| L 311 | L 295 | L Basolateral amygdalar nucleus, posterior | 8.28 ± 2.63 | 13 | 33.23 ± 14.68 | 14 | 8.01 ± 2.91 | 10 | 69.34 ± 14.70 | 10 | -0.15 [-1.95, 1.59] | 2.71 [1.05, 4.36]* | 4.01 [1.5, 6.52]* | -2.84 [-4.83, -0.85]* | -0.17 [-2.24, 1.95] | -7.43 [-9.67, -5.16]* | -2.67 [-4.75, -0.41]* | -4.58 [-6.72, -2.34]* | -7.26 [-9.57, -4.98]* |
| R 451 | R 295 | R Basolateral amygdalar nucleus, ventral p | 10.31 ± 3.25 | 14 | 14.17 ± 4.06 | 12 | 10.49 ± 2.65 | 9 | 64.97 ± 12.25 | 10 | -0.25 [-2.05, 1.54] | 1.13 [-0.54, 2.81] | 4.09 [1.53, 6.63]* | -1.27 [-3.24, 0.81] | -0.07 [-2.25, 2.02] | -5.85 [-8.02, -3.61]* | -1.19 [-3.36, 1.04] | -4.58 [-6.78, -2.38]* | -5.77 [-8.09, -3.49]* |
| L 451 | L 295 | L Basolateral amygdalar nucleus, ventral p | 7.05 ± 2.08 | 12 | 16.31 ± 4.96 | 13 | 12.63 ± 5.91 | 10 | 66.23 ± 17.60 | 10 | 0.69 [-1.12, 2.51] | 1.46 [-0.29, 3.17] | 5.47 [2.87, 8.08]* | -1.59 [-3.57, 0.52] | -1.01 [-3.12, 1.14] | -8.49 [-10.71, -6.26]* | -0.59 [-2.74, 1.56] | -6.89 [-9.12, -4.77]* | -7.48 [-9.79, -5.2]* |
| R 319 | R 703 | R Basomedial amygdalar nucleus | 10.85 ± 2.29 | 14 | 20.10 ± 4.75 | 14 | 10.71 ± 3.12 | 9 | 51.90 ± 11.72 | 12 | -0.19 [-1.96, 1.56] | 0.6 [-1.06, 2.26] | 2.85 [0.37, 5.35]* | -0.72 [-2.74, 1.32] | -0.14 [-2.26, 2.01] | -4.14 [-6.28, -1.97]* | -0.59 [-2.83, 1.57] | -3.4 [-5.48, -1.18]* | -4 [-6.23, -1.65]* |
| L 319 | L 703 | L Basomedial amygdalar nucleus | 11.38 ± 2.37 | 15 | 23.07 ± 7.24 | 14 | 12.68 ± 3.10 | 11 | 50.19 ± 9.85 | 12 | -0.18 [-1.92, 1.53] | 1.28 [-0.41, 2.95] | 1.81 [-0.7, 4.32] | | | | | | |
| R 327 | R 319 | R Basomedial amygdalar nucleus, anterior | 29.35 ± 5.44 | 14 | 55.53 ± 13.90 | 13 | 28.59 ± 7.83 | 9 | 119.13 ± 24.17 | 12 | -0.2 [-2, 1.59] | 1.08 [-0.62, 2.75] | 1.67 [-0.88, 4.16] | | | | | | |
| L 327 | L 319 | L Basomedial amygdalar nucleus, anterior | 29.25 ± 6.31 | 15 | 65.09 ± 22.99 | 12 | 37.64 ± 11.02 | 11 | 111.96 ± 22.29 | 12 | -0.01 [-1.77, 1.68] | 2.04 [0.36, 3.72]* | 0.29 [-2.13, 2.83] | -2.18 [-4.26, -0.17]* | -0.31 [-2.41, 1.72] | -3.19 [-5.38, -1.08]* | -1.86 [-4.01, 0.33] | -1.01 [-3.2, 1.08] | -2.87 [-5.12, -0.69]* |
| R 334 | R 319 | R Basomedial amygdalar nucleus, posterio | 12.33 ± 5.17 | 13 | 20.80 ± 8.41 | 12 | 13.47 ± 4.90 | 9 | 68.79 ± 18.46 | 9 | 0 [-1.83, 1.82] | 0.97 [-0.77, 2.7] | 2.84 [0.22, 5.43]* | -1.1 [-3.19, 0.88] | -0.32 [-2.54, 1.74] | -4.67 [-6.92, -2.45]* | -0.78 [-3.01, 1.43] | -3.58 [-5.82, -1.41]* | -4.35 [-6.58, -1.94]* |
| L 334 | L 319 | L Basomedial amygdalar nucleus, posterior | 12.23 ± 3.28 | 14 | 32.57 ± 14.32 | 14 | 10.42 ± 3.11 | 10 | 71.97 ± 15.41 | 11 | -0.34 [-2.08, 1.46] | 1.98 [0.31, 3.63]* | 2.81 [0.25, 5.3]* | -2.12 [-4.13, -0.12]* | 0.01 [-2.11, 2.1] | -5.33 [-7.42, -3]* | -2.13 [-4.27, 0.07] | -3.2 [-5.41, -1.14]* | -5.34 [-7.59, -3]* |
| R 780 | R 703 | R Posterior amygdalar nucleus | 11.44 ± 3.52 | 12 | 30.84 ± 10.69 | 12 | 9.94 ± 2.57 | 11 | 73.68 ± 28.57 | 8 | -0.36 [-2.11, 1.4] | 1.85 [0.1, 3.58]* | 3.48 [0.87, 6.07]* | -1.98 [-3.99, 0.16] | 0.04 [-2.11, 2.09] | -5.84 [-8.12, -3.51]* | -2.01 [-4.22, 0.12] | -3.87 [-6.03, -1.57]* | -5.87 [-8.12, -3.51]* |
| L 780 | L 703 | L Posterior amygdalar nucleus | 9.90 ± 3.56 | 12 | 19.16 ± 6.28 | 14 | 11.00 ± 4.82 | 11 | 46.09 ± 17.83 | 9 | -0.11 [-1.87, 1.63] | 1.23 [-0.47, 2.95] | 1.98 [-0.56, 4.51] | | | | | | |
| R 477 | R 623 | R Striatum | 3.90 ± 0.79 | 15 | 8.32 ± 1.57 | 16 | 3.98 ± 0.56 | 12 | 12.29 ± 2.05 | 12 | -0.32 [-1.99, 1.33] | 0.98 [-0.55, 2.54] | 0.8 [-1.54, 3.1] | | | | | | |
| L 477 | L 623 | L Striatum | 3.45 ± 0.60 | 15 | 8.44 ± 1.57 | 16 | 4.53 ± 0.72 | 12 | 13.19 ± 1.87 | 11 | -0.02 [-1.68, 1.64] | 1.3 [-0.24, 2.89] | 0.58 [-1.74, 2.91] | | | | | | |
| R 485 | R 477 | R Striatum dorsal region | 2.65 ± 0.50 | 15 | 8.68 ± 2.08 | 16 | 3.28 ± 0.49 | 12 | 10.09 ± 1.79 | 12 | -0.09 [-1.76, 1.53] | 2.13 [0.57, 3.7]* | 0.07 [-2.2, 2.32] | -2.29 [-3.7, -0.82]* | -0.24 [-1.77, 1.32] | -2.95 [-4.52, -1.31]* | -2.06 [-3.7, -0.5]* | -0.66 [-2.26, 0.9] | -2.71 [-4.31, -0.96]* |
| L 485 | L 477 | L Striatum dorsal region | 2.24 ± 0.42 | 15 | 7.53 ± 1.59 | 16 | 3.05 ± 0.49 | 12 | 9.56 ± 1.97 | 12 | 0.01 [-1.57, 1.67] | 2.21 [0.7, 3.81]* | 0.35 [-1.96, 2.6] | -2.38 [-3.87, -0.96]* | -0.34 [-1.91, 1.2] | -3.4 [-5.05, -1.88]* | -2.03 [-3.64, -0.5]* | -1.04 [-2.59, 0.56] | -3.06 [-4.73, -1.4]* |
| R 672 | R 485 | R Caudoputamen | 5.68 ± 1.00 | 14 | 18.53 ± 4.27 | 15 | 6.55 ± 0.98 | 12 | 22.02 ± 3.37 | 11 | -0.12 [-1.77, 1.58] | 2.09 [0.48, 3.71]* | 0.17 [-2.22, 2.43] | -2.25 [-3.79, -0.79]* | -0.22 [-1.83, 1.32] | -2.98 [-4.63, -1.37]* | -2.03 [-3.57, -0.41]* | -0.73 [-2.28, 0.9] | -2.76 [-4.39, -1.04]* |
| L 672 | L 485 | L Caudoputamen | 5.17 ± 0.82 | 13 | 15.06 ± 3.18 | 16 | 6.09 ± 0.98 | 12 | 22.94 ± 3.61 | 10 | -0.06 [-1.73, 1.6] | 1.87 [0.33, 3.45]* | 0.62 [-1.75, 2.93] | -2.03 [-3.54, -0.55]* | -0.28 [-1.87, 1.27] | -3.26 [-4.98, -1.6]* | -1.74 [-3.36, -0.19]* | -1.24 [-2.9, 0.34] | -2.98 [-4.73, -1.28]* |
| R 493 | R 477 | R Striatum ventral region | 5.72 ± 1.23 | 15 | 7.32 ± 1.39 | 15 | 4.60 ± 0.70 | 12 | 14.77 ± 3.25 | 12 | -0.54 [-2.19, 1.11] | 0.29 [-1.26, 1.87] | 1.13 [-1.18, 3.41] | | | | | | |
| L 493 | L 477 | L Striatum ventral region | 4.46 ± 0.85 | 15 | 8.57 ± 1.50 | 15 | 6.01 ± 1.05 | 12 | 14.42 ± 2.91 | 12 | 0.01 [-1.65, 1.66] | 0.93 [-0.65, 2.53] | 0.6 [-1.7, 2.87] | | | | | | |
| R 56 | R 493 | R Nucleus accumbens | 9.77 ± 1.91 | 12 | 16.98 ± 3.92 | 13 | 9.70 ± 1.49 | 12 | 36.39 ± 8.05 | 10 | -0.21 [-1.91, 1.5] | 0.79 [-0.86, 2.47] | 1.74 [-0.71, 4.12] | | | | | | |
| L 56 | L 493 | L Nucleus accumbens | 6.96 ± 1.19 | 13 | 17.07 ± 3.95 | 14 | 11.88 ± 2.70 | 12 | 37.05 ± 8.24 | 10 | 0.41 [-1.27, 2.11] | 1.38 [-0.24, 3.02] | 2.03 [-0.37, 4.39] | | | | | | |
| R 754 | R 493 | R Olfactory tubercle | 17.84 ± 5.20 | 13 | 16.84 ± 2.66 | 12 | 8.86 ± 1.94 | 10 | 27.80 ± 7.05 | 11 | -0.64 [-2.4, 1.11] | -0.18 [-1.82, 1.48] | 0.71 [-1.73, 3.13] | | | | | | |
| L 754 | L 493 | L Olfactory tubercle | 13.76 ± 3.44 | 13 | 18.63 ± 3.87 | 13 | 11.84 ± 2.05 | 12 | 22.29 ± 3.61 | 9 | -0.37 [-2.01, 1.31] | 0.52 [-1.1, 2.17] | -0.28 [-2.68, 2.1] | | | | | | |
| R 275 | R 477 | R Lateral septal complex | 12.94 ± 3.68 | 14 | 21.87 ± 5.28 | 13 | 13.45 ± 1.87 | 12 | 28.37 ± 5.05 | 12 | -0.3 [-1.91, 1.36] | 0.46 [-1.14, 2.07] | 0.33 [-1.99, 2.57] | | | | | | |
| L 275 | L 477 | L Lateral septal complex | 11.84 ± 2.38 | 14 | 24.52 ± 5.16 | 13 | 11.39 ± 1.63 | 12 | 27.39 ± 6.30 | 12 | -0.36 [-2.04, 1.3] | 0.86 [-0.74, 2.47] | 0.13 [-2.26, 2.47] | | | | | | |
| R 242 | R 275 | R Lateral septal nucleus | 19.97 ± 5.55 | 14 | 33.33 ± 8.11 | 13 | 20.45 ± 2.67 | 12 | 44.13 ± 8.68 | 12 | -0.31 [-1.98, 1.36] | 0.45 [-1.12, 2.06] | 0.38 [-1.91, 2.71] | | | | | | |
| L 242 | L 275 | L Lateral septal nucleus | 18.88 ± 3.84 | 14 | 38.14 ± 7.65 | 13 | 17.90 ± 2.61 | 12 | 43.11 ± 10.51 | 12 | -0.4 [-2.03, 1.29] | 0.78 [-0.83, 2.44] | 0.2 [-2.18, 2.55] | | | | | | |
| R 258 | R 242 | R Lateral septal nucleus, rostral (rostrover | 35.85 ± 9.38 | 14 | 68.00 ± 16.64 | 12 | 38.69 ± 5.30 | 12 | 79.48 ± 18.22 | 12 | -0.24 [-1.93, 1.44] | 0.52 [-1.12, 2.17] | 0.24 [-2.16, 2.61] | | | | | | |
| L 258 | L 242 | L Lateral septal nucleus, rostral (rostroven | 35.22 ± 7.40 | 14 | 75.02 ± 16.20 | 12 | 33.98 ± 4.59 | 12 | 78.72 ± 20.94 | 12 | -0.38 [-2.06, 1.29] | 0.74 [-0.88, 2.37] | 0.17 [-2.18, 2.54] | | | | | | |
| R 266 | R 242 | R Lateral septal nucleus, ventral part | 68.77 ± 28.74 | 11 | 75.81 ± 26.88 | 7 | 63.34 ± 11.90 | 10 | 116.88 ± 24.77 | 10 | -0.38 [-2.14, 1.42] | 0.17 [-1.62, 2.07] | 0.32 [-2.3, 2.9] | | | | | | |
| L 266 | L 242 | L Lateral septal nucleus, ventral part | 46.13 ± 15.73 | 10 | 79.99 ± 24.55 | 9 | 52.91 ± 19.22 | 11 | 139.14 ± 31.81 | 9 | -0.27 [-2.03, 1.48] | 0.6 [-1.21, 2.4] | 1.04 [-1.45, 3.56] | | | | | | |
| R 333 | R 275 | R Septohippocampal nucleus | 13.96 ± 6.89 | 14 | 17.06 ± 8.99 | 8 | 19.58 ± 12.76 | 10 | 6.21 ± 4.56 | 7 | -0.01 [-1.73, 1.68] | 0.75 [-1.03, 2.55] | -1.73 [-4.31, 0.81] | | | | | | |
| L 333 | L 275 | L Septohippocampal nucleus | 3.28 ± 2.62 | 13 | 22.64 ± 10.95 | 8 | 2.73 ± 1.84 | 10 | 19.55 ± 11.11 | 7 | -0.58 [-2.29, 1.15] | 6.42 [4.65, 8.2]* | -1.31 [-3.93, 1.29] | -6.58 [-8.26, -4.89]* | 0.25 [-1.38, 1.83] | -5.36 [-7.17, -3.59]* | -6.83 [-8.56, -5.02]* | 1.21 [-0.7, 3.07] | -5.62 [-7.48, -3.74]* |
| R 278 | R 477 | R Striatum-like amygdalar nuclei | 9.22 ± 2.09 | 15 | 16.64 ± 3.09 | 15 | 9.69 ± 2.11 | 12 | 35.13 ± 7.03 | 12 | -0.29 [-1.94, 1.35] | 0.84 [-0.72, 2.42] | 1.55 [-0.78, 3.81] | | | | | | |
| L 278 | L 477 | L Striatum-like amygdalar nuclei | 8.81 ± 1.63 | 15 | 20.04 ± 4.80 | 15 | 12.96 ± 2.61 | 12 | 35.56 ± 5.61 | 12 | 0.13 [-1.53, 1.79] | 1.32 [-0.24, 2.91] | 0.89 [-1.44, 3.22] | | | | | | |
| R 23 | R 278 | R Anterior amygdalar area | 12.99 ± 4.25 | 13 | 18.16 ± 6.36 | 9 | 13.37 ± 4.13 | 9 | 32.16 ± 5.54 | 8 | -0.31 [-2.07, 1.47] | 0.42 [-1.29, 2.17] | 0.99 [-1.57, 3.48] | | | | | | |
| R 536 | R 278 | R Central amygdalar nucleus | 6.43 ± 1.26 | 15 | 15.87 ± 3.50 | 15 | 9.12 ± 2.33 | 11 | 27.61 ± 5.56 | 12 | 0.02 [-1.61, 1.65] | 1.51 [-0.07, 3.11] | 1.07 [-1.22, 3.34] | | | | | | |
| L 536 | L 278 | L Central amygdalar nucleus | 6.81 ± 1.33 | 14 | 16.85 ± 4.25 | 14 | 10.95 ± 2.52 | 12 | 33.78 ± 5.93 | 12 | 0.32 [-1.33, 1.96] | 1.75 [0.19, 3.34]* | 1.24 [-1.09, 3.56] | -1.91 [-3.44, -0.45]* | -0.67 [-2.26, 0.85] | -4.14 [-5.76, -2.56]* | -1.25 [-2.86, 0.3] | -2.24 [-3.85, -0.64]* | -3.5 [-5.17, -1.86]* |
| R 544 | R 536 | R Central amygdalar nucleus, capsular par | 9.29 ± 2.31 | 14 | 40.44 ± 11.74 | 15 | 23.77 ± 8.71 | 11 | 74.55 ± 14.22 | 12 | 1.22 [-0.43, 2.87] | 3.46 [1.86, 5.05]* | 1.72 [-0.57, 4.02] | -3.62 [-5.07, -2.15]* | -1.56 [-3.1, 0.02] | -7.23 [-8.92, -5.67]* | -2.07 [-3.59, -0.42]* | -3.6 [-5.29, -2.11]* | -5.68 [-7.38, -4]* |
| L 544 | L 536 | L Central amygdalar nucleus, capsular part | 12.53 ± 2.94 | 13 | 34.52 ± 7.58 | 14 | 34.44 ± 12.31 | 12 | 84.46 ± 17.21 | 12 | 1.53 [-0.16, 3.2] | 2.09 [0.47, 3.75]* | 1.54 [-0.81, 3.91] | -2.25 [-3.8, -0.72]* | -1.86 [-3.41, -0.23]* | -6 [-7.62, -4.39]* | -0.39 [-1.96, 1.22] | -3.74 [-5.39, -2.19]* | -4.14 [-5.77, -2.49]* |
| R 551 | R 536 | R Central amygdalar nucleus, lateral part | 13.69 ± 4.08 | 14 | 40.88 ± 7.38 | 15 | 14.10 ± 3.96 | 11 | 73.62 ± 17.83 | 12 | -0.31 [-2, 1.35] | 2.09 [0.52, 3.7]* | 1.97 [-0.37, 4.28] | -2.26 [-3.69, -0.73]* | -0.02 [-1.61, 1.54] | -4.58 [-6.22, -2.99]* | -2.22 [-3.92, -0.7]* | -2.33 [-3.88, -0.74]* | -4.56 [-6.22, -2.86]* |
| L 551 | L 536 | L Central amygdalar nucleus, lateral part | 11.88 ± 3.31 | 13 | 39.91 ± 10.47 | 14 | 19.32 ± 4.71 | 12 | 83.61 ± 18.71 | 12 | 0.41 [-1.26, 2.1] | 2.71 [1.15, 4.32]* | 2.34 [0.04, 4.64]* | -2.87 [-4.39, -1.36]* | -0.76 [-2.26, 0.93] | -6.3 [-7.99, -4.69]* | -2.12 [-3.7, -0.49]* | -3.42 [-5.04, -1.82]* | -5.56 [-7.29, -3.93]* |
| R 559 | R 536 | R Central amygdalar nucleus, medial part | 14.63 ± 2.87 | 15 | 21.14 ± 5.89 | 14 | 18.21 ± 4.40 | 11 | 42.00 ± 8.89 | 12 | -0.16 [-1.81, 1.54] | 0.66 [-0.91, 2.27] | 0.67 [-1.69, 3.01] | | | | | | |
| L 559 | L 536 | L Central amygdalar nucleus, medial part | 15.76 ± 3.68 | 14 | 31.16 ± 9.31 | 14 | 18.71 ± 4.73 | 12 | 54.77 ± 7.78 | 12 | -0.11 [-1.76, 1.57] | 1.25 [-0.35, 2.84] | 0.69 [-1.67, 3.01] | | | | | | |
| R 1105 | R 278 | R Intercalated amygdalar nucleus | 21.10 ± 5.77 | 15 | 36.40 ± 12.31 | 14 | 30.68 ± 8.46 | 12 | 90.37 ± 39.04 | 12 | 0.13 [-1.51, 1.78] | 0.96 [-0.64, 2.56] | 1.5 [-0.83, 3.81] | | | | | | |
| L 1105 | L 278 | L Intercalated amygdalar nucleus | 17.67 ± 4.93 | 15 | 42.19 ± 10.44 | 14 | 33.67 ± 5.81 | 12 | 78.20 ± 16.76 | 12 | 0.57 [-1.05, 2.22] | 1.37 [-0.19, 2.97] | 0.78 [-1.53, 3.09] | | | | | | |
| R 403 | R 278 | R Medial amygdalar nucleus | 29.77 ± 7.14 | 15 | 47.16 ± 9.39 | 15 | 27.44 ± 6.10 | 12 | 106.46 ± 19.59 | 12 | -0.41 [-2.07, 1.26] | 0.63 [-0.9, 2.24] | 1.66 [-0.65, 3.97] | | | | | | |
| L 403 | L 278 | L Medial amygdalar nucleus | 28.63 ± 5.64 | 15 | 60.02 ± 16.42 | 15 | 40.35 ± 9.28 | 12 | 106.53 ± 18.44 | 12 | 0.07 [-1.59, 1.77] | 1.13 [-0.42, 2.75] | 0.82 [-1.5, 3.12] | | | | | | |
| R 803 | R 623 | R Pallidum | 2.37 ± 0.54 | 15 | 3.96 ± 1.08 | 15 | 2.62 ± 0.57 | 11 | 5.47 ± 1.05 | 12 | -1.77 [-32.51, 29.11] | -6.65 [-35.95, 23.3] | -11.62 [-54.89, 32.46] | | | | | | |
| L 803 | L 623 | L Pallidum | 2.22 ± 0.41 | 15 | 2.88 ± 0.54 | 15 | 2.72 ± 0.34 | 12 | 6.70 ± 1.54 | 12 | -1.81 [-31.65, 28.8] | -7.16 [-36.59, 22.15] | -10.7 [-53.96, 32.61] | | | | | | |
| R 818 | R 803 | R Pallidum, dorsal region | 1.85 ± 0.48 | 15 | 2.67 ± 0.77 | 14 | 2.02 ± 0.56 | 12 | 4.63 ± 0.80 | 12 | -1.55 [-31.59, 28.18] | -6.77 [-36.37, 22.64] | -11.44 [-55.05, 31.2] | | | | | | |
| L 818 | L 803 | L Pallidum, dorsal region | 2.10 ± 0.44 | 15 | 2.49 ± 0.57 | 14 | 1.88 ± 0.43 | 12 | 4.10 ± 0.94 | 12 | -1.95 [-31.25, 27.95] | -7.33 [-36.64, 22.07] | -11.2 [-54.68, 31.63] | | | | | | |
| R 1022 | R 818 | R Globus pallidus, external segment | 4.07 ± 1.02 | 15 | 6.94 ± 1.94 | 14 | 5.64 ± 1.41 | 11 | 13.23 ± 2.57 | 12 | -1.45 [-31.71, 28.85] | -6.74 [-37.13, 22.75] | -11.17 [-55.41, 32.86] | | | | | | |
| L 1022 | L 818 | L Globus pallidus, external segment | 4.04 ± 0.95 | 14 | 7.01 ± 1.65 | 14 | 4.53 ± 1.53 | 12 | 9.86 ± 2.12 | 12 | -1.72 [-31.81, 29.07] | -6.84 [-36.92, 23.37] | -11.53 [-56.58, 32.63] | | | | | | |
| R 1031 | R 818 | R Globus pallidus, internal segment | 3.73 ± 2.32 | 14 | 3.12 ± 1.18 | 13 | 1.47 ± 0.71 | 11 | 2.20 ± 1.09 | 12 | -2.44 [-34.61, 28.37] | -7.8 [-38.53, 23.45] | -11.78 [-57.64, 33] | | | | | | |
| L 1031 | L 818 | L Globus pallidus, internal segment | 5.93 ± 1.57 | 13 | 1.71 ± 0.74 | 14 | 2.90 ± 1.18 | 12 | 4.29 ± 1.57 | 11 | -2.05 [-33.41, 29.36] | -7.85 [-38.57, 23.2] | -11.64 [-55.54, 33.5] | | | | | | |
| R 835 | R 803 | R Pallidum, ventral region | 3.73 ± 0.90 | 15 | 5.28 ± 1.46 | 15 | 3.04 ± 0.84 | 12 | 9.73 ± 3.03 | 12 | -1.93 [-31.56, 27.58] | -7.11 [-36.41, 22.54] | -10.85 [-53.3, 32.37] | | | | | | |
| L 835 | L 803 | L Pallidum, ventral region | 3.82 ± 0.86 | 15 | 5.14 ± 1.59 | 15 | 4.55 ± 0.79 | 12 | 11.38 ± 3.20 | 12 | -1.47 [-31.77, 28.96] | -7.13 [-36.82, 22.51] | -10.76 [-53.47, 32.92] | | | | | | |
| R 342 | R 835 | R Substantia innominata | 7.29 ± 1.90 | 15 | 10.69 ± 2.93 | 15 | 6.66 ± 1.77 | 11 | 21.87 ± 6.41 | 11 | -2.13 [-33.51, 28.97] | -7.09 [-36.66, 22.42] | -10.56 [-54.62, 33.34] | | | | | | |
| L 342 | L 835 | L Substantia innominata | 7.75 ± 1.78 | 15 | 10.80 ± 3.38 | 14 | 9.40 ± 1.57 | 12 | 25.70 ± 7.04 | 11 | -1.63 [-31.83, 28.49] | -6.89 [-36.98, 23.52] | -10.47 [-53.47, 33.46] | | | | | | |
| R 826 | R 803 | R Pallidum, medial region | 4.45 ± 1.87 | 14 | 7.85 ± 2.83 | 12 | 4.13 ± 1.32 | 11 | 5.12 ± 1.25 | 11 | -1.79 [-32.99, 29.74] | -6.65 [-37.92, 25.18] | -12.87 [-58.57, 33.68] | | | | | | |
| L 826 | L 803 | L Pallidum, medial region | 2.59 ± 1.01 | 14 | 3.62 ± 1.07 | 11 | 6.12 ± 2.18 | 11 | 7.46 ± 1.69 | 11 | -0.39 [-31.39, 31.49] | -7.17 [-39.29, 25.09] | -12.29 [-58.62, 33.08] | | | | | | |
| R 904 | R 826 | R Medial septal complex | 6.67 ± 2.81 | 14 | 10.27 ± 4.10 | 12 | 5.84 ± 1.95 | 11 | 5.71 ± 1.39 | 11 | -1.54 [-32.4, 29.8] | -6.72 [-39.16, 25.2] | -13.24 [-58.47, 32.4] | | | | | | |
| L 904 | L 826 | L Medial septal complex | 3.89 ± 1.52 | 14 | 5.43 ± 1.60 | 11 | 7.90 ± 3.26 | 11 | 10.40 ± 2.17 | 11 | -1.01 [-32.02, 30.27] | -7.25 [-39.9, 24.99] | -11.75 [-57.31, 35.31] | | | | | | |
| R 564 | R 904 | R Medial septal nucleus | 0.00 ± 0.00 | 9 | 9.07 ± 2.48 | 6 | 2.85 ± 1.46 | 8 | 22.38 ± 7.90 | 8 | 26.79 [-10.89, 63.91] | 83.21 [41.39, 125.66]* | 92.36 [36.15, 149.33]* | 90.65 [-131.96, -50.11] | -28.73 [-66.17, 7.9] | 224.16 [-262.44, -186.92] | -62.25 [-102.59, -19.03]* | -133.6 [-174.89, -91.15]* | -195.42 [-233.17, -154.74]* |
| L 564 | L 904 | L Medial septal nucleus | 0.00 ± 0.00 | 8 | 12.46 ± 5.86 | 6 | 7.26 ± 3.70 | 8 | 26.15 ± 12.72 | 8 | 70.97 [31.84, 110.18]* | 117 [73.64, 159.78]* | 52.02 [-6.47, 109.94] | 124.47 [-167.21, -83.36] | -72.81 [-111.67, -34.35]* | 261.94 [-299.99, -222.68] | -51.79 [-95.44, -11.74]* | 137.17 [-178.18, -94.54] | -188.95 [-228.38, -151.22]* |
| R 596 | R 904 | R Diagonal band nucleus | 14.21 ± 5.68 | 14 | 20.28 ± 8.27 | 12 | 12.99 ± 4.01 | 11 | 7.69 ± 2.65 | 11 | -2.03 [-33.64, 29.22] | -7.17 [-39.19, 23.79] | -12.88 [-57.79, 33.44] | | | | | | |
| L 596 | L 904 | L Diagonal band nucleus | 8.83 ± 3.37 | 14 | 9.98 ± 3.28 | 11 | 15.12 ± 6.62 | 11 | 20.50 ± 4.05 | 11 | -1.07 [-33.07, 29.94] | -7.35 [-39.53, 24.95] | -11.91 [-56.8, 34.36] | | | | | | |
| R 549 | R 1129 | R Thalamus | 2.84 ± 0.73 | 15 | 3.20 ± 0.60 | 16 | 4.49 ± 1.05 | 12 | 7.52 ± 1.26 | 12 | -0.19 [-2.76, 2.37] | 0.51 [-2.04, 2.97] | -0.2 [-3.84, 3.48] | | | | | | |
| L 549 | L 1129 | L Thalamus | 2.67 ± 0.55 | 14 | 3.03 ± 0.56 | 16 | 5.07 ± 1.15 | 12 | 7.56 ± 1.14 | 12 | 0.27 [-2.31, 2.82] | 0.69 [-1.84, 3.21] | -0.52 [-4.18, 3.02] | | | | | | |
| R 864 | R 549 | R Thalamus, sensory-motor cortex related | 2.26 ± 0.51 | 15 | 2.96 ± 0.75 | 16 | 5.72 ± 1.42 | 12 | 9.68 ± 1.91 | 12 | 0.66 [-1.93, 3.24] | 0.46 [-2.11, 3] | -0.17 [-3.8, 3.46] | | | | | | |
| L 864 | L 549 | L Thalamus, sensory-motor cortex related | 2.40 ± 0.60 | 14 | 2.80 ± 0.59 | 16 | 6.25 ± 1.52 | 12 | 9.21 ± 1.51 | 12 | 1.05 [-1.47, 3.64] | 1.15 [-1.38, 3.69] | -1.03 [-4.69, 2.52] | | | | | | |
| R 637 | R 864 | R Ventral group of the dorsal thalamus | 2.48 ± 0.55 | 15 | 3.19 ± 0.72 | 16 | 6.52 ± 1.52 | 12 | 13.07 ± 2.64 | 12 | 0.61 [-1.94, 3.13] | 0.2 [-2.27, 2.7] | 0.28 [-3.35, 3.8] | | | | | | |
| L 637 | L 864 | L Ventral group of the dorsal thalamus | 2.47 ± 0.49 | 14 | 3.26 ± 0.62 | 16 | 7.26 ± 1.86 | 12 | 11.95 ± 2.05 | 12 | 1.33 [-1.23, 3.95] | 1.54 [-1.04, 4.03] | -1.29 [-4.95, 2.37] | | | | | | |
| R 629 | R 637 | R Ventral anterior-lateral complex of the th | 2.80 ± 0.81 | 15 | 2.11 ± 0.87 | 14 | 11.25 ± 2.71 | 12 | 30.56 ± 7.90 | 12 | 1.93 [-0.6, 4.52] | -1.06 [-3.61, 1.48] | 1.71 [-2.02, 5.27] | | | | | | |
| L 629 | L 637 | L Ventral anterior-lateral complex of the th | 3.02 ± 1.04 | 14 | 4.92 ± 1.32 | 14 | 11.96 ± 2.48 | 12 | 26.00 ± 8.50 | 12 | 3.44 [0.88, 6.06]* | 1.73 [-0.86, 4.27] | -1.48 [-5.26, 2.05] | -1.56 [-3.8, 0.56] | -4.38 [-6.56, -2.09]* | -5.12 [-7.53, -2.97]* | 2.79 [0.51, 5.13]* | -3.58 [-5.83, -1.25]* | -0.76 [-3.2, 1.68] |
| R 685 | R 637 | R Ventral medial nucleus of the thalamus | 6.40 ± 2.88 | 15 | 6.00 ± 1.37 | 15 | 9.16 ± 3.45 | 12 | 13.05 ± 3.01 | 12 | 1.17 [-1.36, 3.7] | 1.58 [-0.91, 4.08] | -1.32 [-4.95, 2.22] | | | | | | |
| L 685 | L 637 | L Ventral medial nucleus of the thalamus | 5.75 ± 2.19 | 14 | 5.87 ± 1.38 | 15 | 14.30 ± 5.40 | 12 | 12.47 ± 2.51 | 12 | 1.95 [-0.63, 4.53] | 1.75 [-0.81, 4.3] | -2.93 [-6.63, 0.68] | | | | | | |
| R 709 | R 637 | R Ventral posterior complex of the thalamu | 3.80 ± 0.75 | 15 | 5.26 ± 1.23 | 16 | 10.82 ± 2.53 | 12 | 21.24 ± 4.27 | 12 | 0.7 [-1.83, 3.27] | 0.02 [-2.48, 2.55] | 0.47 [-3.18, 4.05] | | | | | | |
| L 709 | L 637 | L Ventral posterior complex of the thalamu | 4.24 ± 0.78 | 13 | 4.95 ± 1.07 | 16 | 11.39 ± 3.00 | 12 | 19.35 ± 3.19 | 12 | 1.12 [-1.47, 3.76] | 1.23 [-1.34, 3.84] | -0.93 [-4.63, 2.67] | | | | | | |
| R 718 | R 709 | R Ventral posterolateral nucleus of the tha | 2.83 ± 0.57 | 15 | 5.78 ± 1.58 | 16 | 13.50 ± 3.51 | 12 | 28.98 ± 6.36 | 12 | 0.82 [-1.68, 3.41] | -0.16 [-2.67, 2.33] | 1.7 [-2.03, 5.32] | | | | | | |
| L 718 | L 709 | L Ventral posterolateral nucleus of the thal | 4.68 ± 1.51 | 13 | 6.86 ± 1.64 | 16 | 12.00 ± 2.17 | 12 | 29.05 ± 4.85 | 12 | 1.71 [-0.9, 4.36] | 1.21 [-1.37, 3.73] | -0.12 [-3.79, 3.47] | | | | | | |
| R 725 | R 709 | R Ventral posterolateral nucleus of the tha | 0.00 ± 0.00 | 11 | 3.93 ± 1.98 | 13 | 5.49 ± 2.80 | 8 | 31.11 ± 11.81 | 7 | 3.9 [1.08, 6.77]* | 3.8 [1.09, 6.43]* | -0.15 [-4.22, 3.9] | -3.65 [-5.98, -1.39]* | -4.81 [-7.42, -2.29]* | -8.97 [-11.59, -6.27]* | 1.17 [-1.32, 3.69] | -5.34 [-7.9, -2.72]* | -4.14 [-7.07, -1.32]* |
| L 725 | L 709 | L Ventral posterolateral nucleus of the thal | 1.22 ± 1.22 | 11 | 5.75 ± 3.38 | 12 | 23.15 ± 9.71 | 10 | 10.19 ± 4.66 | 8 | 4.73 [1.96, 7.54]* | 2.12 [-0.59, 4.8] | -4.27 [-8.25, -0.28]* | -1.97 [-4.41, 0.38] | -5.68 [-8.12, -3.08]* | -4.01 [-6.68, -1.43]* | 3.7 [1.26, 6.13]* | -2.06 [-4.58, 0.46] | 1.65 [-1.14, 4.22] |
| R 733 | R 709 | R Ventral posteromedial nucleus of the tha | 12.83 ± 2.68 | 14 | 14.15 ± 3.52 | 16 | 26.58 ± 6.03 | 12 | 52.29 ± 11.45 | 12 | 0.37 [-2.23, 3.01] | -0.22 [-2.8, 2.34] | 0.65 [-3.01, 4.2] | | | | | | |
| L 733 | L 709 | L Ventral posteromedial nucleus of the tha | 11.35 ± 1.98 | 13 | 11.51 ± 2.65 | 16 | 30.21 ± 9.60 | 12 | 46.32 ± 7.79 | 12 | 1.04 [-1.48, 3.68] | 1.07 [-1.5, 3.54] | -0.81 [-4.49, 2.77] | | | | | | |
| R 741 | R 709 | R Ventral posteromedial nucleus of the tha | 1.81 ± 1.81 | 9 | 4.52 ± 2.25 | 12 | 7.90 ± 3.37 | 10 | 13.43 ± 4.18 | 9 | 3.8 [1, 6.59]* | 2.71 [-0.09, 5.52] | -2.51 [-6.51, 1.41] | -2.57 [-4.91, 0] | -4.72 [-7.29, -2.25]* | -5.46 [-7.97, -2.68]* | 2.16 [-0.22, 4.62] | -2.89 [-5.43, -0.45]* | -0.7 [-3.28, 1.94] |
| L 741 | L 709 | L Ventral posteromedial nucleus of the tha | 6.37 ± 2.90 | 8 | 5.80 ± 2.60 | 14 | 8.67 ± 4.39 | 10 | 10.89 ± 3.67 | 9 | -0.66 [-3.54, 2.18] | 0.4 [-2.41, 3.16] | 1.51 [-2.43, 5.45] | | | | | | |
| R 563807 | R 637 | R Posterior triangular thalamic nucleus | 11.86 ± 8.67 | 9 | 2.38 ± 1.74 | 10 | 12.81 ± 4.87 | 10 | 22.75 ± 8.19 | 7 | 0.55 [-2.29, 3.37] | -2.01 [-4.9, 0.86] | 3.18 [-1, 7.31] | | | | | | |
| L 563807 | L 637 | L Posterior triangular thalamic nucleus | 4.14 ± 1.71 | 9 | 11.10 ± 5.69 | 8 | 10.23 ± 5.49 | 10 | 19.08 ± 9.12 | 7 | -0.72 [-3.56, 2.14] | 0.48 [-2.52, 3.45] | 1.26 [-2.91, 5.35] | | | | | | |
| R 406 | R 864 | R Subparafascicular nucleus | 12.59 ± 4.03 | 11 | 5.51 ± 2.41 | 14 | 28.21 ± 9.99 | 11 | 18.92 ± 3.47 | 8 | 0.95 [-1.77, 3.64] | -1.98 [-4.67, 0.7] | 1.75 [-2.19, 5.55] | | | | | | |
| L 406 | L 864 | L Subparafascicular nucleus | 11.22 ± 3.51 | 11 | 10.21 ± 3.64 | 13 | 8.89 ± 3.13 | 11 | 14.14 ± 2.09 | 8 | -0.21 [-2.89, 2.47] | -0.37 [-3.03, 2.31] | 1.16 [-2.77, 4.91] | | | | | | |
| R 422 | R 406 | R Subparafascicular nucleus, parvicellular | 21.07 ± 9.39 | 9 | 8.42 ± 4.36 | 10 | 35.70 ± 19.07 | 9 | 18.48 ± 9.30 | 7 | -0.06 [-2.95, 2.87] | -2.07 [-4.98, 0.87] | 0.76 [-3.52, 4.99] | | | | | | |
| L 422 | L 406 | L Subparafascicular nucleus, parvicellular | 17.85 ± 9.94 | 9 | 10.97 ± 5.16 | 9 | 9.39 ± 4.65 | 8 | 25.45 ± 4.49 | 7 | -0.36 [-3.24, 2.58] | 0.38 [-2.51, 3.31] | 2.27 [-1.95, 6.44] | | | | | | |
| R 1008 | R 864 | R Geniculate group, dorsal thalamus | 5.05 ± 1.82 | 14 | 5.70 ± 1.92 | 16 | 11.58 ± 3.38 | 12 | 14.85 ± 3.57 | 11 | 1.59 [-0.97, 4.24] | 0.61 [-1.99, 3.14] | -1.46 [-5.23, 2.2] | | | | | | |
| L 1008 | L 864 | L Geniculate group, dorsal thalamus | 6.10 ± 3.08 | 14 | 4.43 ± 1.32 | 16 | 13.85 ± 3.28 | 12 | 16.86 ± 3.23 | 12 | 1.73 [-0.89, 4.28] | 0.55 [-2, 3.01] | -0.79 [-4.37, 2.82] | | | | | | |
| R 475 | R 1008 | R Medial geniculate complex | 1.34 ± 0.64 | 8 | 0.96 ± 0.64 | 9 | 4.29 ± 1.54 | 10 | 15.94 ± 4.58 | 7 | 1.15 [-1.69, 4.01] | -1.51 [-4.52, 1.45] | 4.78 [0.58, 8.9]* | 1.66 [-0.98, 4.31] | -2.06 [-4.71, 0.45] | -5.87 [-8.65, -3.04]* | 3.73 [1.15, 6.27]* | -7.52 [-10.31, -4.77]* | -3.79 [-6.48, -0.97]* |
| L 475 | L 1008 | L Medial geniculate complex | 5.35 ± 3.32 | 8 | 2.34 ± 0.88 | 9 | 6.03 ± 2.30 | 10 | 17.27 ± 4.69 | 7 | 0.15 [-2.77, 3.03] | 0.3 [-2.7, 3.29] | 2.79 [-1.47, 6.99] | | | | | | |
| R 1072 | R 475 | R Medial geniculate complex, dorsal part | 0.00 ± 0.00 | 6 | 4.26 ± 4.26 | 8 | 10.30 ± 7.29 | 9 | 41.68 ± 13.32 | 6 | 2.36 [-0.71, 5.42] | 1.65 [-1.52, 4.79] | 7.11 [2.71, 11.46]* | -1.49 [-4.27, 1.47] | -3.27 [-6.24, -0.66]* | -12.58 [-15.59, -9.48]* | 1.8 [-1.01, 4.3] | -11.09 [-13.99, -8.1]* | -9.28 [-12.17, -6.37]* |
| L 1072 | L 475 | L Medial geniculate complex, dorsal part | 14.98 ± 12.66 | 7 | 3.88 ± 2.67 | 7 | 11.77 ± 8.44 | 8 | 14.52 ± 6.65 | 7 | -0.4 [-3.42, 2.61] | -0.7 [-3.87, 2.47] | 1.73 [-2.66, 6.02] | | | | | | |
| R 1079 | R 475 | R Medial geniculate complex, ventral part | 0.00 ± 0.00 | 6 | 4.51 ± 3.18 | 7 | 7.70 ± 3.49 | 9 | 36.96 ± 15.52 | 6 | 4.79 [1.66, 7.97]* | 3.03 [-0.3, 6.25] | 1.04 [-3.53, 5.53] | -2.88 [-5.79, 0.13] | -5.72 [-8.69, -2.97]* | -10.3 [-13.43, -7.24]* | 2.84 [0.11, 5.57]* | -7.43 [-10.55, -4.43]* | -4.58 [-7.38, -1.64]* |
| L 1079 | L 475 | L Medial geniculate complex, ventral part | 21.61 ± 12.70 | 6 | 7.03 ± 5.93 | 7 | 11.78 ± 6.94 | 8 | 40.79 ± 16.25 | 7 | -1.43 [-4.54, 1.67] | -2.43 [-5.65, 0.76] | 4.09 [-0.29, 8.44] | | | | | | |
| R 1088 | R 475 | R Medial geniculate complex, medial part | 4.76 ± 2.02 | 8 | 0.00 ± 0.00 | 9 | 9.48 ± 3.92 | 10 | 27.83 ± 6.34 | 7 | 0.76 [-2.06, 3.63] | -3.37 [-6.25, -0.42]* | 5.43 [1.23, 9.49]* | 3.54 [1, 6.28]* | -1.68 [-4.17, 1.05] | -4.27 [-7.16, -1.57]* | 5.21 [2.57, 7.73]* | -7.79 [-10.51, -5.11]* | -2.59 [-5.28, 0.2] |
| L 1088 | L 475 | L Medial geniculate complex, medial part | 2.93 ± 2.03 | 8 | 4.02 ± 1.93 | 9 | 7.21 ± 3.52 | 10 | 36.22 ± 7.04 | 7 | 1.2 [-1.63, 4.13] | 1.58 [-1.31, 4.53] | 3.37 [-0.82, 7.5] | | | | | | |
| R 170 | R 1008 | R Dorsal part of the lateral geniculate com | 8.40 ± 3.21 | 14 | 9.79 ± 3.20 | 16 | 21.62 ± 6.26 | 12 | 27.31 ± 6.97 | 11 | 2.15 [-0.43, 4.8] | 1.08 [-1.43, 3.62] | -2.03 [-5.86, 1.6] | | | | | | |
| L 170 | L 1008 | L Dorsal part of the lateral geniculate com | 9.03 ± 4.67 | 14 | 8.05 ± 2.63 | 16 | 26.75 ± 6.24 | 12 | 30.01 ± 6.23 | 12 | 2.39 [-0.18, 4.98] | 1 [-1.53, 3.51] | -1.43 [-5.06, 2.16] | | | | | | |
| R 496345 | R 170 | R Dorsal part of the lateral geniculate com | 7.16 ± 4.58 | 13 | 23.59 ± 8.53 | 15 | 31.79 ± 11.54 | 12 | 35.23 ± 10.70 | 11 | 1.96 [-0.68, 4.6] | 3.3 [0.64, 5.84]* | -1.92 [-5.63, 1.77] | -3.14 [-5.33, -0.95]* | -2.9 [-5.16, -0.56]* | -4.79 [-7.08, -2.33]* | -0.24 [-2.48, 2.03] | -1.64 [-4.01, 0.6] | -1.9 [-4.38, 0.51] |
| L 496345 | L 170 | L Dorsal part of the lateral geniculate com | 17.45 ± 13.38 | 14 | 15.57 ± 6.56 | 15 | 37.03 ± 14.93 | 11 | 58.73 ± 19.99 | 11 | 1.28 [-1.39, 3.86] | 1.12 [-1.53, 3.66] | 0.08 [-3.63, 3.83] | | | | | | |
| R 496345 | R 170 | R Dorsal part of the lateral geniculate com | 21.04 ± 8.53 | 14 | 18.07 ± 5.92 | 16 | 48.53 ± 18.07 | 12 | 60.55 ± 15.11 | 11 | 1.99 [-0.61, 4.65] | 0.89 [-1.61, 3.41] | -1.83 [-5.56, 1.86] | | | | | | |
| L 496345 | L 170 | L Dorsal part of the lateral geniculate com | 19.42 ± 8.71 | 14 | 18.44 ± 5.48 | 16 | 58.75 ± 15.40 | 12 | 68.44 ± 13.65 | 12 | 2.32 [-0.25, 4.94] | 0.7 [-1.8, 3.23] | -1.08 [-4.71, 2.42] | | | | | | |
| R 496345 | R 170 | R Dorsal part of the lateral geniculate com | 16.65 ± 8.15 | 12 | 21.86 ± 8.26 | 15 | 42.81 ± 14.00 | 12 | 64.98 ± 24.98 | 11 | 1.42 [-1.19, 4.02] | 1.7 [-0.95, 4.3] | -0.88 [-4.61, 2.74] | | | | | | |
| L 496345 | L 170 | L Dorsal part of the lateral geniculate com | 18.23 ± 9.86 | 12 | 11.84 ± 5.57 | 16 | 51.56 ± 15.07 | 11 | 48.48 ± 13.97 | 11 | 2.08 [-0.62, 4.73] | 0 [-2.58, 2.53] | -0.42 [-4.21, 3.18] | | | | | | |
| R 856 | R 549 | R Thalamus, polymodal association cortex | 4.33 ± 1.14 | 15 | 4.67 ± 0.83 | 16 | 5.83 ± 1.34 | 12 | 9.64 ± 1.52 | 12 | -0.37 [-2.9, 2.23] | 0.47 [-2.04, 2.94] | -0.15 [-3.73, 3.46] | | | | | | |
| L 856 | L 549 | L Thalamus, polymodal association cortex | 3.82 ± 0.75 | 15 | 4.49 ± 0.83 | 16 | 6.55 ± 1.42 | 12 | 10.01 ± 1.55 | 12 | -0.03 [-2.59, 2.57] | 0.55 [-1.99, 3.08] | -0.36 [-3.99, 3.26] | | | | | | |
| R 138 | R 856 | R Lateral group of the dorsal thalamus | 1.91 ± 0.44 | 14 | 2.70 ± 0.72 | 16 | 6.32 ± 1.77 | 12 | 15.60 ± 2.89 | 12 | 1.01 [-1.53, 3.64] | 0.18 [-2.37, 2.69] | 0.66 [-3.04, 4.33] | | | | | | |
| L 138 | L 856 | L Lateral group of the dorsal thalamus | 1.53 ± 0.36 | 14 | 2.62 ± 0.68 | 16 | 6.35 ± 1.46 | 12 | 14.76 ± 2.26 | 12 | 2.1 [-0.49, 4.68] | 1.43 [-1.13, 3.97] | -0.72 [-4.36, 2.87] | | | | | | |
| R 218 | R 138 | R Lateral posterior nucleus of the thalamus | 4.77 ± 1.37 | 14 | 5.76 ± 1.77 | 16 | 12.85 ± 3.02 | 12 | 22.53 ± 4.21 | 12 | 1.34 [-1.28, 3.88] | 0.54 [-2.04, 3.07] | -0.25 [-3.98, 3.36] | | | | | | |
| L 218 | L 138 | L Lateral posterior nucleus of the thalamus | 3.64 ± 0.88 | 14 | 7.45 ± 2.19 | 16 | 14.30 ± 3.64 | 12 | 24.49 ± 3.91 | 12 | 1.96 [-0.58, 4.54] | 0.86 [-1.71, 3.39] | -0.38 [-4.13, 3.15] | | | | | | |
| R 1020 | R 138 | R Posterior complex of the thalamus | 2.33 ± 0.69 | 14 | 5.69 ± 2.87 | 16 | 13.38 ± 4.98 | 12 | 40.32 ± 8.78 | 12 | 2.49 [-0.1, 5.12] | 1.31 [-1.26, 3.82] | -0.28 [-3.94, 3.37] | | | | | | |
| L 1020 | L 138 | L Posterior complex of the thalamus | 1.71 ± 0.68 | 14 | 3.71 ± 1.03 | 16 | 10.88 ± 1.92 | 12 | 37.87 ± 6.97 | 12 | 3.14 [0.55, 5.72]* | 2.04 [-0.53, 4.56] | -0.59 [-4.22, 3.04] | -1.89 [-3.97, 0.27] | -4.08 [-6.3, -1.79]* | -6.02 [-8.33, -3.68]* | 2.19 [0.05, 4.52]* | -4.15 [-6.35, -1.86]* | -1.97 [-4.34, 0.51] |
| R 1029 | R 138 | R Posterior limiting nucleus of the thalamu | 20.44 ± 8.29 | 10 | 7.34 ± 3.74 | 10 | 12.48 ± 5.15 | 10 | 14.38 ± 5.68 | 8 | -0.89 [-3.6, 1.88] | -1.78 [-4.64, 1] | 1.86 [-2.18, 5.74] | | | | | | |
| L 1029 | L 138 | L Posterior limiting nucleus of the thalamus | 4.61 ± 2.37 | 11 | 1.65 ± 1.08 | 8 | 16.04 ± 7.65 | 10 | 20.63 ± 9.37 | 7 | 0.87 [-1.88, 3.64] | -1.14 [-4.06, 1.73] | 1.98 [-2.22, 6.03] | | | | | | |
| R 325 | R 138 | R Suprageniculate nucleus | 7.16 ± 4.64 | 6 | 0.00 ± 0.00 | 7 | 0.84 ± 0.84 | 9 | 27.18 ± 15.73 | 6 | -2.39 [-5.51, 0.71] | -3.01 [-6.22, 0.21] | 5.57 [1.15, 9.98]* | 3.16 [0.2, 6.01]* | 1.45 [-1.44, 4.22] | -1.63 [-4.68, 1.45] | 1.7 [-1.13, 4.43] | -4.79 [-7.8, -1.86]* | -3.08 [-6.07, -0.23]* |
| L 325 | L 138 | L Suprageniculate nucleus | 11.73 ± 7.45 | 7 | 2.90 ± 2.90 | 6 | 7.00 ± 5.67 | 8 | 12.08 ± 5.03 | 7 | -1.49 [-4.54, 1.58] | -2.32 [-5.58, 0.91] | 3.56 [-0.95, 7.97] | | | | | | |
| R 560581 | R 138 | R Ethmoid nucleus of the thalamus | 5.04 ± 2.59 | 13 | 4.60 ± 1.93 | 16 | 8.95 ± 3.43 | 12 | 11.13 ± 4.30 | 12 | 0.33 [-2.22, 2.95] | 0.31 [-2.25, 2.84] | 0.04 [-3.61, 3.61] | | | | | | |
| L 560581 | L 138 | L Ethmoid nucleus of the thalamus | 4.24 ± 1.68 | 13 | 3.64 ± 1.81 | 16 | 13.11 ± 7.06 | 12 | 15.87 ± 5.39 | 12 | 0.38 [-2.22, 3.05] | -0.2 [-2.82, 2.39] | 1.41 [-2.35, 5.05] | | | | | | |
| R 239 | R 856 | R Anterior group of the dorsal thalamus | 4.14 ± 1.26 | 15 | 4.15 ± 1.08 | 14 | 4.71 ± 1.02 | 12 | 8.40 ± 1.27 | 12 | -0.18 [-2.71, 2.37] | 0.19 [-2.35, 2.7] | 0.11 [-3.56, 3.71] | | | | | | |
| L 239 | L 856 | L Anterior group of the dorsal thalamus | 3.48 ± 0.86 | 15 | 2.84 ± 0.52 | 14 | 6.35 ± 1.43 | 12 | 9.19 ± 1.57 | 12 | 1.35 [-1.2, 3.89] | 1.22 [-1.38, 3.76] | -1.08 [-4.74, 2.52] | | | | | | |
| R 255 | R 239 | R Anteroventral nucleus of thalamus | 11.55 ± 3.89 | 13 | 8.49 ± 4.41 | 10 | 7.41 ± 2.22 | 12 | 10.15 ± 3.53 | 10 | -0.65 [-3.24, 1.93] | -0.59 [-3.35, 2.08] | 0.11 [-3.72, 3.84] | | | | | | |
| L 255 | L 239 | L Anteroventral nucleus of thalamus | 18.61 ± 6.69 | 12 | 17.17 ± 11.25 | 10 | 5.72 ± 1.52 | 12 | 14.95 ± 9.01 | 10 | -1.31 [-3.91, 1.34] | 0.25 [-2.54, 2.97] | -0.48 [-4.42, 3.39] | | | | | | |
| R 127 | R 239 | R Anteromedial nucleus | 3.31 ± 1.75 | 13 | 2.19 ± 0.82 | 9 | 3.75 ± 1.11 | 12 | 2.57 ± 1.34 | 10 | 0.3 [-2.3, 2.95] | 0.22 [-2.51, 2.97] | -2.97 [-6.84, 0.83] | | | | | | |
| L 127 | L 239 | L Anteromedial nucleus | 2.65 ± 0.96 | 13 | 1.23 ± 0.57 | 10 | 2.47 ± 0.79 | 12 | 4.01 ± 1.73 | 10 | -1.81 [-4.41, 0.84] | -1.64 [-4.38, 1.11] | 2.73 [-1.16, 6.56] | | | | | | |
| R 1096 | R 127 | R Anteromedial nucleus, dorsal part | 6.83 ± 4.66 | 13 | 4.41 ± 2.37 | 9 | 4.20 ± 1.70 | 12 | 5.42 ± 3.10 | 10 | -1.1 [-3.75, 1.48] | -0.14 [-2.94, 2.65] | -0.25 [-4.17, 3.51] | | | | | | |
| L 1096 | L 127 | L Anteromedial nucleus, dorsal part | 7.06 ± 2.75 | 13 | 2.18 ± 1.16 | 10 | 6.29 ± 2.66 | 12 | 7.36 ± 3.96 | 10 | -2.46 [-5.08, 0.19] | -2.45 [-5.17, 0.24] | 2.6 [-1.27, 6.33] | | | | | | |
| R 1104 | R 127 | R Anteromedial nucleus, ventral part | 6.14 ± 2.36 | 13 | 4.16 ± 2.65 | 9 | 12.98 ± 4.40 | 12 | 4.68 ± 2.41 | 10 | 1.27 [-1.36, 3.9] | -1.23 [-4.03, 1.52] | -2.52 [-6.44, 1.34] | | | | | | |
| L 1104 | L 127 | L Anteromedial nucleus, ventral part | 2.77 ± 1.31 | 13 | 2.81 ± 1.57 | 10 | 2.56 ± 1.35 | 12 | 10.62 ± 5.43 | 9 | -1.48 [-4.11, 1.17] | -0.04 [-2.82, 2.63] | 2.51 [-1.4, 6.33] | | | | | | |
| R 64 | R 239 | R Anterodorsal nucleus | 15.70 ± 8.13 | 12 | 11.60 ± 2.75 | 10 | 5.27 ± 2.17 | 11 | 11.75 ± 5.85 | 10 | -1.59 [-4.35, 1.11] | 0.91 [-1.89, 3.68] | -1.69 [-5.57, 2.13] | | | | | | |
| L 64 | L 239 | L Anterodorsal nucleus | 11.58 ± 3.18 | 11 | 4.07 ± 2.52 | 10 | 4.66 ± 1.87 | 11 | 13.38 ± 4.30 | 9 | -2.67 [-5.36, 0.05] | -2.58 [-5.42, 0.18] | 4.34 [0.36, 8.26]* | 2.73 [0.24, 5.17]* | 1.74 [-0.57, 4.28] | -0.54 [-3.06, 1.96] | 0.98 [-1.37, 3.53] | -3.27 [-5.89, -0.74]* | -2.28 [-4.76, 0.37] |
| R 1120 | R 239 | R Interanteromedial nucleus of the thalamu | 29.66 ± 14.46 | 12 | 18.62 ± 5.17 | 9 | 30.89 ± 24.71 | 10 | 16.69 ± 9.48 | 9 | -1.98 [-4.68, 0.68] | 1.35 [-1.44, 4.12] | -1.21 [-5.18, 2.69] | | | | | | |
| L 1120 | L 239 | L Interanteromedial nucleus of the thalamu | 19.96 ± 8.03 | 11 | 14.30 ± 7.61 | 9 | 29.34 ± 16.90 | 10 | 31.85 ± 9.77 | 9 | -1.53 [-4.27, 1.19] | -0.4 [-3.2, 2.38] | 1.72 [-2.25, 5.62] | | | | | | |
| R 1113 | R 239 | R Interanterodorsal nucleus of the thalamu | 37.85 ± 12.13 | 12 | 12.36 ± 8.18 | 9 | 23.33 ± 6.38 | 12 | 34.43 ± 14.13 | 10 | -1.26 [-3.84, 1.38] | -3.64 [-6.46, -0.83]* | 2.76 [-1.11, 6.63] | 3.79 [1.34, 6.25]* | 0.34 [-1.99, 2.69] | 0.69 [-1.84, 3.13] | 3.47 [1.01, 5.96]* | -3.1 [-5.66, -0.49]* | 0.36 [-2.08, 2.89] |
| L 1113 | L 239 | L Interanterodorsal nucleus of the thalamu | 26.86 ± 7.53 | 12 | 10.11 ± 4.04 | 10 | 44.01 ± 13.80 | 12 | 23.73 ± 9.49 | 9 | -1.02 [-3.64, 1.58] | -2.26 [-4.96, 0.42] | 1.08 [-2.82, 4.93] | | | | | | |
| R 155 | R 239 | R Lateral dorsal nucleus of thalamus | 3.97 ± 1.01 | 15 | 10.51 ± 4.64 | 14 | 11.37 ± 3.04 | 12 | 23.01 ± 3.56 | 12 | 0.78 [-1.76, 3.36] | 0.52 [-2.06, 3.08] | 0.5 [-3.22, 4.06] | | | | | | |
| L 155 | L 239 | L Lateral dorsal nucleus of thalamus | 3.83 ± 1.44 | 15 | 5.77 ± 1.45 | 14 | 16.65 ± 4.23 | 12 | 22.12 ± 4.55 | 12 | 2.08 [-0.49, 4.64] | 1.31 [-1.24, 3.83] | -0.66 [-4.38, 3.02] | | | | | | |
| R 444 | R 856 | R Medial group of the dorsal thalamus | 5.91 ± 1.46 | 15 | 6.57 ± 1.47 | 15 | 8.56 ± 2.26 | 12 | 12.70 ± 2.41 | 12 | 0.02 [-2.54, 2.6] | 0.59 [-1.97, 3.07] | -0.38 [-4.02, 3.22] | | | | | | |
| L 444 | L 856 | L Medial group of the dorsal thalamus | 5.96 ± 1.29 | 15 | 6.53 ± 1.22 | 15 | 11.16 ± 2.74 | 12 | 13.02 ± 2.40 | 12 | 0.02 [-2.53, 2.6] | 0.64 [-1.87, 3.11] | -0.77 [-4.41, 2.78] | | | | | | |
| R 59 | R 444 | R Intermediodorsal nucleus of the thalamu | 24.04 ± 6.07 | 14 | 31.81 ± 6.36 | 15 | 40.41 ± 8.20 | 12 | 31.86 ± 7.55 | 12 | 0.94 [-1.72, 3.54] | 0.83 [-1.72, 3.38] | -1.81 [-5.42, 1.83] | | | | | | |
| L 59 | L 444 | L Intermediodorsal nucleus of the thalamus | 17.53 ± 4.95 | 14 | 28.35 ± 7.05 | 15 | 46.41 ± 14.84 | 12 | 35.06 ± 8.03 | 12 | 1.05 [-1.52, 3.74] | 2.13 [-0.49, 4.67] | -2.43 [-6.13, 1.2] | | | | | | |
| R 362 | R 444 | R Mediodorsal nucleus of thalamus | 10.70 ± 2.83 | 15 | 8.92 ± 2.11 | 15 | 14.33 ± 4.81 | 12 | 21.80 ± 4.63 | 12 | -0.28 [-2.81, 2.24] | 0.23 [-2.35, 2.79] | 0.55 [-3.18, 4.12] | | | | | | |
| L 362 | L 444 | L Mediodorsal nucleus of thalamus | 12.01 ± 2.64 | 15 | 8.94 ± 1.51 | 14 | 21.58 ± 5.95 | 12 | 23.56 ± 4.89 | 12 | -0.09 [-2.64, 2.48] | 0.42 [-2.18, 2.9] | -0.55 [-4.22, 3.03] | | | | | | |
| R 366 | R 444 | R Submedial nucleus of the thalamus | 2.94 ± 0.79 | 14 | 6.51 ± 1.53 | 14 | 17.93 ± 5.81 | 12 | 37.27 ± 8.24 | 11 | 0.33 [-2.28, 2.93] | 1.51 [-1.05, 4.07] | 2.11 [-1.61, 5.77] | | | | | | |
| L 366 | L 444 | L Submedial nucleus of the thalamus | 4.73 ± 1.47 | 14 | 9.34 ± 2.28 | 14 | 16.37 ± 5.22 | 11 | 33.61 ± 8.17 | 12 | 1.7 [-0.94, 4.32] | 1.93 [-0.67, 4.47] | -1.55 [-5.26, 2.14] | | | | | | |
| R 1077 | R 444 | R Perireunensis nucleus | 30.30 ± 9.24 | 15 | 14.38 ± 4.29 | 14 | 15.03 ± 5.20 | 12 | 28.50 ± 5.11 | 12 | -1.64 [-4.18, 0.94] | -0.05 [-2.66, 2.48] | 1.82 [-1.88, 5.46] | | | | | | |
| L 1077 | L 444 | L Perireunensis nucleus | 33.24 ± 13.90 | 15 | 21.16 ± 6.68 | 14 | 20.68 ± 7.71 | 12 | 23.80 ± 8.54 | 12 | -1.46 [-4.07, 1.07] | 0.51 [-2.11, 3.08] | -0.22 [-3.85, 3.4] | | | | | | |
| R 571 | R 856 | R Midline group of the dorsal thalamus | 20.60 ± 5.93 | 15 | 19.61 ± 5.68 | 16 | 18.55 ± 3.12 | 12 | 28.07 ± 4.26 | 12 | -0.68 [-3.2, 1.88] | -0.46 [-2.98, 2] | 0.7 [-3, 4.25] | | | | | | |
| L 571 | L 856 | L Midline group of the dorsal thalamus | 16.71 ± 3.47 | 15 | 20.96 ± 5.75 | 16 | 22.16 ± 3.66 | 12 | 29.80 ± 5.67 | 12 | 0.38 [-2.15, 2.97] | 0.12 [-2.44, 2.67] | -0.15 [-3.82, 3.49] | | | | | | |
| R 149 | R 571 | R Paraventricular nucleus of the thalamus | 80.87 ± 18.53 | 15 | 72.73 ± 21.61 | 16 | 81.97 ± 14.86 | 12 | 98.93 ± 16.90 | 12 | -0.49 [-3.05, 2.04] | -0.64 [-3.13, 1.86] | 0.19 [-3.38, 3.75] | | | | | | |

| | | | | | | | | | | | | | | | | | | |
|---|---|---|---|---|---|---|---|---|---|---|---|---|---|---|---|---|---|---|
| L 149 | L 571 | L Paraventricular nucleus of the thalamus | 70.48 ± 15.30 | 15 | 83.54 ± 23.49 | 16 | 104.84 ± 15.81 | 12 | 103.12 ± 20.20 | 12 | 0.47 [-2.05, 3.01] | -0.31 [-2.81, 2.16] | -0.11 [-3.7, 3.45] | | | | | |
| R 15 | R 571 | R Parataenial nucleus | 64.81 ± 26.99 | 10 | 35.41 ± 22.71 | 6 | 48.57 ± 17.58 | 7 | 24.53 ± 18.02 | 6 | -0.76 [-3.72, 2.24] | -1.43 [-4.52, 1.62] | -1.4 [-5.88, 2.98] | | | | | |
| L 15 | L 571 | L Parataenial nucleus | 32.56 ± 9.97 | 10 | 43.41 ± 28.05 | 7 | 24.20 ± 12.26 | 8 | 34.10 ± 22.72 | 7 | -1.94 [-4.81, 0.97] | 0.17 [-2.86, 3.18] | -0.61 [-5.03, 3.57] | | | | | |
| R 181 | R 571 | R Nucleus of reuniens | 14.67 ± 6.28 | 15 | 20.38 ± 5.75 | 14 | 12.19 ± 3.48 | 12 | 29.37 ± 7.72 | 12 | -0.46 [-2.99, 2.14] | 0.41 [-2.13, 2.98] | 0.47 [-3.3, 4.03] | | | | | |
| L 181 | L 571 | L Nucleus of reuniens | 12.58 ± 3.94 | 15 | 17.13 ± 4.44 | 14 | 17.01 ± 5.00 | 12 | 27.06 ± 9.05 | 12 | 0.64 [-1.87, 3.25] | 0.4 [-2.14, 2.95] | -0.26 [-3.95, 3.39] | | | | | |
| R 56058 | R 571 | R Xiphoid thalamic nucleus | 27.49 ± 7.45 | 15 | 31.08 ± 7.51 | 14 | 22.39 ± 6.77 | 12 | 47.78 ± 12.73 | 12 | -0.86 [-3.43, 1.68] | 1.3 [-1.25, 3.86] | -0.47 [-4.12, 3.16] | | | | | |
| L 56058 | L 571 | L Xiphoid thalamic nucleus | 18.05 ± 5.14 | 15 | 44.65 ± 14.15 | 14 | 28.33 ± 8.83 | 12 | 50.01 ± 14.14 | 12 | -0.52 [-3.04, 2.08] | 1.79 [-0.73, 4.27] | -1.17 [-4.83, 2.44] | | | | | |
| R 51 | R 856 | R Intralaminar nuclei of the dorsal thalamu | 7.28 ± 1.84 | 15 | 6.51 ± 1.45 | 16 | 8.27 ± 2.06 | 12 | 14.10 ± 2.70 | 12 | 0.13 [-2.41, 2.7] | 0.47 [-2.08, 3] | -0.31 [-4.02, 3.21] | | | | | |
| L 51 | L 856 | L Intralaminar nuclei of the dorsal thalamus | 6.72 ± 1.51 | 15 | 6.54 ± 2.17 | 16 | 9.63 ± 2.55 | 12 | 14.24 ± 2.25 | 12 | 0.18 [-2.35, 2.75] | -0.59 [-3.07, 1.9] | 0.82 [-2.9, 4.42] | | | | | |
| R 189 | R 51 | R Rhomboid nucleus | 17.21 ± 5.97 | 15 | 21.71 ± 8.29 | 14 | 22.79 ± 8.31 | 12 | 40.50 ± 13.81 | 12 | -0.01 [-2.59, 2.59] | 0.68 [-1.92, 3.21] | -0.09 [-3.72, 3.48] | | | | | |
| L 189 | L 51 | L Rhomboid nucleus | 15.23 ± 4.47 | 15 | 14.24 ± 8.74 | 14 | 28.18 ± 9.49 | 12 | 40.04 ± 10.79 | 12 | 0 [-2.53, 2.58] | -1.22 [-3.77, 1.28] | 1.75 [-1.97, 5.37] | | | | | |
| R 599 | R 51 | R Central medial nucleus of the thalamus | 32.59 ± 7.54 | 15 | 27.98 ± 6.71 | 15 | 32.01 ± 10.18 | 12 | 41.33 ± 10.64 | 12 | -0.14 [-2.69, 2.42] | 0.22 [-2.43, 2.72] | -0.83 [-4.48, 2.79] | | | | | |
| L 599 | L 51 | L Central medial nucleus of the thalamus | 26.49 ± 7.30 | 15 | 24.98 ± 8.35 | 15 | 40.46 ± 14.42 | 12 | 29.16 ± 5.45 | 12 | 0.22 [-2.35, 2.8] | -0.45 [-3, 2.08] | 0.42 [-3.27, 4.01] | | | | | |
| R 907 | R 51 | R Paracentral nucleus | 4.62 ± 2.02 | 14 | 8.84 ± 4.18 | 14 | 7.58 ± 2.96 | 12 | 32.53 ± 7.10 | 12 | -0.55 [-3.15, 2.03] | 0.82 [-1.76, 3.38] | 2.81 [-0.88, 6.47] | | | | | |
| L 907 | L 51 | L Paracentral nucleus | 11.24 ± 4.29 | 14 | 8.11 ± 5.68 | 14 | 12.83 ± 3.18 | 12 | 43.34 ± 14.09 | 12 | 0.06 [-2.46, 2.68] | -1.08 [-3.7, 1.53] | 3.63 [0.02, 7.25]* | 1.23 [-0.97, 3.39] | -0.97 [-3.26, 1.38] | -4.06 [-6.35, -1.71]* | 2.22 [-0.00, 4.52] | -5.29 [-7.63, -3.07]* | -3.06 [-5.41, -0.59]* |
| R 575 | R 51 | R Central lateral nucleus of the thalamus | 10.24 ± 4.95 | 14 | 14.48 ± 4.75 | 15 | 8.33 ± 1.53 | 12 | 21.85 ± 4.31 | 12 | 1.32 [-1.25, 3.92] | 1.36 [-1.23, 3.92] | 0.08 [-3.59, 3.66] | | | | | |
| L 575 | L 51 | L Central lateral nucleus of the thalamus | 8.98 ± 2.14 | 14 | 15.37 ± 5.82 | 15 | 12.86 ± 3.90 | 12 | 23.07 ± 5.08 | 12 | -0.42 [-3.01, 2.19] | -0.11 [-2.63, 2.45] | 1.97 [-1.69, 5.54] | | | | | |
| R 930 | R 51 | R Parafascicular nucleus | 6.97 ± 3.47 | 11 | 5.36 ± 2.22 | 11 | 19.65 ± 6.76 | 10 | 15.05 ± 6.17 | 7 | 3.05 [0.29, 5.84]* | 1.86 [-0.93, 4.53] | -3.93 [-7.97, -0.06]* | -1.7 [-4.07, 0.74] | -3.99 [-6.47, -1.55]* | -2.43 [-5.07, 0.29] | 2.27 [-0.12, 4.85] | -0.72 [-3.29, 2.02] | 1.54 [-1.34, 4.21] |
| L 930 | L 51 | L Parafascicular nucleus | 7.84 ± 3.03 | 10 | 5.04 ± 1.86 | 11 | 15.59 ± 5.87 | 9 | 21.45 ± 6.41 | 7 | 2.19 [-0.64, 5.02] | -0.1 [-2.88, 2.66] | -1.03 [-5.17, 3.04] | | | | | |
| R 262 | R 856 | R Reticular nucleus of the thalamus | 2.46 ± 0.56 | 15 | 7.98 ± 1.89 | 15 | 10.40 ± 3.75 | 12 | 24.72 ± 5.19 | 12 | 0.44 [-2.12, 3.01] | 0.77 [-1.76, 3.29] | 0.46 [-3.25, 4] | | | | | |
| L 262 | L 856 | L Reticular nucleus of the thalamus | 2.86 ± 0.88 | 15 | 6.17 ± 1.88 | 15 | 12.59 ± 4.08 | 12 | 24.32 ± 4.81 | 12 | 1.7 [-0.86, 4.27] | 1.27 [-1.28, 3.77] | -0.9 [-4.57, 2.62] | | | | | |
| R 1014 | R 856 | R Geniculate group, ventral thalamus | 22.45 ± 9.08 | 13 | 18.61 ± 4.93 | 16 | 29.20 ± 7.39 | 12 | 21.30 ± 6.72 | 11 | 0.96 [-1.62, 3.65] | 0.36 [-2.22, 2.85] | -1.94 [-5.62, 1.69] | | | | | |
| L 1014 | L 856 | L Geniculate group, ventral thalamus | 17.71 ± 4.99 | 14 | 15.31 ± 4.82 | 16 | 24.37 ± 7.00 | 12 | 27.64 ± 7.49 | 12 | 1.18 [-1.39, 3.71] | 0.04 [-2.56, 2.59] | -1.15 [-4.87, 2.51] | | | | | |
| R 27 | R 1014 | R Intergeniculate leaflet of the lateral genic | 60.84 ± 24.16 | 9 | 43.19 ± 15.11 | 13 | 109.07 ± 30.72 | 10 | 61.17 ± 23.29 | 8 | 1.32 [-1.6, 4.12] | 0.05 [-2.8, 2.89] | -1.68 [-5.68, 2.31] | | | | | |
| L 27 | L 1014 | L Intergeniculate leaflet of the lateral genic | 45.24 ± 18.11 | 12 | 43.99 ± 17.84 | 11 | 85.14 ± 26.39 | 10 | 63.76 ± 28.60 | 8 | 2.37 [-0.33, 5.1] | 0.44 [-2.3, 3.16] | -3.13 [-7.12, 0.82] | | | | | |
| R 178 | R 1014 | R Ventral part of the lateral geniculate com | 47.75 ± 20.66 | 13 | 40.05 ± 11.19 | 16 | 57.04 ± 15.39 | 12 | 45.07 ± 14.45 | 11 | 0.82 [-1.72, 3.41] | 0.37 [-2.21, 2.9] | -1.8 [-5.49, 1.84] | | | | | |
| L 178 | L 1014 | L Ventral part of the lateral geniculate com | 37.14 ± 10.87 | 14 | 34.44 ± 10.85 | 16 | 49.97 ± 14.77 | 12 | 59.16 ± 15.34 | 12 | 1.11 [-1.49, 3.74] | 0.08 [-2.46, 2.65] | -1.09 [-4.88, 2.52] | | | | | |
| L 321 | L 1014 | L Subgeniculate nucleus | 0.00 ± 0.00 | 8 | 2.17 ± 2.17 | 11 | 3.08 ± 3.08 | 10 | 0.00 ± 0.00 | 7 | 0.51 [-2.38, 3.43] | 0.93 [-1.97, 3.82] | -2.91 [-7.15, 1.15] | | | | | |
| R 958 | R 856 | R Epithalamus | 8.47 ± 1.63 | 15 | 14.20 ± 3.32 | 16 | 7.78 ± 3.47 | 12 | 14.43 ± 2.40 | 12 | -2.27 [-4.81, 0.3] | 0.49 [-1.98, 2.99] | 1.46 [-2.17, 4.98] | | | | | |
| L 958 | L 856 | L Epithalamus | 6.10 ± 1.58 | 15 | 13.65 ± 3.24 | 16 | 7.64 ± 1.84 | 12 | 12.98 ± 1.92 | 12 | -0.28 [-2.81, 2.34] | 1.18 [-1.28, 3.66] | -0.71 [-4.38, 2.83] | | | | | |
| R 483 | R 958 | R Medial habenula | 21.01 ± 5.46 | 15 | 36.92 ± 9.78 | 16 | 13.66 ± 4.30 | 12 | 41.79 ± 8.02 | 12 | -1.94 [-4.54, 0.6] | 1.4 [-1.15, 3.91] | 1.21 [-2.42, 4.8] | | | | | |
| L 483 | L 958 | L Medial habenula | 15.09 ± 5.40 | 15 | 37.64 ± 9.04 | 16 | 18.42 ± 4.55 | 12 | 39.76 ± 6.31 | 12 | 1.23 [-1.27, 3.78] | 2.57 [0.05, 5.08]* | -1.91 [-5.57, 1.64] | -2.42 [-4.56, -0.34]* | -2.16 [-4.4, -0.01]* | -3.32 [-5.54, -1.03]* | -0.25 [-2.51, 1.97] | -0.92 [-3.12, 1.29] | -1.18 [-3.56, 1.18] |
| R 186 | R 958 | R Lateral habenula | 11.89 ± 3.02 | 15 | 17.42 ± 5.83 | 15 | 17.73 ± 11.75 | 12 | 15.47 ± 4.94 | 12 | -1.6 [-4.21, 0.92] | 0.29 [-2.3, 2.8] | 1.21 [-2.42, 4.85] | | | | | |
| L 186 | L 958 | L Lateral habenula | 8.92 ± 2.50 | 15 | 12.63 ± 4.44 | 15 | 11.94 ± 3.48 | 12 | 12.31 ± 2.77 | 12 | -0.13 [-2.67, 2.48] | 0.33 [-2.22, 2.84] | -0.54 [-4.26, 3.03] | | | | | |
| R 1097 | R 1129 | R Hypothalamus | 9.67 ± 2.14 | 14 | 10.14 ± 2.00 | 15 | 11.38 ± 2.35 | 12 | 13.66 ± 1.86 | 12 | -0.11 [-2.66, 2.42] | -0.07 [-2.44, 2.33] | -0.19 [-3.68, 3.24] | | | | | |
| L 1097 | L 1129 | L Hypothalamus | 8.57 ± 1.57 | 14 | 11.14 ± 2.28 | 14 | 11.64 ± 2.17 | 12 | 13.68 ± 1.95 | 12 | 0.08 [-2.5, 2.61] | 0.13 [-2.29, 2.57] | -0.39 [-3.9, 3.1] | | | | | |
| R 157 | R 1097 | R Periventricular zone | 16.58 ± 5.26 | 14 | 21.85 ± 6.78 | 15 | 23.54 ± 5.66 | 12 | 31.46 ± 5.42 | 12 | 0.14 [-2.4, 2.67] | 0.24 [-2.14, 2.64] | -0.25 [-3.68, 3.17] | | | | | |
| L 157 | L 1097 | L Periventricular zone | 14.32 ± 3.24 | 15 | 27.32 ± 7.97 | 15 | 26.68 ± 5.89 | 12 | 29.95 ± 5.43 | 12 | 0.53 [-1.98, 3.05] | 0.78 [-1.58, 3.17] | -1.03 [-4.5, 2.42] | | | | | |
| R 390 | R 157 | R Supraoptic nucleus | 83.65 ± 36.42 | 10 | 72.34 ± 21.83 | 7 | 79.37 ± 36.24 | 10 | 17.92 ± 9.22 | 8 | -0.15 [-2.89, 2.61] | -0.44 [-3.35, 2.42] | -0.74 [-4.74, 3.27] | | | | | |
| L 390 | L 157 | L Supraoptic nucleus | 39.56 ± 19.55 | 9 | 73.99 ± 35.32 | 6 | 69.59 ± 32.01 | 9 | 24.75 ± 11.90 | 7 | 0.68 [-2.16, 3.57] | 0.43 [-2.64, 3.48] | -2.1 [-6.34, 2.06] | | | | | |
| R 38 | R 157 | R Paraventricular hypothalamic nucleus | 61.56 ± 22.81 | 13 | 140.29 ± 32.59 | 9 | 73.92 ± 19.24 | 11 | 162.74 ± 30.03 | 10 | -0.04 [-2.65, 2.54] | 0.99 [-1.66, 3.63] | -0.06 [-3.73, 3.66] | | | | | |
| L 38 | L 157 | L Paraventricular hypothalamic nucleus | 64.58 ± 16.26 | 13 | 146.01 ± 32.11 | 9 | 114.74 ± 26.80 | 11 | 143.40 ± 20.36 | 10 | 0.54 [-2.03, 3.1] | 0.98 [-1.67, 3.59] | -1.05 [-4.73, 2.68] | | | | | |
| R 30 | R 157 | R Periventricular hypothalamic nucleus, an | 9.16 ± 4.68 | 8 | 93.78 ± 54.91 | 6 | 58.19 ± 28.05 | 7 | 67.79 ± 21.76 | 8 | 5.29 [2.23, 8.29]* | 8.72 [5.63, 11.8]* | -8.41 [-12.6, -4.15]* | -8.9 [-11.6, -6.09]* | -5.62 [-8.26, -2.96]* | -6.49 [-9.06, -3.93]* | -3.31 [-6.26, -0.53]* | 2.42 [-0.17, 5.38] | -0.88 [-3.53, 1.77] |
| L 30 | L 157 | L Periventricular hypothalamic nucleus, ant | 6.15 ± 6.12 | 8 | 123.40 ± 50.10 | 6 | 32.05 ± 11.41 | 7 | 74.64 ± 24.67 | 8 | 4.15 [1.12, 7.15]* | 18.56 [15.51, 21.62]* | -12.37 [-16.62, -8.12]* | -18.73 [-21.57, -16.07]* | -4.49 [-7.2, -1.92]* | -11.23 [-13.84, -8.69]* | -14.27 [-17.12, -11.4]* | 7.51 [4.71, 10.23]* | -6.75 [-9.42, -4.11]* |
| R 118 | R 157 | R Periventricular hypothalamic nucleus, int | 16.91 ± 8.68 | 13 | 24.28 ± 8.43 | 15 | 16.23 ± 6.61 | 12 | 37.83 ± 10.30 | 12 | -0.27 [-2.83, 2.32] | 0.41 [-2, 2.85] | 0.38 [-3.08, 3.85] | | | | | |
| L 118 | L 157 | L Periventricular hypothalamic nucleus, inte | 13.61 ± 6.53 | 13 | 30.71 ± 12.87 | 15 | 21.70 ± 8.08 | 12 | 40.78 ± 12.21 | 12 | 0.38 [-2.22, 2.9] | 1.25 [-1.18, 3.69] | -0.33 [-3.86, 3.08] | | | | | |
| R 223 | R 157 | R Arcuate hypothalamic nucleus | 30.70 ± 9.90 | 13 | 29.02 ± 11.45 | 14 | 31.36 ± 11.41 | 11 | 30.42 ± 8.68 | 11 | -0.2 [-2.78, 2.41] | -0.01 [-2.49, 2.48] | -0.49 [-4.05, 3.05] | | | | | |
| L 223 | L 157 | L Arcuate hypothalamic nucleus | 27.88 ± 10.08 | 13 | 48.79 ± 15.14 | 15 | 20.26 ± 3.18 | 11 | 28.59 ± 7.21 | 11 | -0.48 [-3.1, 2.09] | 0.74 [-1.73, 3.19] | -0.9 [-4.43, 2.61] | | | | | |
| R 141 | R 1097 | R Periventricular region | 13.47 ± 2.79 | 15 | 17.95 ± 3.13 | 16 | 23.72 ± 4.85 | 12 | 28.61 ± 4.24 | 12 | 0.44 [-2.05, 2.9] | 0.15 [-2.24, 2.56] | -0.27 [-3.67, 3.23] | | | | | |
| L 141 | L 1097 | L Periventricular region | 12.68 ± 2.64 | 14 | 15.90 ± 4.74 | 16 | 23.57 ± 7.24 | 12 | 26.53 ± 4.53 | 12 | 0.58 [-1.98, 3.09] | 0.11 [-2.29, 2.53] | -0.37 [-3.82, 3.08] | | | | | |
| R 830 | R 141 | R Dorsomedial nucleus of the hypothalamu | 38.31 ± 9.00 | 12 | 39.45 ± 11.08 | 14 | 50.79 ± 11.53 | 11 | 69.68 ± 14.14 | 11 | 0.07 [-2.58, 2.66] | 0.02 [-2.48, 2.5] | 0.01 [-3.59, 3.64] | | | | | |
| L 830 | L 141 | L Dorsomedial nucleus of the hypothalamu | 26.31 ± 9.22 | 12 | 34.14 ± 12.81 | 14 | 57.21 ± 17.63 | 11 | 58.78 ± 11.62 | 11 | 0.93 [-1.7, 3.55] | 0.29 [-2.19, 2.76] | -0.69 [-4.25, 2.83] | | | | | |
| R 126 | R 141 | R Periventricular hypothalamic nucleus, po | 8.07 ± 3.99 | 9 | 19.93 ± 7.10 | 10 | 32.22 ± 9.79 | 7 | 27.37 ± 11.16 | 8 | 2.65 [-0.31, 5.64] | 1.04 [-1.7, 3.77] | -1.95 [-5.95, 2.03] | | | | | |
| L 126 | L 141 | L Periventricular hypothalamic nucleus, pos | 9.65 ± 3.33 | 9 | 7.91 ± 2.83 | 10 | 27.75 ± 10.73 | 7 | 30.91 ± 9.94 | 8 | 1.52 [-1.45, 4.5] | -0.62 [-3.4, 2.15] | 0.64 [-3.42, 4.62] | | | | | |
| R 347 | R 141 | R Subparaventricular zone | 18.00 ± 7.27 | 13 | 59.50 ± 18.50 | 9 | 51.13 ± 15.84 | 11 | 85.62 ± 22.13 | 10 | 1.61 [-0.97, 4.21] | 2.02 [-0.63, 4.71] | -0.62 [-4.28, 3.05] | | | | | |
| L 347 | L 141 | L Subparaventricular zone | 28.63 ± 12.54 | 13 | 61.05 ± 18.12 | 9 | 61.93 ± 25.98 | 10 | 67.29 ± 24.74 | 9 | 0.83 [-1.82, 3.48] | 0.86 [-1.86, 3.52] | -1.07 [-4.89, 2.69] | | | | | |
| R 286 | R 141 | R Suprachiasmatic nucleus | 37.05 ± 12.49 | 10 | 39.65 ± 12.95 | 6 | 132.14 ± 79.88 | 8 | 51.64 ± 37.11 | 7 | 2.52 [-0.34, 5.34] | -0.37 [-3.33, 2.57] | -2.58 [-6.81, 1.59] | | | | | |
| L 286 | L 141 | L Suprachiasmatic nucleus | 27.10 ± 11.76 | 9 | 32.98 ± 16.34 | 6 | 127.07 ± 66.33 | 8 | 56.97 ± 40.23 | 7 | 3.62 [0.71, 6.54]* | -0.23 [-3.26, 2.81] | -3.13 [-7.35, 1.11] | 0.03 [-2.66, 2.7] | -3.93 [-6.44, -1.45]* | -1.15 [-3.72, 1.48] | 3.98 [1.28, 6.78]* | -1.19 [-4.09, 1.53] | 2.78 [0.21, 5.6]* |
| R 467 | R 1097 | R Hypothalamic medial zone | 16.01 ± 3.87 | 15 | 15.95 ± 3.57 | 16 | 16.12 ± 3.26 | 12 | 17.82 ± 2.60 | 12 | -0.33 [-2.83, 2.17] | -0.19 [-2.53, 2.18] | -0.19 [-3.58, 3.25] | | | | | |
| L 467 | L 1097 | L Hypothalamic medial zone | 13.95 ± 3.08 | 15 | 18.37 ± 4.04 | 16 | 15.77 ± 3.19 | 12 | 19.27 ± 2.85 | 12 | -0.2 [-2.74, 2.3] | 0.14 [-2.21, 2.54] | -0.37 [-3.77, 3.1] | | | | | |
| R 88 | R 467 | R Anterior hypothalamic nucleus | 26.22 ± 5.25 | 15 | 42.47 ± 11.33 | 12 | 41.77 ± 14.50 | 12 | 41.18 ± 6.93 | 10 | 0.27 [-2.24, 2.78] | 0.42 [-2.02, 2.91] | -0.91 [-4.48, 2.62] | | | | | |
| L 88 | L 467 | L Anterior hypothalamic nucleus | 29.34 ± 6.99 | 14 | 47.55 ± 12.63 | 12 | 44.18 ± 11.01 | 12 | 37.89 ± 8.89 | 11 | 0.23 [-2.31, 2.73] | 0.48 [-2.02, 2.96] | -1.09 [-4.59, 2.45] | | | | | |
| R 331 | R 467 | R Mammillary body | 11.81 ± 4.85 | 9 | 7.57 ± 2.55 | 13 | 7.98 ± 2.58 | 11 | 9.47 ± 1.37 | 7 | -0.5 [-3.34, 2.27] | -0.64 [-3.36, 2.05] | 0.31 [-3.56, 4.25] | | | | | |
| L 331 | L 467 | L Mammillary body | 7.43 ± 1.73 | 10 | 7.76 ± 2.41 | 12 | 5.51 ± 1.33 | 11 | 11.43 ± 1.79 | 8 | -0.33 [-3.05, 2.36] | -0.22 [-2.86, 2.42] | 0.51 [-3.27, 4.29] | | | | | |
| R 210 | R 331 | R Lateral mammillary nucleus | 1.98 ± 1.98 | 6 | 23.78 ± 21.68 | 8 | 11.70 ± 6.28 | 9 | 10.12 ± 7.19 | 7 | 4.56 [1.53, 7.64]* | 10.48 [7.37, 13.57]* | -11.59 [-15.81, -7.37]* | -10.66 [-13.47, -7.95]* | -4.88 [-7.58, -2.2]* | -4.33 [-7.25, -1.56]* | -5.77 [-8.28, -3.21]* | 6.32 [3.63, 8.98]* | 0.55 [-2.06, 3.16] |
| L 210 | L 331 | L Lateral mammillary nucleus | 10.63 ± 10.63 | 7 | 11.75 ± 6.45 | 9 | 10.91 ± 9.01 | 8 | 19.10 ± 5.25 | 8 | -0.3 [-3.31, 2.69] | -0.34 [-3.27, 2.57] | 0.7 [-3.34, 4.81] | | | | | |
| R 491 | R 331 | R Medial mammillary nucleus | 29.37 ± 22.66 | 7 | 12.32 ± 7.14 | 10 | 4.77 ± 1.92 | 8 | 10.85 ± 2.79 | 7 | -1.16 [-4.23, 1.84] | -1.06 [-3.99, 1.8] | 0.88 [-3.27, 5.06] | | | | | |
| L 491 | L 331 | L Medial mammillary nucleus | 6.62 ± 2.72 | 7 | 10.61 ± 4.53 | 9 | 4.45 ± 1.32 | 8 | 12.54 ± 2.57 | 7 | -0.65 [-3.7, 2.35] | 0.15 [-2.78, 3.03] | 0.7 [-3.46, 4.81] | | | | | |
| R 732 | R 491 | R Medial mammillary nucleus, median part | 91.09 ± 69.77 | 7 | 11.61 ± 6.72 | 9 | 5.98 ± 4.30 | 7 | 15.38 ± 4.99 | 7 | -1.2 [-4.31, 1.88] | -1.33 [-4.33, 1.6] | 1.02 [-3.21, 5.2] | | | | | |
| L 732 | L 491 | L Medial mammillary nucleus, median part | 24.04 ± 9.30 | 7 | 11.33 ± 5.68 | 9 | 11.22 ± 7.87 | 7 | 17.87 ± 9.93 | 7 | -0.82 [-3.9, 2.24] | -0.99 [-3.9, 1.94] | 0.83 [-3.37, 5.06] | | | | | |
| R 60682 | R 491 | R Medial mammillary nucleus, lateral part | 52.94 ± 45.72 | 7 | 28.16 ± 15.74 | 10 | 11.67 ± 3.63 | 8 | 13.29 ± 5.65 | 7 | -1.1 [-4.12, 1.91] | -0.94 [-3.8, 1.92] | 0.59 [-3.51, 4.67] | | | | | |
| L 606826 | L 491 | L Medial mammillary nucleus, lateral part | 9.39 ± 5.74 | 7 | 32.56 ± 13.80 | 9 | 5.34 ± 2.24 | 8 | 21.18 ± 7.23 | 7 | -0.76 [-3.8, 2.25] | 2.02 [-0.92, 4.95] | -0.69 [-4.91, 3.52] | | | | | |
| R 60682 | R 491 | R Medial mammillary nucleus, medial part | 10.02 ± 6.87 | 7 | 6.95 ± 6.12 | 10 | 4.93 ± 3.23 | 8 | 35.54 ± 11.13 | 6 | -0.82 [-3.8, 2.19] | -0.78 [-3.64, 2.11] | 3.38 [-0.83, 7.62] | | | | | |
| L 606826 | L 491 | L Medial mammillary nucleus, medial part | 10.68 ± 4.36 | 7 | 11.94 ± 9.02 | 9 | 2.83 ± 2.11 | 8 | 50.26 ± 6.94 | 6 | -1.05 [-4.07, 1.99] | -0.33 [-3.26, 2.6] | 4.3 [0.12, 8.49]* | 0.13 [-2.47, 2.69] | 0.74 [-1.85, 3.42] | -3.81 [-6.63, -0.93]* | -0.59 [-3.19, 1.95] | -3.94 [-6.61, -1.2]* | -4.53 [-7.37, -1.8]* |
| R 525 | R 331 | R Supramammillary nucleus | 29.33 ± 10.20 | 8 | 23.01 ± 7.44 | 10 | 34.60 ± 12.25 | 10 | 41.62 ± 6.09 | 7 | 0 [-2.87, 2.85] | -0.59 [-3.49, 2.2] | 0.41 [-3.58, 4.47] | | | | | |
| L 525 | L 331 | L Supramammillary nucleus | 31.64 ± 9.01 | 9 | 25.51 ± 8.41 | 10 | 30.39 ± 9.72 | 10 | 56.99 ± 17.11 | 8 | -0.23 [-3.08, 2.57] | -0.6 [-3.37, 2.16] | 0.96 [-2.94, 4.87] | | | | | |
| R 557 | R 331 | R Tuberomammillary nucleus | 8.77 ± 3.86 | 8 | 13.58 ± 4.28 | 12 | 15.27 ± 5.85 | 9 | 12.36 ± 4.08 | 6 | 0.54 [-2.28, 3.42] | 0.17 [-2.56, 2.91] | -1.03 [-5.16, 2.91] | | | | | |
| L 557 | L 331 | L Tuberomammillary nucleus | 6.55 ± 2.91 | 10 | 8.52 ± 3.29 | 10 | 7.24 ± 2.01 | 9 | 3.66 ± 3.05 | 7 | -0.01 [-2.84, 2.76] | -0.21 [-2.91, 2.51] | -0.86 [-4.78, 3.09] | | | | | |
| L 1126 | L 557 | L Tuberomammillary nucleus, dorsal part | 19.79 ± 14.23 | 8 | 37.19 ± 16.34 | 8 | 31.94 ± 11.87 | 6 | 10.26 ± 10.26 | 6 | 0.07 [-3.08, 3.22] | 0.14 [-2.76, 3.09] | -1.6 [-5.92, 2.77] | | | | | |
| R 1 | R 557 | R Tuberomammillary nucleus, ventral part | 17.91 ± 8.58 | 7 | 23.32 ± 9.49 | 10 | 16.18 ± 7.19 | 9 | 30.73 ± 12.62 | 6 | -0.22 [-3.2, 2.78] | 0 [-2.97, 2.93] | 0.28 [-3.87, 4.47] | | | | | |
| L 1 | L 557 | L Tuberomammillary nucleus, ventral part | 11.93 ± 4.90 | 10 | 8.34 ± 5.76 | 8 | 8.79 ± 2.96 | 9 | 1.77 ± 1.77 | 7 | -0.37 [-3.17, 2.37] | -0.4 [-3.2, 2.38] | -0.72 [-4.76, 3.27] | | | | | |
| R 980 | R 467 | R Dorsal premammillary nucleus | 64.21 ± 26.94 | 7 | 33.02 ± 13.58 | 8 | 24.54 ± 7.59 | 7 | 34.63 ± 10.17 | 6 | -0.85 [-3.97, 2.22] | -1.12 [-4.11, 1.91] | 0.89 [-3.43, 5.15] | | | | | |
| L 980 | L 467 | L Dorsal premammillary nucleus | 43.19 ± 13.06 | 10 | 34.97 ± 16.02 | 9 | 26.05 ± 12.81 | 7 | 30.63 ± 5.05 | 8 | -0.44 [-3.4, 2.52] | -0.64 [-3.4, 2.12] | 0.25 [-3.82, 4.26] | | | | | |
| R 1004 | R 467 | R Ventral premammillary nucleus | 28.38 ± 12.38 | 8 | 32.63 ± 10.52 | 11 | 22.40 ± 8.21 | 9 | 39.38 ± 12.17 | 7 | -0.41 [-3.34, 2.48] | -0.37 [-3.14, 2.4] | 0.41 [-3.54, 4.41] | | | | | |
| L 1004 | L 467 | L Ventral premammillary nucleus | 26.98 ± 11.20 | 10 | 53.74 ± 19.02 | 10 | 32.81 ± 8.63 | 10 | 34.54 ± 9.46 | 8 | 0.2 [-2.55, 2.89] | 0.5 [-2.16, 3.22] | -0.98 [-4.81, 2.87] | | | | | |
| R 63 | R 467 | R Paraventricular hypothalamic nucleus, de | 68.25 ± 16.55 | 13 | 40.81 ± 7.64 | 12 | 78.80 ± 19.75 | 9 | 63.70 ± 8.70 | 11 | -0.19 [-2.9, 2.45] | -0.48 [-2.98, 2.04] | -0.16 [-3.78, 3.54] | | | | | |
| L 63 | L 467 | L Paraventricular hypothalamic nucleus, de | 58.38 ± 9.90 | 12 | 63.71 ± 13.47 | 12 | 110.43 ± 30.13 | 9 | 51.19 ± 11.03 | 10 | 0.62 [-2.12, 3.32] | 0.1 [-2.45, 2.63] | -1.44 [-5.13, 2.29] | | | | | |
| R 693 | R 467 | R Ventromedial hypothalamic nucleus | 17.70 ± 3.93 | 14 | 23.87 ± 6.12 | 13 | 17.78 ± 7.02 | 12 | 45.46 ± 8.38 | 11 | -0.27 [-2.81, 2.28] | 0.6 [-1.87, 3.07] | 0.51 [-3.02, 4.03] | | | | | |
| L 693 | L 467 | L Ventromedial hypothalamic nucleus | 20.90 ± 4.19 | 14 | 28.53 ± 9.54 | 13 | 16.73 ± 4.27 | 11 | 48.49 ± 14.74 | 11 | -0.47 [-3.05, 2.12] | 0.61 [-1.87, 3.11] | 0.46 [-3.09, 4.02] | | | | | |
| R 946 | R 467 | R Posterior hypothalamic nucleus | 57.59 ± 15.87 | 14 | 47.73 ± 12.78 | 15 | 66.80 ± 15.09 | 11 | 45.16 ± 8.84 | 9 | 0.03 [-2.58, 2.59] | -0.25 [-2.66, 2.19] | -0.77 [-4.32, 2.8] | | | | | |
| L 946 | L 467 | L Posterior hypothalamic nucleus | 48.44 ± 13.08 | 14 | 49.74 ± 11.82 | 15 | 62.74 ± 17.50 | 11 | 43.28 ± 7.54 | 9 | 0.18 [-2.42, 2.73] | -0.07 [-2.44, 2.37] | -1 [-4.55, 2.54] | | | | | |
| R 290 | R 1097 | R Hypothalamic lateral zone | 8.60 ± 1.98 | 15 | 9.30 ± 2.01 | 16 | 9.94 ± 2.26 | 12 | 13.04 ± 1.91 | 12 | -0.17 [-2.66, 2.31] | -0.1 [-2.48, 2.27] | -0.04 [-3.47, 3.38] | | | | | |
| L 290 | L 1097 | L Hypothalamic lateral zone | 6.93 ± 1.31 | 15 | 9.35 ± 1.86 | 16 | 10.27 ± 1.86 | 12 | 12.79 ± 1.97 | 12 | 0.16 [-2.36, 2.64] | 0.16 [-2.18, 2.5] | -0.29 [-3.71, 3.12] | | | | | |
| R 194 | R 290 | R Lateral hypothalamic area | 19.30 ± 4.49 | 15 | 24.86 ± 6.70 | 16 | 23.45 ± 5.60 | 10 | 23.41 ± 3.79 | 12 | -0.24 [-2.81, 2.3] | 0.11 [-2.24, 2.48] | -0.47 [-3.9, 3.06] | | | | | |
| L 194 | L 290 | L Lateral hypothalamic area | 16.10 ± 3.25 | 15 | 19.93 ± 5.00 | 16 | 25.93 ± 5.40 | 11 | 25.47 ± 5.20 | 12 | 0.25 [-2.27, 2.79] | 0.06 [-2.34, 2.44] | -0.56 [-4.07, 2.91] | | | | | |
| R 364 | R 290 | R Parasubthalamic nucleus | 31.29 ± 8.07 | 12 | 51.81 ± 12.95 | 13 | 40.23 ± 12.24 | 11 | 85.00 ± 15.14 | 8 | 0.12 [-2.5, 2.71] | 0.41 [-2.12, 2.9] | 0.54 [-3.13, 4.28] | | | | | |
| L 364 | L 290 | L Parasubthalamic nucleus | 26.34 ± 10.25 | 12 | 55.39 ± 19.23 | 13 | 26.40 ± 6.64 | 11 | 63.91 ± 12.08 | 9 | -0.11 [-2.73, 2.45] | 0.9 [-1.62, 3.41] | -0.13 [-3.77, 3.57] | | | | | |
| R 57607 | R 290 | R Perifornical nucleus | 24.78 ± 5.39 | 15 | 23.21 ± 6.17 | 14 | 35.65 ± 12.36 | 12 | 31.24 ± 7.87 | 11 | 0.12 [-2.41, 2.61] | -0.12 [-2.56, 2.29] | -0.62 [-4.1, 2.92] | | | | | |
| L 576073 | L 290 | L Perifornical nucleus | 12.82 ± 3.93 | 15 | 32.87 ± 12.64 | 14 | 32.68 ± 7.02 | 12 | 47.44 ± 9.73 | 10 | 1.23 [-1.3, 3.73] | 1.5 [-0.94, 3.96] | -0.82 [-4.41, 2.75] | | | | | |
| L 173 | L 290 | L Retrochiasmatic area | 5.61 ± 3.08 | 12 | 16.88 ± 12.58 | 7 | 9.31 ± 3.49 | 8 | 12.81 ± 8.15 | 7 | 0.28 [-2.51, 3.04] | 1.34 [-1.47, 4.16] | -1.41 [-5.47, 2.64] | | | | | |
| R 470 | R 290 | R Subthalamic nucleus | 5.86 ± 2.30 | 12 | 6.59 ± 3.03 | 13 | 11.35 ± 6.63 | 11 | 43.86 ± 23.55 | 9 | 0.68 [-1.99, 3.31] | 0.05 [-2.5, 2.6] | 4.97 [1.29, 8.61]* | -0.23 [-2.36, 1.84] | -0.99 [-3.19, 1.21] | -6.58 [-8.91, -4.3]* | 0.76 [-1.41, 2.85] | -6.35 [-8.51, -3.98]* | -5.57 [-7.94, -3.2]* |
| L 470 | L 290 | L Subthalamic nucleus | 9.24 ± 3.82 | 10 | 6.24 ± 2.37 | 16 | 7.13 ± 2.98 | 11 | 24.28 ± 11.33 | 11 | -0.48 [-3.17, 2.2] | -0.44 [-2.96, 2.1] | 1.83 [-1.76, 5.44] | | | | | |
| R 614 | R 290 | R Tuberal nucleus | 17.77 ± 5.06 | 14 | 16.74 ± 3.76 | 14 | 18.02 ± 4.43 | 11 | 36.96 ± 7.86 | 11 | -0.26 [-2.8, 2.3] | -0.06 [-2.49, 2.36] | 0.66 [-2.86, 4.21] | | | | | |
| L 614 | L 290 | L Tuberal nucleus | 9.68 ± 2.13 | 14 | 29.43 ± 8.89 | 14 | 24.78 ± 5.34 | 11 | 29.23 ± 6.80 | 11 | 1.29 [-1.26, 3.83] | 2.04 [-0.37, 4.47] | -2.03 [-5.56, 1.5] | | | | | |
| R 797 | R 290 | R Zona incerta | 14.20 ± 3.56 | 15 | 14.92 ± 4.15 | 16 | 17.70 ± 3.68 | 12 | 21.95 ± 3.63 | 12 | -0.08 [-2.59, 2.42] | -0.13 [-2.49, 2.24] | -0.06 [-3.52, 3.36] | | | | | |
| L 797 | L 290 | L Zona incerta | 12.57 ± 2.36 | 15 | 10.50 ± 3.03 | 16 | 16.06 ± 3.43 | 12 | 19.38 ± 2.99 | 12 | -0.05 [-2.59, 2.46] | -0.35 [-2.71, 2.02] | 0.13 [-3.29, 3.54] | | | | | |
| R 804 | R 797 | R Fields of Forel | 19.98 ± 9.38 | 10 | 18.41 ± 8.89 | 11 | 18.24 ± 8.34 | 9 | 24.85 ± 7.61 | 8 | -0.12 [-2.92, 2.62] | -0.25 [-2.92, 2.39] | 0.06 [-3.72, 3.9] | | | | | |
| L 804 | L 797 | L Fields of Forel | 9.00 ± 3.30 | 10 | 21.36 ± 9.59 | 10 | 19.94 ± 6.87 | 10 | 18.68 ± 7.87 | 8 | 1.2 [-1.57, 3.97] | 1.08 [-1.63, 3.79] | -1.76 [-5.67, 2.14] | | | | | |
| R 10671 | R 1097 | R Median eminence | 27.35 ± 15.86 | 12 | 11.59 ± 5.45 | 14 | 10.28 ± 6.11 | 10 | 18.75 ± 8.98 | 10 | -0.69 [-3.36, 1.98] | -0.55 [-3.05, 1.96] | 0.09 [-3.51, 3.76] | | | | | |
| L 10671 | L 1097 | L Median eminence | 13.07 ± 13.07 | 12 | 8.60 ± 4.74 | 14 | 7.11 ± 5.83 | 11 | 18.56 ± 11.18 | 10 | -0.68 [-3.29, 1.9] | -0.32 [-2.84, 2.16] | 0.58 [-3.04, 4.21] | | | | | |
| R 313 | R 343 | R Midbrain | 13.92 ± 3.61 | 13 | 10.76 ± 4.06 | 13 | 10.85 ± 3.13 | 12 | 11.33 ± 2.57 | 10 | -3.02 [-133.71, 127.62] | 3.11 [-131.31, 122.12 | -1.17 [-193.86, 189.46] | | | | | |
| L 313 | L 343 | L Midbrain | 10.46 ± 2.52 | 13 | 11.28 ± 3.78 | 15 | 12.52 ± 3.99 | 12 | 11.91 ± 3.27 | 11 | | | | | | | | |
| R 339 | R 313 | R Midbrain, sensory related | 1.56 ± 1.05 | 11 | 16.73 ± 8.28 | 11 | 18.99 ± 15.77 | 10 | 28.05 ± 6.58 | 7 | 9.6 [-134.33, 153.42] | 3.39 [-133.47, 149.22 | -7.81 [-216.13, 203.07] | | | | | |
| L 339 | L 313 | L Midbrain, sensory related | 13.92 ± 8.73 | 11 | 10.29 ± 5.43 | 11 | 4.57 ± 2.91 | 10 | 19.71 ± 4.02 | 7 | -3.3 [-144.11, 142.05] | 2.39 [-141.85, 136.59 | -1.36 [-217.22, 211.33] | | | | | |
| L 302 | L 339 | L Superior colliculus, sensory related | 3.62 ± 2.45 | 6 | 25.89 ± 12.65 | 7 | 9.45 ± 6.26 | 7 | 34.78 ± 4.42 | 6 | 1.9 [-180.58, 182.77] | 3.19 [-177.05, 189.46 | -6.14 [-265.95, 249.74] | | | | | |
| R 59962 | R 339 | R Subcommissural organ | 0.00 ± 0.00 | 11 | 19.75 ± 19.75 | 9 | 32.00 ± 32.00 | 10 | 112.82 ± 61.17 | 7 | 18.61 [174.82, 464.98] | 95.42 [46.97, 342.41 | 607.22 [389.49, 829.49]* | 197.24 [-345.02, -49.76 | -320.09 [-464.47, -174.62]* | 128.13 [-1293.09, -974.8 | 123.11 [-30.06, 272.43] | 31.26 [-1096.35, -764.77 | -807.95 [-973.31, -651.92]* |
| L 599626 | L 339 | L Subcommissural organ | 25.25 ± 17.73 | 11 | 0.00 ± 0.00 | 9 | 6.40 ± 6.40 | 10 | 8.47 ± 8.47 | 7 | -2.31 [-145.36, 139.38] | 2.42 [-149.44, 145.37 | -3.86 [-222.75, 212.66] | | | | | |
| R 323 | R 313 | R Midbrain, motor related | 12.80 ± 3.77 | 13 | 9.93 ± 3.72 | 14 | 11.31 ± 3.46 | 12 | 11.02 ± 2.26 | 10 | -3.24 [-133.21, 126.52] | 2.91 [-127.47, 122.15 | -2.34 [-189.68, 186.19] | | | | | |
| L 323 | L 313 | L Midbrain, motor related | 8.10 ± 1.78 | 13 | 8.19 ± 3.03 | 15 | 17.16 ± 7.01 | 12 | 12.37 ± 4.02 | 10 | -0.65 [-131.04, 130.83] | 1.67 [-128.17, 120.96 | -3.86 [-189.88, 181.71] | | | | | |
| L 381 | L 323 | L Substantia nigra, reticular part | 0.00 ± 0.00 | 6 | 2.46 ± 2.08 | 8 | 1.32 ± 1.32 | 8 | 3.87 ± 1.78 | 7 | 11.4 [-169.66, 189.43] | 2.58 [-153.55, 201.62 | -2.79 [-253.49, 244.06] | | | | | |
| R 128 | R 323 | R Midbrain reticular nucleus | 11.21 ± 3.94 | 8 | 9.30 ± 3.98 | 10 | 12.88 ± 5.13 | 10 | 19.43 ± 3.14 | 6 | -0.83 [-159.87, 159.47] | -1.9 [-158.52, 155.5] | -3.54 [-229.93, 227.06] | | | | | |
| L 128 | L 323 | L Midbrain reticular nucleus | 8.21 ± 2.11 | 8 | 6.40 ± 2.12 | 10 | 9.82 ± 4.01 | 10 | 18.74 ± 4.60 | 7 | -1.82 [-159.5, 151.42] | 1.51 [-156.21, 155.85 | -1.96 [-223.72, 221.27] | | | | | |
| R 795 | R 323 | R Periaqueductal gray | 37.03 ± 14.58 | 13 | 24.15 ± 7.42 | 14 | 24.91 ± 6.73 | 12 | 20.97 ± 4.74 | 10 | -3.38 [-132.66, 125.7] | 2.73 [-128.72, 123.12 | -0.87 [-188.96, 189.11] | | | | | |
| L 795 | L 323 | L Periaqueductal gray | 21.49 ± 4.62 | 13 | 20.37 ± 7.09 | 14 | 35.34 ± 11.26 | 12 | 21.77 ± 6.05 | 10 | -2.17 [-134.17, 132.26] | -1.58 [-131.74, 126] | -3.46 [-198.28, 187.84] | | | | | |
| R 50 | R 795 | R Precommissural nucleus | 82.12 ± 31.21 | 10 | 54.36 ± 15.87 | 13 | 66.33 ± 14.97 | 12 | 42.45 ± 11.48 | 10 | -2.19 [-139.61, 142.02] | -1.9 [-141.98, 136.23 | -3.72 [-199.54, 194.99] | | | | | |
| L 50 | L 795 | L Precommissural nucleus | 50.80 ± 12.39 | 10 | 42.81 ± 15.30 | 13 | 64.84 ± 20.75 | 12 | 99.93 ± 36.10 | 10 | -1.01 [-140.23, 141.13] | 0.79 [-136.39, 134.57 | -4.37 [-205.26, 193.6] | | | | | |
| R 1100 | R 323 | R Pretectal region | 9.03 ± 2.68 | 11 | 15.24 ± 7.46 | 12 | 10.46 ± 3.16 | 11 | 10.98 ± 2.12 | 8 | -1.46 [-145.99, 139.12] | -0.45 [-136.85, 138.3 | -3.84 [-212.29, 202.82] | | | | | |
| L 1100 | L 323 | L Pretectal region | 8.00 ± 1.87 | 12 | 15.56 ± 6.90 | 11 | 10.30 ± 4.35 | 10 | 13.15 ± 2.48 | 7 | -1.37 [-144.02, 139.39] | 0.32 [-136.35, 138.69 | -4.38 [-218.24, 206.69] | | | | | |
| R 215 | R 1100 | R Anterior pretectal nucleus | 12.71 ± 3.87 | 11 | 25.42 ± 14.17 | 12 | 20.25 ± 6.15 | 11 | 21.81 ± 5.23 | 8 | -1.31 [-138.32, 136.47] | -0.8 [-135.94, 136.54 | -3.72 [-212.91, 202.12] | | | | | |
| L 215 | L 1100 | L Anterior pretectal nucleus | 11.74 ± 3.08 | 12 | 24.09 ± 13.07 | 11 | 19.79 ± 6.68 | 10 | 26.81 ± 6.84 | 7 | -0.53 [-143.89, 140.24] | -1.4 [-139.46, 137.53 | -3.83 [-213.6, 206.28] | | | | | |
| R 628 | R 1100 | R Nucleus of the optic tract | 15.98 ± 6.28 | 8 | 11.08 ± 5.65 | 8 | 17.33 ± 6.37 | 8 | 21.68 ± 7.47 | 7 | -1.43 [-166.75, 162.93] | 1.83 [-165.76, 162.76 | -2.87 [-243.92, 229.87] | | | | | |
| L 628 | L 1100 | L Nucleus of the optic tract | 16.47 ± 4.81 | 8 | 23.91 ± 8.96 | 8 | 10.44 ± 6.98 | 9 | 24.39 ± 3.43 | 7 | -2.52 [-160.21, 153.26] | 2.11 [-166.22, 160.13 | -1.28 [-229.31, 232.48] | | | | | |
| R 634 | R 1100 | R Nucleus of the posterior commissure | 38.89 ± 15.69 | 9 | 57.23 ± 35.39 | 8 | 16.90 ± 8.36 | 8 | 22.61 ± 8.16 | 7 | -4.43 [-165.82, 150.9] | 2.66 [-162.31, 155.79 | -0.75 [-230.15, 229.57] | | | | | |
| L 634 | L 1100 | L Nucleus of the posterior commissure | 24.99 ± 10.14 | 9 | 52.78 ± 33.71 | 8 | 26.19 ± 12.23 | 8 | 25.68 ± 7.98 | 7 | -2.58 [-163.75, 157.49] | -0.3 [-162.88, 158.72 | -4.69 [-238.86, 231.37] | | | | | |
| R 706 | R 1100 | R Olivary pretectal nucleus | 16.68 ± 7.28 | 8 | 10.62 ± 4.76 | 8 | 13.39 ± 5.28 | 7 | 22.15 ± 8.52 | 7 | -0.19 [-165.9, 169.17] | 1.26 [-165.03, 162.98 | -4.52 [-245.06, 233.62] | | | | | |
| L 706 | L 1100 | L Olivary pretectal nucleus | 18.43 ± 9.41 | 6 | 8.94 ± 4.73 | 8 | 3.63 ± 2.33 | 6 | 25.77 ± 7.78 | 7 | -3.41 [-191.55, 188.85] | 2.73 [-179.72, 171.22 | -0.77 [-258.33, 246.73] | | | | | |
| R 1061 | R 1100 | R Posterior pretectal nucleus | 33.81 ± 19.98 | 6 | 48.73 ± 28.74 | 7 | 42.69 ± 20.45 | 7 | 18.11 ± 8.25 | 6 | -1.85 [-183.98, 180.57] | -2.25 [-185.4, 183.14 | -3.53 [-255.14, 254.51] | | | | | |
| L 1061 | L 1100 | L Posterior pretectal nucleus | 36.68 ± 15.04 | 8 | 30.34 ± 15.06 | 8 | 23.00 ± 17.77 | 9 | 22.33 ± 8.36 | 7 | -2.79 [-163.26, 160.02] | 3.18 [-169.48, 162.15 | -2.56 [-235.03, 229.04] | | | | | |
| R 54900 | R 1100 | R Retroparafascicular nucleus | 10.16 ± 10.16 | 6 | 1.40 ± 1.40 | 8 | 14.28 ± 8.01 | 6 | 14.69 ± 8.61 | 6 | -2.13 [-187.46, 191.56] | -2.98 [-178.5, 174.73 | -2.29 [-255.61, 254.15] | | | | | |

**Table S2**. **Results from parent and subregion level tdT+ mapping.** Mean tdT+ cells/mm$^2$ +/- standard error of the mean and sample size for each experimental group (n) for each sampled brain region. Parent ID indicates the hierarchical "parent" area for each region. Results from Bayesian statistical models (genotype, treatment, and brain region as fixed effects, animal as random effect) show the estimated relative change (normalized to regional WT-Sal) and 95% credible intervals (CI) in tdT+ cells due to interactions between brain region and 5XFAD genotype (genotype effect), PTZ treatment (treatment effect), and 5XFAD genotype and PTZ treatment (Interaction). Estimated marginal means and 95% CIs were used for group contrasts at each brain region, where credible genotype, treatment, or interaction effects were found. Separate models were performed for the isocortex, olfactory areas, hippocampal formation, cortical subplate, striatum, pallidum, thalamus, hypothalamus, and midbrain. *=Results are considered credible when the 95% CI does not cross 0.

| Brain region | Dependent variable | Independent variable | | | | | |
|---|---|---|---|---|---|---|---|
| | | Sex | | Genotype | | Kindling | |
| | | t statistic (DF) (M) | p value | t statistic (WT) | p value | t statistic (Sal) | p value |
| R isocortex | **AT8+/mm^2** | -0.743(46) | 0.46 | -4.58(46) | <0.0001 | -1.838(46) | 0.073 |
| R olfactory areas | | -0.3(48) | 0.77 | -2.55(48) | 0.014 | -0.143(48) | 0.89 |
| R hippocampus | | -2.52(48) | 0.015 | -5.014(48) | <0.0001 | -0.355(48) | 0.72 |
| R cortical subplate | | -1.162(48) | 0.25 | -5.533(48) | <0.0001 | -0.437(48) | 0.66 |
| R striatum | | -1.582(48) | 0.12 | -8.122(48) | <0.0001 | -0.49(48) | 0.63 |
| R thalamus | | -1.43(48) | 0.16 | -1.55(48) | 0.13 | -0.766(48) | 0.48 |
| R hypothalamus | | -0.456(46) | 0.65 | -1.187(46) | 0.24 | -0.381(46) | 0.71 |
| R midbrain | | -1.948(41) | 0.0583 | -0.395(41) | 0.7 | 0.144(41) | 0.88 |
| R fiber tracts | | -2.249(48) | 0.016 | -6.894(48) | <0.0001 | -0.783(48) | 0.44 |
| L isocortex | | -1.558(46) | 0.13 | -5.103(46) | <0.0001 | -4.089(46) | 0.073 |
| L olfactory areas | | -0.194(46) | 0.85 | -3.213(46) | 0.0024 | -0.757(46) | 0.453 |
| L hippocampus | | -3.002 (48) | 0.0043 | -5.559(48) | <0.0001 | -0.468(48) | 0.64 |
| L cortical subplate | | -0.28(48) | 0.79 | -4.773(48) | <0.0001 | -1.026(48) | 0.3 |
| L striatum | | 0.353(48) | 0.73 | -5.746(48) | <0.0001 | -1.756(48) | 0.085 |
| L Thalamus | | -0.463(47) | 0.65 | -4.681(47) | <0.0001 | -1.825(47) | 0.074 |
| L hypothalamus | | -0.0781(45) | 0.94 | -0.52(45) | 0.6 | 0.271(45) | 0.79 |
| L midbrain | | -1.464(44) | 0.15 | -1.171(44) | 0.25 | -0.781(44) | 0.44 |
| L fiber tracts | | -1.285(48) | 0.2 | -4.275(48) | <0.0001 | -2.012(48) | 0.05 |
| R isocortex | **tdT+/mm^2** | 1.844(50) | 0.071 | -2.573(50) | 0.013 | -4.872(50) | <0.0001 |
| R olfactory areas | | 2.164(49) | 0.035 | -2.585(49) | 0.013 | -4.296(49) | <0.0001 |
| R hippocampus | | 1.091(48) | 0.2803 | -1.809(48) | 0.076 | -4.476(48) | <0.0001 |
| R cortical subplate | | 1.824(51) | 0.074 | -2.883(51) | 0.0057 | -4.29(51) | <0.0001 |
| R striatum | | 1.937(51) | 0.058 | -1.877(51) | 0.066 | -4.644(51) | <0.0001 |
| R thalamus | | 0.118(51) | 0.91 | -3.235(51) | 0.0021 | -1.676(51) | 0.01 |
| R hypothalamus | | 1.306(49) | 0.17 | -1.576(49) | 0.12 | -0.712(49) | 0.48 |
| R midbrain | | 0.156(44) | 0.88 | 0.336(44) | 0.74 | 0.391(44) | 0.7 |
| L isocortex | | 1.61(50) | 0.11 | -2.66(50) | 0.011 | -4.848(50) | <0.0001 |
| L olfactory areas | | 1.762(49) | 0.084 | -2.802(49) | 0.0073 | -4.103(49) | 0.00015 |
| L hippocampus | | 0.887(51) | 0.38 | -1.504(51) | 0.14 | -4.844(51) | <0.0001 |
| L cortical subplate | | 1.555(51) | 0.13 | -3.057(51) | 0.0035 | -4.519(51) | <0.0001 |
| L striatum | | 1.142(50) | 0.16 | -2.455(50) | 0.018 | -5.148(50) | <0.0001 |
| L Thalamus | | -0.0743(50) | 0.94 | -3.939(50) | 0.00025 | -1.519(50) | 0.14 |
| L hypothalamus | | 1.753(48) | 0.086 | -1.805(48) | 0.077 | -1.326(48) | 0.19 |
| L midbrain | | 0.322(47) | 0.75 | -0.453(47) | 0.65 | -0.099(47) | 0.92 |
| | **Seizure severity** | 0.428(27) | 0.67 | -2.59(27) | 0.015 | NA | NA |
| | **NOR** | -0.138(52) | 0.89 | 1.925(52) | 0.06 | 1.142(52) | 0.26 |
| | **OF distance** | -1.24(56) | 0.22 | 2.425(56) | 0.019 | 1.03(56) | 0.31 |
| | **OF center time** | -0.181(56) | 0.86 | -0.408(56) | 0.68 | 0.0131(56) | 0.99 |
| | **OF rearing** | 3.165(56) | 0.0025 | 4.585(56) | <0.0001 | -0.143(56) | 0.89 |
| | **CFC recent** | -0.948(56) | 0.35 | -1.309(56) | 0.2 | -1.36(56) | 0.18 |
| | **CFC remote** | -0.441(55) | 0.66 | -0.887(55) | 0.38 | -1.469(55) | 0.15 |
| R cortex | **Fluorojade-B** | -1.282(54) | 0.21 | -7.895(54) | <0.0001 | -0.162(54) | 0.87 |
| R hippocampus | | -1.303(54) | 0.198 | -6.412(54) | <0.0001 | -1.741(54) | 0.087 |
| L cortex | | -1.092(55) | 0.389 | -9.865(55) | <0.0001 | -0.869(55) | 0.29 |
| L hippocampus | | -1.654(55) | 0.1 | -4.957(55) | <0.0001 | -0.857(55) | 0.39 |

**Table S3. Multiple linear regressions for sex effects in mouse studies.** Multiple linear regression was performed including sex, genotype, and kindling as independent variables for AT8+ levels and tdTomato (tdT)+ levels, and behavioral results. We found that female sex (reference variable) was associated with increased AT8+ levels in the hippocampus ipsilateral (right, R) and contralateral (left, L) to AD-tau injection compared to male mice and that male sex was associated with increased tdT labelling in the R olfactory areas. Females displayed less rearing behavior in the open field assay.

| Dependent variable | Independent variable | | | | | |
|---|---|---|---|---|---|---|
| | Cohort | | Genotype | | Kindling | |
| | F (DF) | p value | F (DF) | p value | F (DF) | p value |
| **NOR** | 1.206 (11, 39) | 0.32 | 1.307 (1, 39) | 0.26 | 1.613 (1, 39) | 0.21 |
| **CFC recent** | 1.232 (11, 43) | 0.30 | 1.422 (1, 43) | 0.24 | 1.286 (1, 43) | 0.26 |
| **CFC remote** | 1.114 (11, 42) | 0.38 | 0.993 (1, 42) | 0.32 | 1.745 (1, 42) | 0.19 |

**Table S4. Multiple linear regressions for cohort effects in memory studies.** Multiple linear regression was performed including cohort, genotype, and kindling as independent variables for performance in novel object recognition (NOR) and CFC. No significant cohort effects were found.

| Brain Region | Dependent variable | Independent variable | | | |
|---|---|---|---|---|---|
| | | Sex | | Seizure history | |
| | | z value (M) | p value | z value (+Sz) | p value |
| Amygdala | Tau pathology score | 0.845 | 0.4 | NA | NA |
| Dentate gyrus | | -0.41 | 0.68 | -0.59 | 0.56 |
| CA/Subiculum | | -1.58 | 0.12 | NA | NA |
| Entorhinal cortex | | NA | NA | NA | NA |
| Middle frontal gyrus | | 1.07 | 0.28 | 1.75 | 0.08 |
| Angular gyrus | | 1.05 | 0.29 | NA | NA |
| SMT gyrus | | 1.22 | 0.22 | 1.13 | 0.26 |
| Cingulate gyrus | | 1.58 | 0.11 | 1.6 | 0.11 |
| Occipital cortex | | 0.32 | 0.75 | 0.27 | 0.79 |
| Striatum | | 0.53 | 0.59 | 0.92 | 0.36 |
| Globus pallidus | | -0.99 | 0.32 | 0.035 | 0.97 |
| Thalamus | | -0.22 | 0.82 | 0.98 | 0.33 |
| Midbrain | | 0.72 | 0.47 | 0.22 | 0.83 |
| Substantia nigra | | 3.03 | 0.0024 | 1.4 | 0.16 |
| Pons | | 1.5 | 0.13 | 0.22 | 0.82 |
| Locus Coeruleus | | 2.09 | 0.036 | 0.45 | 0.65 |
| Medulla | | 0.56 | 0.57 | -1.29 | 0.2 |
| Cerebellum | | -0.45 | 0.65 | NA | NA |

**Table S5. Statistical analyses for sex effects in human studies.** Ordinal linear regressions for regional tau pathology scores with seizure history and sex as independent variables. NA indicates that there were not enough levels of the dependent variable for comparison. Male sex was associated with worsened tau scores in the substantia nigra and locus coeruleus.

| **Antibody** | **Ref (Distributor)** | **Isotype** | **Size (kDa)** |
|---|---|---|---|
| NeuN | ABN78 (EMD Millipore) | Rabbit polyclonal IgG | 48/42 |
| phospho-tau AT8 [Ser202; Thr205] | MN1020 (Millipore) | Mouse monoclonal IgG1κ | 40-80 |

**Table S6. Primary antibody information.**

| ID | Seizure history | Age at death (years) | PMI | Braak stage | Sex | Age of onset (years) | Brain weight (grams) | Ventricular enlargement | Brain atrophy |
|---|---|---|---|---|---|---|---|---|---|
| AD-Sz1 | Absent | 84 | 24 | 6 | Female | 78 | 1204 | Mild | None/Normal |
| AD-Sz2 | Absent | 90+ | 19 | 6 | Male | 76 | 1309 | Severe | Mild |
| AD-Sz3 | Absent | 78 | 6 | 6 | Male | 70 | 1190 | Severe | Severe |
| AD-Sz4 | Absent | 70 | 5 | 6 | Male | 61 | 1256 | Moderate | Moderate |
| AD-Sz5 | Absent | 90+ | 84 | 6 | Female | 70 | 908 | Moderate | Moderate |
| AD-Sz6 | Absent | 79 | 17 | 6 | Male | 72 | 1345 | Moderate | Mild |
| AD-Sz7 | Absent | 90+ | 10 | 6 | Male | 82 | 1366 | None/Normal | Mild |
| AD-Sz8 | Absent | 88 | 20 | | Female | 76 | 1094 | Mild | Moderate |
| AD-Sz9 | Absent | 90+ | 5 | 3 | Female | 89 | 1123 | Mild | Mild |
| AD-Sz10 | Absent | 87 | 19 | 6 | Male | 73 | 1198 | Mild | Mild |
| AD-Sz11 | Absent | 86 | 10 | 6 | Male | 72 | 1278 | Moderate | Mild |
| AD-Sz12 | Absent | 62 | 8 | 6 | Female | 56 | 988 | None/Normal | Mild |
| AD-Sz13 | Absent | 90+ | 17 | | Female | 83 | 1124 | None/Normal | Mild |
| AD-Sz14 | Absent | 79 | 4 | 6 | Female | 69 | 909 | Mild | Moderate |
| AD-Sz15 | Absent | 83 | 4 | 6 | Male | 69 | 1155 | Severe | Severe |
| AD-Sz16 | Absent | 85 | 9 | 5 | Female | 76 | 1014 | Moderate | Moderate |
| AD-Sz17 | Absent | 84 | 21 | 3 | Male | 72 | 1265 | Mild | Mild |
| AD-Sz18 | Absent | 44 | 44 | 6 | Male | 42 | 1510 | None/Normal | None/Normal |
| AD-Sz19 | Absent | 85 | 14 | 5 | Female | 78 | 1138 | None/Normal | Moderate |
| AD-Sz20 | Absent | 63 | 16 | 6 | Male | 54 | 1243 | None/Normal | Mild |
| AD-Sz21 | Absent | 74 | 4 | 6 | Female | 62 | 989 | Moderate | Moderate |
| AD-Sz22 | Absent | 90+ | 48 | 3 | Male | 88 | 1438 | Moderate | Moderate |
| AD-Sz23 | Absent | 84 | 21.5 | 6 | Female | 69 | 1151 | Mild | Mild |
| AD-Sz24 | Absent | 70 | 6 | | Female | 57 | 978 | Severe | Severe |
| AD-Sz25 | Absent | 71 | 4 | 6 | Male | 59 | 1330 | Moderate | Moderate |
| AD-Sz26 | Absent | 63 | 5 | | Female | 53 | 1137 | None/Normal | None/Normal |
| AD-Sz27 | Absent | 84 | 18 | 5 | Male | 77 | 1281 | Moderate | None/Normal |
| AD-Sz28 | Absent | 85 | 20 | | Female | 80 | 1026 | Moderate | Moderate |
| AD-Sz29 | Absent | 78 | 18 | 6 | Male | 68 | 1100 | None/Normal | Moderate |
| AD-Sz30 | Absent | 65 | 8.5 | 6 | Female | 55 | 997 | Severe | Severe |
| AD-Sz31 | Absent | 84 | 18.5 | 6 | Male | 68 | 1185 | Moderate | Severe |
| AD-Sz32 | Absent | 90+ | 24 | 6 | Female | 86 | 938 | Mild | Mild |
| AD-Sz33 | Absent | 90+ | 12 | 6 | Female | 79 | 1269 | Mild | Mild |
| AD-Sz34 | Absent | 74 | 6 | 6 | Female | 64 | 1149 | Mild | Mild |
| AD-Sz35 | Absent | 70 | 16 | 6 | Female | 51 | 845 | Severe | Severe |
| AD-Sz36 | Absent | 64 | 16.5 | 6 | Male | 55 | 1320 | None/Normal | None/Normal |
| AD-Sz37 | Absent | 84 | 7 | 6 | Male | 69 | 1309 | Moderate | Mild |
| AD-Sz38 | Absent | 78 | 6.5 | 6 | Male | 64 | 926 | Severe | Severe |
| AD-Sz39 | Absent | 65 | 17 | 6 | Male | 59 | 960 | Moderate | Severe |

| | | | | | | | | | |
|---|---|---|---|---|---|---|---|---|---|
| AD-Sz40 | Absent | 64 | 20 | 3 | Female | 59 | 1098 | Moderate | Severe |
| AD-Sz41 | Absent | 72 | 8 | 6 | Male | 63 | 1167 | Moderate | Moderate |
| AD-Sz42 | Absent | 68 | 7 | 6 | Male | 54 | 1058 | Moderate | Moderate |
| AD-Sz43 | Absent | 88 | 15 | 5 | Female | 78 | 1145 | Mild | None/Normal |
| AD-Sz44 | Absent | 89 | 12 | 6 | Female | 72 | 1080 | Severe | Moderate |
| AD-Sz45 | Absent | 90+ | 12 | 6 | Female | 76 | 1283 | Moderate | Moderate |
| AD-Sz46 | Absent | 89 | 5 | | Female | 76 | 1060 | Severe | Severe |
| AD-Sz47 | Absent | 85 | 12 | 6 | Female | 70 | 1069 | Moderate | Mild |
| AD-Sz48 | Absent | 90+ | 11 | 6 | Male | 82 | 1459 | Moderate | Moderate |
| AD-Sz49 | Absent | 90+ | 4 | 6 | Male | 84 | 1105 | Moderate | Moderate |
| AD-Sz50 | Absent | 75 | 18.5 | 6 | Male | 61 | 1210 | Severe | Mild |
| AD-Sz51 | Absent | 84 | 5 | | Female | 74 | 1053 | Severe | Severe |
| AD-Sz52 | Absent | 81 | 18 | 6 | Male | 73 | 1427 | Moderate | Mild |
| AD-Sz53 | Absent | 83 | 4.5 | 6 | Male | 66 | 1013 | Moderate | Severe |
| AD-Sz54 | Absent | 84 | 11 | 6 | Female | 78 | 1043 | Moderate | Moderate |
| AD-Sz55 | Absent | 90+ | 20 | 5 | Male | 83 | 1237 | Moderate | Moderate |
| AD-Sz56 | Absent | 81 | 26 | 6 | Female | 65 | 1260 | Moderate | Mild |
| AD-Sz57 | Absent | 64 | 14 | 6 | Female | 53 | 1099 | Mild | Moderate |
| AD-Sz58 | Absent | 61 | 18 | 6 | Male | 53 | 1203 | Severe | Severe |
| AD-Sz59 | Absent | 86 | 6.5 | 6 | Male | 75 | 1147 | Severe | Moderate |
| AD-Sz60 | Absent | 71 | 4.5 | 6 | Male | 62 | 1234 | Mild | Moderate |
| AD-Sz61 | Absent | 85 | 4 | 6 | Female | 66 | 969 | Severe | Severe |
| AD-Sz62 | Absent | 82 | 7 | 6 | Female | 71 | 1113 | Moderate | Moderate |
| AD-Sz63 | Absent | 90+ | 18 | 4 | Female | 75 | 1012 | Moderate | Mild |
| AD-Sz64 | Absent | 86 | 6 | 5 | Female | 76 | 889 | Mild | Moderate |
| AD-Sz65 | Absent | 90+ | 9 | 3 | Male | 86 | 1232 | Mild | Moderate |
| AD-Sz66 | Absent | 79 | 20 | 6 | Male | 72 | 1147 | Moderate | Mild |
| AD-Sz67 | Absent | 85 | 16 | 5 | Male | 65 | 1404 | Severe | Moderate |
| AD-Sz68 | Absent | 83 | 11 | | Female | 66 | 845 | Severe | Severe |
| AD-Sz69 | Absent | 75 | 18 | 6 | Female | 65 | 1207 | Moderate | Severe |
| AD-Sz70 | Absent | 70 | 4 | | Male | 56 | 1201 | Severe | Severe |
| AD-Sz71 | Absent | 74 | 8 | 6 | Male | 59 | 1173 | Moderate | Mild |
| AD-Sz72 | Absent | 90+ | 10 | 6 | Female | 83 | 1214 | Mild | Mild |
| AD-Sz73 | Absent | 85 | 4 | 6 | Female | 61 | 967 | Severe | Severe |
| AD-Sz74 | Absent | 88 | 27 | 5 | Female | 76 | 1205 | Moderate | Mild |
| AD-Sz75 | Absent | 63 | 27 | | Female | 51 | 999 | Severe | Moderate |
| AD-Sz76 | Absent | 83 | 18 | 6 | Male | 73 | 1336 | Moderate | Mild |
| AD-Sz77 | Absent | 89 | 18 | 6 | Female | 74 | 1208 | Moderate | Severe |
| AD-Sz78 | Absent | 88 | 7 | | Female | 72 | 1016 | Severe | Severe |
| AD-Sz79 | Absent | 88 | 17 | 4 | Male | 79 | 1177 | Mild | Mild |
| AD-Sz80 | Absent | 85 | 20 | N/A | Female | 79 | 1202 | Moderate | Moderate |

| | | | | | | | | | |
|---|---|---|---|---|---|---|---|---|---|
| AD-Sz81 | Absent | 70 | 3.5 | 6 | Female | 52 | 1068 | Moderate | Moderate |
| AD-Sz82 | Absent | 86 | 21 | 6 | Female | | 1128 | Mild | Moderate |
| AD-Sz83 | Absent | 75 | 19 | 6 | Male | 65 | 1193 | Moderate | Moderate |
| AD-Sz84 | Absent | 73 | 22 | 6 | Female | 63 | 1022 | Moderate | Moderate |
| AD-Sz85 | Absent | 73 | 20 | 5 | Male | 66 | 1319 | Mild | Mild |
| AD-Sz86 | Absent | 77 | 17 | 6 | Female | 70 | 1102 | Moderate | Mild |
| AD-Sz87 | Absent | 79 | 11 | 5 | Female | 68 | 1049 | Moderate | Moderate |
| AD-Sz88 | Absent | 68 | 20 | 6 | Male | | 1134 | Mild | Mild |
| AD-Sz89 | Absent | 72 | 24 | 6 | Male | 68 | 1176 | Moderate | Moderate |
| AD-Sz90 | Absent | 55 | 37 | 6 | Female | | 868 | | |
| AD-Sz91 | Absent | 75 | 5 | 6 | Male | 64 | 1235 | Severe | Mild |
| AD-Sz92 | Absent | 53 | 14 | 6 | Female | | 1146 | None/Normal | Moderate |
| AD-Sz93 | Absent | 75 | 12 | 6 | Female | 70 | 1140 | Mild | Moderate |
| AD-Sz94 | Absent | 72 | 17 | 5 | Male | 63 | 1388 | Mild | Mild |
| AD-Sz95 | Absent | 57 | 9 | 6 | Male | 52 | 1331 | Mild | Mild |
| AD+Sz1 | Yes | 83 | 7 | 6 | Female | 59 | 783 | Moderate | Severe |
| AD+Sz2 | Yes | 83 | 11 | 6 | Male | 71 | 1077 | Severe | Severe |
| AD+Sz3 | Yes | 81 | 5 | 6 | Female | 71 | 847 | Severe | Moderate |
| AD+Sz4 | Yes | 77 | 3 | 6 | Female | 71 | 1010 | Mild | Moderate |
| AD+Sz5 | Yes | 76 | 45 | 6 | Female | 66 | 1139 | Mild | Mild |
| AD+Sz6 | Yes | 65 | 11 | 6 | Female | 55 | 1125 | Moderate | Moderate |
| AD+Sz7 | Yes | 80 | 12 | 6 | Female | 67 | 1063 | Moderate | Moderate |
| AD+Sz8 | Yes | 68 | 12 | 6 | Female | 56 | 942 | Severe | Severe |
| AD+Sz9 | Yes | 89 | 3 | 6 | Male | 77 | 1034 | Mild | Mild |
| AD+Sz10 | Yes | 71 | 23 | 6 | Male | 65 | 1206 | Mild | Moderate |
| AD+Sz11 | Yes | 68 | 9 | 6 | Female | 60 | 1000 | Mild | Moderate |

**Table S7. Clinical information for human subjects.** PMI=postmortem interval.

| AT8/mm$^2$ ~ Genotype * Treatment + (1 + Genotype * Treatment \| Parent.ID/Region.ID) + (1 \| Ms.ID) | | | |
|---|---|---|---|
| **Major brain region** | **Rhat** | **Bulk ESS Range** | **Tail ESS Range** |
| Isocortex | <1.01 | 1993 - 18922 | 1738 - 10904 |
| Olfactory areas | <1.01 | 2712 - 16265 | 2930 - 11270 |
| Hippocampus | <1.01 | 1007 - 18811 | 898 - 11036 |
| Cortical subplate | <1.01 | 1220 - 19825 | 3209 - 11026 |
| Striatum | <1.01 | 837 - 14707 | 1240 - 11230 |
| Pallidum | <1.01 | 919 - 16733 | 1256 - 11205 |
| Thalamus | <1.01 | 1172 - 14301 | 2007 - 10439 |
| Hypothalamus | <1.01 | 1568 - 14125 | 2111 - 10665 |
| Midbrain | <1.01 | 2141 - 19508 | 2029 - 11177 |
| Fiber tracts | <1.01 | 944 - 21268 | 1268 - 10952 |
| **AT8/mm$^2$ (WT/Sal norm) ~ Genotype * Treatment * region + (1 \| Ms.ID)** | | | |
| **Major brain region** | **Rhat** | **Bulk ESS Range** | **Tail ESS Range** |
| Isocortex | <1.01 | 1579 - 15452 | 3983 - 9145 |
| Olfactory areas | <1.01 | 3187 - 15853 | 5676 - 9886 |
| Hippocampus | <1.01 | 3083 - 18569 | 5592 - 11169 |
| Cortical subplate | <1.01 | 3166 - 13096 | 5307 - 10225 |
| Striatum | <1.01 | 3341 - 16352 | 6174 - 10907 |
| Pallidum | <1.01 | 2840 - 11702 | 5383 - 9888 |
| Thalamus | <1.01 | 1259 - 12518 | 2607 - 8169 |
| Hypothalamus | <1.01 | 1193 - 12079 | 2220 - 8767 |
| Midbrain | <1.01 | 3511 - 19084 | 6321 - 10395 |
| Fiber tracts | <1.01 | 1098 - 12365 | 2501 - 7641 |
| **tdT/mm$^2$ ~ Genotype * Treatment + (1 + Genotype * Treatment \| Parent.ID/Region.ID) + (1 \| Ms.ID)** | | | |
| **Major brain region** | **Rhat** | **Bulk ESS Range** | **Tail ESS Range** |
| Isocortex | <1.01 | 568 - 19395 | 1473 - 11109 |
| Olfactory areas | <1.01 | 2115 - 15291 | 2880 - 11167 |
| Hippocampus | <1.01 | 1291 - 19055 | 2063 - 11061 |
| Cortical subplate | <1.01 | 1299 - 20982 | 2855 - 11230 |
| Striatum | <1.01 | 2685 - 20878 | 3321 - 11448 |
| Pallidum | <1.01 | 2705 - 20670 | 4453 - 12086 |
| Thalamus | <1.01 | 1983 - 21508 | 2432 - 11444 |
| Hypothalamus | <1.01 | 1673 - 27406 | 2893 - 17714 |
| Midbrain | <1.01 | 1756 - 25477 | 3024 - 11631 |
| **tdT/mm$^2$ (WT/Sal norm) ~ Genotype * Treatment * region + (1 \| Ms.ID)** | | | |
| **Major brain region** | **Rhat** | **Bulk ESS Range** | **Tail ESS Range** |
| Isocortex | <1.01 | 404 - 13523 | 622 - 8643 |
| Olfactory areas | <1.01 | 1355 - 9235 | 2334 - 8571 |
| Hippocampus | <1.01 | 4702 - 36338 | 7764 - 13882 |
| Cortical subplate | <1.01 | 2779 - 12546 | 4884 - 8253 |
| Striatum | <1.01 | 1532 - 8901 | 2498 - 7379 |
| Pallidum | <1.01 | 3526 - 23707 | 6144 - 10865 |
| Thalamus | <1.01 | 349 - 13533 | 968 - 11308 |
| Hypothalamus | <1.01 | 1783 - 22733 | 3947 - 17378 |
| Midbrain | <1.01 | 4344 - 22070 | 5982 - 10918 |

**Table S8. Bayesian model diagnostics.** Diagnostics from Bayesian models incorporating parent region/subregion as random effects, or each brain region independently as a fixed effect and genotype and treatment as fixed effects. Ms.ID = subject. Rhat of less than 1.01 indicates strong Markov chain Monte Carlo chain convergence. Bulk effective sample size (ESS) >400 and tail ESS >400 indicate reliable estimates of median and tails of the posterior distribution, respectively.
